# Molecular atlas of the human brain vasculature at the single-cell level

**DOI:** 10.1101/2021.10.18.464715

**Authors:** Thomas Wälchli, Moheb Ghobrial, Marc Schwab, Shigeki Takada, Hang Zhong, Samuel Suntharalingham, Sandra Vetiska, Daymé Rodrigues Gonzalez, Hubert Rehrauer, Ruilin Wu, Kai Yu, Jeroen Bisschop, Fiona Farnhammer, Luca Regli, Karl Schaller, Karl Frei, Troy Ketela, Mark Bernstein, Paul Kongkham, Peter Carmeliet, Taufik Valiante, Peter B. Dirks, Mario L. Suva, Gelareh Zadeh, Viviane Tabar, Ralph Schlapbach, Katrien De Bock, Jason E. Fish, Philippe P. Monnier, Gary D. Bader, Ivan Radovanovic

## Abstract

A broad range of brain pathologies critically relies on the vasculature, and cerebrovascular disease is a leading cause of death worldwide. However, the cellular and molecular architecture of the human brain vasculature remains poorly understood. Here, we performed single-cell RNA sequencing of 599,215 freshly isolated endothelial, perivascular and other tissue-derived cells from 47 fetuses and adult patients to construct a molecular atlas of the developing fetal, adult control and diseased human brain vasculature. We uncover extensive molecular heterogeneity of healthy fetal and adult human brains and across eight vascular-dependent CNS pathologies including brain tumors and brain vascular malformations. We identify alteration of arteriovenous differentiation and reactivated fetal as well as conserved dysregulated pathways in the diseased vasculature. Pathological endothelial cells display a loss of CNS-specific properties and reveal an upregulation of MHC class II molecules, indicating atypical features of CNS endothelial cells. Cell-cell interaction analyses predict numerous endothelial-to-perivascular cell ligand-receptor crosstalk including immune-related and angiogenic pathways, thereby unraveling a central role for the endothelium within brain neurovascular unit signaling networks. Our single-cell brain atlas provides insight into the molecular architecture and heterogeneity of the developing, adult/control and diseased human brain vasculature and serves as a powerful reference for future studies.

## INTRODUCTION

The brain vasculature is important for both the proper functioning of the normal brain as well as for a variety of vascular-dependent CNS pathologies such as brain tumors, brain vascular malformations, stroke and neurodegenerative diseases^1–8^. A better understanding of the underlying cellular and molecular mechanisms and architecture of the brain vasculature during brain development, in the healthy adult brain, as well as in vascular-dependent brain diseases, has broad implications for both the biological understanding as well as the therapeutic targeting of the pathological brain vasculature^9–13^. Vascular growth and network formation, involving endothelial cells (ECs) and other cells of the neurovascular unit (NVU), are highly dynamic during brain development, almost quiescent in the healthy adult brain, and reactivated in a variety of angiogenesis-dependent brain pathologies such as brain tumors and brain vascular malformations^2, 6, 14, 15^, thereby activating ECs and perivascular cells (PVCs) of the NVU and other tissue-derived cells (collectively referred to hereafter as PVCs). However, which molecular signaling cascades are reactivated and how they regulate brain tumor and brain vascular malformation vascularization and growth is largely unknown.

The CNS vasculature has unique features such as the blood-brain barrier (BBB) and the NVU^16–18^. During development, various CNS-specific and general signaling pathways drive CNS angiogenesis^2,196, 17, 20, 21^. The brain vasculature also displays an arteriovenous endothelial hierarchy similar to peripheral vascular beds^22–24^. Developmentally regulated signaling axes in endothelial cells are thought to contribute to the establishment of CNS-specific properties as well as arteriovenous specification of the endothelium in the healthy adult brain and potentially to their alteration in disease^12, 13^. Over the past years, single-cell transcriptome atlases of human peripheral organs^25,26,27^ as well as of mouse brain and peripheral vasculature^22, 28^ were established. Nevertheless, a landscape of the human brain vasculature at the single-cell level has not been achieved until now. Thus, we created a comprehensive molecular atlas of the human brain vasculature using single-cell RNA sequencing (scRNA-seq) analysis in the developing, adult/control, and diseased human brain (Figure 1a,b, Extended Data Figure 1). We discovered extensive heterogeneity among ECs as well as common hallmarks across multiple important brain pathologies, including altered arteriovenous specification and CNS-specificity, upregulation of major histocompatibility complex (MHC) class II signaling and strong EC-EC/EC-PVC communication networks.

**Figure 1.**
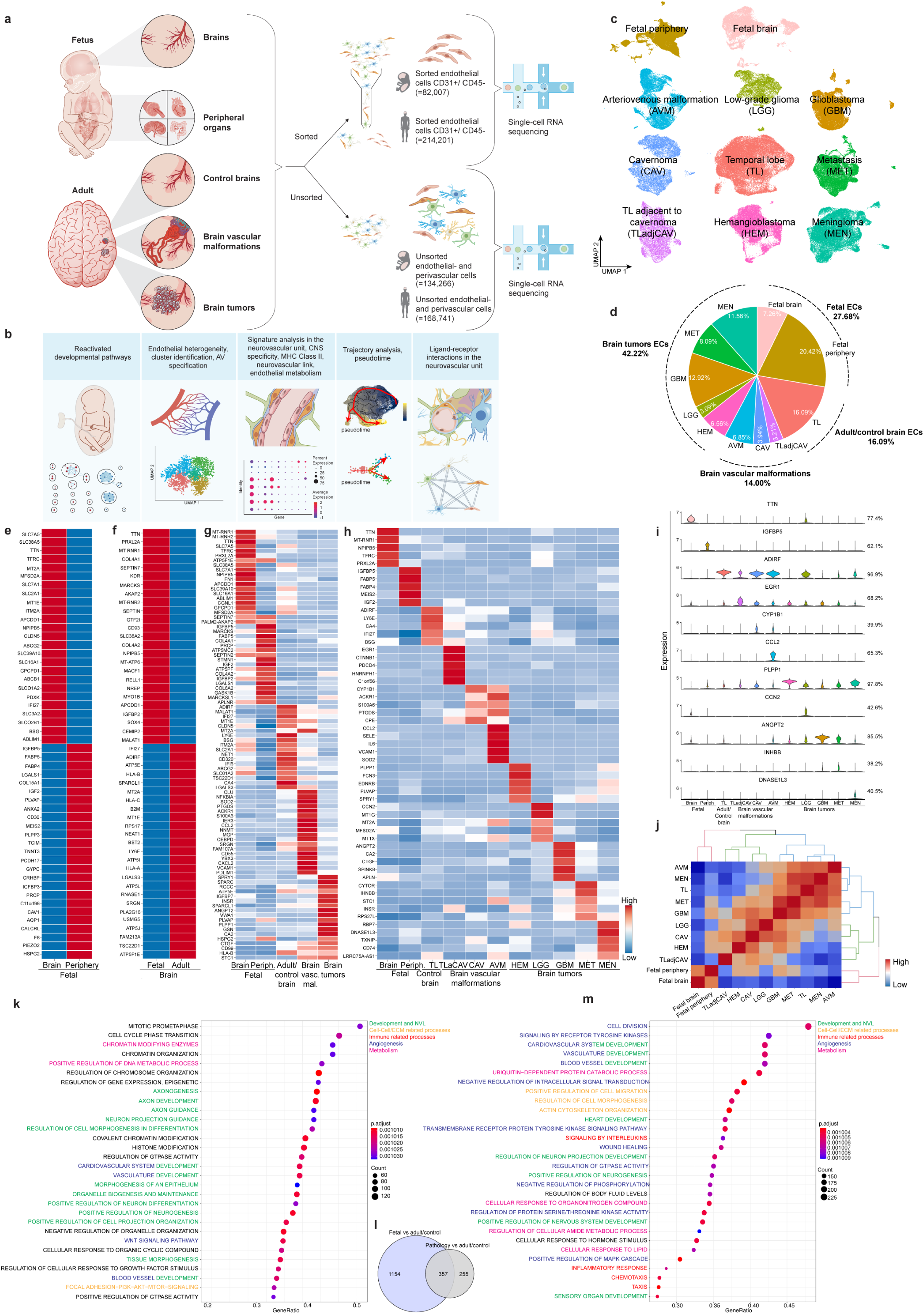
Construction of a molecular single-cell atlas of the human brain vasculature. **a,b,** Scheme of the experimental workflow (**a**) and computational analysis summary (**b**). **c,** UMAP plots of 296,208 sorted and in silico-quality checked (ECs, colored by tissue of origin. **d,** Piechart showing relative abundance and percentage of ECs from each tissue collected. **e-h,** Expression heatmap of the top 25 (**e,f**), the top 18 (**g**), and the top 5 ranking marker genes in the indicated tissues. Color scale: red, high expression; white, intermediate expression; blue, low expression. **i,** Violin plots of the expression of the top marker genes of each tissue type (percentage of cells expressing the marker gene is indicated on the right; in case of marker gene enrichment in multiple tissues, the violin plot with the highest expression is indicated by an asterisk). **j,** Endothelial cells transcriptome correlation heatmap and hierarchical clustering of all tissues. **k,m,** Over-representation (enrichment) analysis shown as dotplots representing the top 30 pathways enriched in fetal brain ECs as compared to adult/control brains (**k**) and in pathological brain ECs over adult/control brain ECs (**m**). Pathway analysis was performed using gene-set enrichment analysis (GSEA). Pathways are color-coded for the biological processes indicated. **l,** Venn diagram showing the overlap between the 1511 significant pathways enriched in fetal brain ECs as compared to adult/control brain ECs and the 612 significant gene sets enriched in pathological brain ECs as compared to adult/control brain ECs.

## RESULTS

### Construction of a molecular single-cell atlas of the human brain vasculature

We constructed a human brain vasculature single-cell atlas using samples from fetal as well as adult control) and diseased brains. The neocortical resection of temporal lobectomy epilepsy surgeries served as adult control brain tissue and diseased brain tissue was collected from various brain vascular malformations and brain tumors (Figure 1a, Extended Data Figure 1, Supplementary Tables 1,2). We acquired freshly isolated fetal cells from 8 individual fetuses (5 fetal brains and peripheral tissue from 7 fetuses) and from 41 adult brain specimens (derived from 39 individual adult patients), covering temporal lobe (TL) controls and cavernoma (CAV), adjacent to cavernoma (TLadjCAV), brain arteriovenous malformation (AVM), hemangioblastoma (HEM), low-grade glioma (LGG), high-grade gliomas/glioblastoma (GBM), lung cancer brain metastasis (MET) and meningioma (MEN) pathologies (Figure 1a, Extended Data Figure 1, Supplementary Tables 1,2).

Brain tissue samples were dissociated into single-cell suspensions, which were either FACS-sorted for ECs (CD31^+^/CD45^-^) or processed as unsorted samples to examine the NVU (Figure 1a). Single-cell transcriptomes were collected using the 10x Genomics Chromium system^29^ and analyzed using a range of computational methods (Figure 1b). CD31^+^/CD45^-^ ECs showed consistent expression of classical endothelial markers, such as *CD31*, *VWF* and *CLDN5*, while not expressing the pericyte markers *PDGFRB* (except for some PDGFRB^+^ GBM ECs, as previously reported^30–32^) and *NG2*, the monocyte markers *CD68* and *CD45,* the astrocyte marker *GFAP*, and the platelet marker *CD61* (Extended Data Figure 2a-n), thereby confirming the purity of our EC isolations. In summary, 599,215 single cells, including 296,208 sorted ECs and 303,007 unsorted ECs and PVCs, passed our quality control criteria considering the number of detected genes, mitochondrial read counts and doublets (Figure 1a, Supplementary Table 2).

To broadly map the neurovasculature and to address the NVU across different brain entities, 44,116 fetal, 47,305 adult/control and 121,436 pathological unsorted EC and PVC transcriptomes from 33 patients were pooled, clustered, annotated using known marker genes and comparison to public datasets^22, 28, 33^ and visualized (Figure 1a, Extended Data Figure 3c,e). We identified 18 major cell types, including all known human brain vascular and perivascular, and other tissue-derived cell types in the human brain (Extended Data Figures 3e-g,4). The detected cell type distributions differed between the fetal, adult control and pathological brain sample types and the detected EC frequency was highest in the hemangioblastoma (Extended Data Figures 3f,4), reflecting its nature as a vascular tumor^34^. Individual cell types clustered together across tissue entities more often than different cell types within entities (Extended Data Figure 3g), indicating conserved cell-specific transcriptomes for each cell type.

We next compared ECs in the sorted samples between fetal brains, fetal peripheral organs, adult temporal lobes and vascular pathologies and found that ECs from different entities exhibited prominent transcriptomic heterogeneity (Figure 1c,d, Supplementary Figure 1). Differential expression and heatmap visualization of top-ranking marker genes revealed distinct gene expression signatures of ECs for each of the 11 entities (Figure 1e-h, Extended Data Figures 5,6, Supplementary Figures 1-13). To define major EC signatures, we compared fetal brain vs. fetal periphery ECs, fetal brain vs. adult/control brain ECs, and adult/control vs. pathological brain ECs. Notably, comparison of fetal brain versus fetal periphery ECs defined a human fetal CNS signature characterizing CNS and periphery-specific markers of the fetal vasculature (Figure 1e, Supplementary Table 3), whereas comparison of the fetal vs. adult brain ECs defined a human fetal/developmental brain signature revealing properties of the developing and mature human brain vasculature (Figure 1f, Supplementary Table 4). Moreover, fetal periphery, fetal brain, adult/control brain (TL), brain vascular malformations (TLadjCAV, CAV, AVM) and brain tumors (HEM, LGG, GBM, MET, MEN) all revealed distinct EC markers, with the exception of some overlapping marker genes between AVMs and CAVs (Figure 1g-i, Supplementary Tables 5,6). Fetal brain and periphery revealed specific markers (e.g. *TTN*, *IGFBP5*). In the adult/control, some EC markers were conserved across two or more entities (e.g. *ADIRF*, *EGR1*, *PLPP1, ANGPT2*), whereas other top marker genes showed more specificity for a single disease entity (e.g. *CCL2*, *CCN2*, and *DNASE1L3* in AVM, LGG and MEN, respectively) (Figure 1h,i, Supplementary Table 6). While most of these markers were expressed by a substantial fraction of ECs within a particular entity (Figure 1h,i), the expression of common markers by ECs from various vascular beds indicates transcriptomic correlations among different entities. Using hierarchical clustering to address correlations of ECs revealed that ECs from certain tissues (fetal brain and fetal periphery; TLadjCAV and HEM; CAV and LGG; intraaxial tumors (GBM) and MET; extraaxial tumor MEN and AVM) clustered together (Figure 1j), suggesting partially overlapping transcriptome signatures.

We next assessed differences in ECs between fetal, adult/control and diseased human brains to investigate how ECs are affected by developmental stage and in pathological conditions (Figure 1k-m, Extended Data Figures 5,6). We performed differential gene expression analysis followed by pathway analysis using gene-set enrichment analysis (GSEA) on the ranked gene lists. DEGs between the fetal and adult stage and between the adult/control and pathological brain showed developmental and pathology-specific pathway enrichments (Figure 1k,m, Extended Data Figures 5,6, Supplementary Figures 1-13), providing insight into functional specialization of the human brain vasculature across development, homeostasis and disease. We identified numerous DEGs across the entities, namely 764 in fetal brain vs. fetal periphery, 780 in fetal vs. adult/control, 286 in adult/control vs. pathology, 223 in brain vascular malformations vs. adult/control, 321 in brain tumors vs. adult/control, highlighting heterogeneity within the EC compartments across developmental stages and pathological conditions (Extended Data Figures 5,6).

The top regulated pathways in both fetal vs. adult/control as well as in pathological vs. adult/control brain EC signatures included development and neurovascular link (NVL)^2^, cell-cell/extracellular matrix (ECM)-related processes, immune-related processes, angiogenesis and metabolism (Figure 1k,m, Extended Data Figures 5,6). Notably, of the 612 differentially regulated pathways in pathology vs. adult/control, more than half (357) also showed differential regulation in fetal vs. adult/control brain ECs (Figure 1l, Extended Data Figure 6d), highlighting the importance of developmental pathways in vascular-dependent brain pathologies. Together, these data indicate that signaling axes driving vascular growth during fetal brain development are silenced in the adult control brain and (re)activated in the vasculature of brain tumors and brain vascular malformations.

### Inter-tissue heterogeneity of brain vascular endothelial cells

Next, to further address EC heterogeneity across different brain entities at the single-cell level, we pooled, batch-corrected^35–38^, clustered and visualized all fetal (21,512), adult/control (47,652) and pathological (166,549) sorted brain EC transcriptomes from 38 patients (Figure 2a-d, Extended Data Figure 7a-e). Brain vascular ECs are organized along the human brain arteriovenous axis, referred to as zonation^22, 39–44^. To address arteriovenous zonation, endothelial clusters were biologically annotated using top-ranking marker genes/DEGs in fetal, adult and pathological ECs, and we identified 44 clusters (Figure 2e,f, Extended Data Figure 7a-e,h, Supplementary Table 7) that we then grouped into 14 major EC subtypes for further downstream analysis (Figure 2a-d).

**Figure 2.**
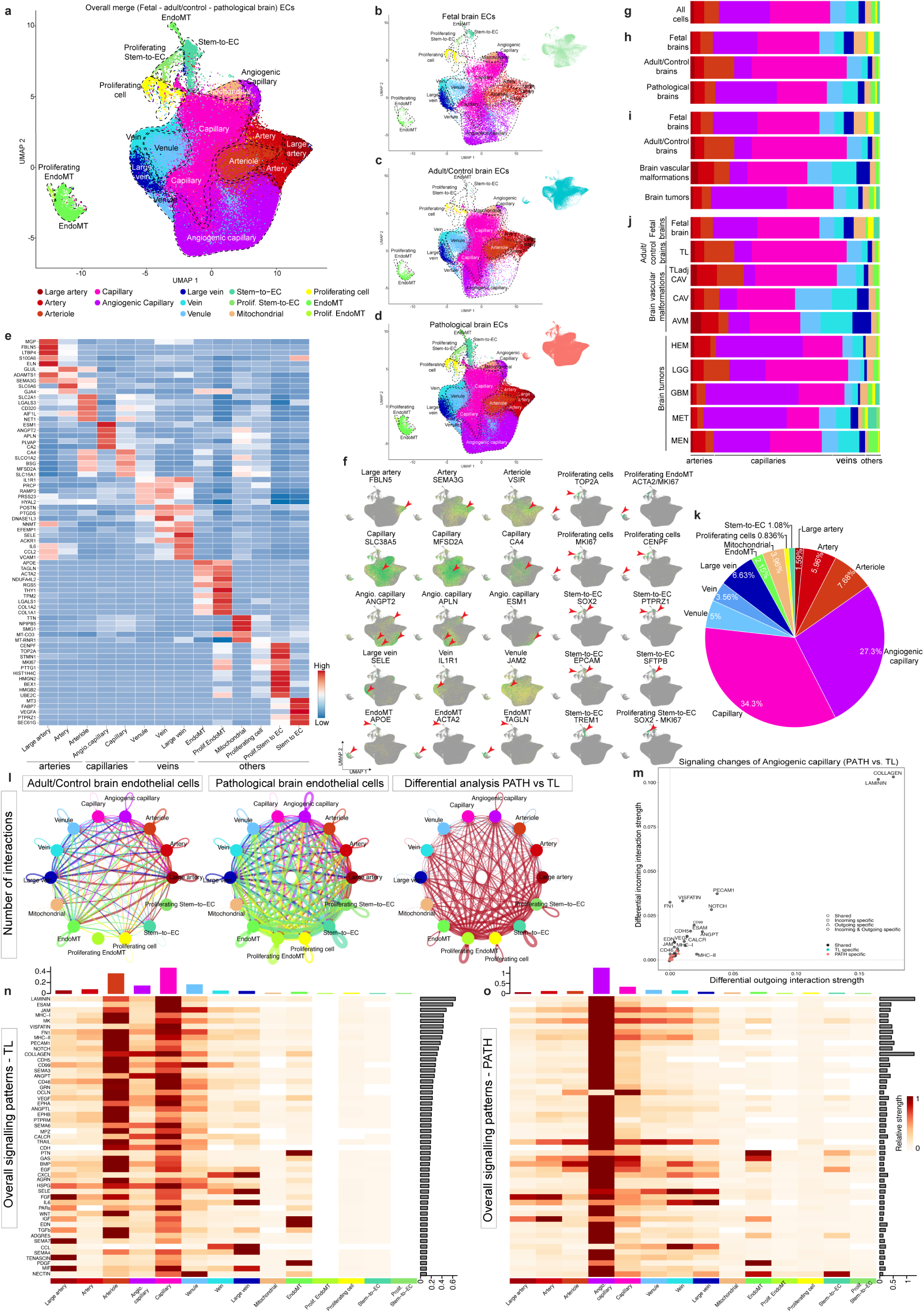
Inter-tissue heterogeneity of brain vascular endothelial cells. **a,** UMAP plot of the 296,208 batch corrected ECs, color-coded by ECs arteriovenous (AV) specification. **b-d,** UMAP plot shown in (**a**) split by tissue of origin: fetal brain (**b**), adult/control brain (**c**) and brain pathologies (**d**). **e,** Heatmap of the top 5 ranking marker gene expression levels in different EC subtypes. Color scale: red, high expression; white, intermediate expression; blue, low expression. **f,** UMAPs plots, color-coded for expression of indicated marker genes (red arrowheads). **g-j,** Relative abundance of EC subtypes (AV specification cluster) from the indicated tissue of origin. Color-code corresponds to (**a**). **k,** Piechart showing the relative abundance of each EC subtype according to AV specification. **l,** Circle plot showing the number of statistically significant ligand-receptor interactions between ECs subtypes in adult/control brain (left panel) and pathological brain ECs (middle panel). Differential analysis of the intercellular signaling interactions (right panel, pathology over control), red indicating upregulation, while blue indicating downregulation. **m,** Scatter plot showing the differential incoming and outgoing interaction strength of pathways in angiogenic capillaries - identifying signaling changes in those cells in pathological as compared to control conditions. **n,o,** Heatmap showing overall signaling patterns of different EC subtypes in adult/control (**n**) and pathological (**o**) brains.

Known vessel segments were identified by established arterio-venous zonation markers^22, 45^, with arterial and venous clusters located at opposite ends of the UMAP, separated by major capillary clusters (Figure 2a-d,f, Extended Data Figures 7a-e,h, Supplementary Tables 7,8). We also assigned EC clusters outside arteriovenous zonation, notably (proliferating) stem-to-endothelial cell transdifferentiating (stem-to-EC) clusters, (proliferating) endothelial-to-mesenchymal transition (EndoMT) clusters (Figure 2f, Extended Data Figure 7h). Multiple clusters showed substantial multi-tissue contributions from the fetal, adult/control and pathological entities, whereas other clusters were either mostly/predominantly pathological or mostly fetal and pathological (i.e. no notable contribution of control/adult brain), namely multiple angiogenic capillary clusters, (proliferating) stem-to-EC clusters, (proliferating) EndoMT clusters, and proliferating cells (Extended Data Figure 7f,g).

Heatmap visualization revealed distinct markers of EC clusters along arteriovenous zonation and confirmed robust differential expression of known marker genes of arteriovenous specification^22, 28^ (Figure 2e,f, Extended Data Figure 7h, Supplementary Table 7,8). Notably, several of these top marker genes have not previously been identified as markers for AV-zonation in the human brain: *LTBP4* (large arteries), *ADAMTS1* (arteries), *VSIR*, *AIF1L*, *CD320*, and others (arterioles), *SLC38A5*, *BSG*, *SLC16A1*, *SLCO1A2* (capillaries), *JAM2*, *PRCP*, *PRSS23*, *RAMP3* (venules), *PTGDS*, *POSTN*, *DNASE1L3* (veins), *CCL2* (large veins) and *PLVAP* and *CA2* (angiogenic capillaries) (Figure 2e,f, Extended Data Figure 7h, Supplementary Table 8). We confirmed high *PLVAP* expression in diseased brain entities (brain tumors>brain vascular malformations) and a slight elevation in the fetal brain (Extended Data Figure 8a-m), indicating its role in developmental and pathological vascular growth^46, 47^. Indeed, *PLVAP* exhibited RNA and protein expression in human brain vascular malformation/tumor ECs by RNA scope and immunofluorescence (Extended Data Figure 8n-n‘).

We found that arteriolar ECs expressed both arterial and capillary markers whereas venular ECs showed co-expression of venous and capillary markers (Figure 2f, Extended Data Figure 7h, Supplementary Table 8)^28^, in line with their topographical location along the AV-vascular tree. We identified specific molecular markers defining EC clusters outside the arteriovenous zone. We found proliferating ECs (e.g. *TOP2A*, *MKI67*) in the fetal, adult (surprisingly, 0.63%) and pathological brains (Figure 2e,f, Extended Data Figure 7h, Supplementary Tables 7,8). We identified EndoMT clusters expressing both mesenchymal (e.g. *APOE*, *ACTA2*, *TAGLN*) and endothelial markers (Extended Data Figure 9m-x)^48^. Notably, we observed two subsets of EndoMT ECs (proliferating EndoMT and EndoMT) in all entities, but increased in pathologies. Proliferating EndoMT ECs expressed both EndoMT and proliferation markers (e.g. *ACTA2*, *MKI67*) (Figure 2e,f, Extended Data Figure 7h, Supplementary Tables 7,8).

In GBMs and METs, we observed stem-to-EC clusters that expressed classical EC markers (e.g. *CD31*, *CLDN5*, *CDH5*, *VWF*, to a lower level compared to other EC clusters) as well as some markers of (tumor) stem cells (Figure 2e,f, Extended Data Figures 7h,10), suggesting that these ECs undergo stem-to-EC transdifferentiation. In GBM, we identified a stem-to-EC cluster expressing GBM stemness markers *SOX2*, *PTPRZ1*, *POUR3F2* and *OLIG1*^49, 50^ in addition to classical EC markers (Extended Data Figures 7h,10a-i)^51, 52^. In METs, we noted a previously undescribed stem-to-EC population that co-expressed EC markers and stem cell markers of lung cancers (e.g. *SOX2*, *EPCAM*, *CD44*, *SFTPB*)^53^ (Extended Data Figures 7h,10n-v). In GBM and METs, we identified groups of stem-to-ECs that co-expressed stemness (e.g. *SOX2, PTPRZ1, EPCAM1* and *SFTPB*) and proliferation markers (e.g. *MKI67*, *BEX1*, *HMGB2* and *UBE2C*) (Figure 2e,f, Extended Data Figure 7h).

To validate the stem-to-EC clusters in GBM and MET, we used double immunostaining for EC and stemness markers. In GBM, we found SOX2^+^/CD31^+^ and PTPRZ1^+^/CD31^+^ co-expressing ECs, whereas in MET, we observed EPCAM^+^/CD31^+^ and SFTBP^+^/CD31^+^ co-expressing ECs (Extended Data Figure 10j-m,w-z). The confirmation of tumor stemness marker enrichment in a subset of tumor ECs indeed suggests the presence of stem-to-ECs in GBM and MET vasculature.

We next addressed the distributions of EC clusters between the fetal, adult/control and pathological brains. Notably, capillaries accounted for ∼61.6% of ECs, arterial ECs for 15.23%, and venous ECs for 15.19%, in agreement with^2, 14^. Stem-to-EC clusters accounted for 1.92%, EndoMT clusters for 2.1% (Figure 2g,k, Extended Data Figure 11). Angiogenic capillaries were more prominent in the fetal and pathological brains (Figure 2g-k, Extended Data Figures 11,9a-l), indicating the angiogenic capacity of the human brain vasculature in development and disease^2, 12, 16, 54^. We further uncovered previously unrecognized EC heterogeneity across a wide range of human brain tissues (Figure 2i,j, Extended Data Figure 11). Angiogenic capillary proportions were higher in the fetal brain and brain tumors (GBM>MET∼HEM>MEN>LGG) (Figure 2i,j, Extended Data Figure 11). Brain vascular malformations revealed elevated proportions of venous clusters (AVM∼CAV), indicating the venous character of these brain vascular malformations^55, 56^.

Because EC clusters reside in close proximity to each other along the arterio-venous tree, we next inferred cell-cell communication pathways^57, 58^ (Figure 2l-o, Extended Data Figures 12-17, Supplementary Figures 14-19). Differential analysis revealed increased cellular crosstalk among EC clusters in pathological ECs, highlighting a key role of angiogenic capillaries (Figure 2l-o, Extended Data Figures 12-17). Angiogenic capillaries displayed upregulation of several incoming and outgoing signaling pathways including cell-cell/ECM-related processes (COLLAGEN, LAMININ in pathological and fetal brains), immune-related processes (IL6 and CCL in pathologies and fetal brains, MHC class II in pathological but not fetal brains), development and NVL (SEMA6A, EPHA/EPHB in pathological and fetal brains), angiogenesis (VEGF, NOTCH in pathological and fetal brains), and metabolism (ACTIVIN, VISFATIN in pathological and fetal brains) in both the diseased vs. adult/control brain and fetal vs. adult/control brain (Figure 2m-o, Extended Data Figures 12-17, Supplementary Figures 14-17), highlighting the angiogenic capillary EC cluster as a major signaling mediator within brain EC networks.

### Alteration of AV-specification in pathological brain vascular endothelial cells

Failure of proper arteriovenous specification (predominantly of the capillary bed) in brain vascular malformations as well as formation of dilated and tortuous arteries and veins in brain tumors has been reported^54, 60^, but alteration of AV-specification in brain pathologies and during fetal (brain) development remains poorly understood. We therefore ordered ECs along a one-dimensional transcriptional gradient using Monocle^61^ to examine the arteriovenous axis in the different entities (Figure 3a-c,e-g,i-k, Extended Data Figures 18a-c,e-g,i-k,m-o,19a-c,e-g,i-k,m-o,q-s,u-w). Whereas arterial and venous markers peaked at opposite ends, capillary markers showed peaks throughout all entities (Extended Data Figure 20, Figure 3b,f,j, Extended Data Figures 18b,f,j,n,19b,f,j,n,r,v), indicating that in silico computational pseudospace recapitulates the in vivo anatomical topography of EC clusters in the human brain vasculature^22, 28^. We observed AV-zonation throughout the fetal, control and pathological brains, but observed a partial alteration of endothelial cell ordering along the AV-axis in disease, in both brain tumors (to different degrees e.g. GBM>MET>HEM>LGG>MEN) and brain vascular malformations (shift towards the venous compartments observed in both AVM and CAV) (Figure 3b,c,f,g,j,k, Extended Data Figures 18b,c,f,g,j,k,n,o,19b,c,f,g,j,k,n,o,r,s,v,w).

**Figure 3.**
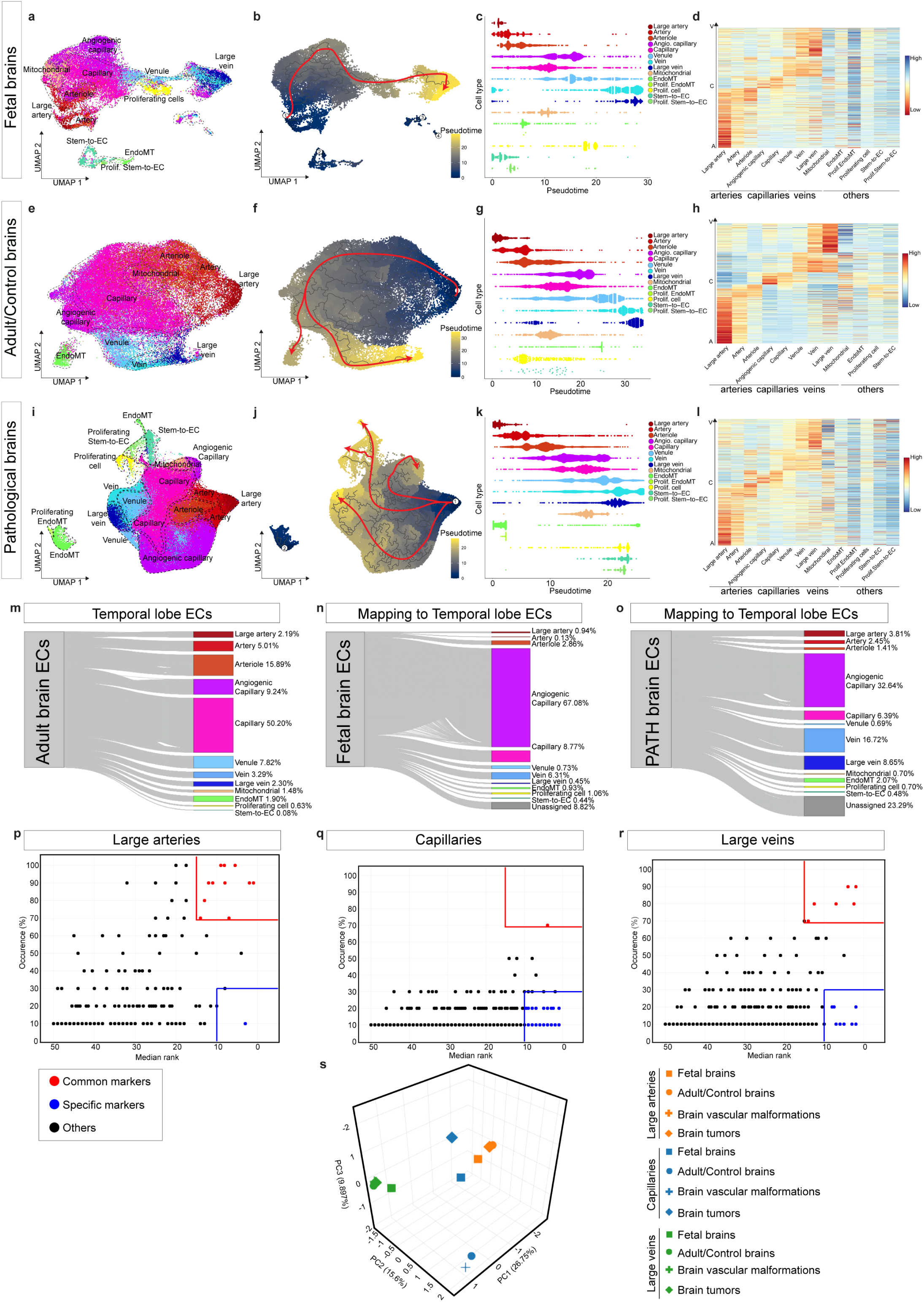
Alteration of AV-specification in brain vascular endothelial cells. **a,b,e,f,i,j,** UMAP plots of human brain ECs isolated from fetal brains (**a,b**), adult/control brains (**e,f**) and pathological brains (**i,j**), colored by AV specification (**a,e,i**) and by pseudotime (**b,f,j**). **c,g,k,** Pseudotime order of ECs color-coded according to AV specification from fetal brains (**c**), control adult/control brains (**g**), and pathological brains (**k**). **d,h,l,** Heatmap of adult/control brain ECs AV specification signature gene expression in human brain ECs isolated from fetal brains (**d**), adult/control brains (**h**), and pathological brains (**l**). **m,n,o,** Sankey plot showing the relative abundance of EC phenotypes in TL (**m**), predicted annotation of fetal ECs (**n**) and brain pathologies ECs (**o**) ECs as mapped to TL ECs. **p,q,r,** Dotplots of common and tissue-specific markers in ECs from large arteries (p), capillaries (**q**) and large veins (**r**) in different tissue types (fetal brain, adult control brain, brain vascular malformations and brain tumors). Red boxes highlight conserved markers between ECs from different tissues; blue boxes highlight tissue-specific markers dots are colored as in the legend. **s,** Three-dimensional PCA visualization of pairwise Jaccard similarity coefficients between indicated ECs from the different tissues.

We defined an AV-signature comprising 1,021 genes revealing significant expression gradients along the arteriovenous axis. Ordering of ECs according to this AV-signature resulted in gradually changing gene expression patterns along the AV-tree (Figure 3d,h,l, Extended Data Figures 18d,h,l,p,19d,h,l,p,t,x). The seamless zonation continuum was recapitulated in all entities but again showed alteration in pathologies. Whereas AV-markers revealed clear distinction between the AV-compartments in the fetal and adult/control brain, notably showing specific markers of large arteries (e.g. *VEGFC*, *FBLN5*), arterioles (e.g. *LGALS3*, *AIF1L*), capillaries (e.g. *SLC35A5*, *MFSD2A*), angiogenic capillaries (e.g. *ESM1*, *ANGPT2*,), venules (e.g. *JAM2*, *PRCP*) and large veins (e.g. *SELE*, *SELP*), some zonation markers showed a less specific/more widespread presence across AV-clusters in pathologies: *JAM2* and *PRCP* were expressed in both capillaries and venules, *LGALS3* and *AIF1L* showed expression in both capillaries and arterioles, indicating partial alteration of AV-specification (Figure 2,3d,h,l, Extended Data Figures 18d,h,l,p,19d,h,l,p,t,x).

We next explored how fetal and pathological ECs map to temporal lobe ECs^35^. Whereas almost all fetal and pathological ECs could be assigned to temporal lobe EC clusters, some ECs were “unassigned”, indicating alteration of AV-specification in fetal and pathological EC transcriptomes. Notably, these “unassigned” ECs mainly belonged to small caliber vessels (Figure 3m-o, Extended Data Figure 21a-j).

We next addressed whether EC markers of AV-clusters were conserved between vascular beds of fetal, adult and pathological brains or expressed in a more tissue-specific manner^28^. Whereas we identified multiple conserved markers for large arteries and large veins, capillaries were more tissue/entity-specific, indicating a more pronounced transcriptional heterogeneity of the capillary bed across the different brain tissues (Figure 3p-s)^28^. Accordingly, capillaries showed more tissue-specific markers than large caliber vessels (Figure 3p-r), indicating a higher susceptibility of capillary ECs to the local tissue microenvironment to respond to tissue-specific requirements.

### Alteration of CNS-specificity in pathological brain vascular endothelial cells

We next examined CNS-specific properties distinguishing brain ECs from ECs outside the CNS^2, 4^ across brain development, adulthood and in disease. Bulk RNA-sequencing in the mouse previously revealed a BBB-enriched transcriptome^62^, but how human brain EC CNS properties differ at the single cell level and whether they are heterogeneous across developmental stages and in disease remains largely unknown.

To address molecular differences of CNS and peripheral ECs at the single-cell level, we compared transcriptomes between human brain and peripheral ECs from computationally extracted ECs of adult human tissues^25^ as well as in our sorted fetal EC dataset (Extended Data Figure 22, Supplementary Table 9). We defined a human adult and fetal CNS signature comprising the top 50 genes enriched in brain ECs compared with ECs of peripheral organs (Figure 1a, Extended Data Figure 23a-f, Supplementary Table 8). These include known BBB markers *MFSD2A*^63^ and *CLDN5*^64^ and known capillary markers *CA4*^22, 61^ and *SPOCK2*^41^, as well as novel, previously unknown genes enriched in the CNS vasculature such as *SPOCK3*, *BSG*, *CD320* and others (Extended Data Figure 22b,e, Supplementary Table 9).

Comparison of the CNS-specific properties between fetal, adult and pathological ECs revealed CNS properties in the fetal and adult brains as well as an alteration of the CNS signature and an acquisition of the peripheral signature in pathologies (Figure 4a-f, Extended Data Figures 23-25). When comparing the CNS signature between pathological and adult/control brain ECs, we found downregulation of *SLC2A1* which is known to be dysregulated in neurodegenerative conditions^44^, as well as of the lipid transporter *MFSD2A*, which is expressed in brain ECs and restricts caveolae-mediated transcytosis at the BBB^63, 65, 66^, and the BBB marker *CLDN5* (Figure 4d-f, Supplementary Table 9), thus suggesting BBB alteration in pathologies^16^. Comparing CNS and peripheral signature expression across entities revealed that the CNS signature was highest in the TL, followed by intra-axial primary brain tumors and fetal brain (LGG>fetal brain>GBM), brain vascular malformations (TLadjCAV>CAV>AVM), intra-axial secondary brain tumor MET, extra-axial brain tumor MEN and the intra-axial primary brain tumor HEM, whereas the peripheral signature followed an inversed pattern (Figure 4e,f, Extended Data Figure 25e,f).

**Figure 4.**
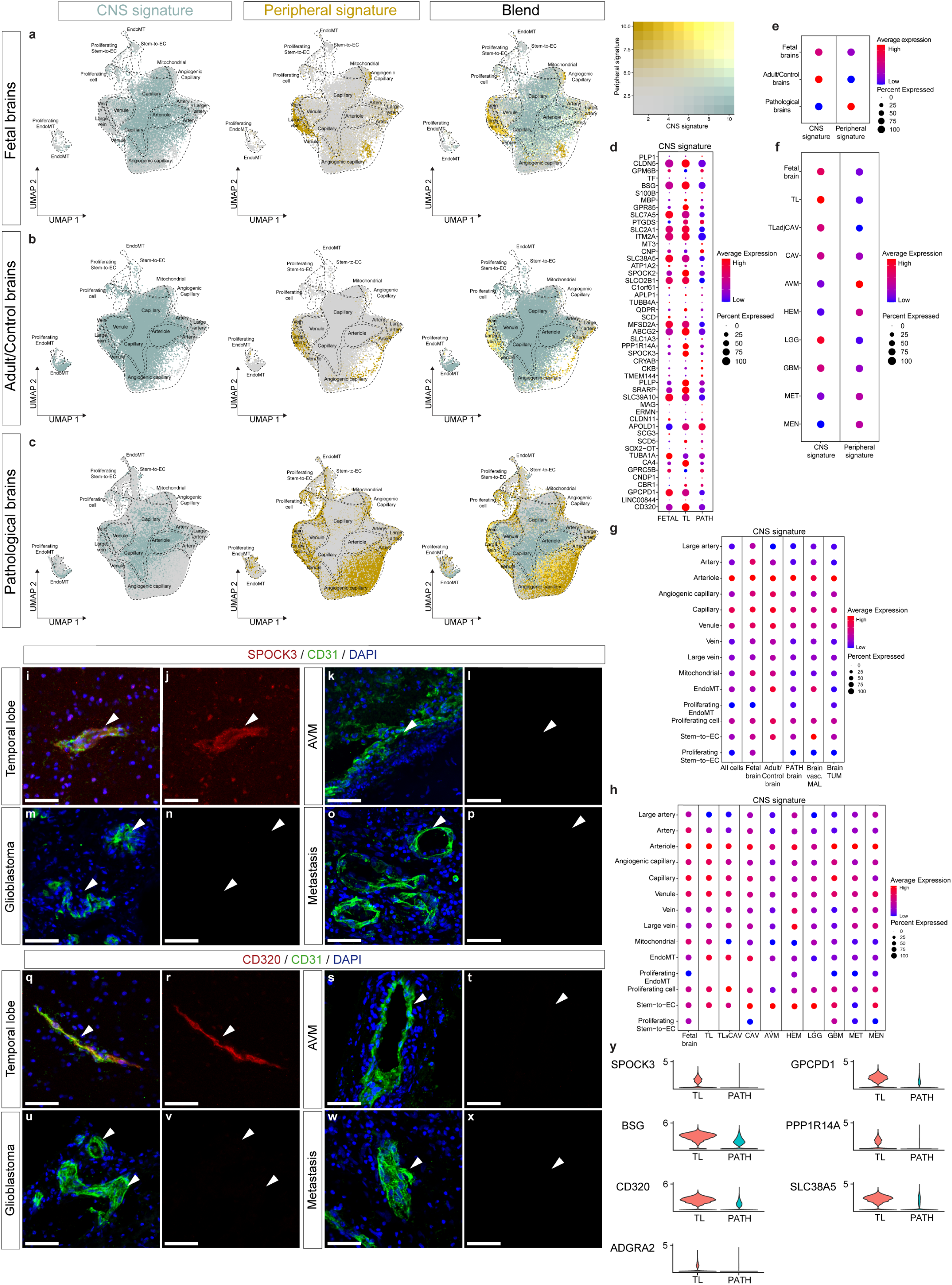
Alteration of CNS-specificity in pathological brain vascular endothelial cells. **a-c**, UMAP plots of the ECs from fetal brains (**a**), adult/control brains (**b**), and pathological brains (**c**). Plots are color-coded for CNS signature (green, left panel), peripheral signature (yellow, middle panel) and a blend of both signatures (right panel). **d,** Dotplot heatmaps of CNS signature genes expression in fetal brain, adult/control brains (temporal lobes) and pathological brain ECs. **e,f,** Dotplot heatmaps of the CNS and peripheral signature expression in fetal brain, adult/control brain, and pathological brain ECs (**e**) and in each individual entity (**f**). **g,h,** Dotplot heatmaps of CNS signature at the level of AV specification for the indicated entities. Color scale: red, high expression; blue, low expression, whereas the dot size represents the percentage expression within the indicated entity. **i**-**x,** Immunofluorescence (IF) images for the protein expression of SPOCK3 and CD320, in temporal lobe (**i,j** and **q,r**), in arteriovenous malformations (**k,l** and **s,t**), in glioblastoma (**m,n** and **u,v**) and in metastasis (**o,p** and **w,x**). Arrowheads identify blood vessels in the different tissues. Scale bars = 50μm. **y,** Violin plots showing the expression of representative CNS specific marker genes for adult/control brain vs. pathological brain ECs.

We next addressed CNS-specific properties of ECs along the arteriovenous axis. In the fetal and adult brain, the CNS signature was mainly expressed by small caliber vessels, while the peripheral signature was predominantly present in large arteries and large veins (Figure 4g, Extended Data Figure 22h). We observed a similar pattern in pathological brains with, however, a notable decrease of cells expressing the CNS signature (most pronounced for angiogenic capillaries>capillaries), paralleled by an increase of cells expressing the peripheral signature predominantly for angiogenic capillaries and large caliber vessels (Figure 4g, Extended Data Figure 22h).

To compare the changes in gene expression in each disease, we determined how the CNS and peripheral signatures responded along the AV-axis in each pathological entity. The CNS signature was downregulated in every pathology with a similar pattern as described above (and reaching highest baseline values of CNS-specificity at the capillary and arteriole levels, with the capillaries being the cluster mostly affected by pathologies) (Figure 4h, Extended Data Figure 22h and^28^), likely pertaining to the influence of the local microenvironment for small caliber vessels. The peripheral signature was upregulated in disease, peaking for AVM, followed by high expression for HEM>MEN>MET>GBM and lower expression for LGG>TLadjCAV, predominantly affecting large caliber vessels and angiogenic capillaries (Figure 4h, Extended Data Figures 22h,23-25). These data indicate that CNS ECs acquire CNS-specific properties during fetal-to-adult transition and take on a peripheral EC signature in disease conditions.

The CNS signature is tightly linked to a functional BBB in vivo^2^. We next investigated the BBB dysfunction module that appears to be upregulated in CNS ECs upon various disease triggers (stroke, multiple sclerosis, traumatic brain injury and seizure) in the mouse brain and that shifts CNS ECs into peripheral EC-like states under these conditions^62^. We found the BBB dysfunction module to be upregulated in brain tumors and brain vascular malformations as well as in the fetal brain (Extended Data Figures 26,28), probably due to pathways related to BBB dysfunction (see methods)^62^. The BBB dysfunction module was highest in AVM, followed by GBM>MET>HEM>CAV>MEN and TLadjCAV (Extended Data Figures 26,28). The expression pattern of these BBB dysfunction module genes along the AV-axis revealed enrichment in large caliber vessels and angiogenic capillaries mimicking the peripheral signature expression pattern, therefore indicating that in brain tumors and brain vascular malformations, CNS ECs take on a “peripheral” endothelial gene expression pattern (Extended Data Figure 28)^62^.

We validated the alteration of transcriptional expression of several CNS signature genes using immunofluorescence. We confirmed decreased expression of *SPOCK3*, *BSG*, *CD320*, *PPP1R14A* and *SLC38A5* in all brain tumors and brain vascular malformations (Figure 4i-y, Extended Data Figures 29-31), thereby highlighting the alteration of CNS properties in the diseased human cerebrovasculature.

### Upregulation of MHC class II receptors in pathological brain vascular endothelial cells

During our analysis of the human brain vasculature, we identified EC populations expressing the MHC class II genes *CD74*, *HLA-DRB5*, *HLA-DMA*, *HLA-DPA1* and *HLA-DRA* in various pathological CNS tissues (and to low levels in adult/control brain), including brain tumors and brain vascular malformations (Extended Data Figures 32-34). This antigen-presenting signature, indicating a possible immune function of ECs of the human brain, prompted us to investigate the heterogeneity of MHC class II transcripts between tissues at the single-cell level.

Recently, genome-wide expression profiling by scRNA-seq has identified endothelial MHC class II expression in a variety of peripheral human and mouse tissues^25, 33^, but assessment of MHC class II expression in developing and aberrant human brain vascular beds at the single-cell level is currently lacking. To assess the molecular heterogeneity of MHC class II gene expression across development and disease, we defined a human MHC class II signature including all known MHC class II genes (Figure 5d, Supplementary Table 10). We observed an upregulated MHC class II signature in pathologies as compared to TL, while the transcriptional MHC class II signature levels in the fetal brain were low (Figure 5a-f, Extended Data Figure 32), in agreement with the recently reported low MHC class II expression in human fetal organs^25^. In brain pathologies, we found that the MHC class II signature was highest in brain vascular malformations (CAV>AVM), the extra-axial brain tumor MEN, the intra-axial secondary brain tumor MET, followed by the intra-axial primary brain tumors HEM>LGG>GBM, TL and TLadjCAV and the fetal brain, grossly following the peripheral signature expression gradient whereas the CNS signature showed an inverse trend (Figure 5f, Extended Data Figures 32-34). We examined MHC class II signature expression patterns according to arteriovenous zonation. Whereas in the fetal brain, only very few ECs (large arteries and arterioles) expressed the MHC class II signature, mainly large caliber vessel (large arteries and large veins) ECs expressed a signature of genes involved in MHC-II-mediated antigen presentation in the adult brain (Figure 5a,b,g,h).

**Figure 5.**
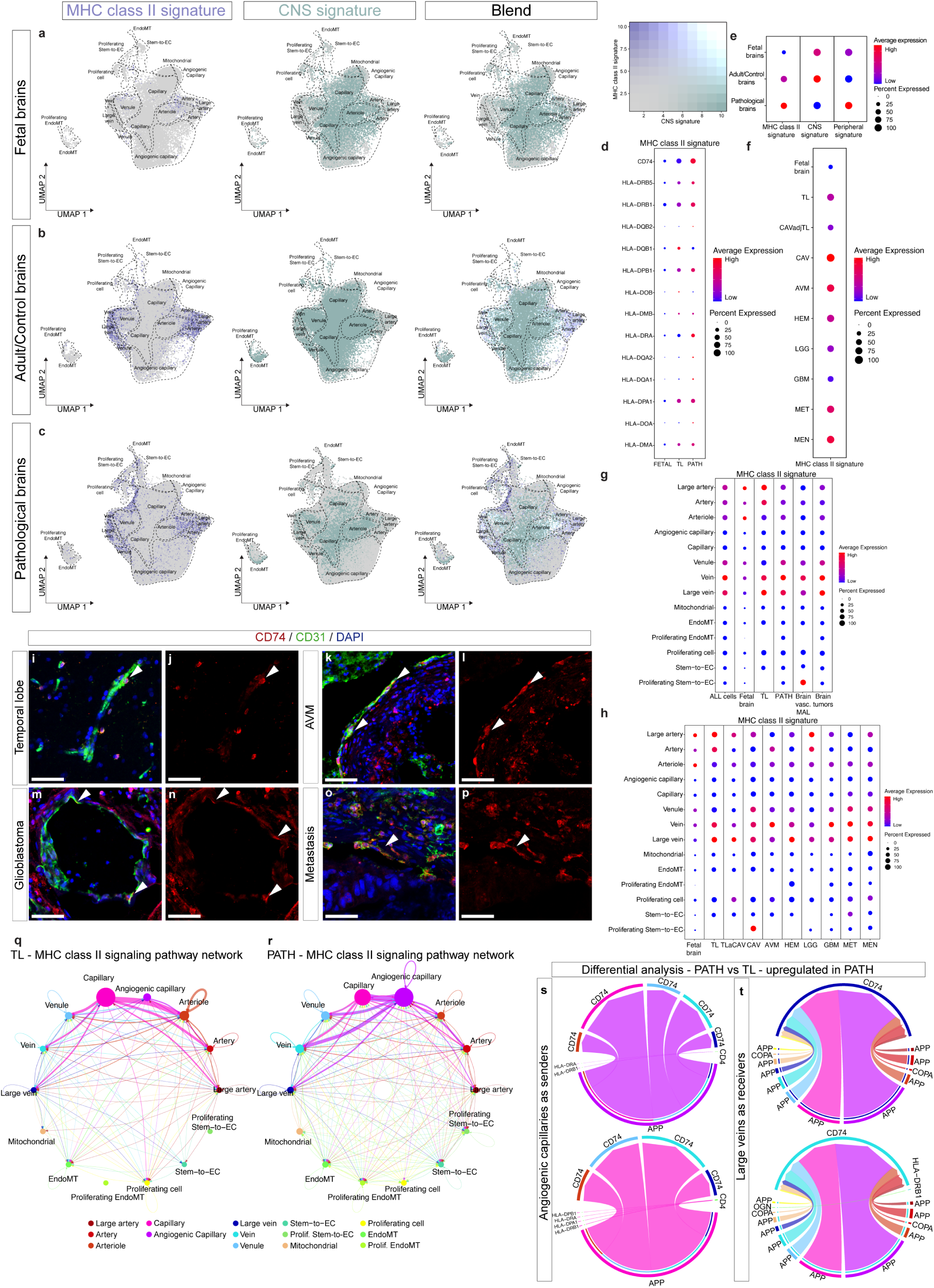
Upregulation of MHC class II receptors in pathological brain vascular endothelial cells. **a-c**, UMAP plots of ECs from fetal brains (**a**), adult/control brains (**b**), and pathological brains (**c**). Plots are color-coded for MHC class II signature (violet, left panel), CNS signature (green, middle panel), and blend of both signatures (right panel). **d,** Dotplot heatmaps of MHC class II signature genes expression in fetal brain, adult/control brains (temporal lobes) and pathological brain ECs. **e,f,** Dotplot heatmaps of the MHC class II, CNS and peripheral signatures expression in fetal brain, adult/control brain, and pathological brain ECs (**e**) and MHC class II signature expression in each individual entity (**f**). **g,h,** Dotplot heatmaps of MHC class II signature at the level of AV specification for the indicated entities. Color scale: red, high expression; blue, low expression, whereas the dot size represents the percentage expression within the indicated entity. **i-p,** Immunofluorescence (IF) images for the protein expression of CD74 in temporal lobe (**i,j**), in arteriovenous malformations (**k,l**), in glioblastoma (**m,n**) and in metastasis (**o,p**). Arrowheads identify blood vessels in the indicated tissues. Scale bars = 50 microns.**q,r,** Circle plot showing the strength of MHC class II signaling interactions between the different EC subtypes of the TL (adult/control brain) (**q**) and pathological brain (**r**) ECs at the AV specification level (color-coded by ECs AV specification). **s,t,** Visualization of the differential analysis of MHC class II ligand-receptor pairs. Chord/circos plots showing upregulated MHC class II signaling in angiogenic capillaries as source and all other cell clusters as targets (**s**, top panel), capillaries as source and all other cell clusters as targets (**s**, bottom panel), large veins (**t**, top panel) and veins as receivers (**t**, bottom panel). Edge thickness represents its weights. Edge color indicates the “sender” cell type.

The MHC class II signature was upregulated in all pathologies according to the pattern described above showing highest expression levels at the level of large veins, veins>large arteries and arteries (Figure 5c,g,h, Extended Data Figures 32 and^28^). We observed a partial overlap of the MHC class II and peripheral signatures and of the BBB dysfunction module with a common predominance in large caliber vessels, but a more widespread/stronger expression of the peripheral signature and BBB dysfunction module in angiogenic capillaries (Extended Data Figures 33,34), in line with the previously-observed links between BBB dysregulation and innate immune activation^44^. Together, these data suggest that pathological CNS ECs upregulate MHC class II receptors in brain tumors and brain vascular malformations.

Using immunofluorescence and RNAscope, we validated the upregulation of several MHC class II genes in diseased tissues. We observed enrichment of MHC class II genes including *CD74* and others in the pathological human cerebrovasculature (Figure 5i-p, Extended Data Figure 36).

We next addressed MHC class II signaling among EC clusters by inferring cell-cell communication pathways^44, 57^. Network centrality analysis suggested capillaries (in fetal and control/adult brains) and angiogenic capillaries (in pathological brains) as “senders” and venous ECs (in fetal, adult/control and pathological brains) as major “recipients” of MHC class II signaling and predicted elevated MHC class II signaling (predominantly in angiogenic capillaries) in brain pathologies (Figure 5q-t, Extended Data Figure 35). Notably, MHC class II signaling seems to be mediated mainly by *APP* and *COPA* ligands (and to a lesser extent by *MIF*) and the *CD74* receptor in AV-clusters across development, adulthood and disease (Extended Data Figure 35), and *APP/COPA/MIF–CD74* have been described as ligand-receptor pairs between monocyte-derived macrophages and tumor ECs in human lung adenocarcinoma^70^ and to be involved in antitumor immune response^83^.

### A key role for endothelial cells in the human brain neurovascular unit

Single-cell transcriptomics of unsorted ECs and PVCs offers the opportunity to address cellular crosstalk within the NVU and enables analysis of corresponding ligand-receptor interactions. To address cell–cell interactions in adult/control and diseased brains, we constructed ligand–receptor interaction maps^57, 58^ (Figure 6, Extended Data Figures 37-47, Supplementary Figures 20-28). In the majority of entities, ECs were at the center of the network displaying numerous interactions with other ECs and PVCs (Figure 6a-i, Extended Data Figures 37a-c,f-h,k-m,p-r,38a-c,f-h,k-m,p-r,u-w,z-z_ii_), indicating a crucial role of ECs in NVU function and EC-PVC crosstalk. In fetal and adult/control brains, ECs showed most interactions with fibroblasts, pericytes and astrocytes (Figure 6a-f). In brain pathologies, ECs displayed increased interaction numbers and in brain tumors additionally interacted with tumor (stem) cells (for LGG, GBM, and MET) as well as increased interactions with immune cells (Figure 6h,i, Extended Data Figures 37a-c,f-h,k-m,p-r,38a-c,f-h,k-m,p-r,u-w,z-z_ii_,40a-f). Intercellular signaling pathways were substantially increased in fetal and pathological ECs (and PVCs) (Figure 6j-n, Extended Data Figures 40a-f,42a-h). Cell-cell communication analysis predicted upregulation of several incoming and outgoing signaling pathways in the developing vs control brain as well as diseased versus control brain NVU, including similar pathways to what we observed in EC-EC networks (Figures 2n,o,6o,p, Extended Data Figures 40g-k, 42l-q, Supplementary Figure 21-24). Intercellular crosstalk analysis predicted a key role for the EC and EndoMT clusters within the fetal, adult/control and diseased brain NVU signaling networks (Figure 6o,p, Extended Data Figures 40d-f,42i-k,43-46, Supplementary Figures 25-28).

**Figure 6.**
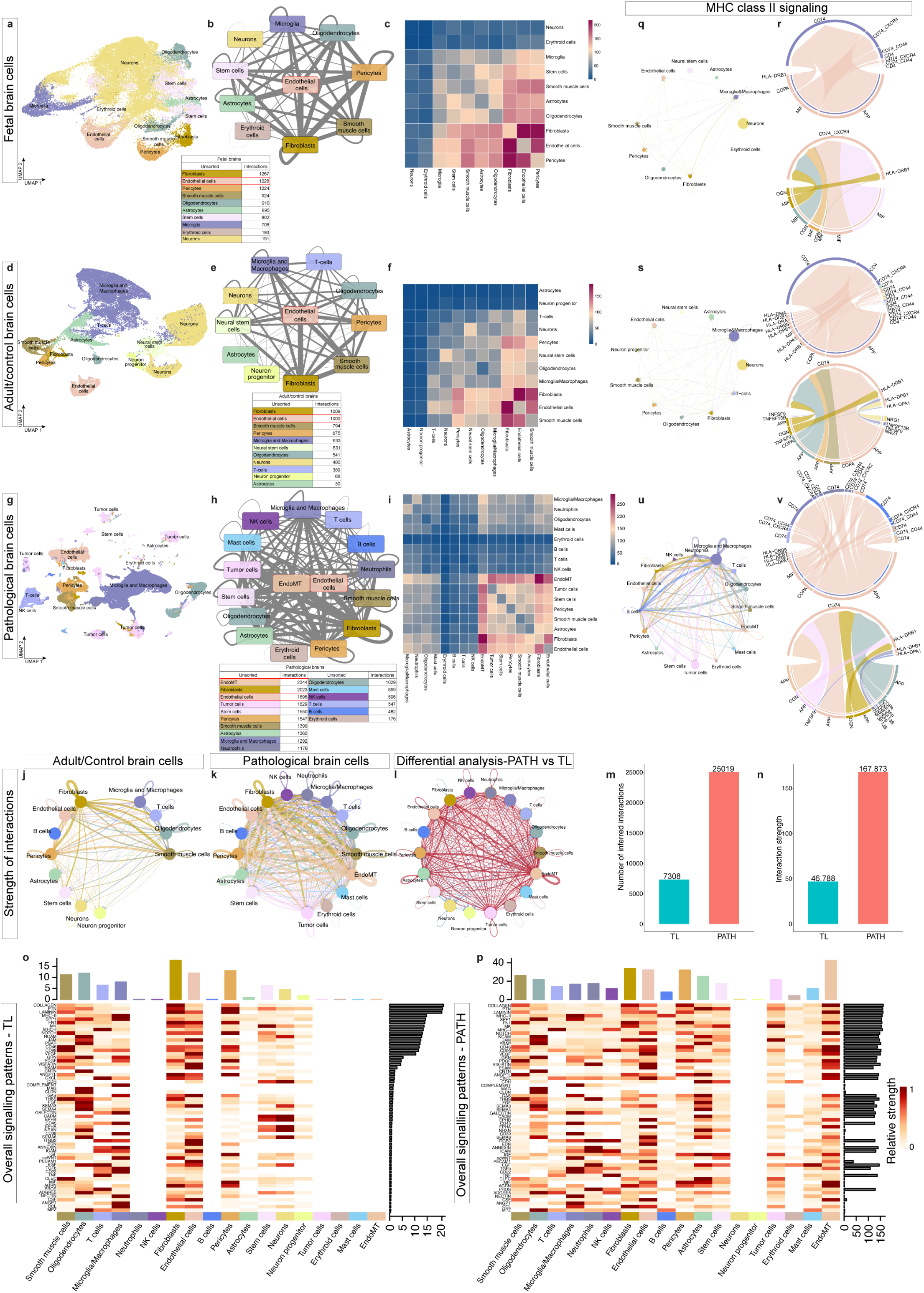
Ligand-receptor interactions of endothelial and perivascular cells in the human brain neurovascular unit. **a,d,g**, UMAP plots of endothelial and perivascular cells derived from fetal brains (**a**), adult/control brains (**d**) and pathological brains (**g**). **b,e,h,** Ligand and receptor analysis of fetal brain (**b**), adult control brain (**e**), and brain pathology (**h**) cells using CellphoneDB. Line thickness indicates the number of interactions between cell types. Tables summarize the number of interactions for each cell type. **c,f,i,** Heatmap showing the number of ligand-receptor interactions between the different cells of fetal brains (**c**), adult/control brains (**f**), and pathological brains (**i**). **j-l,** Circle plot showing the strength of statistically significant interactions between cells of adult/control brains (**j**), and pathological brains (**k**). **l,** Circle plot of the differential analysis of the strength of interactions showed in j and k (pathology over control), red indicating upregulation, while blue indicating downregulation. **m,n,** Barplots showing the number (**m**) and strength of interactions (**n**) in adult/control brains (TL) and pathological brains (PATH). **o,p,** Heatmap showing overall signaling patterns of different cell types in adult control (**o**) and pathological (**p**) brains. **q,s,u,** Circle plots showing the strength of MHC class II signaling interactions between the different cell types of fetal brain (**q**), adult/control brain (**s**), and pathological brain (**u**). **r,t,v,** Visualization of MHC class II connectomic analysis. Cord/circos plots of MHC class II ligand-receptor interactions with EC as “senders” (upper panel) and as “receivers” (lower panel) in fetal brains (**r**), adult/control brains (**t**) and pathological brains (**v**). Edge thickness represents its weights. Edge color indicates the “sender” cell type. In both Circos plots ligands occupy the lower semicircle and corresponding receptors the upper semicircle, and ligands and receptors are colored by the expressing cell type.

To further address the role of ECs in EC-PVC crosstalk in fetal versus adult/control and pathological versus adult/control brains, we identified the cell-cell signaling patterns with ECs on either the “sending” or the “receiving” end (Extended Data Figures 43-46, Supplementary Figures 25-28). During fetal brain development and in brain pathologies, we observed upregulation of ligands and receptors on ECs as well as of the corresponding pathways, which partially overlapped with the ones we found in EC-EC crosstalk (Figure 6o,p, Extended Data Figures 40g-k,41,42l-q,439-46), suggesting that these ligand-receptor interactions may contribute to functional brain EC-PVC signaling and function.

In accordance, we identified large Euclidean distances and substantially changed information flow (see methods) for signaling pathways belonging to the five main groups mentioned above (i.e. immune-related processes, development and NVL, cell-cell/ECM-related processes, angiogenesis) (Extended Data Figure 47c-f),) in both pathologies vs. adult/control and fetal vs. adult/control brains (Extended Data Figure 47a,b), suggesting that these pathways are essential for both EC-EC and EC-PVC crosstalk in fetal and pathological brains where they might critically contribute to vascular growth.

Based on our observation of MHC class II signaling in EC-EC communication, we next addressed MHC class II signaling in EC-PVC intercellular communications. Network centrality analysis corroborated neurons/neural stem cells/ECs (in fetal brains), macrophages and microglia/neurons/ECs (in adult/control brains) and oligodendrocytes/tumor cells/ECs/EndoMT/fibroblasts (in pathological brains) as “senders” and macrophages and microglia as well as ECs (in fetal, adult/control, and pathological brains) as major “recipients” of MHC class II signaling (Figure 6q-v, Extended Data Figure 39) and predicted elevated MHC class II signaling (predominantly in microglia and macrophages, ECs/EndoMT, and tumor cells/oligodendrocytes) in brain pathologies (Figure 6p, Extended Data Figure 39c-m). Of note, the *APP*-*CD74*, *COPA*-*CD74*, and *MIF*-*CD74* ligand-receptor pairs that were predicted to mediate MHC class II signaling in EC-EC interactions (Extended Data Figure 35a-c) were also predicted ligand-receptor pairs in the developing, adult and diseased NVU (Extended Data Figure 39, Supplementary Figure 20), with ECs notably strongly expressing *CD74* (Figure 6r,t,v, Extended Data Figures 43-46). These data indicate that the *APP*/*COPA*/*MIF*–*CD74* ligand-receptor pairs may contribute to NVU signaling.

## DISCUSSION

Here, we performed a large-scale single-cell molecular atlas of the developing fetal, adult/control and diseased human brain vasculature, using single-cell RNA sequencing, composed of 599,215 freshly isolated endothelial, perivascular and other tissue-derived cells from 47 fetuses and adult patients, covering an unprecedented diversity of human brain tissue. Based on genome-wide quantitative single-cell transcriptomes, we have provided molecular definitions of human brain cell types and their differences by brain developmental stage and pathology, thereby unraveling organizational principles of endothelial, perivascular and other tissue-derived cells composing the human brain vasculature. Our experimental methodology relies on transcriptional profiles of human cerebrovascular cells generated from fresh human neurosurgical resections and fresh fetal abortions, that reduces the likelihood of transcriptional alterations associated with post mortem tissue asservation^75–77^.

Our data suggest a paradigm in which developmentally established characteristics and activated pathways of the fetal brain vasculature are silenced in the adult control brain and (re)activated in the vasculature across various brain pathologies, indicating functional plasticity of the endothelial lineage across developmental and disease states.

We uncover the transcriptional basis of the cellular and molecular heterogeneity in the fetal, adult/control and pathological human brain thereby enabling us to identify a treasure trove of novel findings that includes properties conserved across pathologies characterizing a common “backbone” of the diseased human brain vasculature: we observe specific alterations of arteriovenous differentiation, as well as dysregulated/aberrant- and reactivated fetal pathways conserved in the diseased vasculature across multiple pathologies. Pathological ECs display a loss of CNS-specific properties and reveal an upregulation of MHC class II molecules, indicating atypical features of pathological CNS ECs.

CNS-specificity of ECs revealed phenotypic zonation along the arteriovenous axis in the fetus and adult mainly at the level of small-caliber vessels. Zonal characteristics also arise in disease states, where CNS ECs take on a peripheral signature at the level of large>small caliber vessels and angiogenic capillaries. We further identified ECs expressing MHC class II genes mainly in diseased large-caliber vessels, suggesting a role for ECs in immune surveillance^33^. Our work also revealed that upregulation of MHC class II gene expression partially co-occurs with alteration of CNS-specificity and acquisition of a peripheral signature, suggesting that these two observations might be linked, as recently suggested for immune activation and loss of BBB tight junction protein expression in the mouse brain^62^. These similar observations made across species and various diseases indicate their broad applicability. We have thus unveiled a molecular blueprint for zonation and fundamental EC properties (CNS-specificity, MHC class II expression) along the arteriovenous axis at the single-cell level.

Cell-cell interaction analysis predicted strong endothelial-to-perivascular cell ligand-receptor crosstalk involving immune-related (including MHC class II) and angiogenic pathways, thereby unraveling a central role for the endothelium within developing, adult/control and diseased brain NVU signaling networks. Our findings suggest a cellular and molecular environment within the NVU with notable parallels between the fetal and adult brains as well as brain vascular malformations and brain tumors, in which ECs increase their crosstalk with other cell types during development and in various brain diseases.

Our human vascular brain atlas provides a basis for understanding the organizing principles and single-cell heterogeneity of universal, specialized and activated endothelial and perivascular cells with broad implications for physiology and medicine and serves as a powerful publicly available reference for the field.

## METHODS

### Ethics statement

The collection of human samples and research conducted in this study were approved by the institutional research ethics review boards of the University Hospital Zurich, the University Health Network Toronto and the Mount Sinai Hospital Toronto (approval numbers: BASEC 2016-00167, 13-6009, 20-0141-E). Informed consent for fetal tissue collection and research was obtained from each patient after her decision to legally terminate her pregnancy but before the abortive procedure was performed. For adult tissue collection, informed consents for collection and research use of the surgically removed adult brain tissues was obtained from each patient before the operation. Details on patient information and pathology reports are provided in Supplementary Table 1. All the protocols used in this study were in strict compliance with the legal and ethical regulations of the University of Zurich, the University of Toronto and affiliated hospitals.

### Isolation of FACS-sorted human endothelial cells and of unsorted human endothelial and perivascular cells for single cell RNA-seq

Endothelial cells were isolated from human tissues using tissue digestion and subsequent FACS sorting whereas human endothelial and perivascular cells were isolated from the unsorted fraction. Briefly, tissues were quickly minced in a petri dish on ice, using two surgical blades. For FACS sorting a cell suspension was obtained upon digesting the tissue in 2 mg/ml Dispase II (D4693, Sigma-Aldrich, Steinheim, Germany), 2 mg/ml Collagenase IV (#1710401, Thermo Fisher Scientific, Zurich, Switzerland) and 2 mM CaCl2 PBS solution for 40 min at 37°C with occasional shaking. The suspension was filtered sequentially through 100/70/40 μm cell strainers (#431751, Corning, New York, USA) to remove large cell debris. Cells were then centrifuged 500 RCF for 5 min at 4°C. In case of a visible myelin pellet 5 ml 25% BSA (ice cold) was overlayed with 5 ml of the sample, centrifuged at 2000 RCF for 20 min (4°C). The supernatant was removed (including the lipid phase) and the pellet was resuspended in 9 ml PBS, followed by another round of centrifugation at 500 RCF for 5 min (4°C). Supernatant was subsequently discarded, while the cell pellets were resuspended in 3 ml of ACK hemolytic buffer at room temperature for 3 minutes. To stop the reaction, 30 ml of ice-cold PBS was added to the mixture and centrifuged at 500 RCF for 5 min at 4°C. The cell pellets were resuspended in FACS buffer (PBS + 1% Bovine Serum Albumin), a volume is taken for unsorted scRNAseq analysis. For FACS sorting the cells were stained with anti- CD31 PE conjugated antibody in a concentration of 1:20 (#566125, clone MBC78.2, BD Pharmingen) and anti-CD45 APC conjugated antibody in a concentration of 1:20 (#17-0459-42, clone HI30, eBiosciences) for 30 min at 4° C, protected from light. Thereafter the cells were washed with 1ml of FACS buffer, centrifuged in a tabletop centrifuge at 500 RCF at 4°C for 5 min. Finally, the cell pellets were resuspended in appropriate volumes of FACS buffer (PBS + 1% Bovine Serum Albumin) and the suspension was passed through a 35 μm cell strainer of a FACS sorting tube (#352235, Corning). Immediately before sorting, SYTOX^TM^ blue was added in 1:1000 (Thermo Fisher Scientific, #S34857) to exclude dead cells from further analysis. Viable (SYTOX^TM^ blue negative) endothelial cells were FACS-sorted by endothelial marker CD31 positivity and negative selection for the brain microglia and macrophages marker CD45, whereas unsorted endothelial and perivascular cells were obtained from the SYTOX^TM^ blue^-^ fraction Cells were sorted by a FACS Aria III (BD Bioscience) sorter using the four-way purity sorting mode directly in EGM2 medium (#CC-3162, Lonza, Basel, Switzerland).

### Single cell RNA-seq and data analysis

FACS sorted endothelial cells and unsorted endothelial and perivascular cells were resuspended in PBS supplemented with BSA (400 µg/ml, Thermo Fisher Scientific#37525), targeting the required a 1,000 cells/μl concentration. Single-cell RNA-seq libraries were obtained following the 10x Genomics recommended protocol, using the reagents included in the Chromium Single Cell v3 Reagent Kit. Libraries were sequenced on the NextSeq 500 (Illumina) instrument, aiming at 50k reads per cell. The 10x Genomics scRNA-seq data was processed using cellranger-5.0.0 with the Homo sapiens Gencode GRChm38.p13 genome Ensembl release. Based on filtered gene-cell count matrix by CellRanger’s default cell calling algorithm, we performed the standard Seurat clustering (version 4.0.0) workflow; raw expression values were normalized and log transformed. In order to exclude low quality cells and doublets, cells with less than 500 or more than 3000 detected genes were filtered out. scDblFinder (v3.13) was used to validate that doublets cells were minimal. We also filtered cells with > 25% mitochondrial counts. For integration of the unsorted and endothelial cell datasets, Seurat based canonical correlation analysis (CCA) and reciprocal PCA (RPCA) were used for unsorted (endothelial and perivascular cells) and sorted (endothelial cells) datasets batch correction respectively.

To predict the cell identity of the pathological brain endothelial cell clusters of the adult and fetal datasets as compared to the adult/control brain endothelial cells, the cell identity classification and label transfer was done using the standard Seurat workflow using the temporal lobe endothelial cells. as the reference dataset. Illustration of the results was generated using Seurat (v.4.0.0), Sankey plots were done using networkD3 (v.0.4). DESeq2 package (v1.30.1) was used to perform pseudobulk differential expression analysis, volcano plots were done using EnhancedVolcano (v1.8.0) package and heatmaps were plotted using pheatmap (v1.0.12).

Differential expression was computed using the wilcoxauc function implemented in the github package presto. FDR values were calculated using the Benjamini–Hochberg method. Pathway analysis was performed using the Gene Set Enrichment Analysis (GSEA) software from the Broad Institute (software.broadinstitute.org/GSEA) (version 4.0.1)^78, 79^. A permutation-based *P*-value is computed and corrected for multiple testing to produce a permutation based Benjamini – Hochberg correction false-discovery rate q-value that ranges from 1 (not significant) to 0 (highly significant). The resulting pathways were ranked using NES and FDR q-value, *P*-values were reported in the GSEA output reports.

Human_GOBP_AllPathways_no_GO_iea_March_01_2021_symbol.gmt from [http://baderlab.org/GeneSets] was used to identify enriched pathways in GSEA analysis. Highly related pathways were grouped into a themes, labeled by AutoAnnotate (version 1.3) and plotted using Cytoscape (Version 3.7.0) and EnrichmentMap (version 3.3)^80^. Data will be deposited on GEO and an accession number will be provided and code will be provided in github.

### Definition of endothelial fetal/adult, AV, CNS, peripheral and MHC class II signatures

Fetal/adult brain EC signature: we defined a human fetal/developmental and adult brain EC signature - revealing properties of the developing and mature human brain vasculature - comprising the top 50 the genes that passed the threshold of at least log2>0.25 and *P*<0.05 enriched in fetal brain ECs compared with adult brain ECs and vice versa.

Arteriovenous signature: we defined an AV-signature comprising 1,021 genes revealing significant expression gradients along the arteriovenous axis in the adult/control brains, only genes that passed the threshold of at least log2>0.25 and *P*<0.05 were used to construct the signature. CNS and peripheral signatures: we defined a human adult and fetal – endothelial CNS and peripheral signatures comprising the top 50 the genes and at least twofold (log2>1.000) and *P*<0.05 enriched in brain ECs compared with ECs of peripheral organs (heart, kidney, muscle and colon in our fetal dataset) and vice versa. For the mouse derived endothelial CNS and peripheral signature we used the genes described in^62^.

MHC class II signature: comprised of the following list of genes: *CD74*, *HLA-DRB5*, *HLA-DRB1*, *HLA-DQB1*, *HLA-DQB2*, *HLA-DPB1*, *HLA-DOB*, *HLA-DBM*, *HLA-DRA*, *HLA-DQA1*, *HLA-DQA2*, *HLA-DPA1*, *HLA-DOA* and *HLA-DMA*.

The blood-brain barrier (BBB) dysfunction module: comprised the top 50 genes that are upregulated in CNS ECs upon various disease triggers (e.g. stroke, multiple sclerosis, traumatic brain injury and seizure) in the mouse brain and that shifts CNS ECs into peripheral endothelial cell-like states under these conditions^62^. Genes comprising this signature are implicated in such as cell division, blood vessel development, inflammatory response, wound healing, leukocyte migration and focal adhesion^62^.

### Pseudotime trajectory analysis

Pseudotime analysis was performed using Monocle 3^81^ in fetal, adult/control and pathological brain endothelial cells. Endothelial cells were clustered using the standard Seurat (version 4.0.0) clustering procedure and cluster markers were used to AV annotate those clusters, which were used as an input into Monocle to infer trajectory/lineage/arteriovenous relationships within endothelial cells. SeuratWrappers (v.0.3.0) was used to convert the Seurat objects to cell data set objects, while retaining the Seurat generated UMAP embeddings and cell clustering and then trajectory graph learning and pseudo-time measurement with Monocle3.

### Cell-cell communication and ligand-receptor interaction analysis

Cell-cell (ligand receptor) interaction analysis between vascular cell types as well as endothelial and perivascular cells was performed using two published packages: CellPhoneDB^58^ and Cellchat^57^. First, using CellphoneDB ligand-receptor pairing matrix was constructed as follows; only ligands and receptors expressed in at least 10% of the cells in a particular cluster were considered, cluster labels were then permuted randomly 1,000 times to calculate the mean expression values of ligands and receptors, followed by pairwise comparisons between all cell types. The cut-off of expression was set to more that 0.1 and *P*-value to less than 0.05. The number of paired cell-cell interactions was based on the sum of the number of ligand-receptor interactions in each of the cell–cell pairs. Finally, Cytoscape was used to visualize the interaction network as a degree sorted circle layout^80^.

Second, using CellChat (v.1.1.0) we followed the developers’ suggested workflow, briefly applied the pre-processing functions identifyOverExpressedGenes, identifyOverExpressedInteractions, and projectData with standard parameters set. The CellChatDB including the Secreted Signaling pathways, ECM-receptor as well as Cell-Cell contact were analysed, in addition MHC class-II interactions reported in the CellphoneDB. Moreover, the gene expression data was projected onto experimentally validated protein-protein interaction. the standard package functions as computeCommunProb, computeCommunProbPathway and aggregateNet were used with default parameters. Finally, to determine the ligand-receptor contributions and senders/receivers’ roles in the network the functions netAnalysis_contribution and netAnalysis_signalingRole was applied on the netP data slot respectively.

We further compared cell–cell communication patterns by computing the Euclidean distance between ligand-receptor pairs of the shared signaling pathways (a measure of the difference between the signaling networks of datasets, see methods e.g. larger Euclidean distance implying larger difference of the communication networks between two datasets in terms of either functional or structure similarity, termed network architecture)^57^. We compared the information flow for each signaling pathway between, which is defined by the sum of communication probability among all pairs of cell groups for a given signaling pathway in the inferred network^57^.

### Human tissue preparation for immunofluorescence and RNAscope

Freshly resected tissue samples of brain pathologies or the neocortical part of temporal lobes of pharmacoresistant epilepsy patients were obtained from the Division of Neurosurgery, Toronto Western Hospital, University Health Network, University of Toronto and the Department of Neurosurgery, Zurich University Hospital, University of Zurich, whereas fetal tissue was obtained from the Research Centre for Women’s and Infants’ Health (RCWIH) BioBank. Sample age (gestational age for fetal tissues), gender and pathology were documented in Supplementary Table 1. Tissues were fixed at 4°C in 4% paraformaldehyde (PFA) for 12 hours and placed into 30% sucrose in PBS solution overnight. The tissues were then embedded in Optimum Cutting Temperature compound (Tissue-Tek O.C.T. Compound, #4583) and stored in a −80°C freezer.

### Immunofluorescence staining

Fixed, Cryo-embedded, human adult control brains, brain tumors, and brain vascular malformations slices were cut in 40-µm thick sections, using a cryotome (Leica Cryostat 1720 Digital Cryotome), and submitted for single and double staining with the antibodies provided in Supplementary Table 12. Briefly, the sections were: 1) permeabilized with 0.3% Triton X-100 in PBS for 30 min at room temperature (RT); 2) incubated overnight at 4°C with primary Abs, incubated with the appropriate secondary Abs, donkey anti-mouse Alexa Fluor 488 (1:1000, Thermo Fisher Scientific, #A-21202), donkey anti-Guinea pig Alexa Fluor 488 (1:1000, Jackson Immunoresearch Labs, #706-545-148), donkey anti-rabbit Alexa Fluor 555 (1:1000, Thermo Fisher Scientific, #A-31572), donkey anti-goat Alexa Fluor 488 (1:1000, Thermo Fisher Scientific, #A-11055) for 90 min at RT; to quench the autofluorescence signal, tissue sections were treated with 0.1% Sudan Black B (Thermo Fisher Scientific, Cat: J62268) in 70% ethanol for 15 minutes at room temperature. 4) counterstained with the 4, 6- diamidino-2-phenylindole (DAPI) (diluted 1:20,000; BioLegend, #422801). Finally, the sections on glass slides (Fisherbrand, Superfrost Plus Microscope Slides, Fisher Scientific) were coverslipped with VWR micro cover glass (VWR International). Negative controls were prepared by omitting the primary antibodies and mismatching the secondary antibodies. Sections were examined under Zeiss laser scanning confocal microscope (LSM 880). Laser scanning confocal images were taken through the z-axis of the section, with 20x and 40x lenses. Z-stacks of optical planes (maximum intensity projections) and single optical planes were recorded and analyzed by Zeiss Zen software.

### RNAscope

Fixed, cryo-embedded, brain tumor, brain vascular malformations and temporal lobe tissue obtain from adult patients were cut into 40-µm thick sections and subjected to RNAscope in-situ hybridization using the RNAscope HiPlex kit (324100-UM, ACD, Newark, CA) according to the manufacturer’s instructions (pretreatment and RNAscope Multiplex Fluorescent v2 Assay were performed according to protocol 323100-USM). Briefly, after deparaffinization, the slides were incubated with hydrogen peroxide for 10 min at RT. Target retrieval was performed using 1X target retrieval buffer (Ref 322000; ACD, Newark, CA) and distilled water were heated to above 99°C. Slides were dipped in water for 10 seconds before placing into the 1X target retrieval buffer for 5 minutes. Then, slides were quickly washed in distilled water for 15 seconds. An additional Sudan Black step was added for tissues with high auto-immunofluorescence (AVM, LGG, TL, MEN). Slides were submerged in 0.1% Sudan Black in 70% ethanol for 30 minutes at RT, covered with foil. Then, slides were washed three times with distilled water, once in 100% ethanol and dried at 60°C for 5 minutes. Hydrophobic barrier was drawn around the sections and left to dry overnight. On day two, RNAscope Protease III (Ref 322340; ACD, Newark, CA) was added to cover each section and incubated at 40°C for 15 minutes inside the RNAscope HybEZ II oven. For hybridization, 1X probe mixture was added to each section and incubated at 40°C for 2 hours in the RNAscope HybEZ II Oven. For HiPlex Amp 1-3 hybridization, 1X Amp 1 solution was added to each section and incubated at 40°C for 30 minutes in the RNAscope HybEZ II Oven. Amp hybridization steps were sequentially repeated with Amp 2 and Amp 3 solutions. For HiPlex fluorophore hybridization, corresponding fluorophores (1X HiPlex T1-T4 solution) was added to each section and incubated at 40°C for 15 minutes in the RNAscope HybEZ II Oven. Wash steps with the appropriate buffers were done in between each step as indicated in the user manual. HiPlex Amp and fluorophore solutions were included in the RNAscope HiPlex8 Detection (Ref 324110; ACD, Newark, CA) and HiPlex12 Ancillary Kits (Ref 324120; ACD, Newark, CA). After fluorophore hybridization, DAPI (Ref 320858; ACD, Newark, CA) was added for 30 seconds at RT before being replaced with ProLong Gold Antifade Mountant (Ref P36930; Invitrogen, Waltham, MA). We used the following RNAscope probes: Hs-CD31 (Ref 548451-T3), Hs-ESM1 (Ref 586041-T7), Hs-ACTA2 (Ref 311811-T10), Hs-PLVAP (Ref 437461-T1), Hs-HLA-DPA1 (Ref 821641-T6), Hs-CD74 (Ref 477521-T11). Images were acquired using an Olympus FluoView Laser Scanning Confocal Microscope Olympus IX81 inverted stand; 40X objective lens Plan Apo 40x/1.35 NA oil immersion. Laser wavelengths were 405nm, 473nm, 559nm and 635nm. After the first round of imaging, slides were soaked in 4X SSC buffer (#BP1325-1; Fisher Scientific, Waltham, MA) until the cover slip could be removed easily. Fluorophores were cleaved with 10% cleavage solution (Ref 324130; ACD, Newark, CA) at RT for 15 minutes, followed by two washes with PBST (0.5% Tween 20) (repeated cleavage twice). Procedures for round 2 and 3 fluorophore hybridization were the same as the round 1. After three rounds of imaging, image alignment, merging and processing were performed using the RNAscope HiPlex Image Registration Software following the Image Registration Software User Manual (300065-UM) (ACD, Newark, CA).

Visualization was done using the FV10-ASW 4.2 Viewer and ImageJ^82^. Pseudocolors were used for better visualization.

## ACKNOWLEDGMENTS

We thank Niklaus Krayenbühl, Menno Germans, Oliver Bozinov, Philippe Bijlenga, Pierre-Yves Dietrich and Valerie Dutoit for help with the tissue asservation. Edwin Speck and the Flow Cytometry Facility, Krembil Discovery Tower, University Health Network for help with the FACS-sorting. Gurbaksh Basi, Julia Cirlan, Claudia Dumrese and Malgorzata Kisielow for help with the single-cell RNA sequencing experiments. Aiden M. Sababi and Mohammed Mahmoud Saad for help with the computational analysis, Nancy Chu Ji for help with the illustrations and Ashley Thomson for help with English proofreading.

The author(s) disclosed receipt of the following financial support for the research, and/or publication of this article: T.W. was supported by the OPO Foundation, the Swiss Cancer Research foundation (KFS-3880-02-2016-R, KFS-4758-02-2019-R), the Stiftung zur Krebsbekämpfung, the Kurt und Senta Herrmann Foundation, Forschungskredit of the University of Zurich, the Zurich Cancer League, the Theodor und Ida Herzog Egli Foundation, the Novartis Foundation for Medical-Biological Research and the HOPE Foundation. P.M. was supported by the Canadian Institutes of Health Research. I.R. was supported by the Canadian Institutes of Health Research. G.D.B. was supported by NRNB (U.S. National Institute of Health, National Center for Research Resources grant number P41GM103504.

## AUTHOR CONTRIBUTIONS

T.W. had the idea for the study, T.W. and I.R. conceived the study, T.W. designed the experiments, wrote the manuscript, analyzed the data, designed the figures and made the figures with M.G. and the help of M.S., I.R., T.W., L.R., K.S., M.B., P.K., G.Z, T.V. acquired the tissue. M.G., M.S., S.S. S.V. performed the cell isolations experiments. T.W. and K.F. developed the initial isolation experiments. M.G. (majority of the analysis), M.S., H.Z., D.R.Z., H.R. performed the single-cell data processing and analyzed the data with T.W.. T.W. and G.B. supervised the data analysis and interpreted data. R.S. and T.K. helped with single-cell data processing. M.G., M.S., G.B. developed the computational methods. S.T., S.S., R.W., K.Y. performed the immunofluorescent stainings and RNA scope experiments. T.W., I.R., G.B., P.P.M. acquired funding. T.W., P.P.M., I.R. and G.B. edited the final version of the manuscript. M.G., M.S., J.B. helped editing the manuscript. P.P.M., K.B., J.E.F., M.L.S., P.D., P.C., V.T., G.Z., T.V., I.R., G.B. gave critical inputs to the manuscript. T.W. supervised all the research. All authors read and approved the final manuscript.

## EXTENDED DATA FIGURE LEGENDS

**Extended Data Figure 1.**
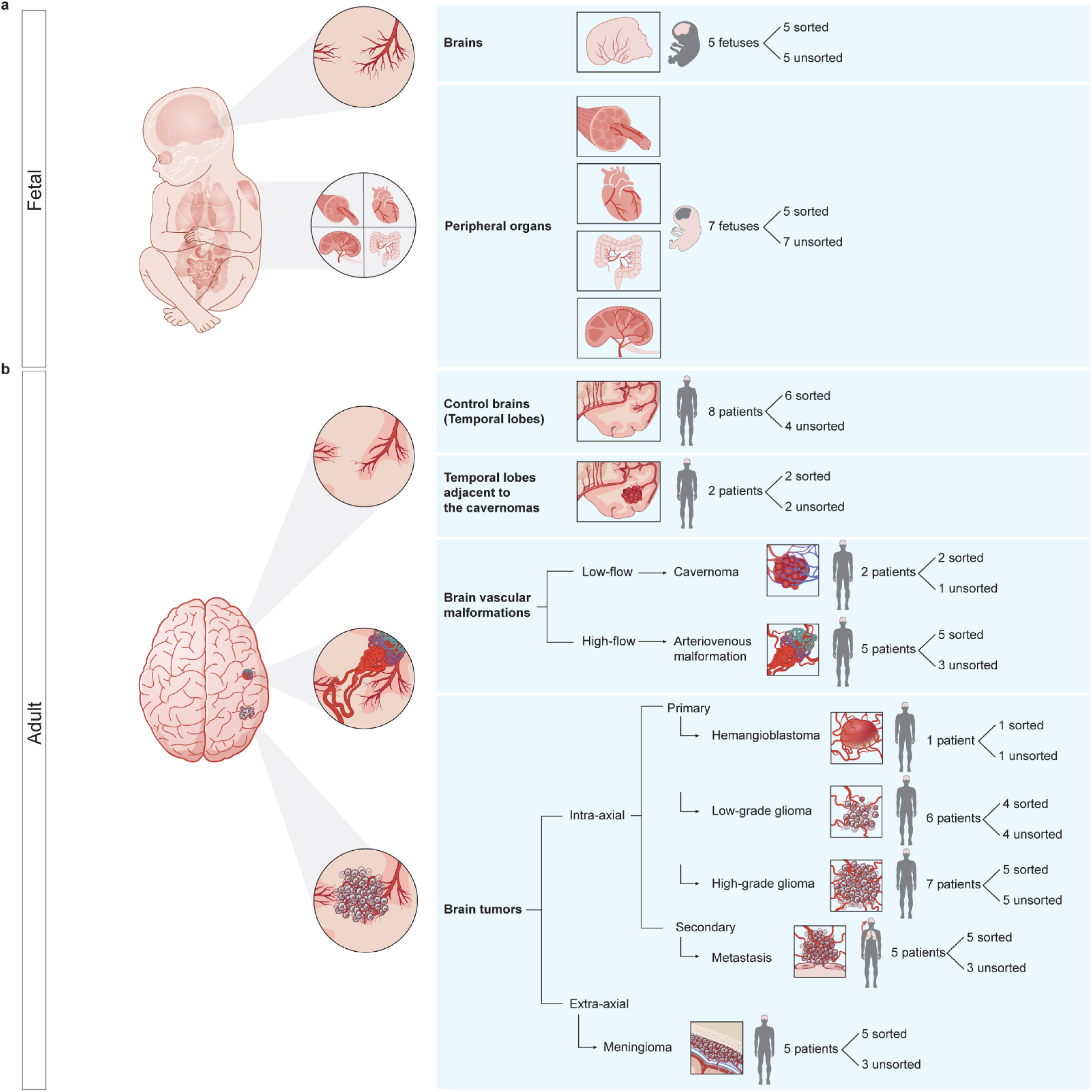
**a,b,** Scheme of the different tissue types present in the study with respective sample/patient numbers of fetal (**a**) and adult (**b**) origins.

**Extended Data Figure 2.**
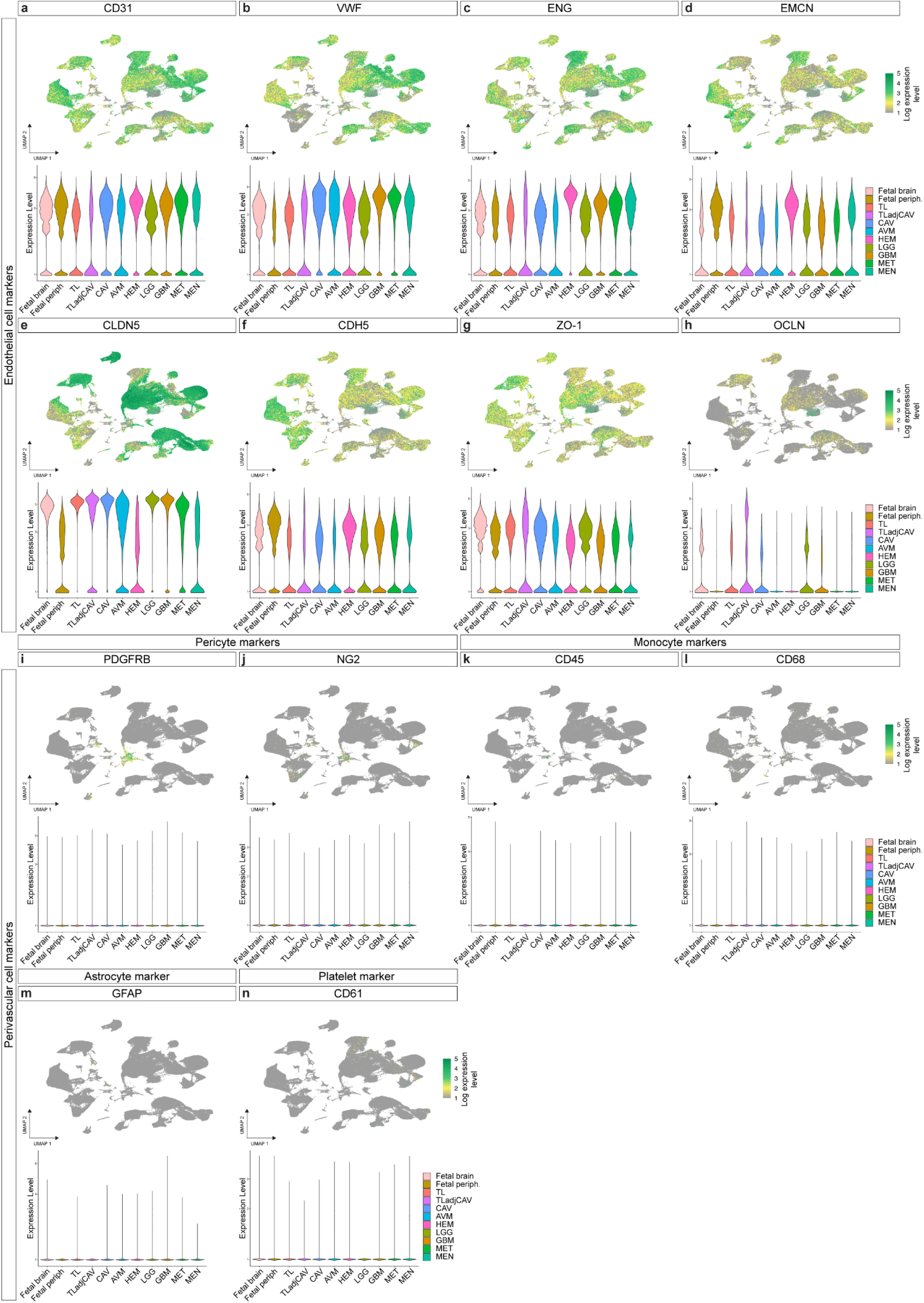
**a-n,** Violin plots showing the expression of endothelial (**a-h**) and perivascular (**i-n**) markers in the isolated endothelial cells from fetal, adult/control and pathological brains.

**Extended Data Figure 3.**
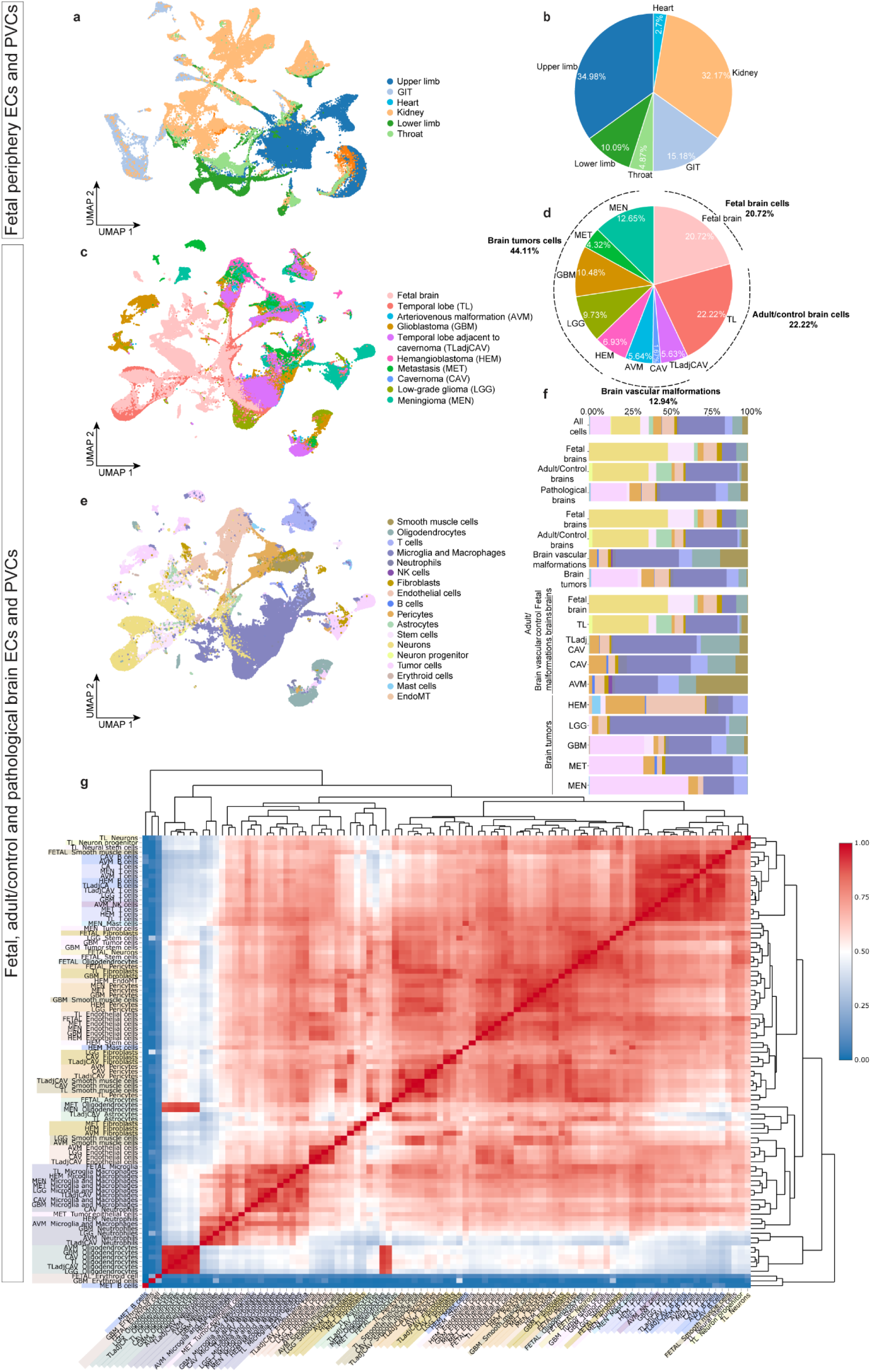
**a,** UMAP plot of 90,150 human fetal periphery endothelial and perivascular cells, colored by tissue of origin. **c,** UMAP plot of 212,857 human brain endothelial and perivascularcells, colored by tissue of origin (**c**) and by cell type (**e**). **b,d,** Piechart showing relative abundance and percentage of cells from each tissue collected. **f,** Relative abundance of cell types from the indicated tissue of origin. Color-code corresponds to (**e**). **g,** Endothelial and perivascular cells transcriptome correlation heatmap and hierarchical clustering.

**Extended Data Figure 4.**
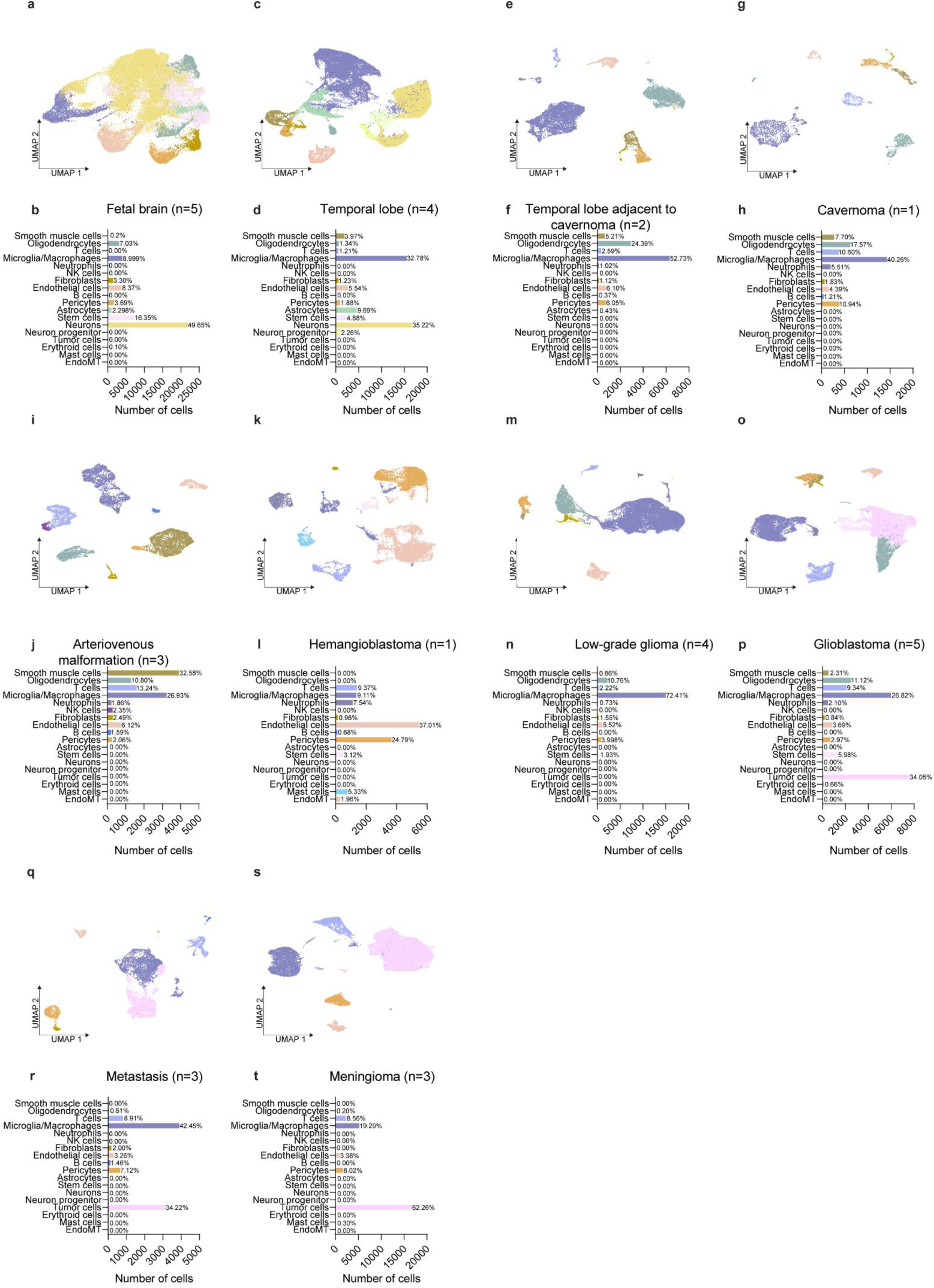
**a-s,** UMAP plots of ECs and PVCsfor each tissue of origin indicated, color-coded by cell type. Bar plots showing the number and proportion of cells of each cell type are shown below each corresponding UMAP.

**Extended Data Figure 5.**
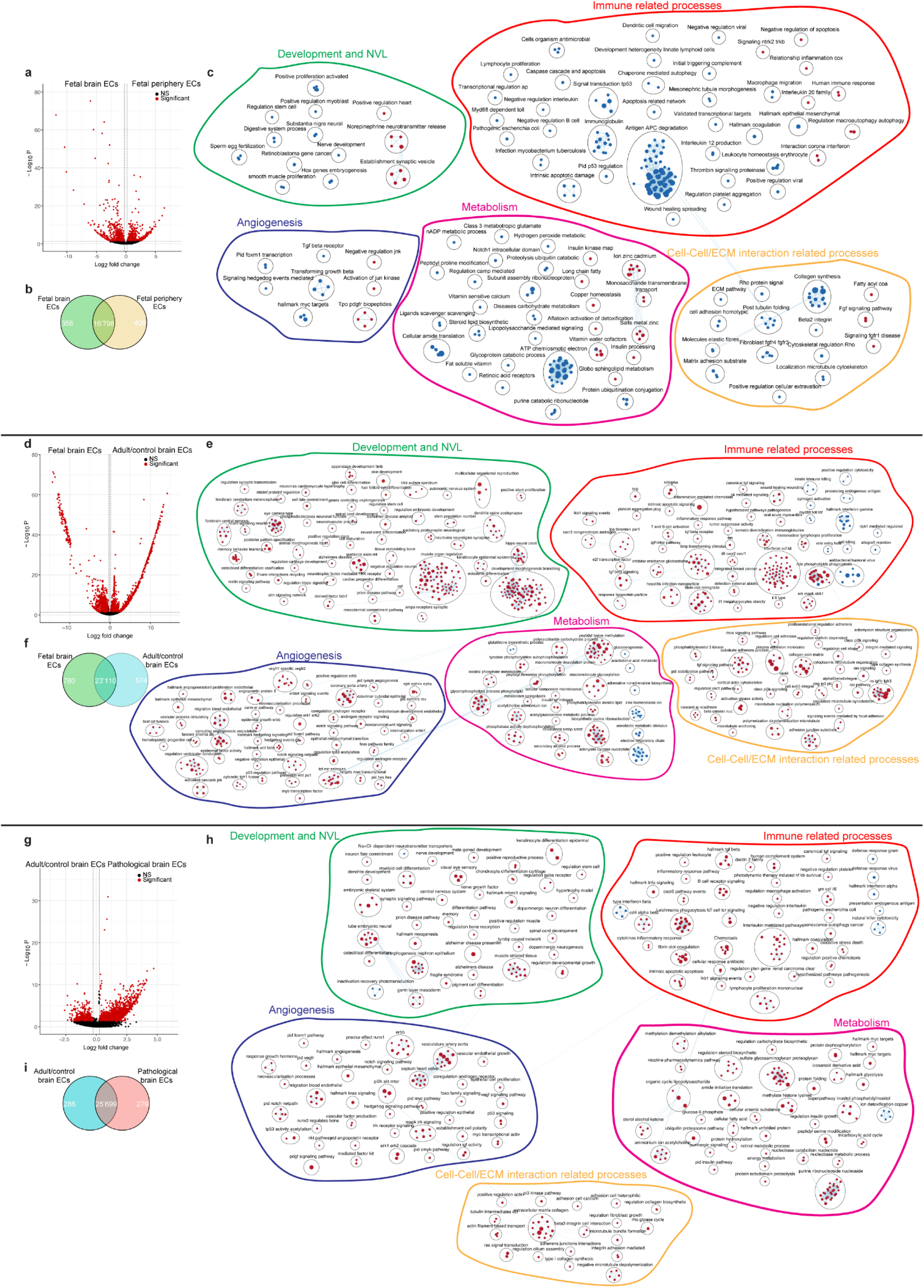
**a,d,g,** Volcano plot showing the differential expression analysis comparing endothelial cells from fetal brain (left) and fetal periphery (right) (**a**), fetal (left) and adult/control brains (right) (**b**), adult/control (left) and pathological brains (right) (**c**) (Benjamini Hochberg correction; *P*-value<0.05 and log2FC≥0.25 colored significant in red). **b,** Heatmap and hierarchical clustering of all significant genes comparing fetal and adult/control brain endothelial cells. **b,f,i,** Venn diagram showing the number of differentially expressed genes between the indicated enities. **c,e,h** Enrichment map visualizing the significantly enriched pathways from the GSEA analysis performed on the ranked list of differentially expressed genes between fetal and adult/control brain endothelial cells. Results include enriched genesets belonging to development and NVL, angiogenesis, cell-cell/extracellular matrix interaction, metabolism, and immune related processes.

**Extended Data Figure 6.**
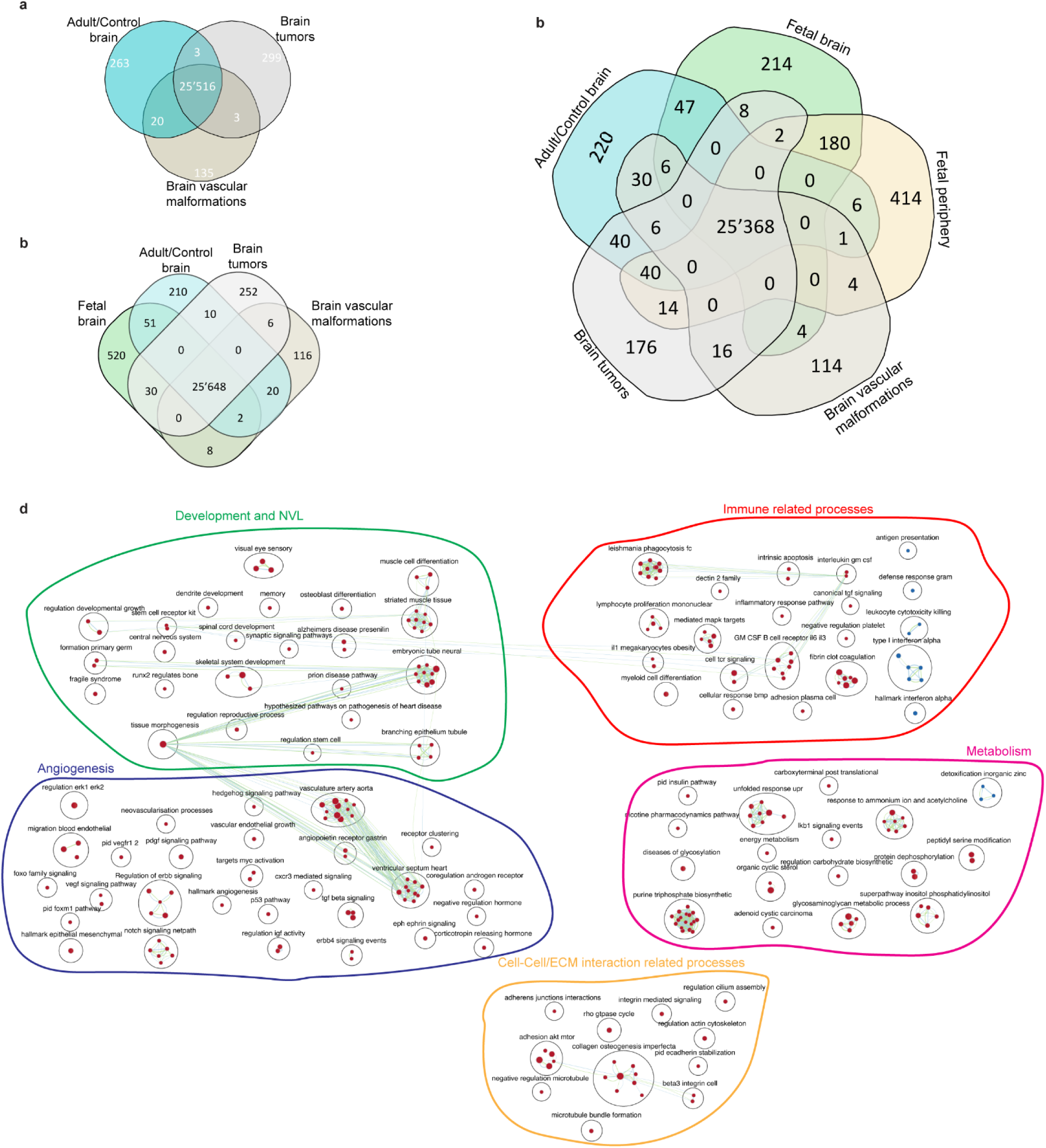
**a-c,** Venn diagram showing the number of differentially expressed genes between the indicated tissue types. **d,** Enrichment map visualizing the 357 commonly enriched pathways from the GSEA pathway analysis, commonly enriched to fetal and pathological brain endothelial cells as compared to adult/control brain endothelial cells. Results include enriched genesets belonging to development and NVL, angiogenesis, cell-cell/extracellular matrix interaction, metabolism, and immune related processes.

**Extended Data Figure 7.**
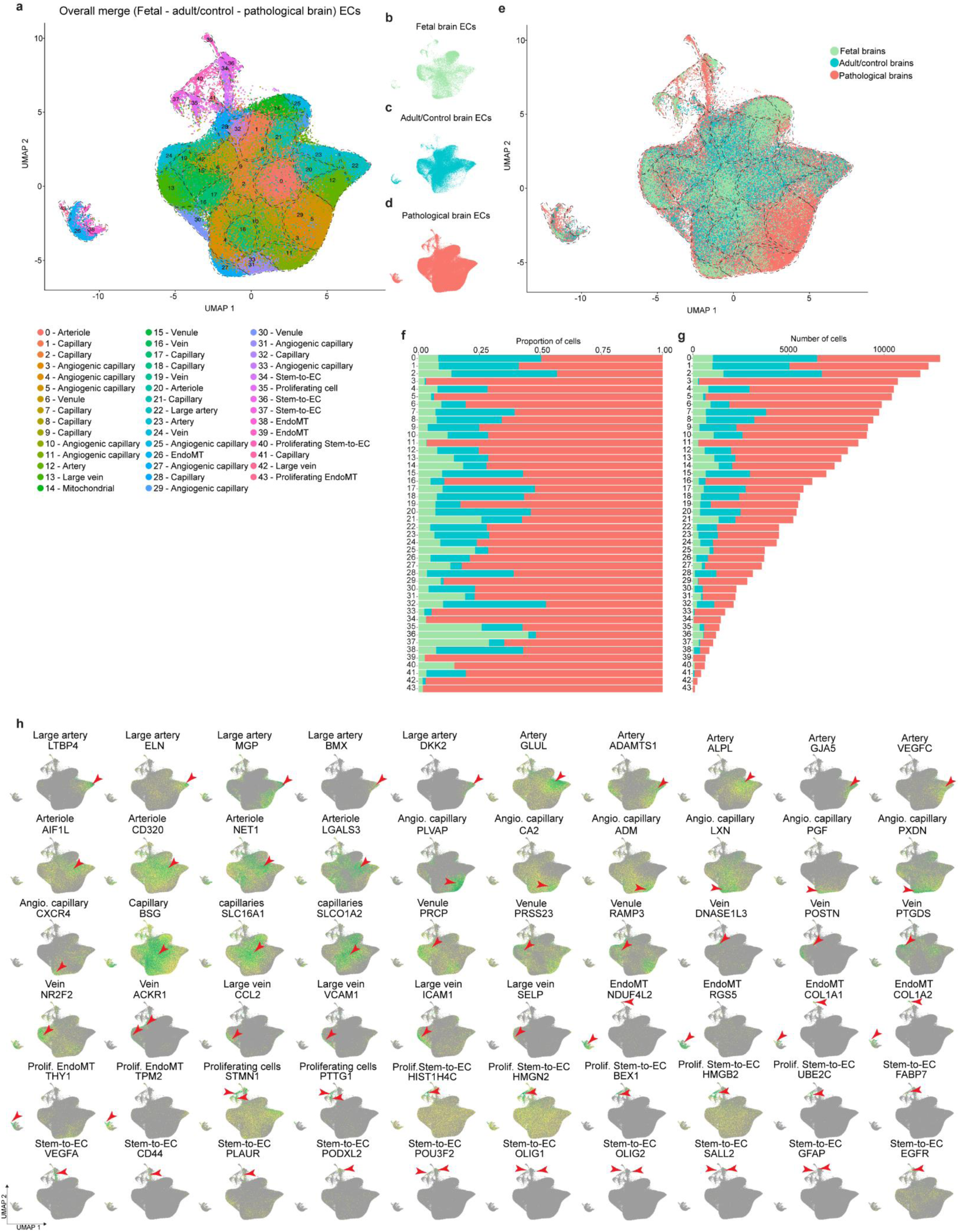
**a,e,** UMAP plot of the 296,208 batch corrected ECs, color-coded by seurat clusters (**a**), cluster annotation in indicated in the legend and by tissue of origin (**e**). **b-d,** UMAP showed in (**a**) split by tissue of origin: fetal brain (**b**), adult/control brain (**c**) and brain pathologies (**d**). **f,g,** Relative abundance (**f**) and absolute number of (**g**) endothelial cells in the different seurat clusters, color-code corresponds to tissue of origin: fetal brain (green), adult/control brain (cyan) and brain pathologies (red). **h,** UMAPs plots, color-coded for expression of indicated marker genes (red arrowheads).

**Extended Data Figure 8.**
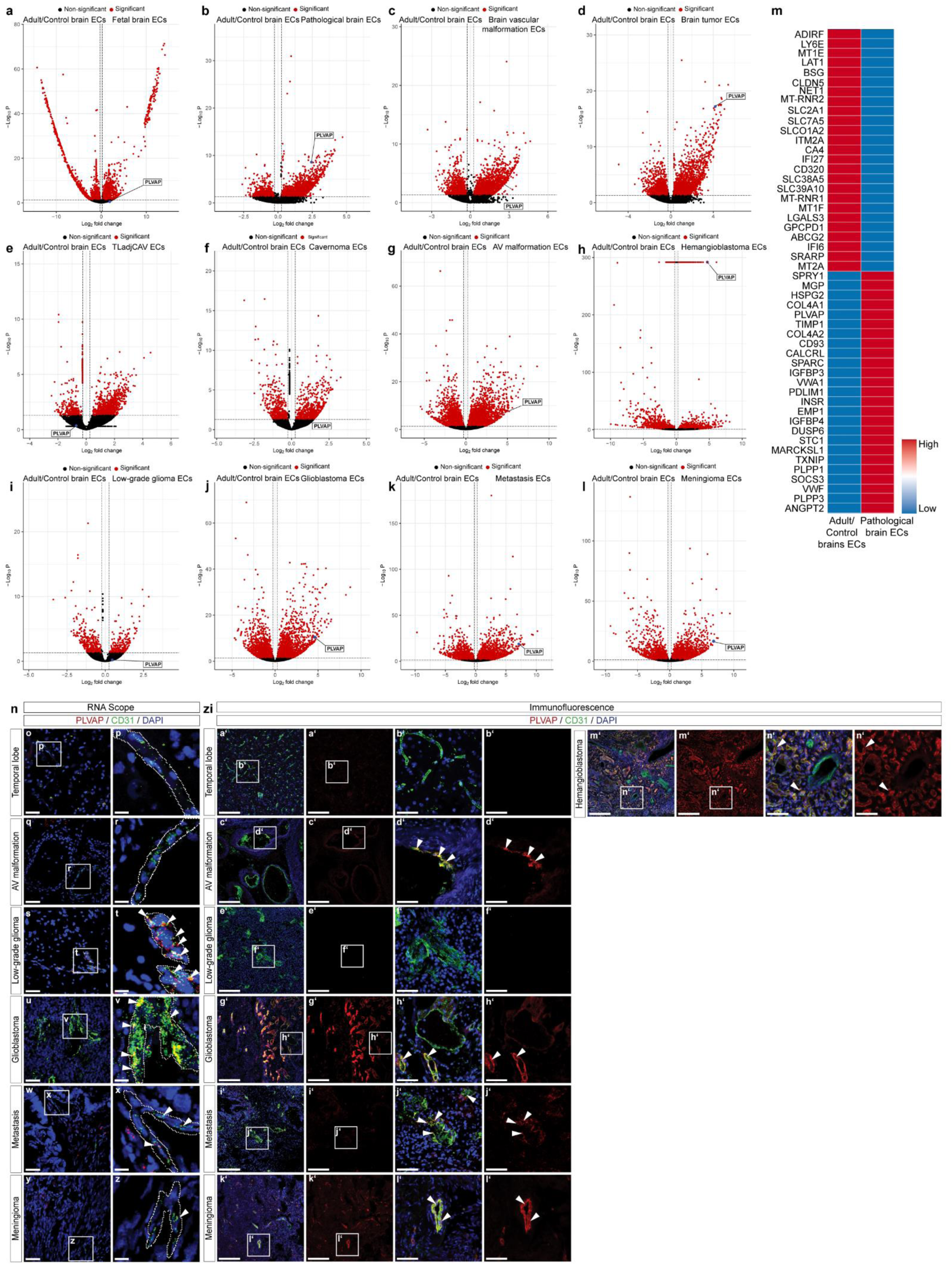
**a-l**, Volcano plot showing the differential expression analysis comparing endothelial cells from adult/control brains (left) and the indicated entity (right) (Benjamini Hochberg correction; *P*-value<0.05 and log2FC≥0.25 colored significant in red, PLVAP is colored blue). **m,** Expression heatmap of the top 25 differentially expressed genes in adult/control vs. pathological brain ECs. **n-n’**, Immunofluorescence (IF) and RNAscope imaging of tissue sections from the indicated entities, stained for PLVAP (red; **a‘-n‘,** RNAscope; **o-z**, IF) and CD31 (green). Nuclei are stained with DAPI (blue). Boxed area is magnified on the right; arrowheads (IF) and dotted lines (RNAscope) indicate vascular structures in the different tissues. Scale bars: 200μm in overviews (IF), 50μm in overviews (RNAscope); 50μm in zooms (IF) and 12.5μm in zooms (RNAscope).

**Extended Data Figure 9.**
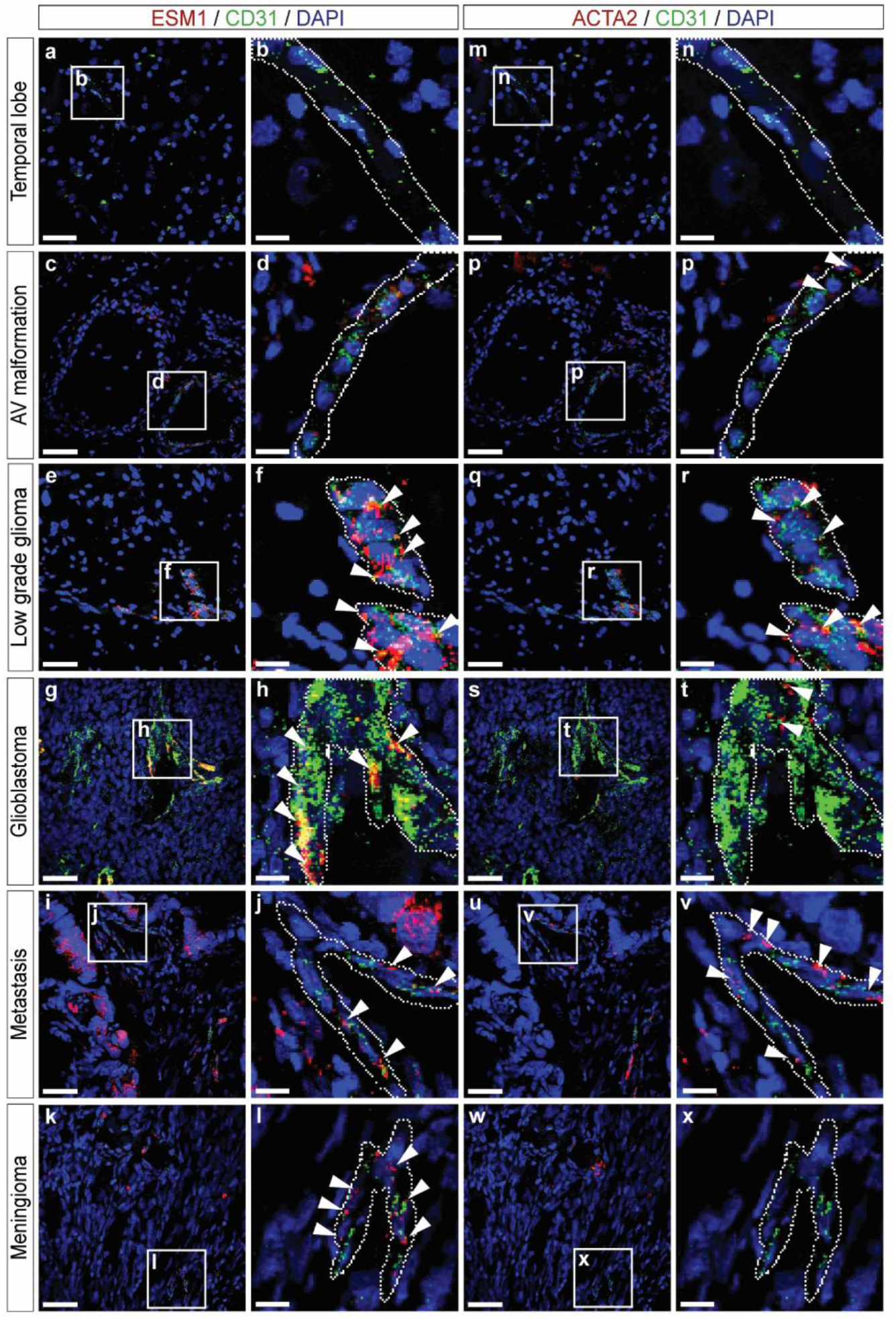
**a-x,** RNAscope imaging of tissue sections from the indicated entities, stained for ESM1 (red; **a-l**), ACTA2 (red; **m-x**) and CD31 (green). Nuclei are stained with DAPI (blue). Boxed area is magnified on the right; dotted lines (RNAscope) indicate vascular structures in the different tissues. Scale bars: 50μm in overviews and 12.5μm in zooms.

**Extended Data Figure 10.**
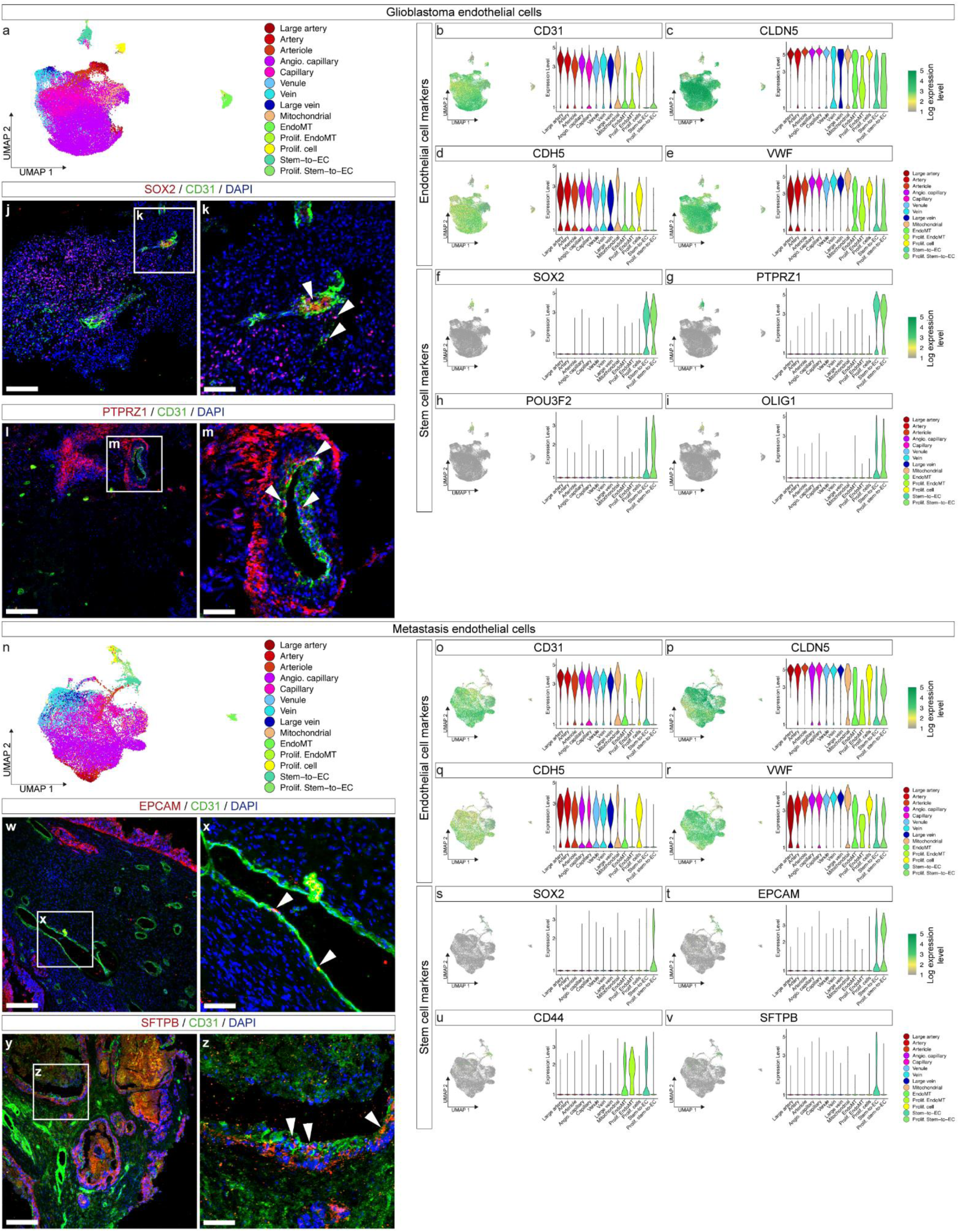
**a,b,** UMAP plot of GBMs and METs endothelial cells, colored by AV specification. **b-i, o-v,** Violin plots showing the expression of the indicated endothelial and stem cell specific markers in the different EC subtypes. **j-m,** Immunofluorescence staining showing co-localization of *SOX2* (**j,k**), *PTPRZ1* (**l,m**), *EPCAM* (**w,x**), *SFTPB* (**y,z**) and *CD31*. Scale bars: 200 µm in overviews (left) and 50 µm in zooms (right), in blood vessels of human GBMs and METs.

**Extended Data Figure 11.**
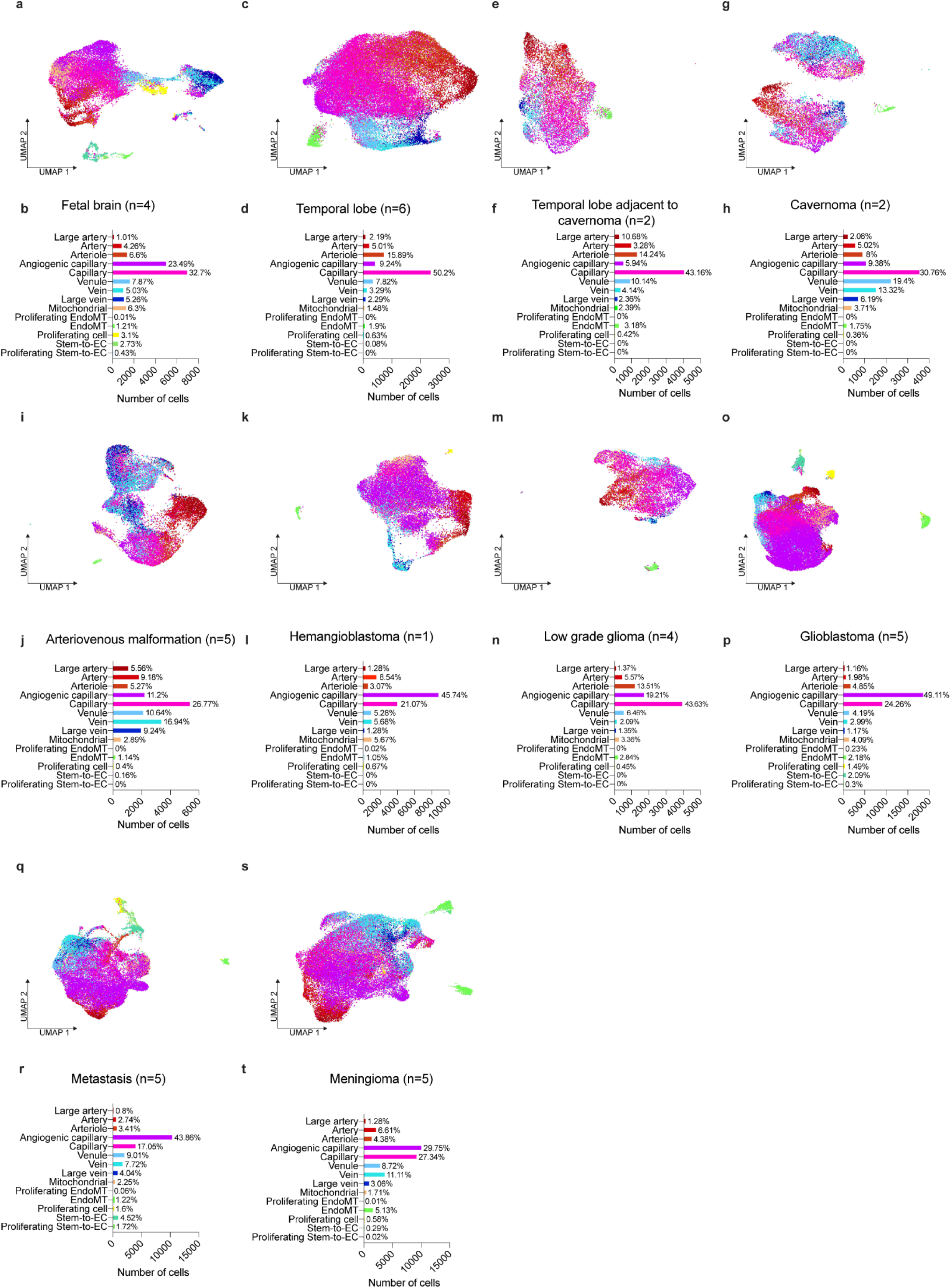
**a-s,** UMAP plots of ECs for each tissue of origin indicated, color-coded by ECs arteriovenous (AV) specification. Barplots showing the number and proportion of each EC subtype are shown below each corresponding UMAP.

**Extended Data Figure 12.**
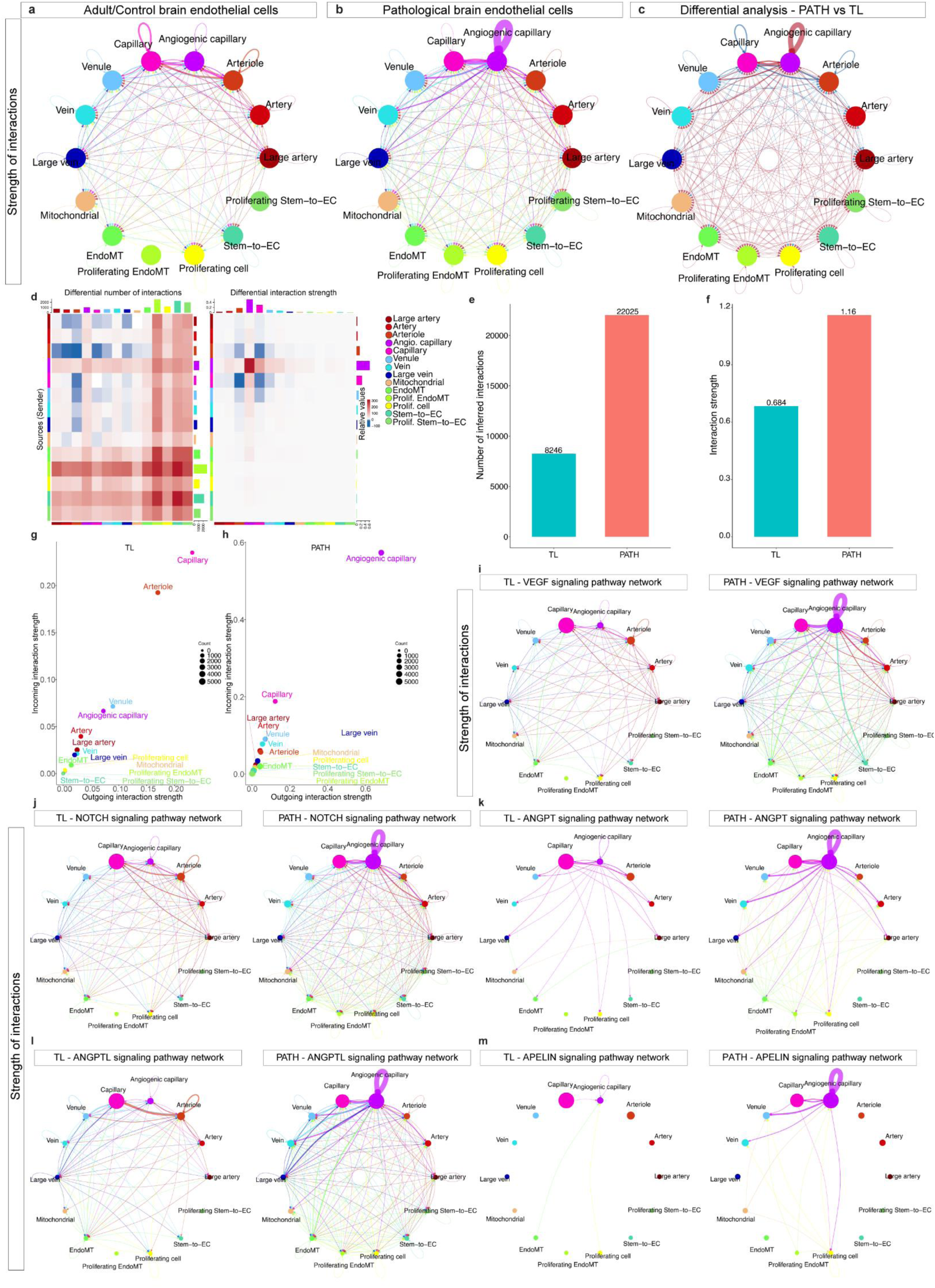
**a-c,** Circle plot showing the strength of statistically significant signaling interactions between EC subtypes of adult/control brain (**a**), pathological brain (**b**) and the differential analysis of the number of interactions (pathology over control) (**c**), red indicating upregulation, while blue indicating downregulation. **d,** Heatmap showing the differential analysis of the number (left) and strength (right) of ligand-receptor interactions for different EC subtypes (pathology over control). **e,f,** Barplots showing the number (**e**) and strength (**n**) of interactions in adult/control brains (TL) and pathological brains (PATH) endothelial cells. **g,h,** Scatter plot showing the strength of outgoing (x-axis) and incoming (y-axis) signaling pathways of different EC subtypes from temporal lobe and pathological brains. **i-m,** Circle plot showing the strength of *VEGF* (**i**), *NOTCH* (**j**), *ANGPT* (**k**), *ANGPTL* (**l**) and *APELIN* (**m**) signaling interactions between temporal lobe and pathological brain endothelial cells (color-coded by AV specification).

**Extended Data Figure 13.**
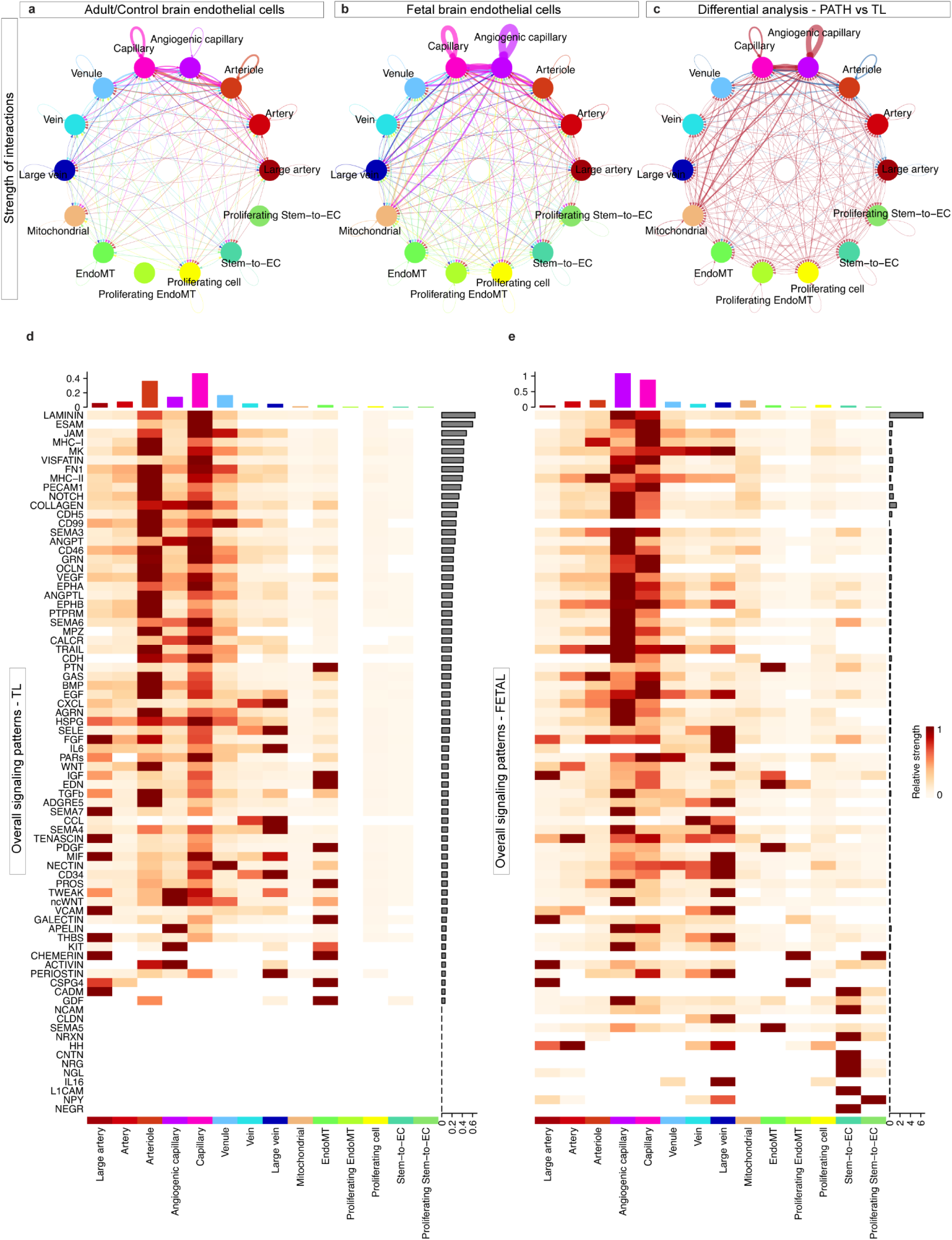
**a-c,** Circle plot showing the strength of statistically significant signaling interactions between EC subtypes of adult/control brain (**a**), fetal brain (**b**) and the differential analysis of the number of interactions (pathology over control) (**c**), red indicating upregulation, while blue indicating downregulation. **d,e,** Heatmap showing overall signaling patterns of different EC subtypes in adult/control (**d**) and fetal (**e**) brains.

**Extended Data Figure 14.**
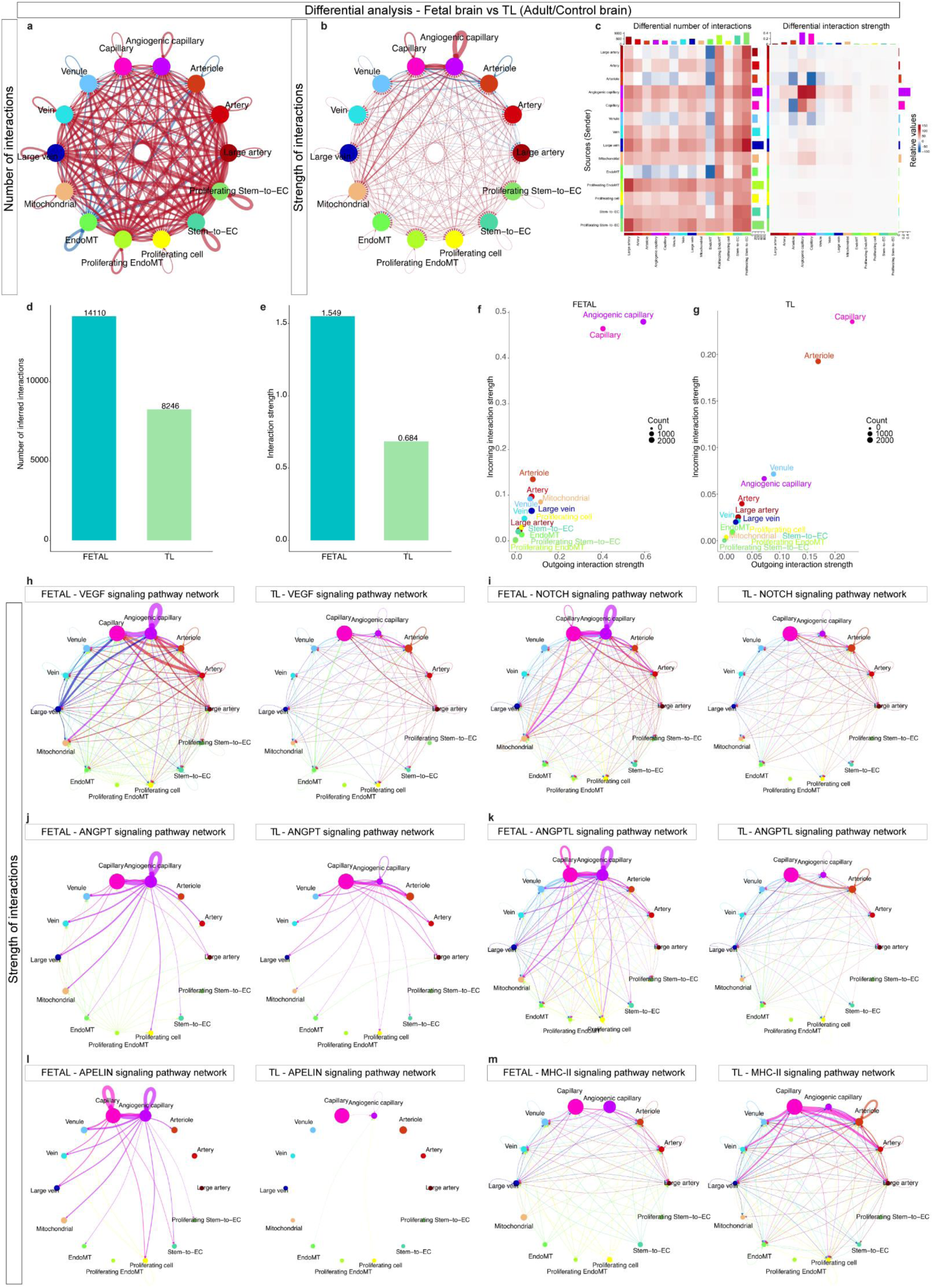
**a,b,** Circle plot showing the differential analysis of the number (**a**) and strength (**b**) of statistically significant ligand-receptor interactions between EC subtypes in adult/control and fetal brains. **c,** Heatmap showing the differential analysis of the number (left) and strength (right) of ligand-receptor interactions for different EC subtypes (fetal over adult/control). **d,e,** Barplots showing the number (**d**) and strength (**e**) of interactions in fetal brain (FETAL) and adult/control brain (TL) endothelial cells. **f,g,** Scatter plot showing the strength of outgoing (x-axis) and incoming (y-axis) signaling pathways of different EC subtypes from fetal and adult/control brains. **h-m,** Circle plot showing the strength of *VEGF* (**h**), *NOTCH* (**i**), *ANGPT* (**j**), *ANGPTL* (**k**), *APELIN* (**l**) and MHC class II signaling interactions between fetal brain and adult/control brain endothelial cells (color-coded by AV specification).

**Extended Data Figure 15.**
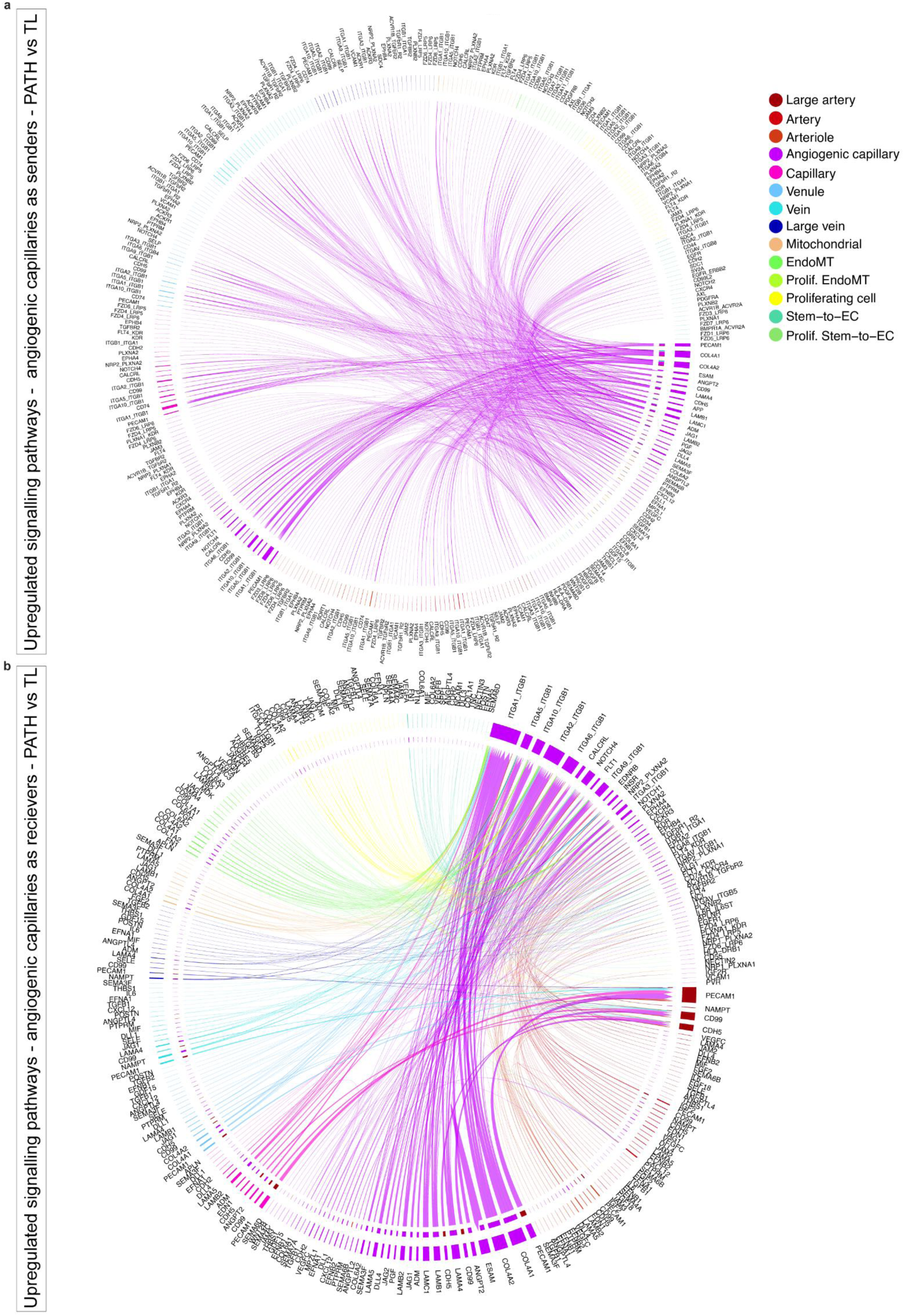
**a,b,** Chord plots showing the ligand-receptor signaling interactions between ECs subtypes upregulated in pathological as compared to adult/control brains; signaling pathways sending from (**a**) and receiving by angiogenic capillaries (**b**). Edge thickness represents edge weights and edge color indicates the sender cell type.

**Extended Data Figure 16.**
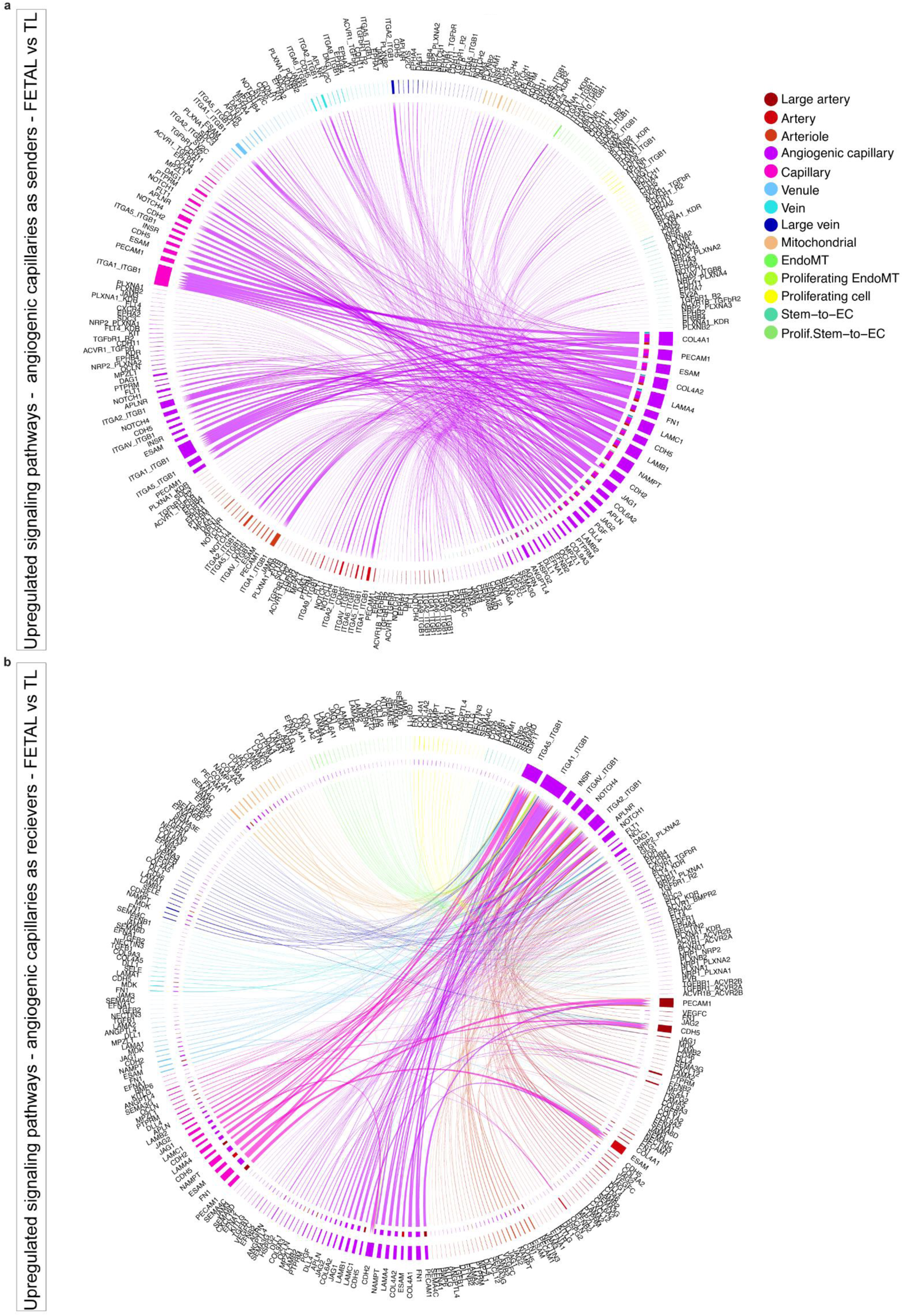
**a,b,** Chord plots showing the ligand-receptor signaling interactions between ECs subtypes upregulated in fetal as compared to adult/control brains; signaling pathways sending from (**a**) and receiving by angiogenic capillaries (**b**). Edge thickness represents edge weights and edge color indicates the sender cell type.

**Extended Data Figure 17.**
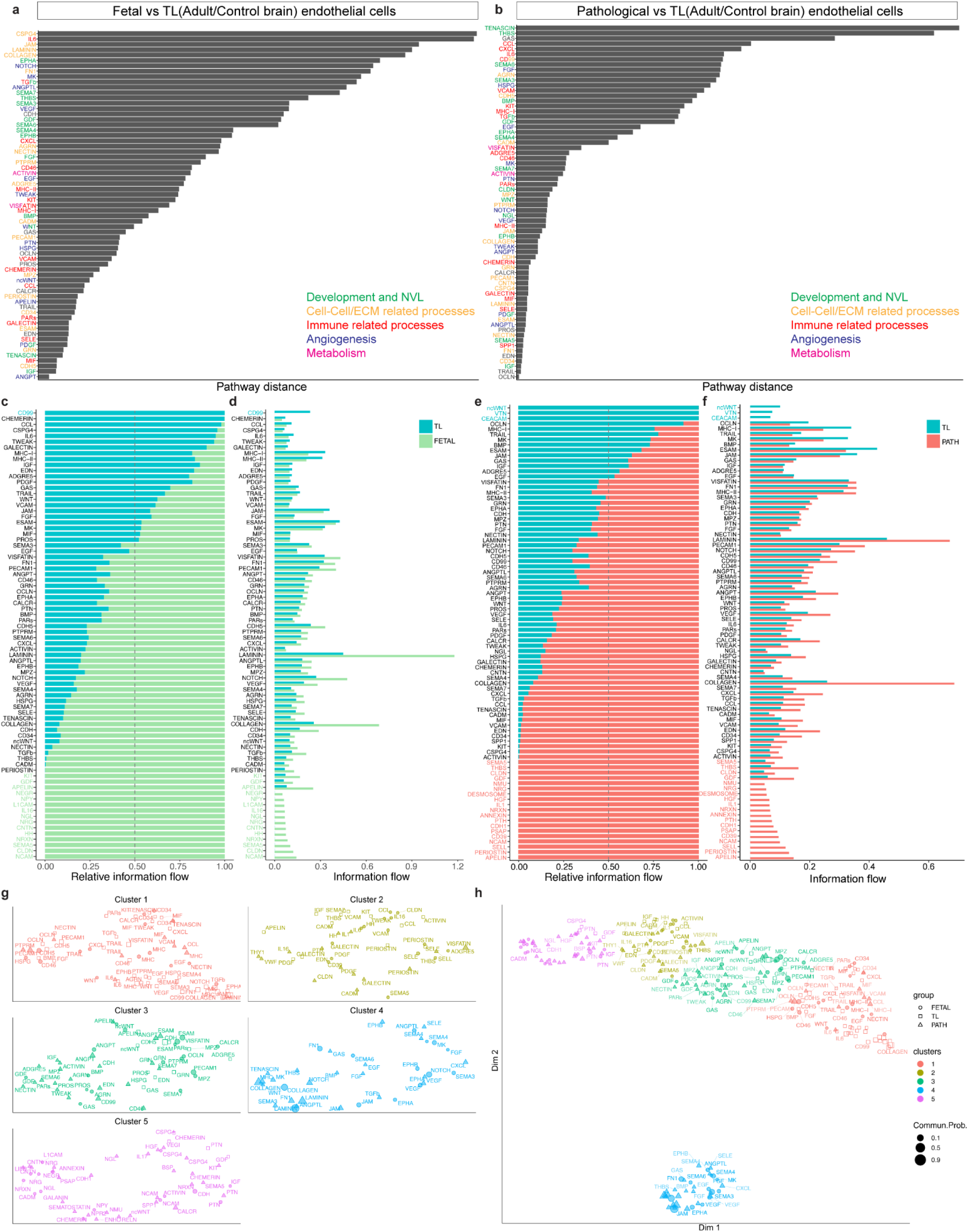
**a,b,** Barplot showing pathway distance; the overlapping signaling pathways between fetal and adult/control brain (**a**), pathological and adult/control brain (**b**) endothelial cells were ranked based on their pairwise Euclidean distance in the shared two-dimensional manifold. **c-f,** Barplot showing the relative (left panel) and absolute (right panel) information flow of all significant signaling pathways within the inferred networks between fetal and adult/control brain endothelial cells (**c,d**), and between pathological and adult/control brain endothelial cells (**e,f**). **g,h,** Jointly projecting and clustering signaling pathways from fetal, adult/control and pathological brain endothelial cells onto shared two-dimensional manifold according to their functional similarity of the inferred networks (**g**) and magnified view in (**h**). Circles, squares and triangle symbols represent the signaling networks from fetal, adult/control and pathological brains. Each circle/square/triangle represents the communication network of one signaling pathway. Symbol size is proportional to the total communication probability of that signaling network. Different colors represent different groups of signaling pathways.

**Extended Data Figure 18.**
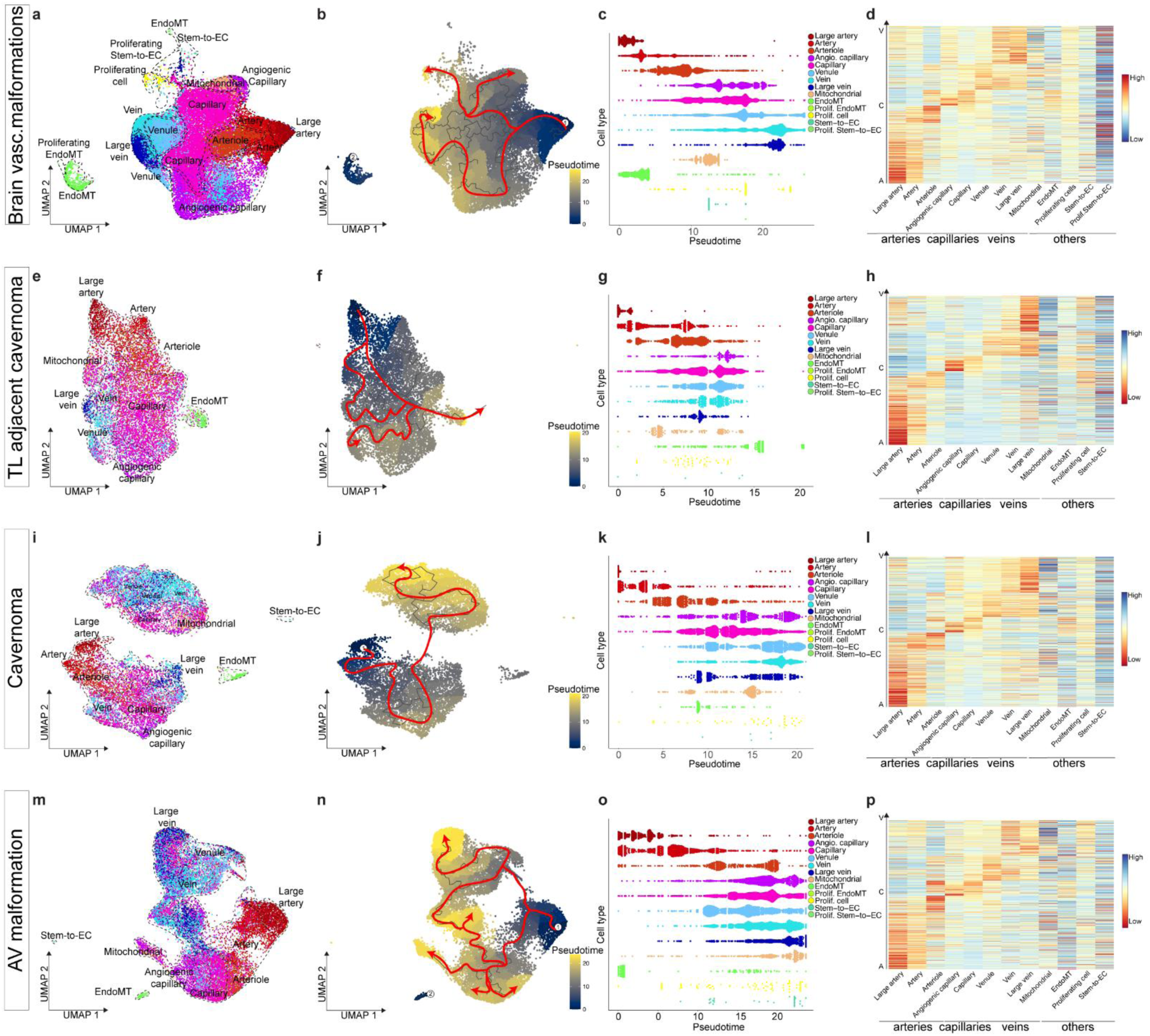
**a,b,f,i,j,m,n,** UMAP plots of human brain ECs isolated from all the brain vascular malformations (**a,b**), temporal lobes adjacent to cavernoma (**e,f**), cavernoma (**i,j**) and arteriovenous malformations (**m,n**) colored by AV specification (**a,e,i,m**) and by pseudotime (**b,f,j,n**). **c,g,k,o,** Pseudotime order of ECs color-coded according to AV specification from all the brain vascular malformations (**c**), temporal lobes adjacent to cavernoma (**g**), cavernoma (**k**) and arteriovenous malformations (**o**). **d,h,l,p,** Heatmap of adult/control brain ECs AV specification signature gene expression in human brain ECs isolated from all the brain vascular malformations (**d**), temporal lobes adjacent to cavernoma (**h**), cavernoma (**l**) and arteriovenous malformations (**p**).

**Extended Data Figure 19.**
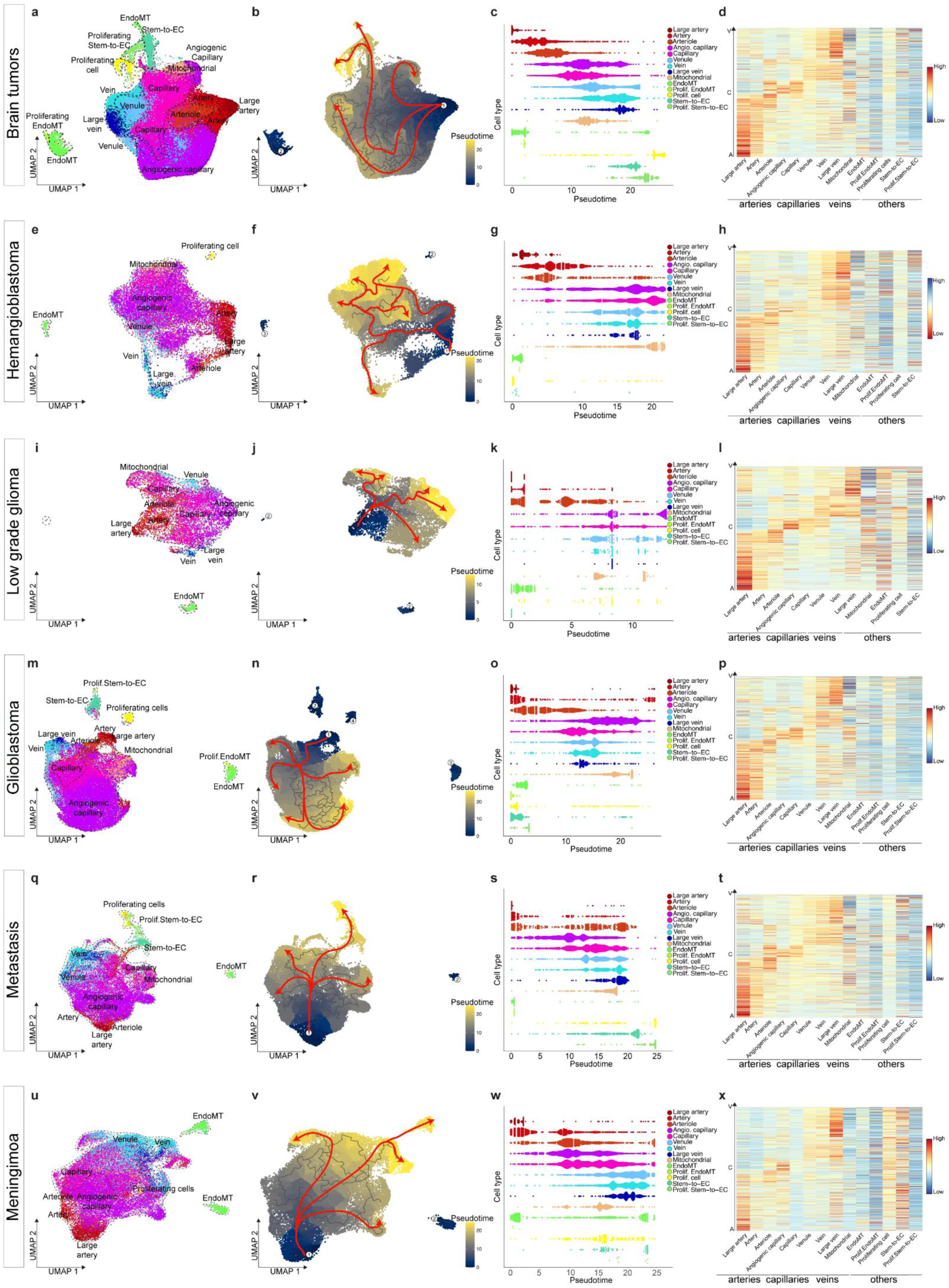
**a,b,e,f,i,j,m,n,q,r,u,v,** UMAP plots of human brain ECs isolated from all the brain tumors (**a,b**), hemangioblastoma (**e,f**), low-grade glioma (**i,j**) glioblastoma (**m,n**), metastasis (**q,r**) and meningioma (**u,v**) colored by AV specification (**a,e,i,m,q,u**) and by pseudotime (**b,f,j,n,r,v**). **c,g,k,o,s,w,** Pseudotime order of ECs color-coded according to AV specification from all the brain tumors (**c**), hemangioblastoma (**g**), low-grade glioma (**k**) glioblastoma (**o**), metastasis (**s**) and meningioma (**w**). **d,h,l,p,t,x,** Heatmap of adult/control brain ECs AV specification signature gene expression in human brain ECs isolated from all the brain tumors (**d**), hemangioblastoma (**h**), low-grade glioma (**l**) glioblastoma (**p**), metastasis (**t**) and meningioma (**x**).

**Extended Data Figure 20.**
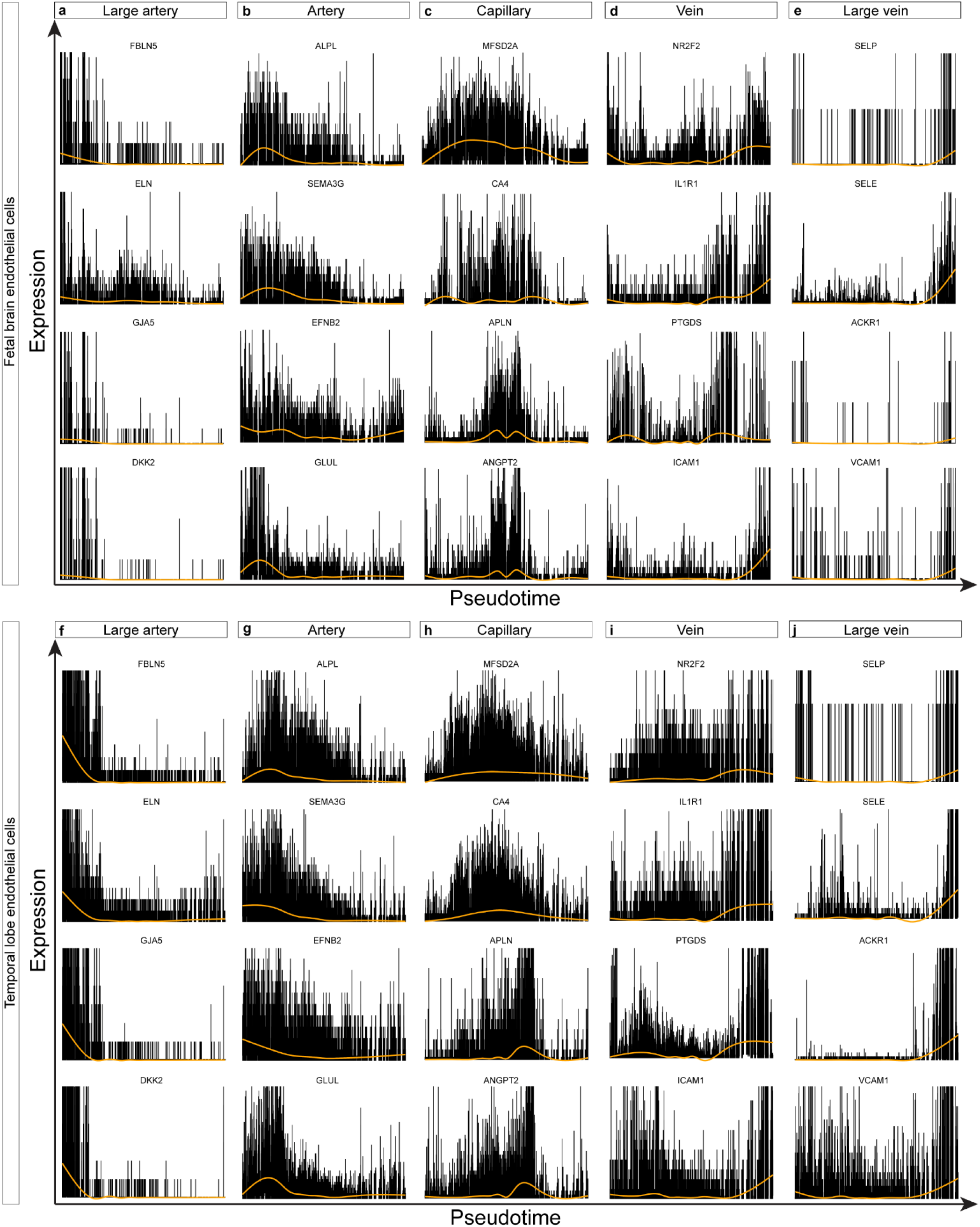
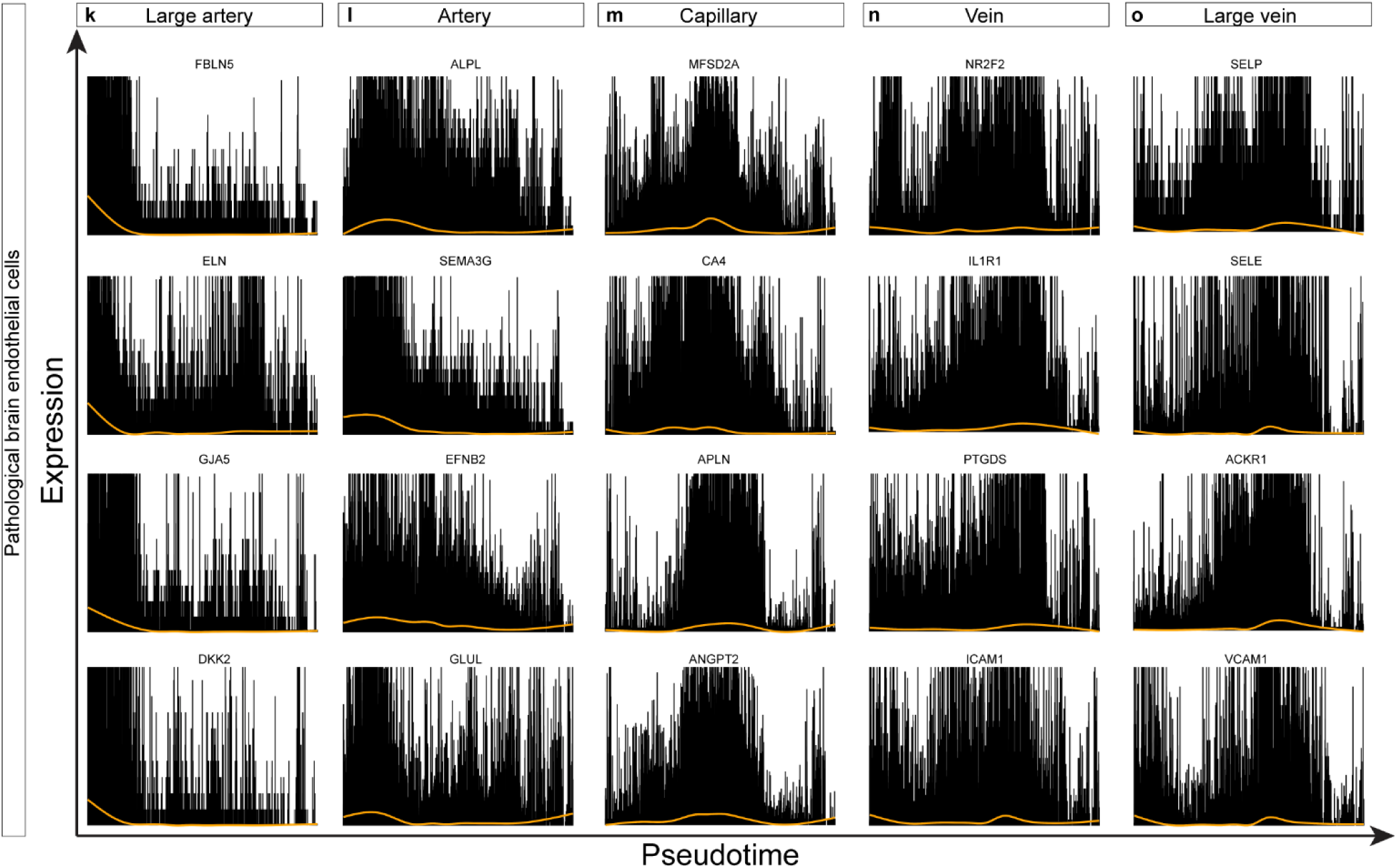
**a-j,** Gene expression of the indicated AV specification markers along the pseudotime trajectory in fetal brain (**a-e**), adult/control brain (**f-g**), and pathological brain (**k-o**) ECs. Spline (orange) and density of black lines (counts) correspond with average expression levels.

**Extended Data Figure 21.**
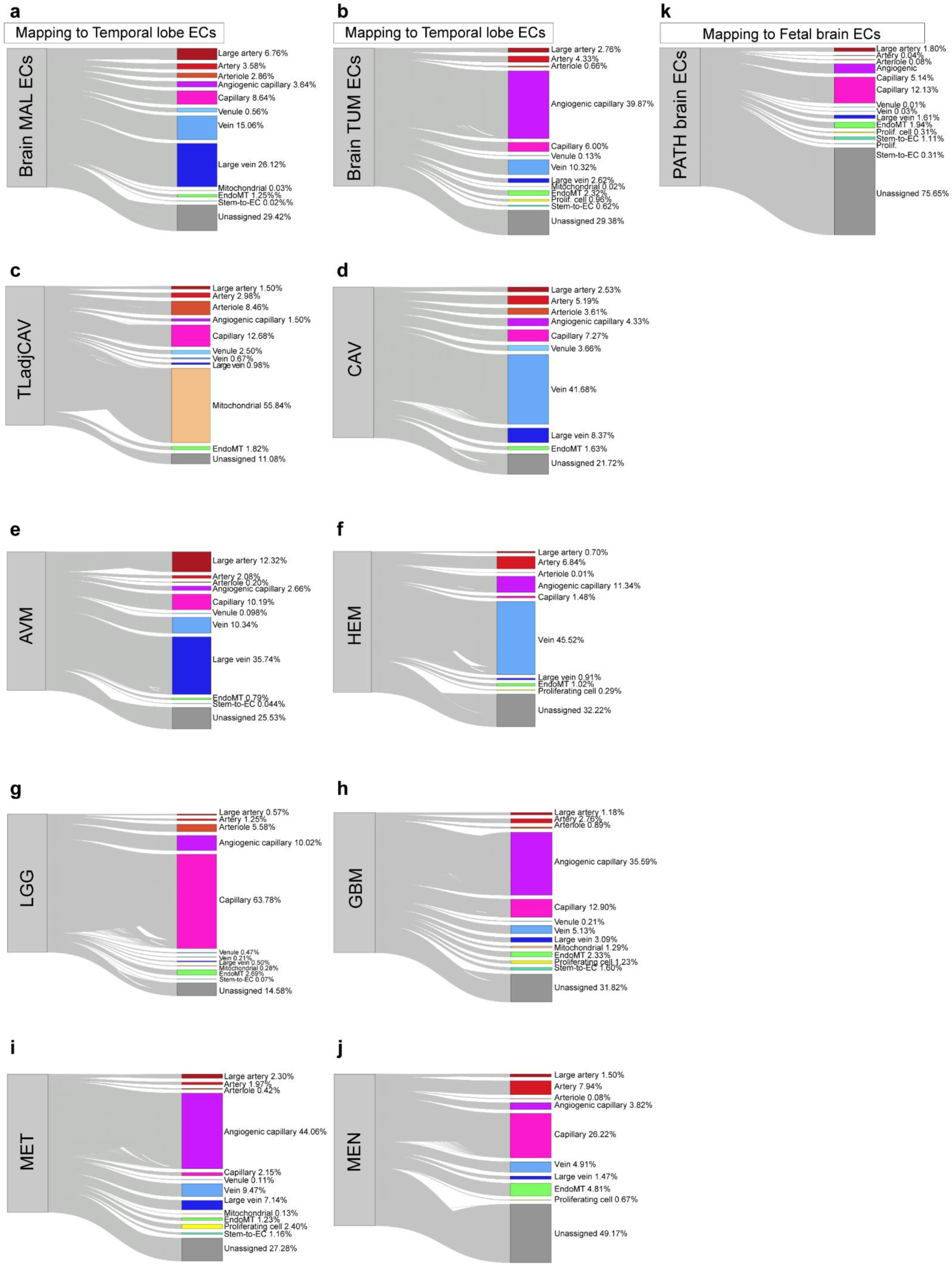
**a-j,** Sankey plot showing the predicted annotation of the ECs of the indicated entities as mapped to adult/control brain (TL) ECs. **k,** Sankey plot showing the predicted annotation of pathological ECs as mapped to fetal brain ECs. Unassigned cells are indicated in grey.

**Extended Data Figure 22.**
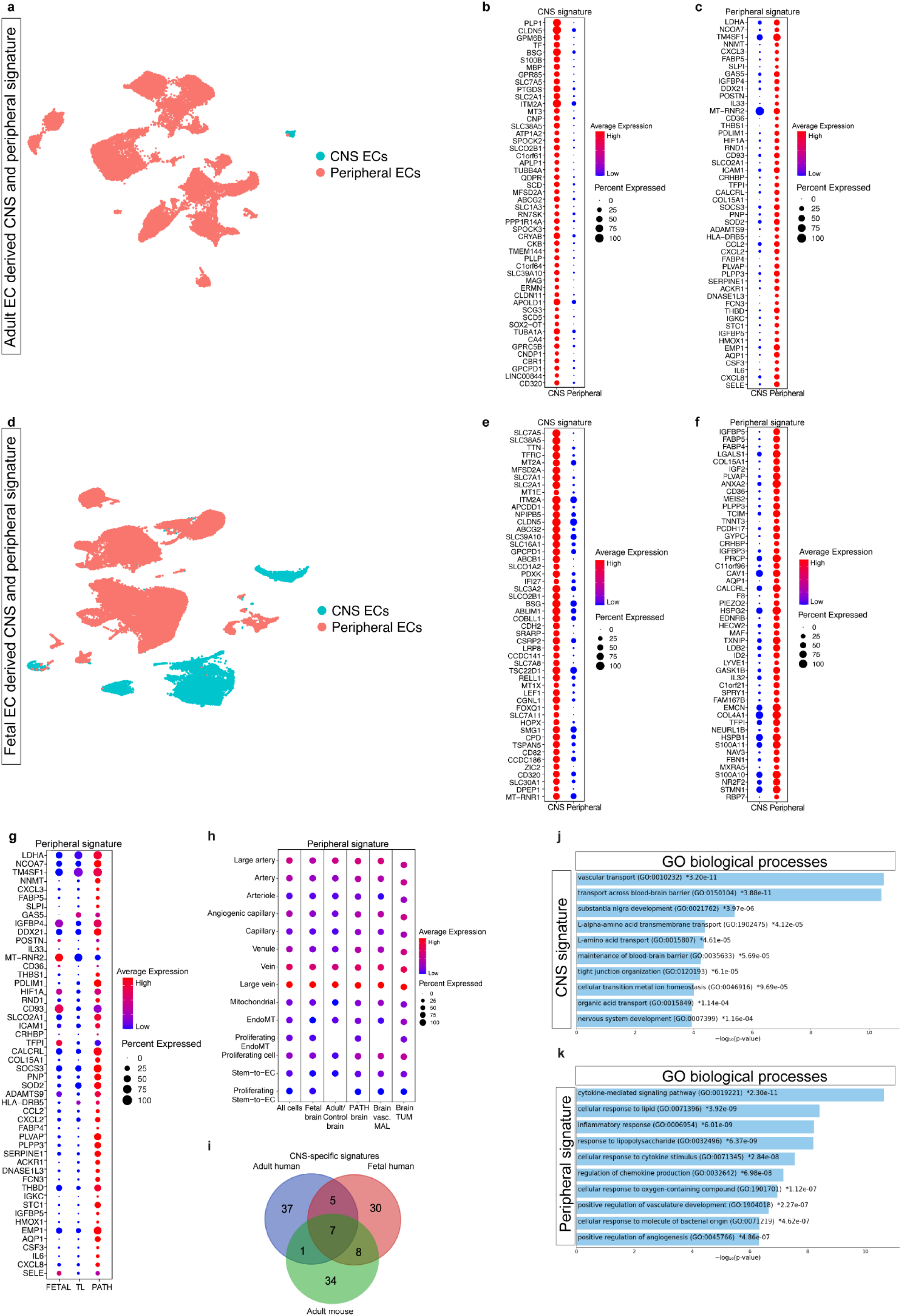
**a,d,** UMAP plot of adult ECs from Han et al., *Nature* 2020^25^ (**a**) and fetal ECs (**d**) colored by CNS and peripheral identity. **b,c,** Dotplot heatmap of the adult endothelial CNS signature (**b**) and peripheral signature (**c**) genes enriched in the respective EC populations. **e,f,** Dotplot heatmap of the fetal endothelial CNS signature (**e**). **g,** Dotplot heatmap of the adult peripheral EC signature genes in fetal, adult/control (temporal lobes) and pathological brain ECs. **h,** Dotplot heatmaps of CNS signature at the level of AV specification for the indicated entities. Color scale: red, high expression; blue, low expression, whereas the dot size represents the percentage expression within the indicated entity. **i,** Venn diagram showing the overlap between the top 50 CNS signature genes obtained from human adult, human fetal and mouse ECs. **j,** Enrichment analysis of human adult CNS endothelial signature showing the top 10 enriched gene ontology biological process (GOBP) genesets. **i,** Enrichment analysis of human adult peripheral endothelial signature. The top 10 enriched GOBP genesets.

**Extended Data Figure 23.**
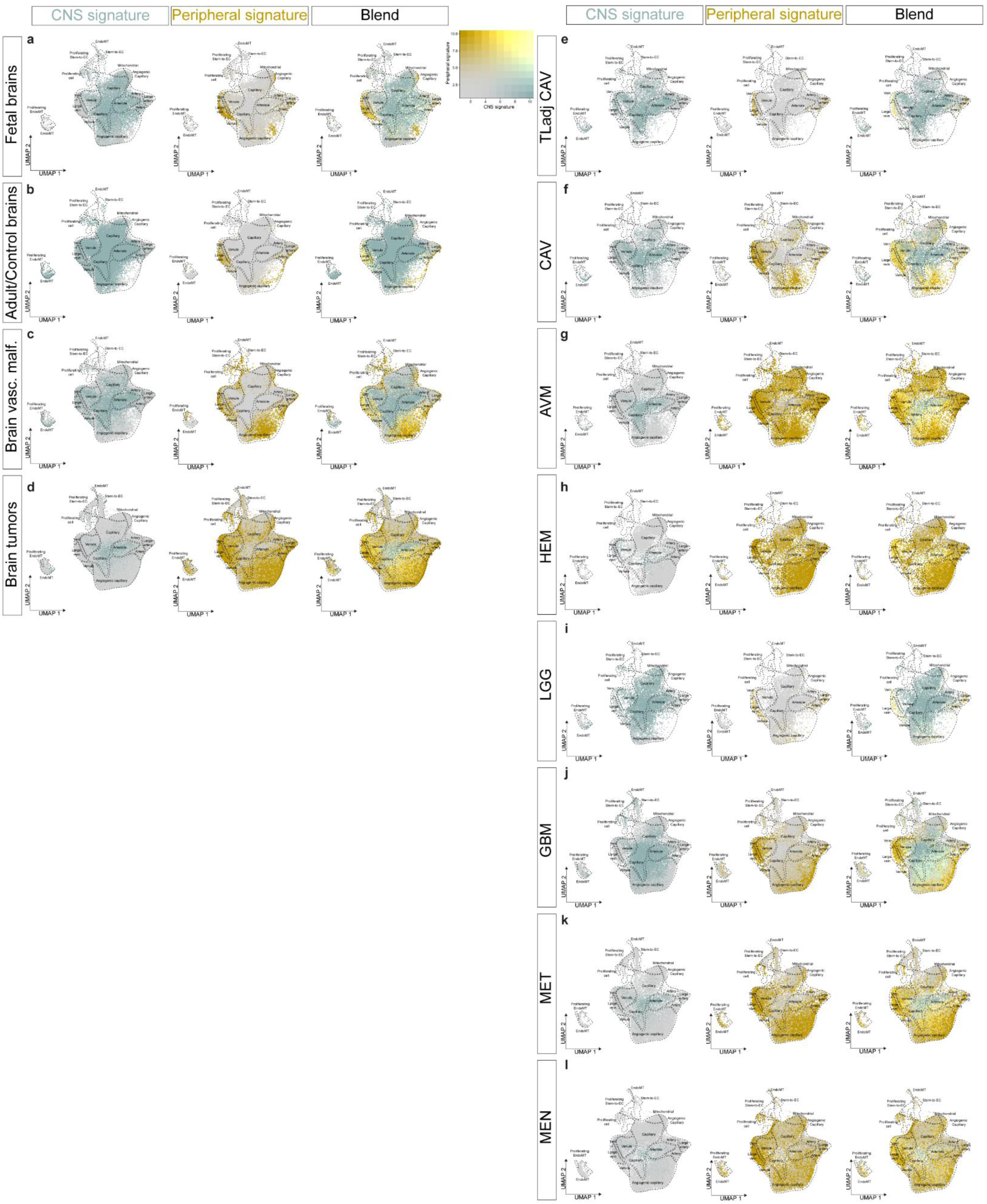
**a-l**, UMAP plots of ECs from fetal brains (**a**), adult/control brains (**b**), brain vascular malformations (**c**), brain tumors (**d**), temporal lobe adjacent to cavernoma (**e**), cavernoma (**f**), arteriovenous malformations (**g**), hemangioblastoma (**h**), low-grade glioma (**i**), glioblastoma (**j**), metastasis (**k**) and meningioma (**l**). Plots are color-coded for adult EC CNS signature (green, left panel), adult EC peripheral signature (yellow, middle panel), and blend of both signatures (right panel).

**Extended Data Figure 24.**
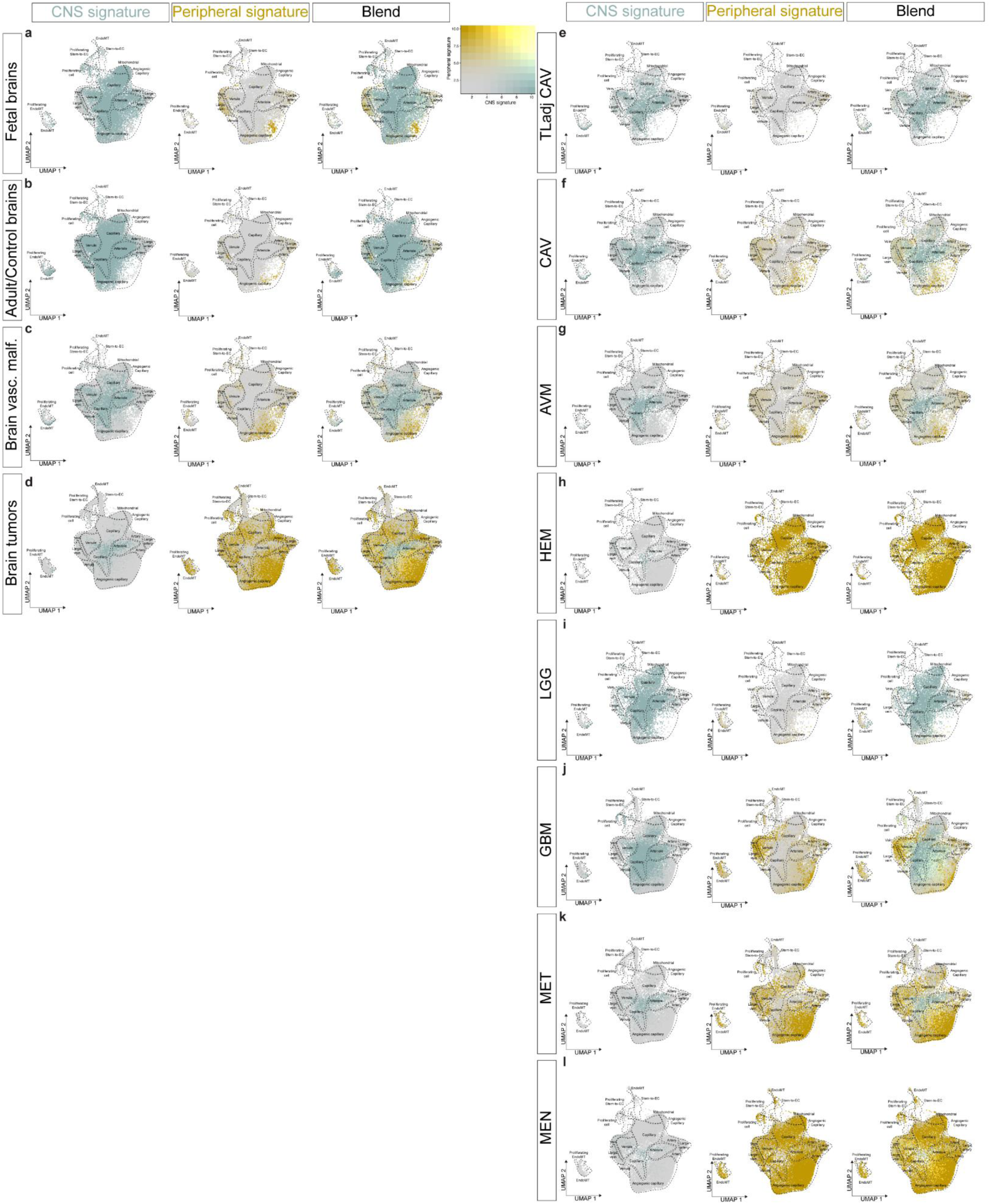
**a-l**, UMAP plots of ECs from fetal brains (**a**), adult/control brains (**b**), brain vascular malformations (**c**), brain tumors (**d**), temporal lobe adjacent to cavernoma (**e**), cavernoma (**f**), arteriovenous malformations (**g**), hemangioblastoma (**h**), low-grade glioma (**i**), glioblastoma (**j**), metastasis (**k**) and meningioma (**l**). Plots are color-coded for fetal EC CNS signature (green, left panel), fetal EC peripheral signature (yellow, middle panel), and blend of both signatures (right panel).

**Extended Data Figure 25.**
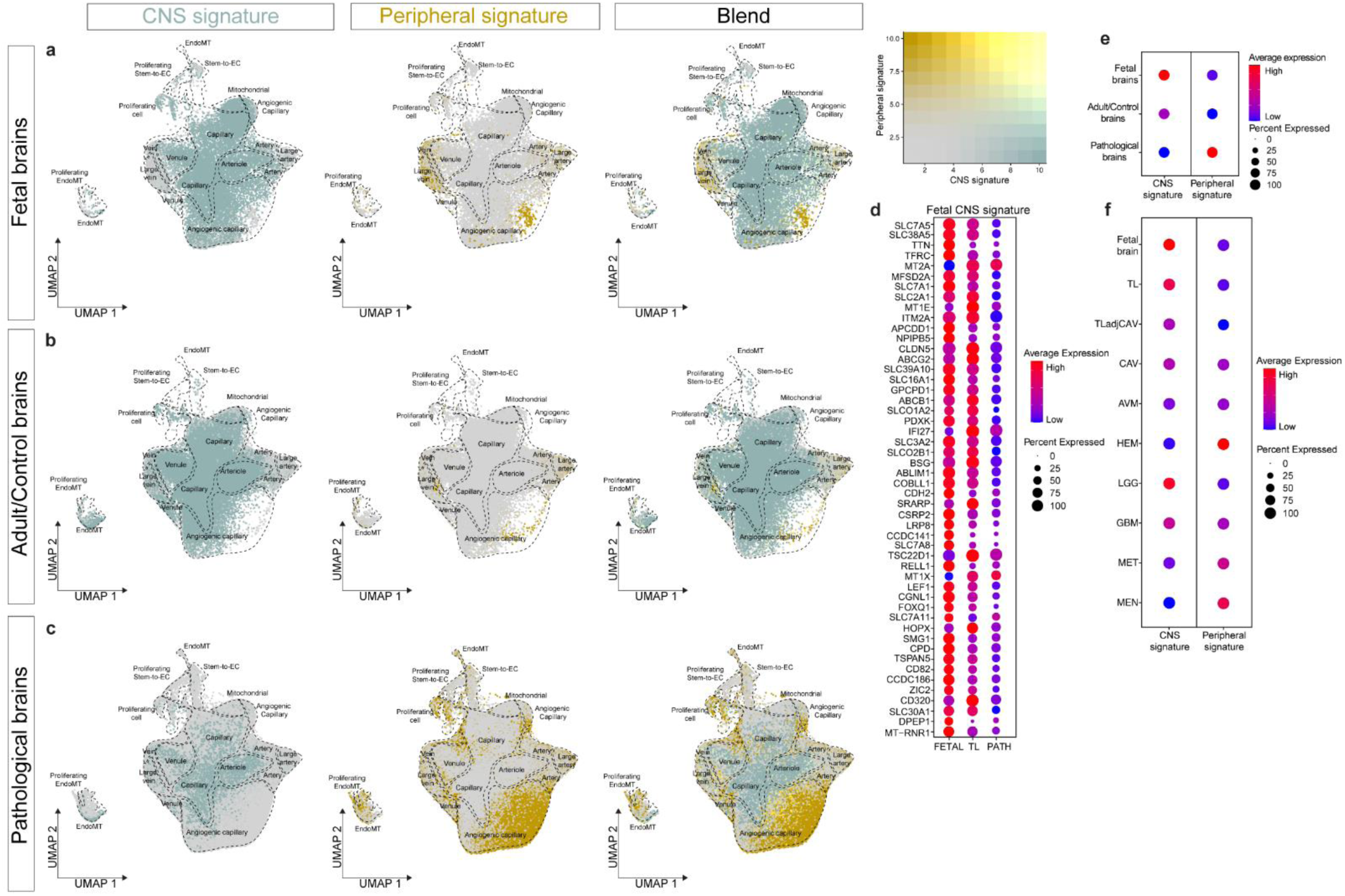
**a-c**, UMAP plots of the ECs from fetal brains (**a**), adult/control brains (**b**), and pathological brains (**c**). Plots are color-coded for fetal ECs CNS signature (green, left panel), fetal ECs peripheral signature (yellow, middle panel) and a blend of both signatures (right panel). **d,** Dotplot heatmaps of fetal CNS signature genes expression in fetal brain, adult/control brains (temporal lobes) and pathological brain ECs. **e,f,** Dotplot heatmaps of the fetal CNS and peripheral signature expression in fetal brain, adult/control brain, and pathological brain ECs (**e**) and in each individual entity (**f**). Color scale: red, high expression; blue, low expression, whereas the dot size represents the percentage expression within the indicated entity.

**Extended Data Figure 26.**
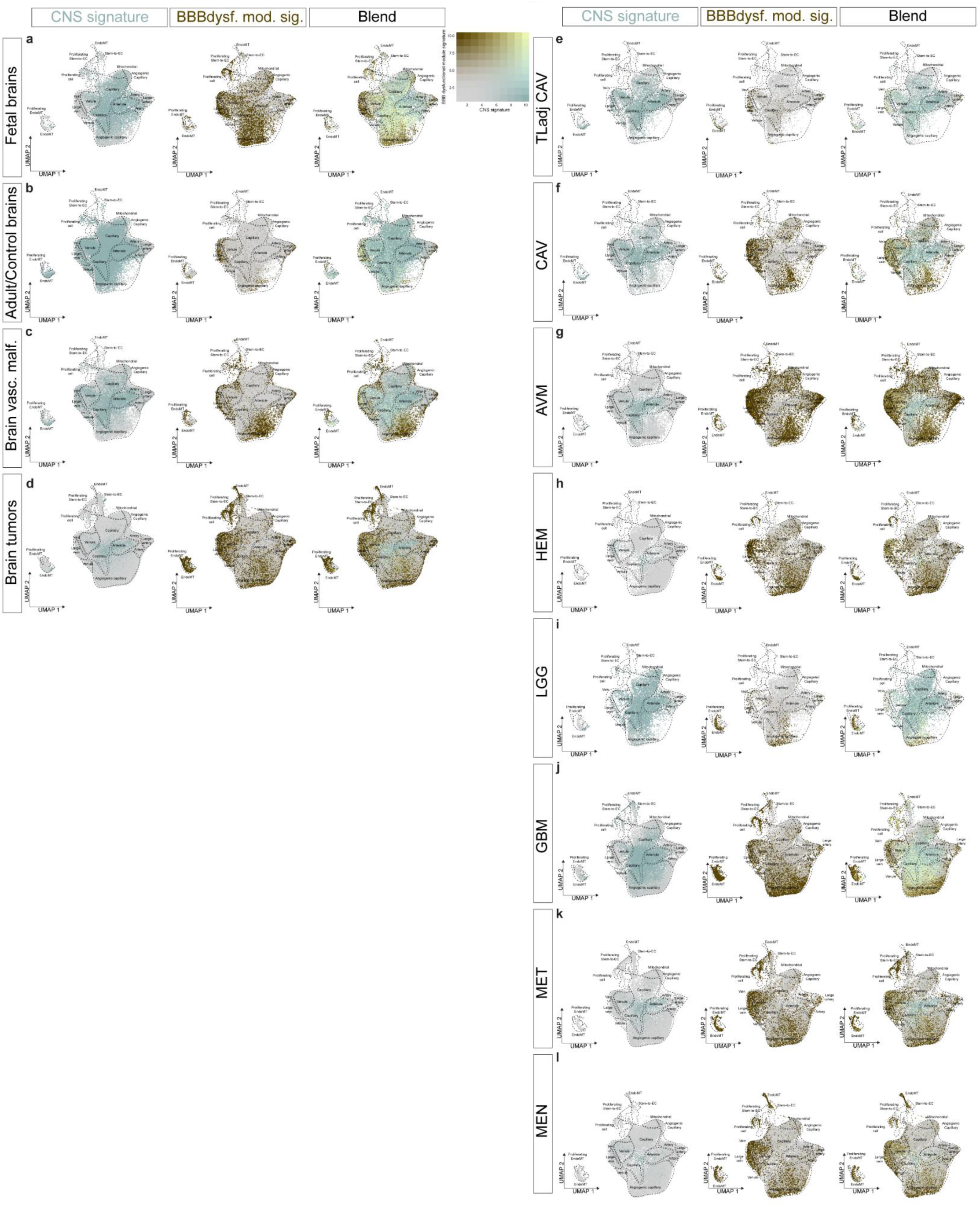
**a-l**, UMAP plots of ECs from fetal brains (**a**), adult/control brains (**b**), brain vascular malformations (**c**), brain tumors (**d**), temporal lobe adjacent to cavernoma (**e**), cavernoma (**f**), arteriovenous malformations (**g**), hemangioblastoma (**h**), low-grade glioma (**i**), glioblastoma (**j**), metastasis (**k**) and meningioma (**l**). Plots are color-coded for adult EC CNS signature (green, left panel), BBB dysfunction module signature (brown, middle panel), and blend of both signatures (right panel).

**Extended Data Figure 27.**
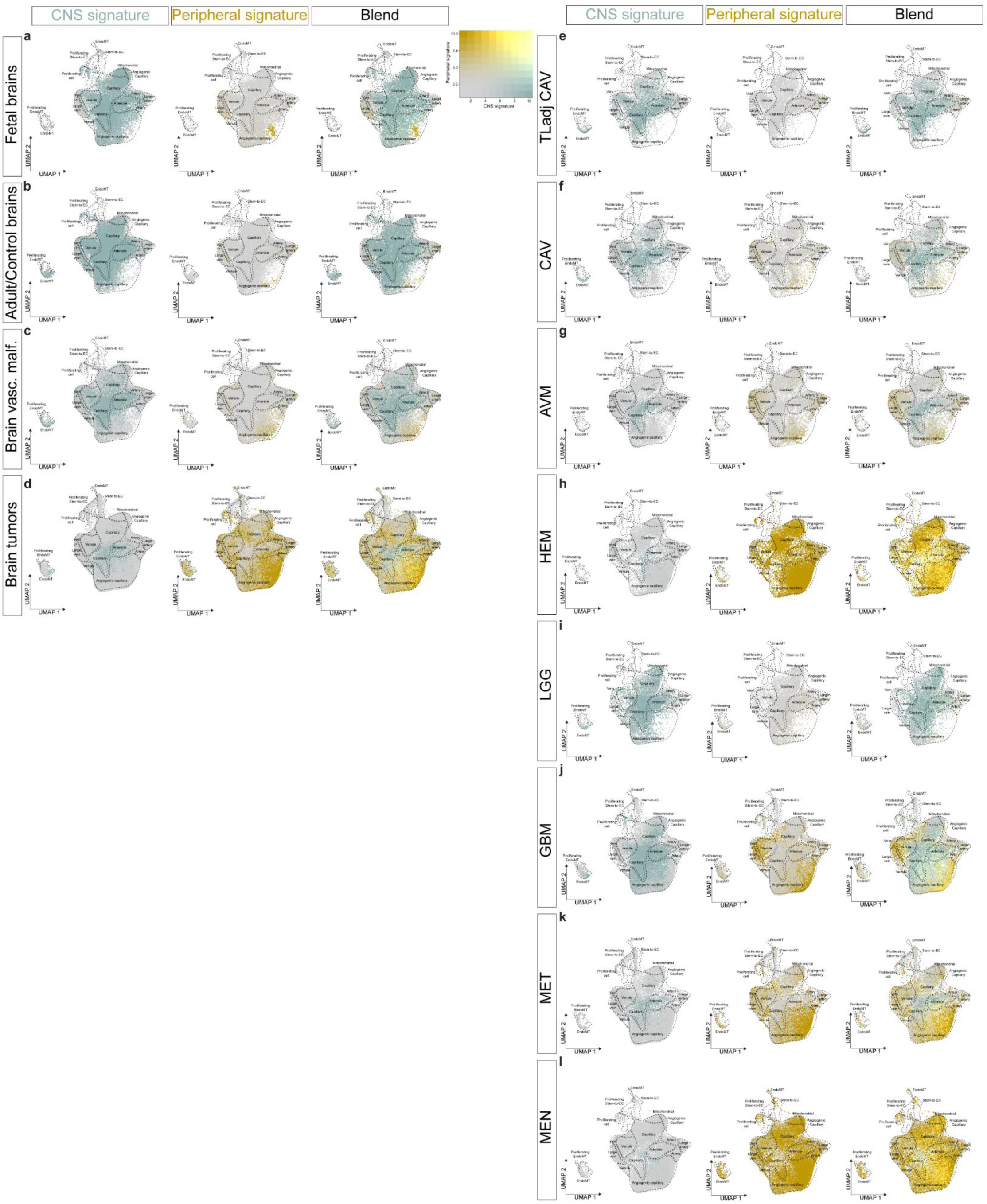
**a-l**, UMAP plots of ECs from fetal brains (**a**), adult/control brains (**b**), brain vascular malformations (**c**), brain tumors (**d**), temporal lobe adjacent to cavernoma (**e**), cavernoma (**f**), arteriovenous malformations (**g**), hemangioblastoma (**h**), low-grade glioma (**i**), glioblastoma (**j**), metastasis (**k**) and meningioma (**l**). Plots are color-coded for mouse EC CNS signature (green, left panel), mouse EC peripheral signature (yellow, middle panel), and blend of both signatures (right panel).

**Extended Data Figure 28.**
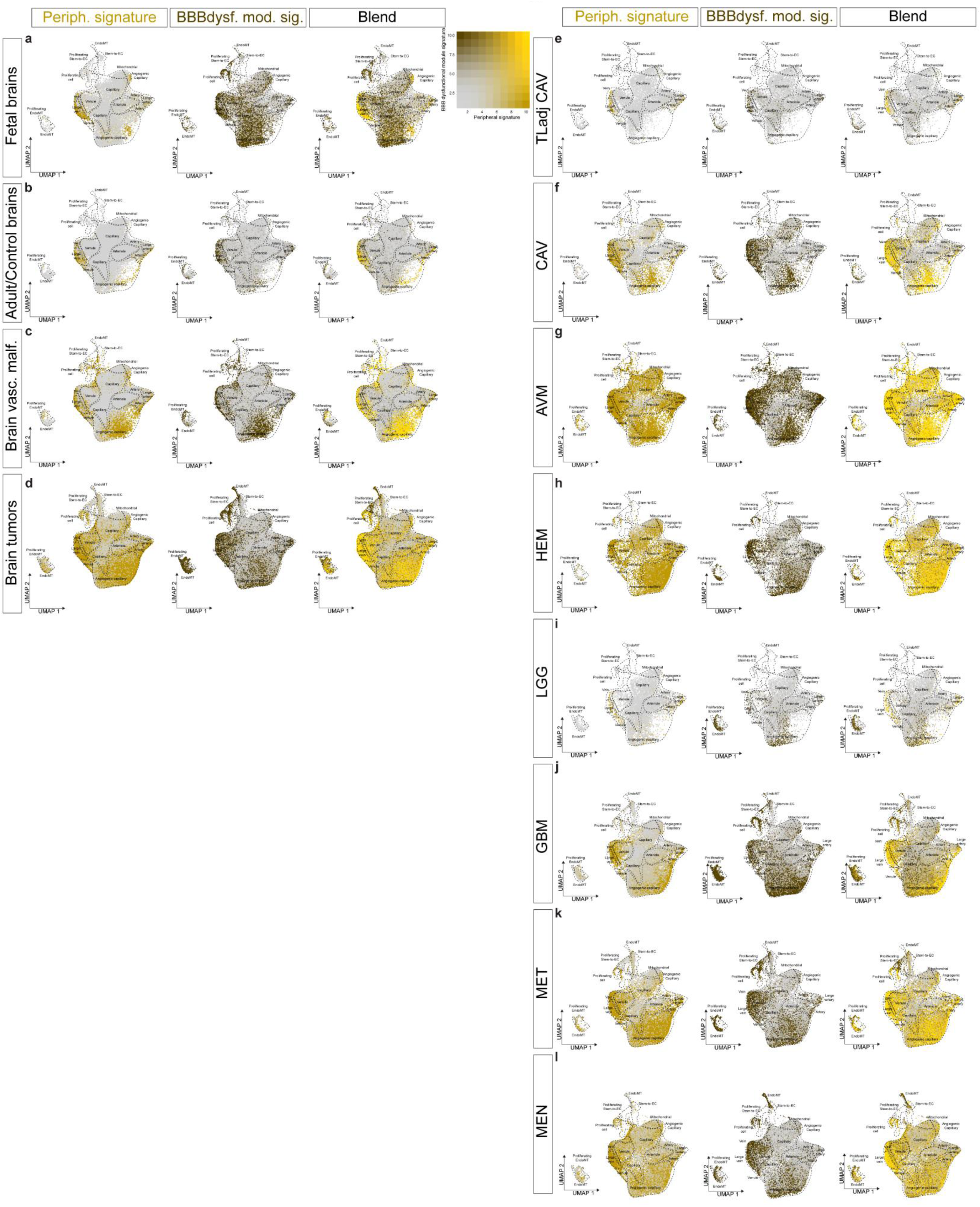
**a-l**, UMAP plots of ECs from fetal brains (**a**), adult/control brains (**b**), brain vascular malformations (**c**), brain tumors (**d**), temporal lobe adjacent to cavernoma (**e**), cavernoma (**f**), arteriovenous malformations (**g**), hemangioblastoma (**h**), low-grade glioma (**i**), glioblastoma (**j**), metastasis (**k**) and meningioma (**l**). Plots are color-coded for adult human peripheral EC signature (yellow, left panel), BBB dysfunction module signature (brown, middle panel), and blend of both signatures (right panel).

**Extended Data Figure 29.**
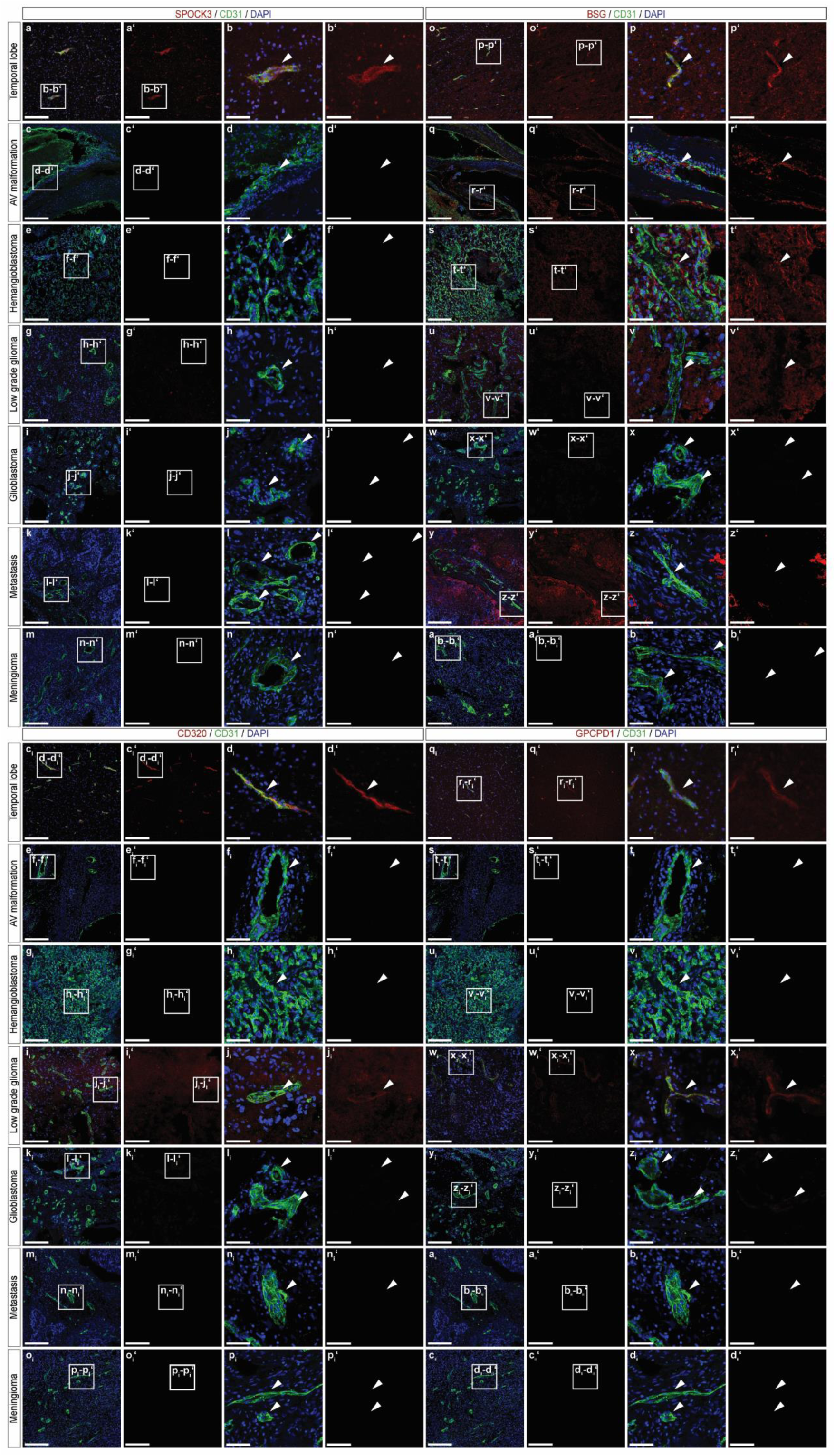
**a-d_ii_‘,** Immunofluorescence (IF) imaging of tissue sections from the indicated entities, stained for *SPOCK3* (red; **a-n‘**), *BSG* (red; **o-b_i_‘**), *CD320* (red; **c_i_-p_i_‘**), *GPCPD1* (red; **q_i_-d_ii_‘**) and *CD31* (green). Nuclei are stained with DAPI (blue). Boxed area is magnified on the right; arrowheads indicate vascular structures in the different tissues. Scale bars: 200μm in overviews, 50μm in zooms.

**Extended Data Figure 30.**
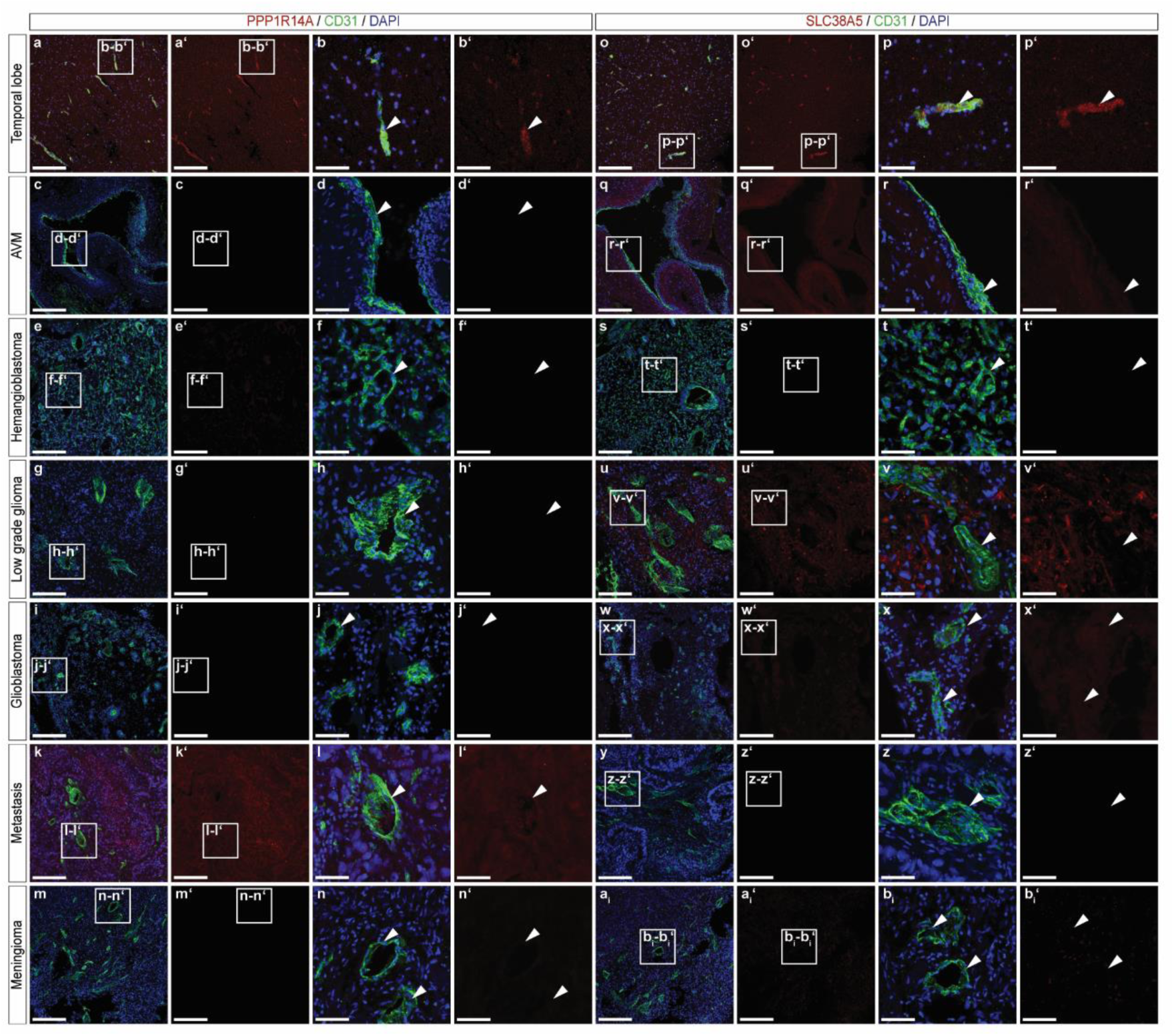
**a-b_i_‘,** Immunofluorescence (IF) imaging of tissue sections from the indicated entities, stained for *PPP1R141* (red; **a-n‘**), *SLC38A5* (red; **o-b_i_‘**) and CD31 (green). Nuclei are stained with DAPI (blue). Boxed area is magnified on the right; arrowheads indicate vascular structures in the different tissues. Scale bars: 200μm in overviews, 50μm in zooms.

**Extended Data Figure 31.**
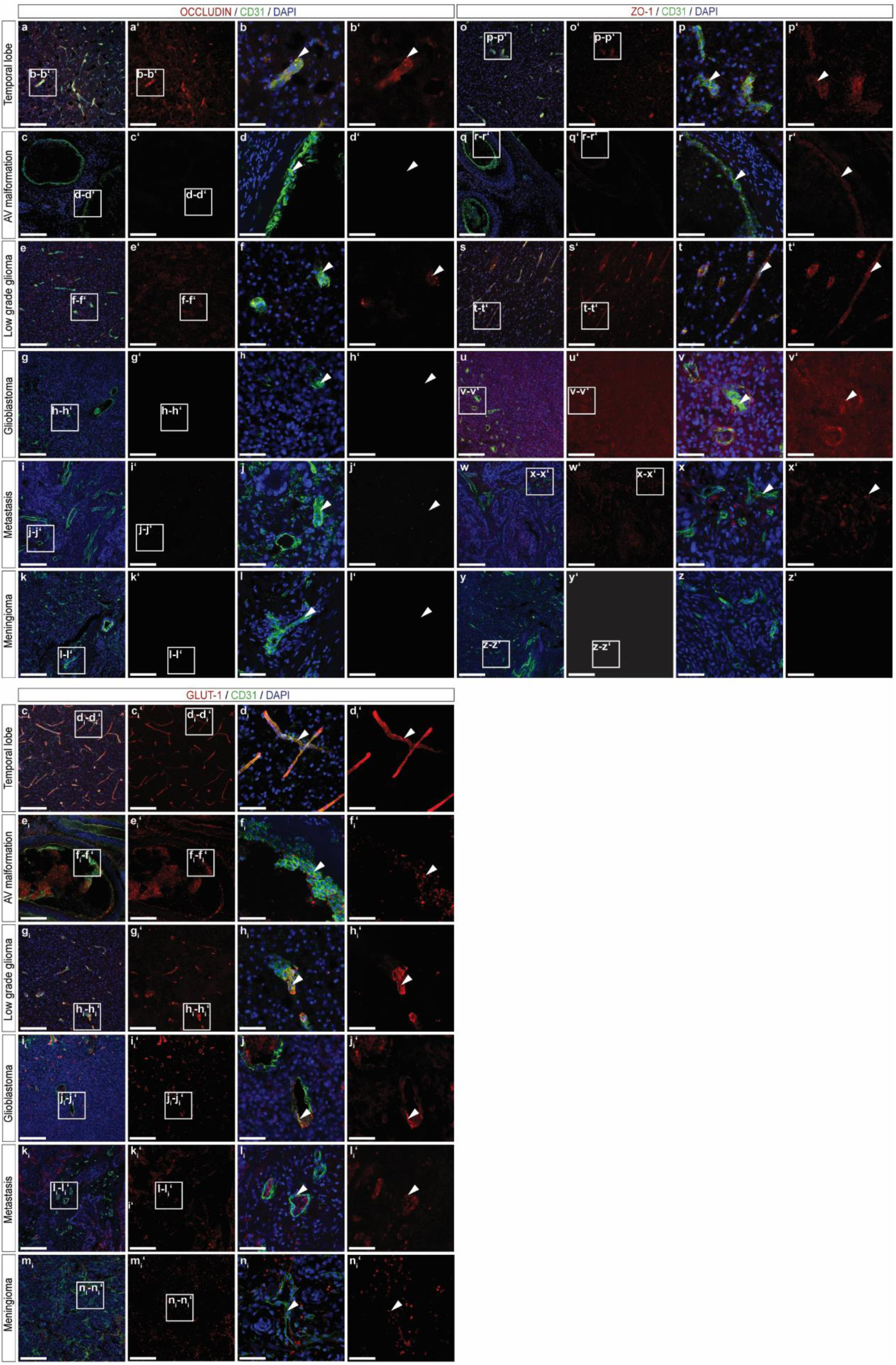
**a-n_i_‘**, Immunofluorescence (IF) imaging of tissue sections from the indicated entities, stained for *OCLN* (red; **a-l‘**), *ZO1* (red; **o-z‘**), *GLUT1* (red; **c_i_-n_i_‘**) and *CD31* (green). Nuclei are stained with DAPI (blue). Boxed area is magnified on the right; arrowheads indicate vascular structures in the different tissues. Scale bars: 200μm in overviews, 50μm in zooms.

**Extended Data Figure 32.**
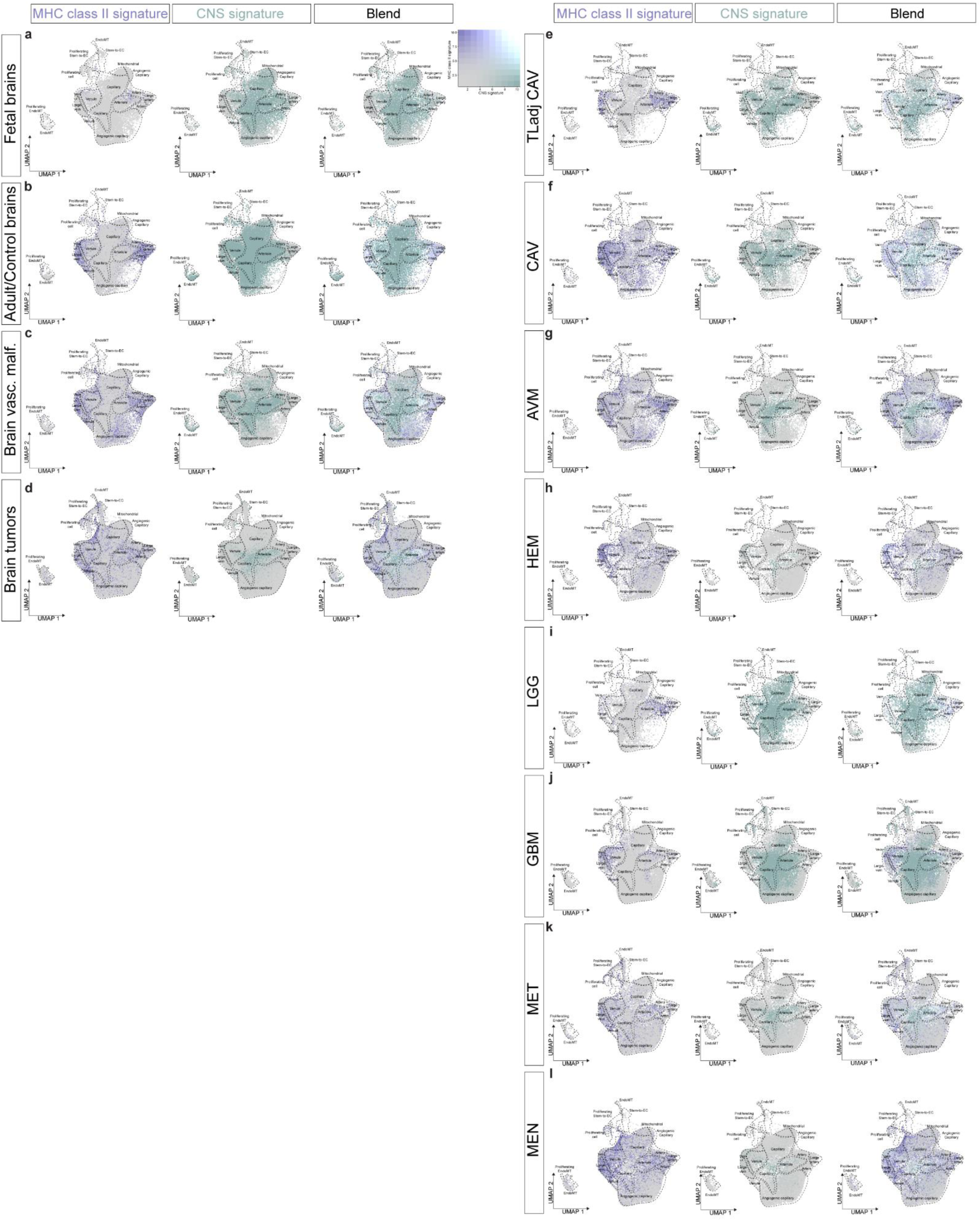
**a-l**, UMAP plots of ECs from fetal brains (**a**), adult/control brains (**b**), brain vascular malformations (**c**), brain tumors (**d**), temporal lobe adjacent to cavernoma (**e**), cavernoma (**f**), arteriovenous malformations (**g**), hemangioblastoma (**h**), low-grade glioma (**i**), glioblastoma (**j**), metastasis (**k**) and meningioma (**l**). Plots are color-coded for MHC class II signature (violet, left panel), CNS signature (green, middle panel), and blend of both signatures (right panel).

**Extended Data Figure 33.**
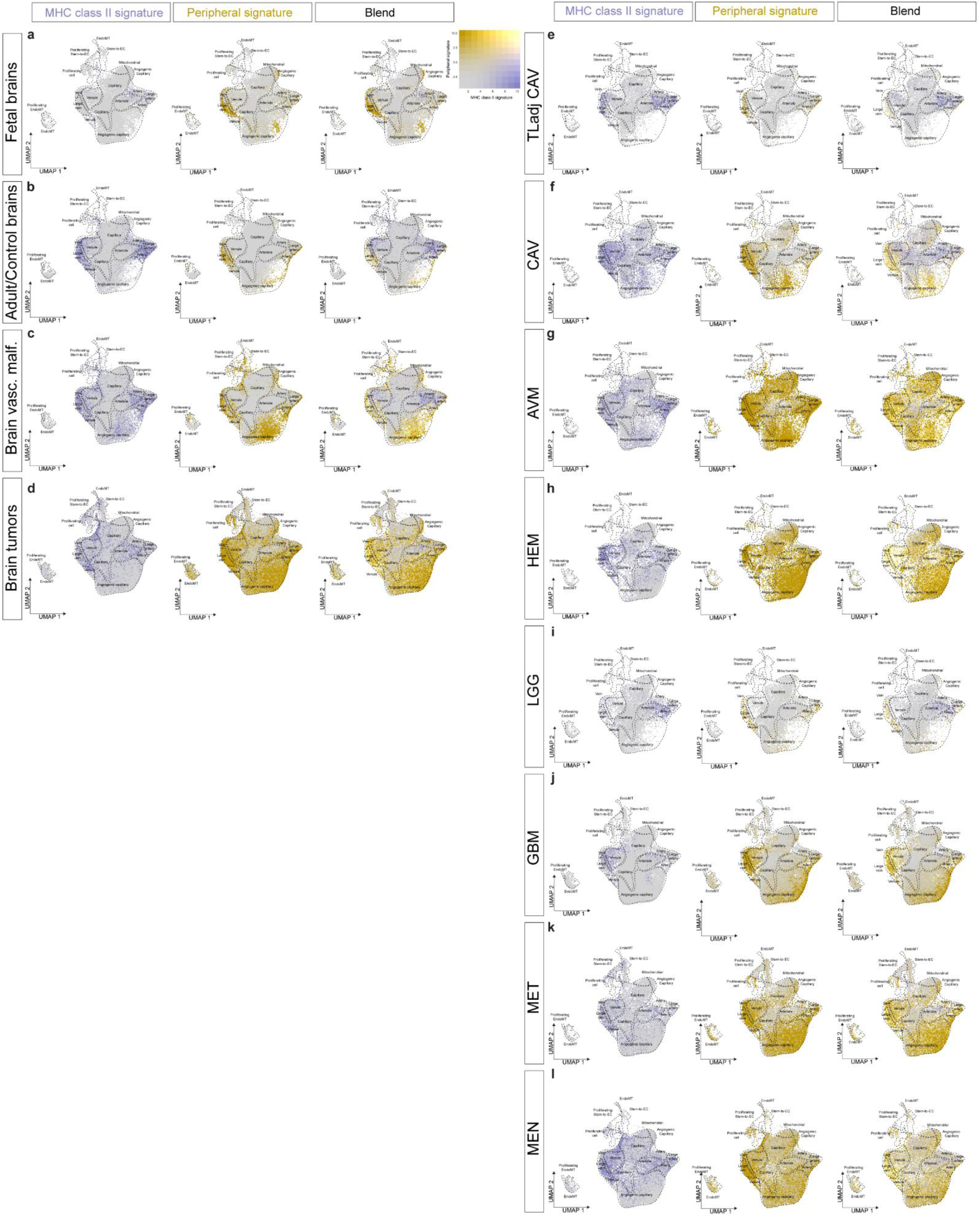
**a-l**, UMAP plots of ECs from fetal brains (**a**), adult/control brains (**b**), brain vascular malformations (**c**), brain tumors (**d**), temporal lobe adjacent to cavernoma (**e**), cavernoma (**f**), arteriovenous malformations (**g**), hemangioblastoma (**h**), low-grade glioma (**i**), glioblastoma (**j**), metastasis (**k**) and meningioma (**l**). Plots are color-coded for MHC class II signature (violet, left panel), peripheral signature (yellow, middle panel), and blend of both signatures (right panel).

**Extended Data Figure 34.**
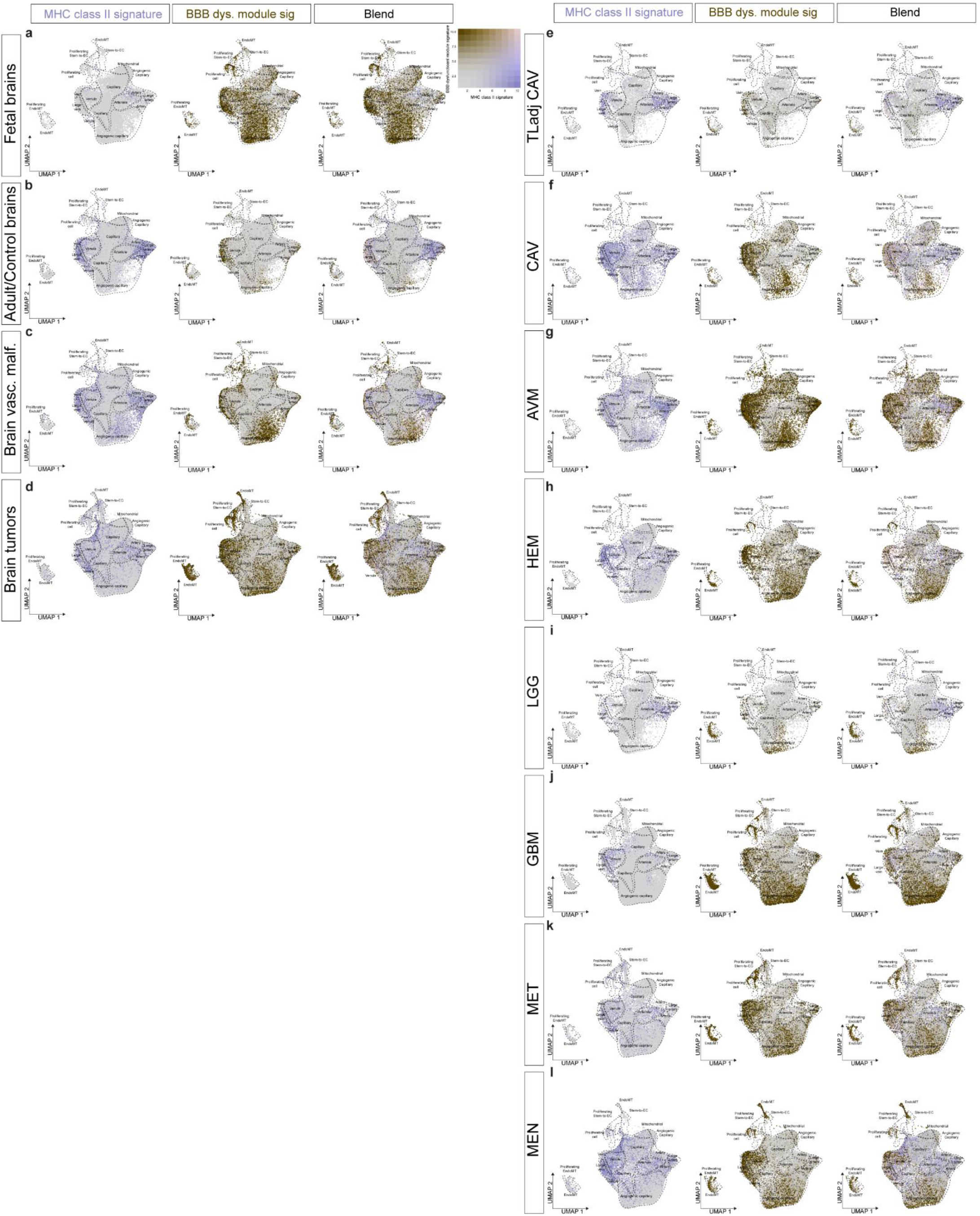
**a-l**, UMAP plots of ECs from fetal brains (**a**), adult/control brains (**b**), brain vascular malformations (**c**), brain tumors (**d**), temporal lobe adjacent to cavernoma (**e**), cavernoma (**f**), arteriovenous malformations (**g**), hemangioblastoma (**h**), low-grade glioma (**i**), glioblastoma (**j**), metastasis (**k**) and meningioma (**l**). Plots are color-coded for MHC class II signature (violet, left panel), BBB dysfunction module signature (brown, middle panel), and blend of both signatures (right panel).

**Extended Data Figure 35.**
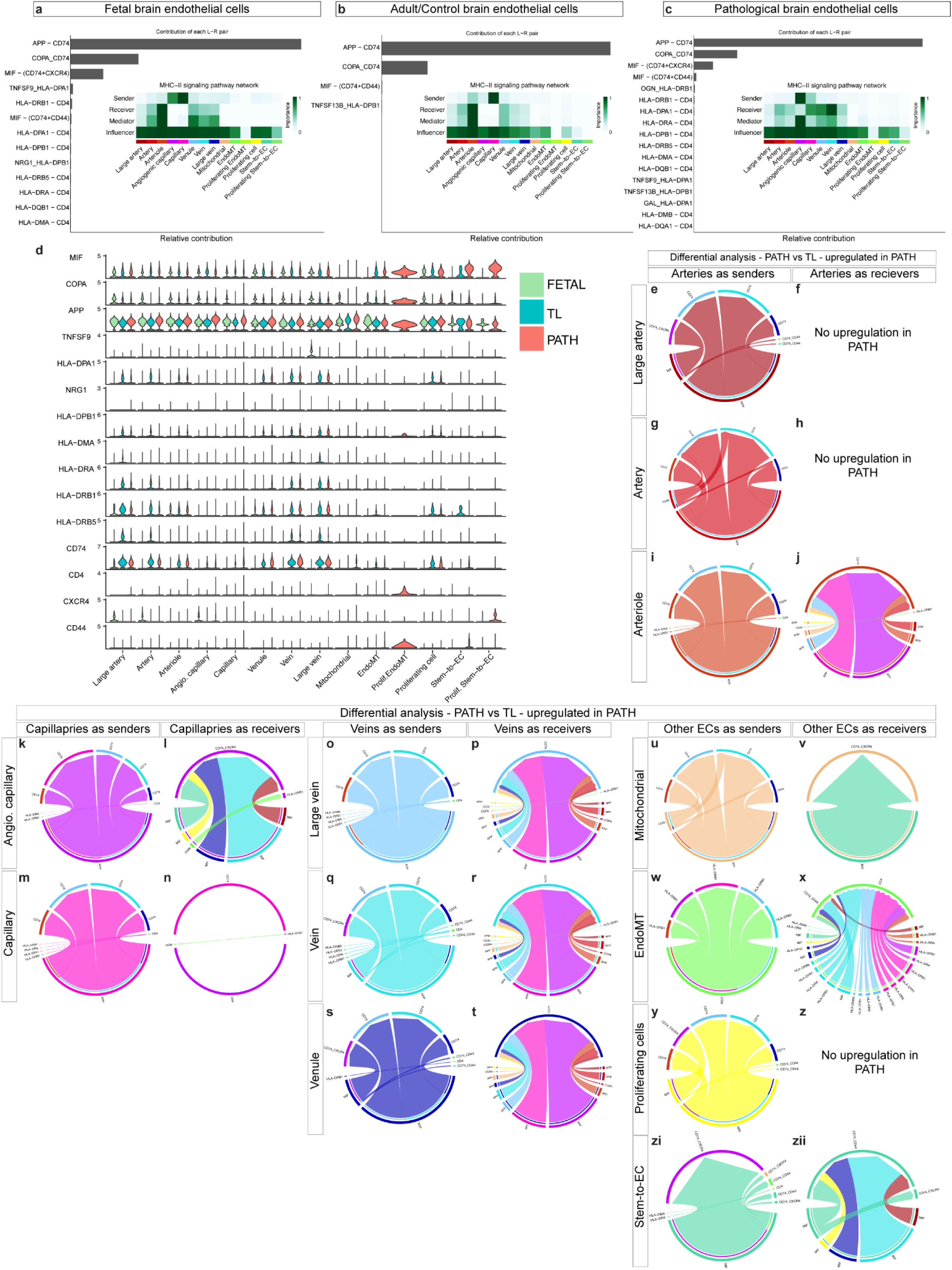
**a-c,** Bar plots showing the relative contribution of each ligand-receptor pair to the overall communication network of MHC class II signaling pathway and heatmaps showing the relative importance of each cell type based on the computed network centrality in the fetal (**a**), adult/control (**b**) and pathological (**c**) brain ECs. **d,** Violin plot showing the expression of the main ligand and receptors participating in MHC class II signaling interactions in EC subtypes of fetal, adult control and pathological brains. **e-z_ii_,** Visualization of the differential analysis - pathological brain ECs over adult/control brain ECs - of MHC class II signaling ligand-receptor pairs in the indicated EC subtypes (left panel as sender, and right panel as receiver). **e,g,i,** Chord/circos plots showing the upregulated MHC class II signaling in arteries as source and all EC clusters as targets (**e**, large arteries; **g**, arteries; **i**, arterioles). **f,h,j,** Chord/circos plots showing the upregulated MHC class II signaling in arteries as target and all EC clusters as source (**f**, large arteries for which there was no upregulation; **h**, arteries for which there was no upregulation; **j**, arterioles). **k,m,** Chord/circos plots showing the upregulated MHC class II signaling in capillaries as source and all EC clusters as targets (**k**, angiogenic capillaries; **m**, capillaries). **l,n,** Chord/circos plots showing the upregulated MHC class II signaling in capillaries as target and all EC clusters as source (**l**, angiogenic capillaries; **n**, capillaries). **o,q,s,** Chord/circos plots showing the upregulated MHC class II signaling in veins as source and all EC clusters as targets (**o**, large veins; **q**, veins; **t**, venules). **p,r,t,** Chord/circos plots showing the upregulated MHC class II signaling in veins as target and all EC clusters as source (**p**, large veins; **r**, veins; **t**, venules). **u,** Chord/circos plots showing the upregulated MHC class II signaling in mitochondrial EC subtype as source and all EC clusters as targets. **v,** Chord/circos plots showing the upregulated MHC class II signaling in mitochondrial EC subtype as target and all EC clusters as source. **w,** Chord/circos plots showing the upregulated MHC class II signaling in EndoMT as source and all EC clusters as targets. **x,** Chord/circos plots showing the upregulated MHC class II signaling in EndoMT as target and all EC clusters as source. **y,** Chord/circos plots showing the upregulated MHC class II signaling in proliferating ECs as source and all EC clusters as targets. **z,** There was no upregulated MHC class II signaling in proliferating ECs as target and all EC clusters as source. **zi,** Chord/circos plots showing the upregulated MHC class II signaling in stem-to-EC as source and all EC clusters as targets. **x,** Chord/circos plots showing the upregulated MHC class II signaling in stem-to-EC as target and all EC clusters as source.

**Extended Data Figure 36.**
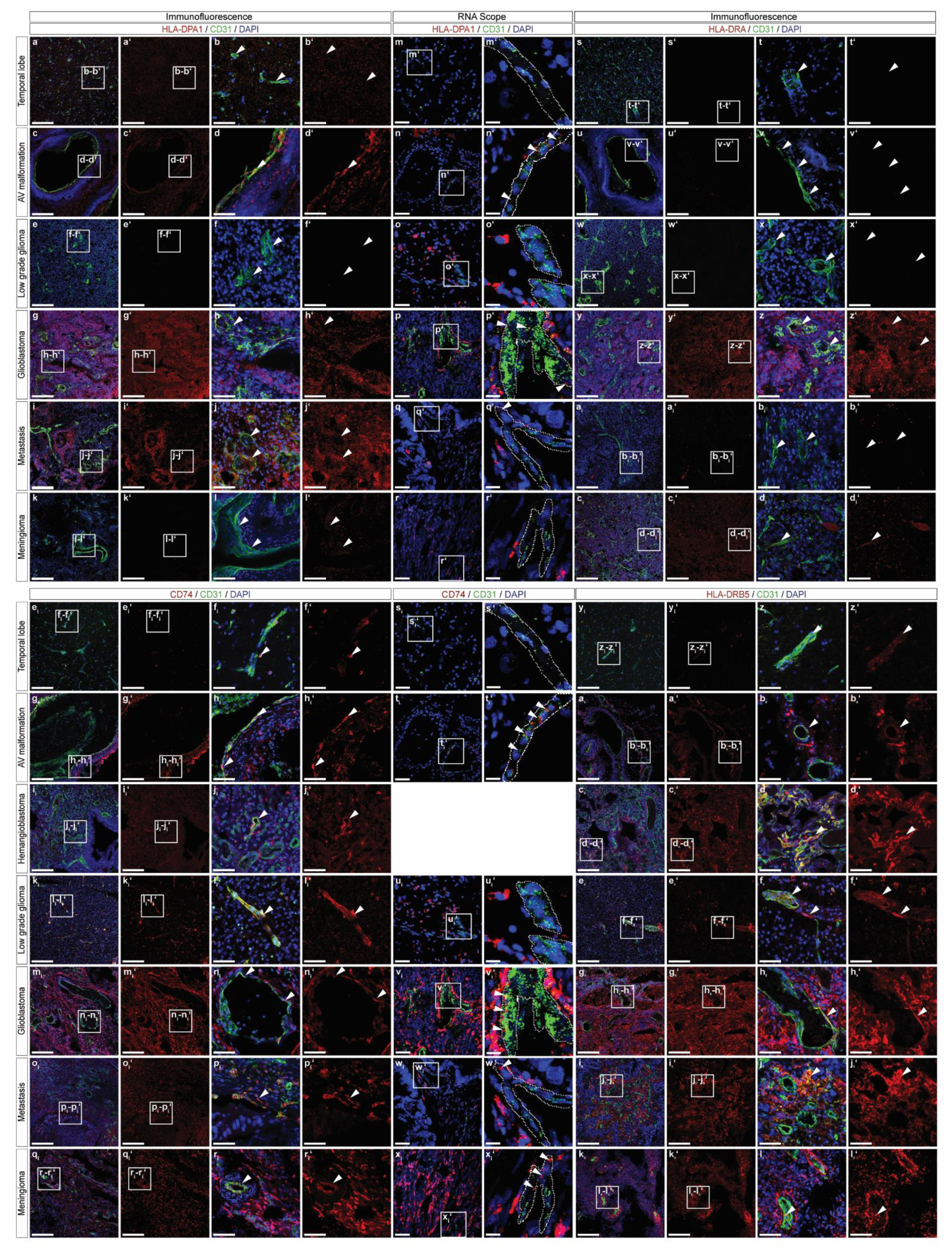
**a-l _ii_‘**, Immunofluorescence (IF) and RNAscope imaging of tissue sections from the indicated entities, stained for HLA-DPA1 (red; **a-l‘**, IF; **m-r‘**, RNAscope), HLA-DRA (red; **s-d_i_‘**, IF), CD74 (red; **e_i_-r_i_‘**, IF; **s_i_-x_i_‘**, RNAscope), HLA-DRB5 (red; **y_i_-l_ii_‘**, IF) and CD31 (green). Nuclei are stained with DAPI (blue). Boxed area is magnified on the right; arrowheads (IF) and dotted lines (RNAscope) indicate vascular structures in the different tissues. Scale bars: 200μm in overviews (IF), 50μm in overviews (RNAscope); 50μm in zooms (IF) and 12.5μm in zooms (RNAscope).

**Extended Data Figure 37.**
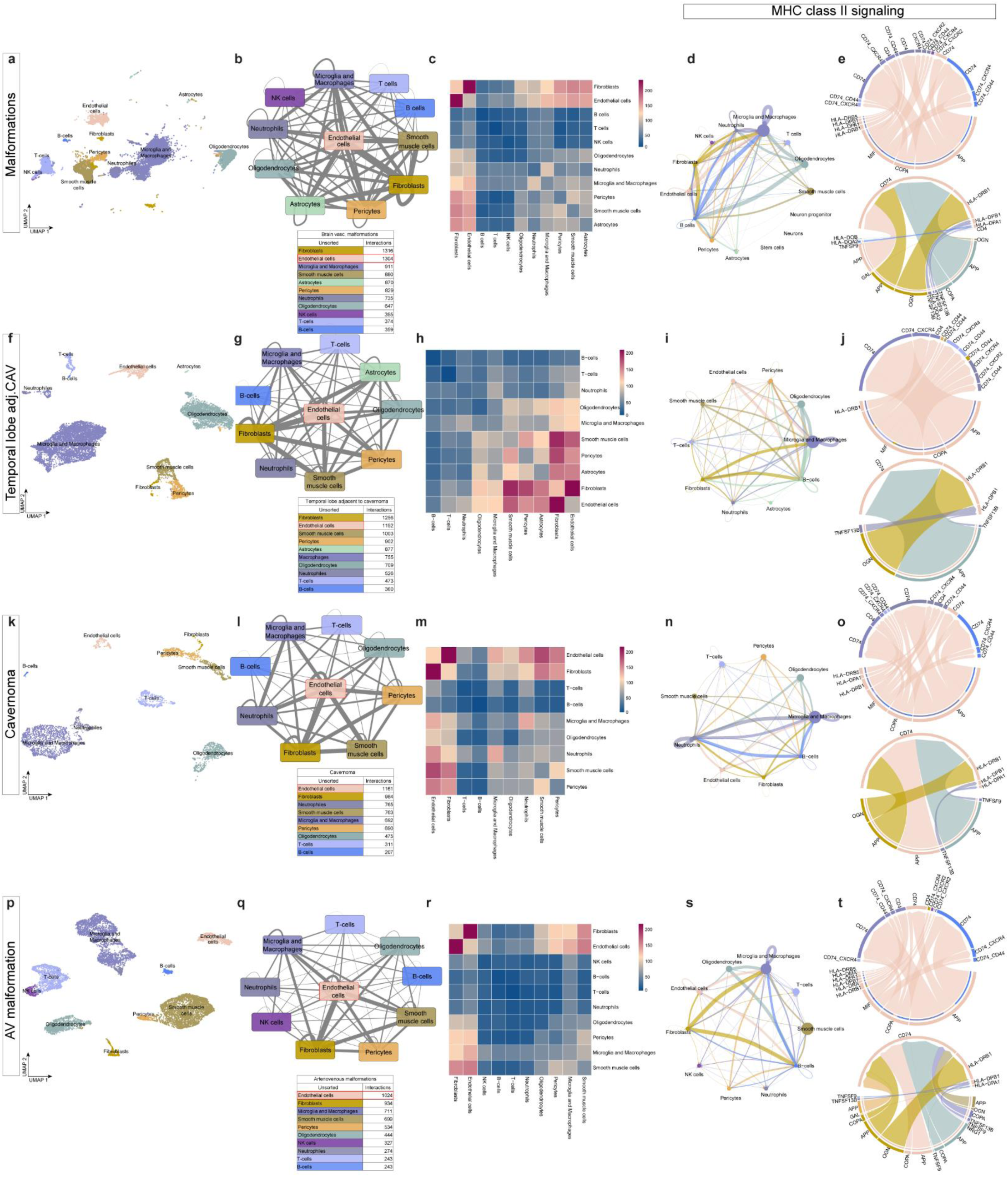
**a,f,k,p,** UMAP plots of endothelial and perivascular cells derived from all brains vascular malformations (**a**), temporal lobes adjacent to cavernomas (**f**), cavernomas (**k**) and arteriovenous malformations (**p**). **b,g,l,q,** Ligand and receptor analysis of all brains vascular malformations (**b**), temporal lobes adjacent to cavernomas (**g**), cavernomas (**l**) and arteriovenous malformations (**q**) cells using CellphoneDB. Line thickness indicates the number of interactions between cell types. Tables summarize the number of interactions for each cell type. **c,h,m,r,** Heatmaps showing the number of ligand-receptor interactions between the different cells of all brains vascular malformations (**c**), temporal lobes adjacent to cavernomas (**h**), cavernomas (**m**) and arteriovenous malformations (**r**). **d,i,n,s,** Circle plots showing the strength of MHC class II signaling interactions between the different cell types of all brain vascular malformations (**d**), temporal lobes adjacent to cavernomas (**i**), cavernomas (**n**) and arteriovenous malformations (**s**) cells. **e,j,o,t,** Visualization of MHC class II connectomic analysis. Cord/circos plots of MHC class II ligand-receptor interactions with EC as “senders” (upper panel) and as “receivers” (lower panel) in all brain vascular malformations (**e**), temporal lobes adjacent to cavernomas (**j**), cavernomas (**o**) and arteriovenous malformations (**t**) cells. Edge thickness represents its weights, whereas edge color indicates the “sender” cell type.

**Extended Data Figure 38.**
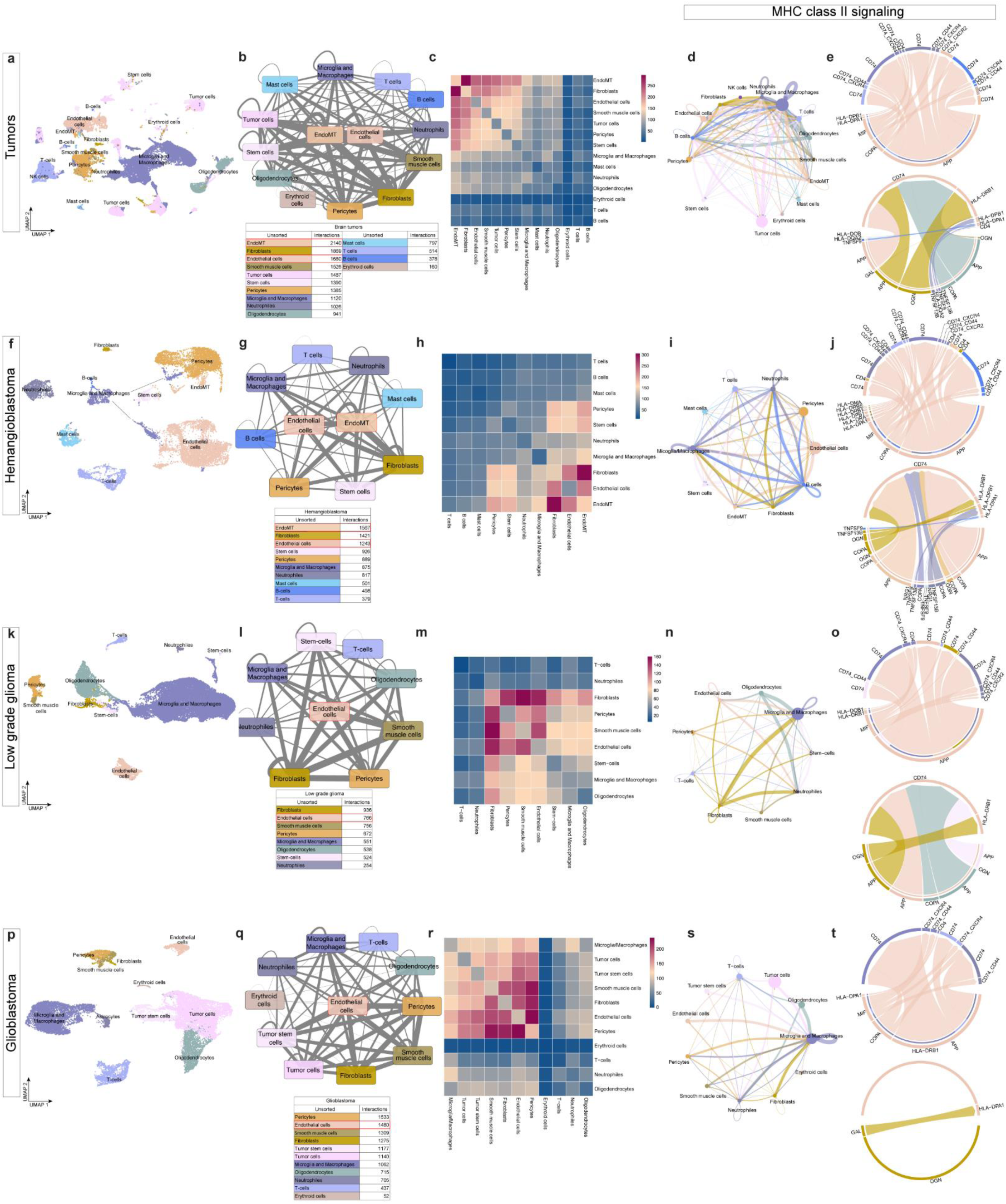
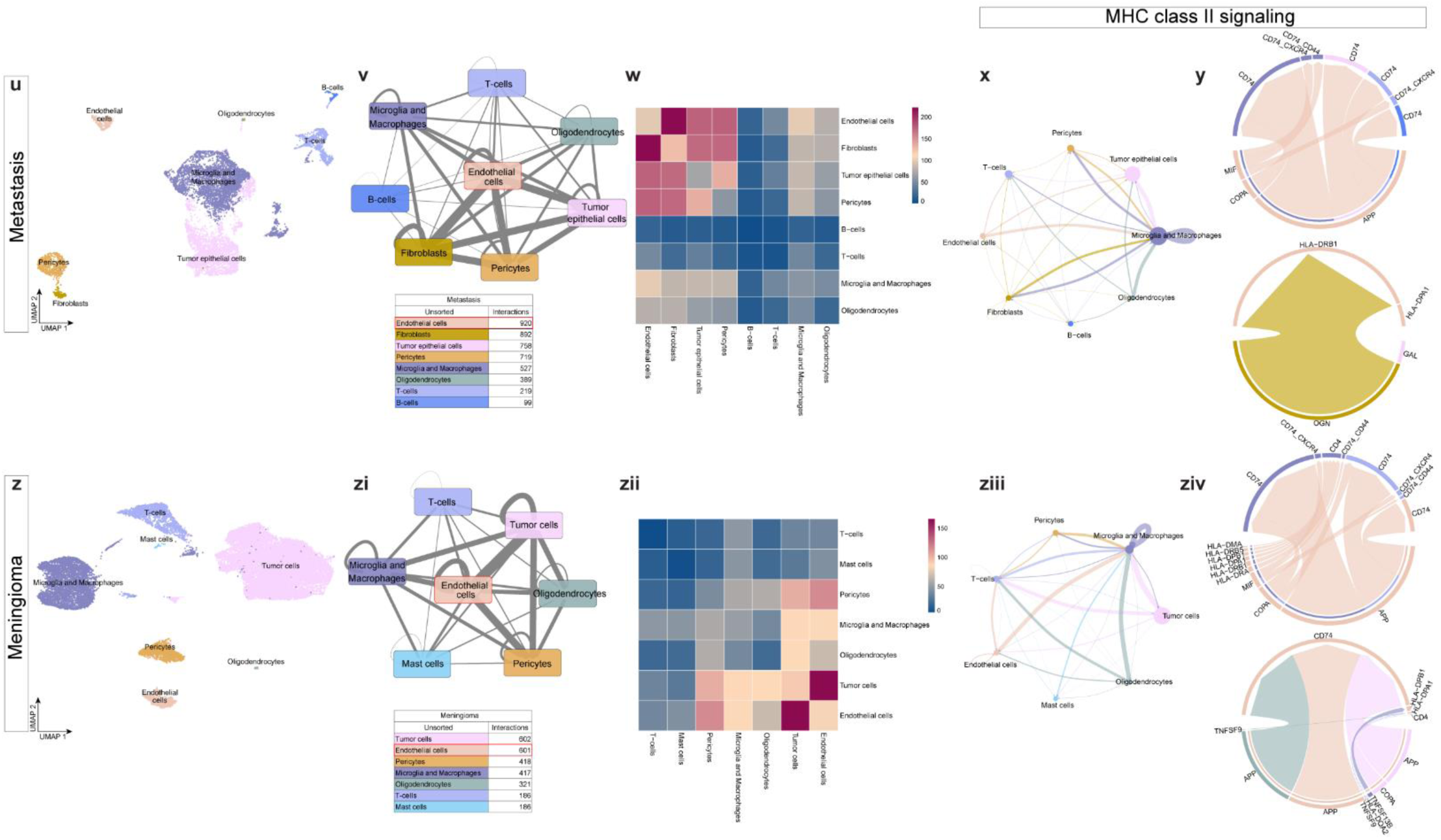
**a,f,k,p,u,z,** UMAP plots of endothelial and perivascular cells derived from all brains tumors (**a**), hemangioblastoma (**f**), low-grade glioma (**k**), glioblastoma (**p**), metastasis (**u**), and meningioma (**z**). **b,g,l,q,v,zi,** Ligand and receptor analysis of all brains tumors (**b**), hemangioblastoma (**g**), low-grade glioma (**l**), glioblastoma (**q**), metastasis (**u**) and meningioma (**zi**) cells using CellphoneDB. Line thickness indicates the number of interactions between cell types. Tables summarize the number of interactions for each cell type. **c,h,m,r,w,zii,** Heatmaps showing the number of ligand-receptor interactions between the different cells of all brains tumors (**c**), hemangioblastoma (**h**), low-grade glioma (**m**), glioblastoma (**r**), metastasis (**w**) and meningioma (**zii**). **d,i,n,s,x,ziii,** Circle plots showing the strength of MHC class II signaling interactions between the different cell types of all brain tumors (**d**), hemangioblastoma (**i**), low-grade glioma (**n**), glioblastoma (**s**), metastasis (**x**) and meningioma (**ziii**) cells. **e,j,o,t,y,ziv,** Visualization of MHC class II connectome analysis. Cord/circos plots of MHC class II ligand-receptor interactions with EC as “senders” (upper panel) and as “receivers” (lower panel) in all brain tumors (**e**), hemangioblastoma (**j**), low-grade glioma (**o**), and glioblastoma (**t**), metastasis (**y**) and meningioma (**ziv**) cells. Edge thickness represents its weights, whereas edge color indicates the “sender” cell type.

**Extended Data Figure 39.**
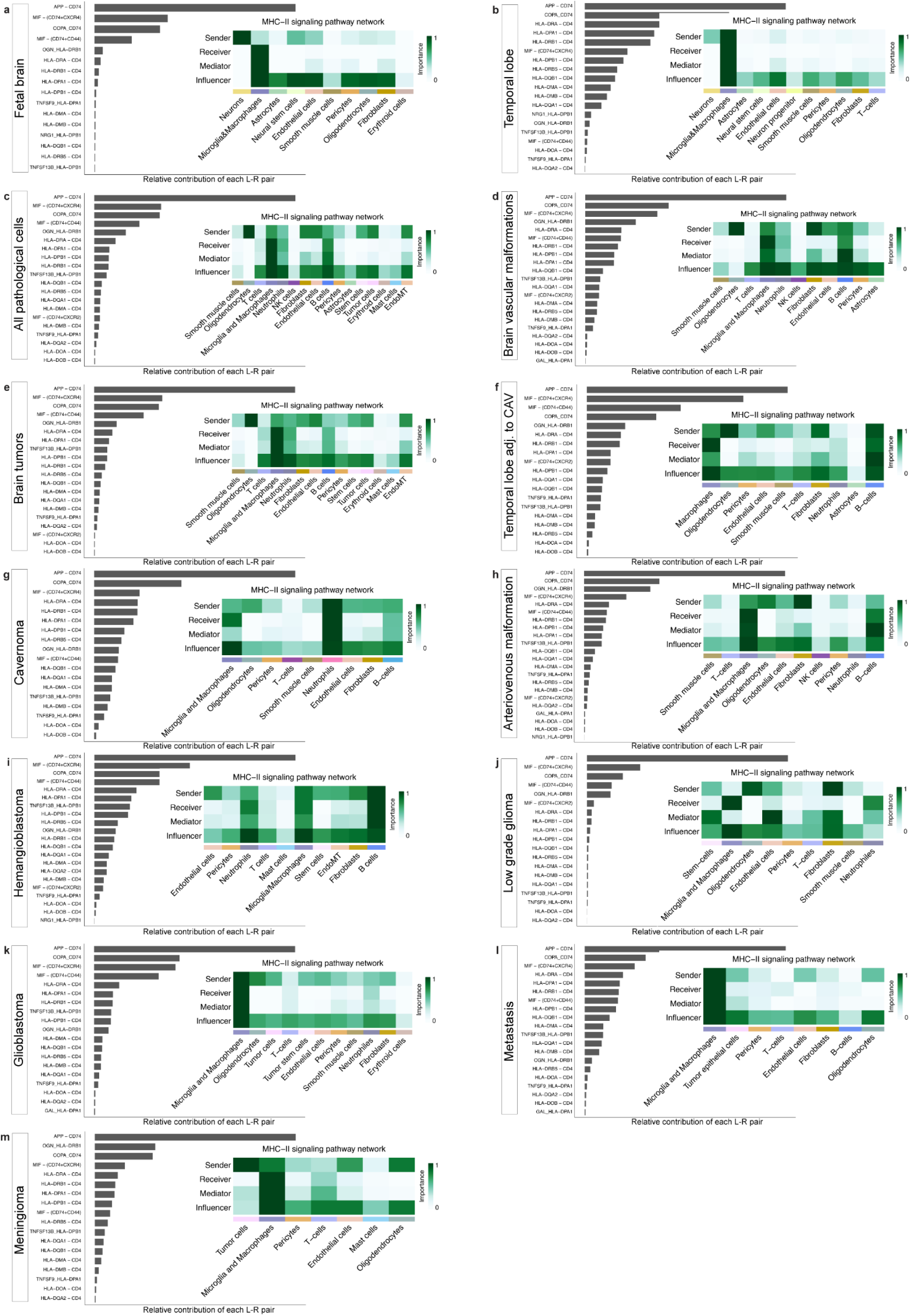
**a-m**, Bar plots showing the relative contribution of each ligand-receptor pair to the overall communication network of MHC class II signaling pathway, and heatmaps showing the relative importance of each cell type based on the computed network centrality in the indicated tissues.

**Extended Data Figure 40.**
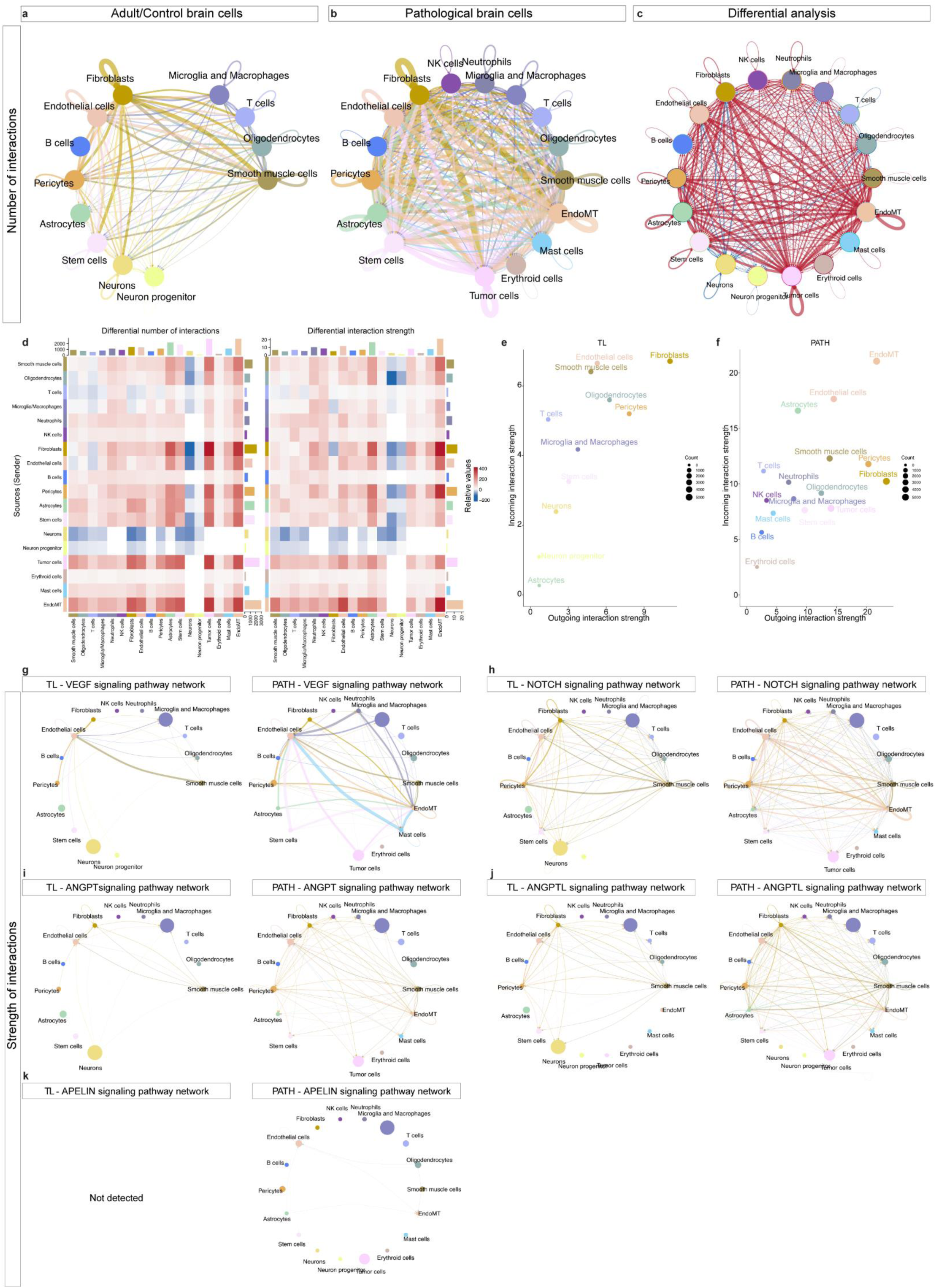
**a-c,** Circle plot showing the number of statistically significant signaling interactions between cells of adult/control brain (**a**), pathological brain (**b**) and the differential analysis of the number of interactions (pathology over control) (**c**), red indicating upregulation, while blue indicating downregulation. **d,** Heatmap showing the differential analysis of the number (left) and strength (right) of ligand-receptor interactions for different cell types (pathology over control). **e,f,** Scatter plot showing the strength of outgoing (x-axis) and incoming (y-axis) signaling pathways of different cell types from temporal lobe and pathological brains. **g-k,** Circle plot showing the strength of *VEGF* (**g**), *NOTCH* (**h**), *ANGPT* (**i**), *ANGPTL* (**j**) and *APELIN* (**k**) signaling interactions between temporal lobe and pathological brain cells (color-coded by cell type).

**Extended Data Figure 41.**
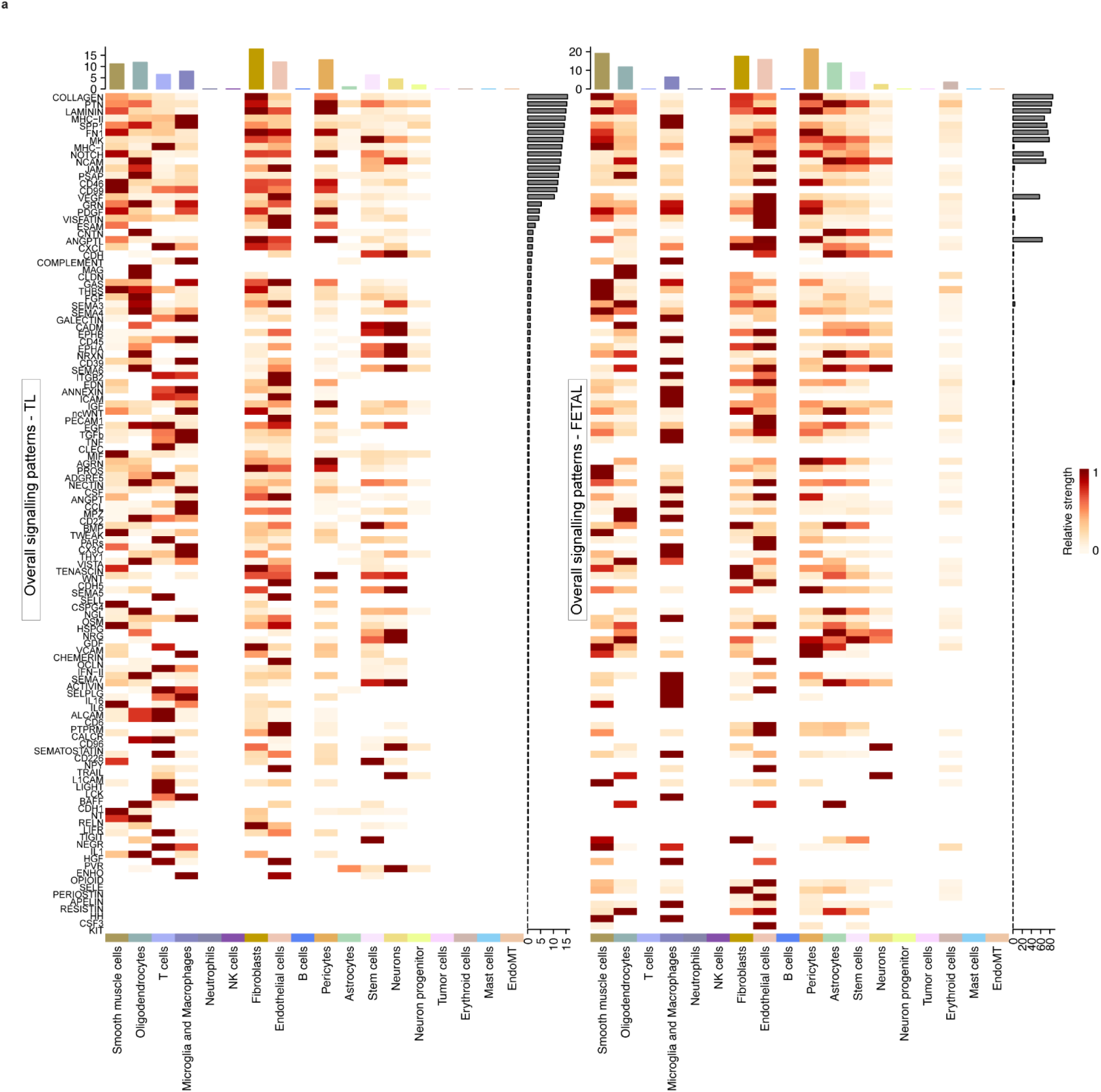
**a,** Heatmap showing overall signaling patters of different cell types in adult/control brains (TL, left panel) and fetal brains (right panel).

**Extended Data Figure 42.**
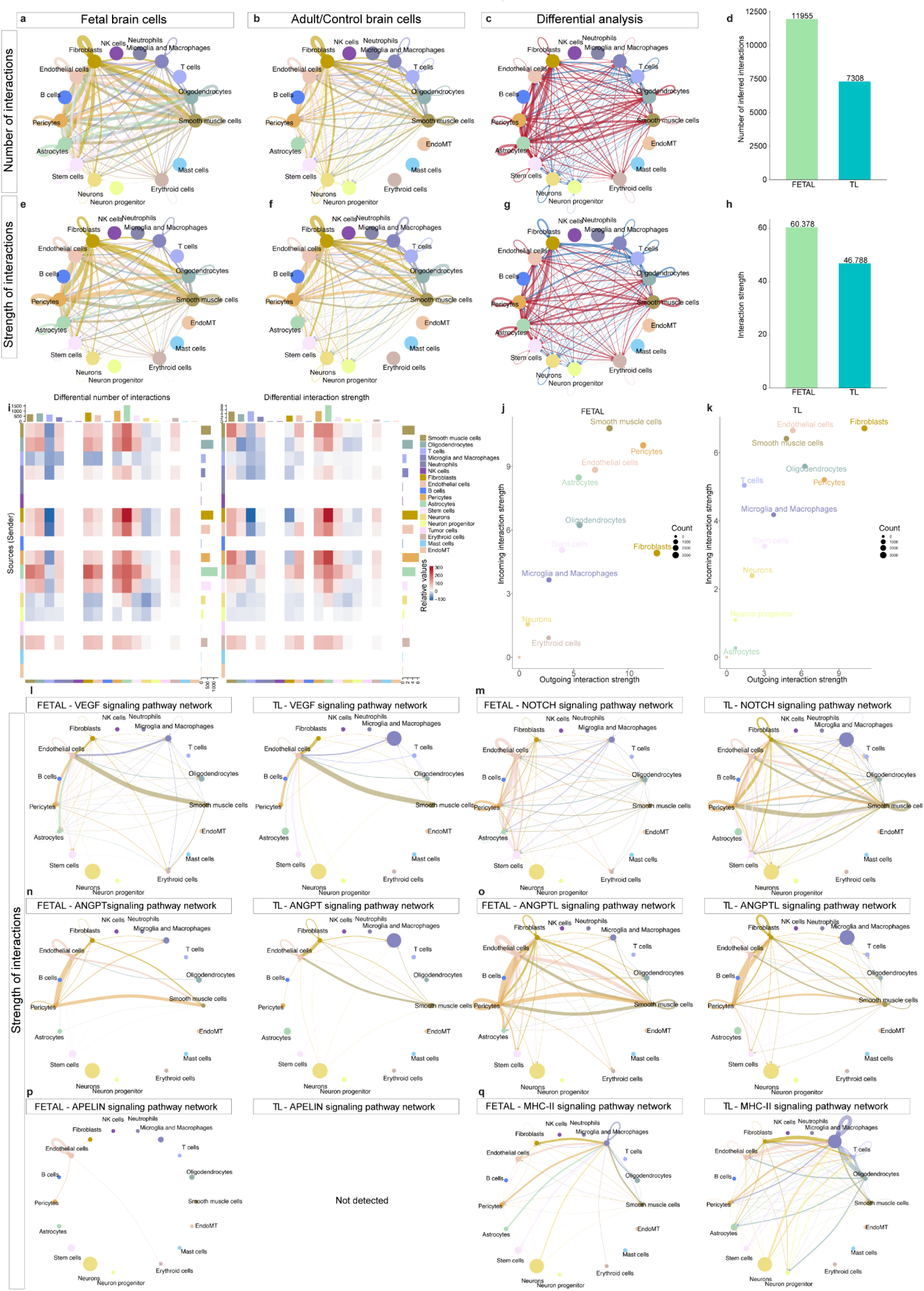
**a-c,** Circle plot showing the number of statistically significant signaling interactions between cells of fetal brains (**a**), adult/control brain (**b**) and the differential analysis of the number of interactions (fetal over adult/control brains), red indicating upregulation, while blue indicating downregulation (**c**). **e-g,** Circle plot showing the strength of signaling interactions between cells of fetal brains (**e**), adult/control brain (**f**) and the differential analysis of the strength of interactions (fetal over adult/control brains), red indicating upregulation, while blue indicating downregulation (**g**). **d,h,** Bar plots showing the number (**d**) and strength of interactions (**h**) in fetal and adult/control brain (TL) cells. **i,** Heatmap showing the differential analysis of the number (left) and strength (right) of ligand-receptor interactions between different cell types (fetal over adult/control brains). **j,k,** Scatter plot showing the strength of of outgoing (x-axis) and incoming (y-axis) signaling pathways of different cell types from fetal and adult/control brains (TL). **l-q,** Circle plot showing the strength of VEGF (**l**), *NOTCH* (**m**), *ANGPT* (**n**), *ANGPTL* (**o**), *APELIN* (**p**) and MHC II (**q**) signaling interactions between fetal and temporal lobe cells (color-coded by cell type).

**Extended Data Figure 43.**
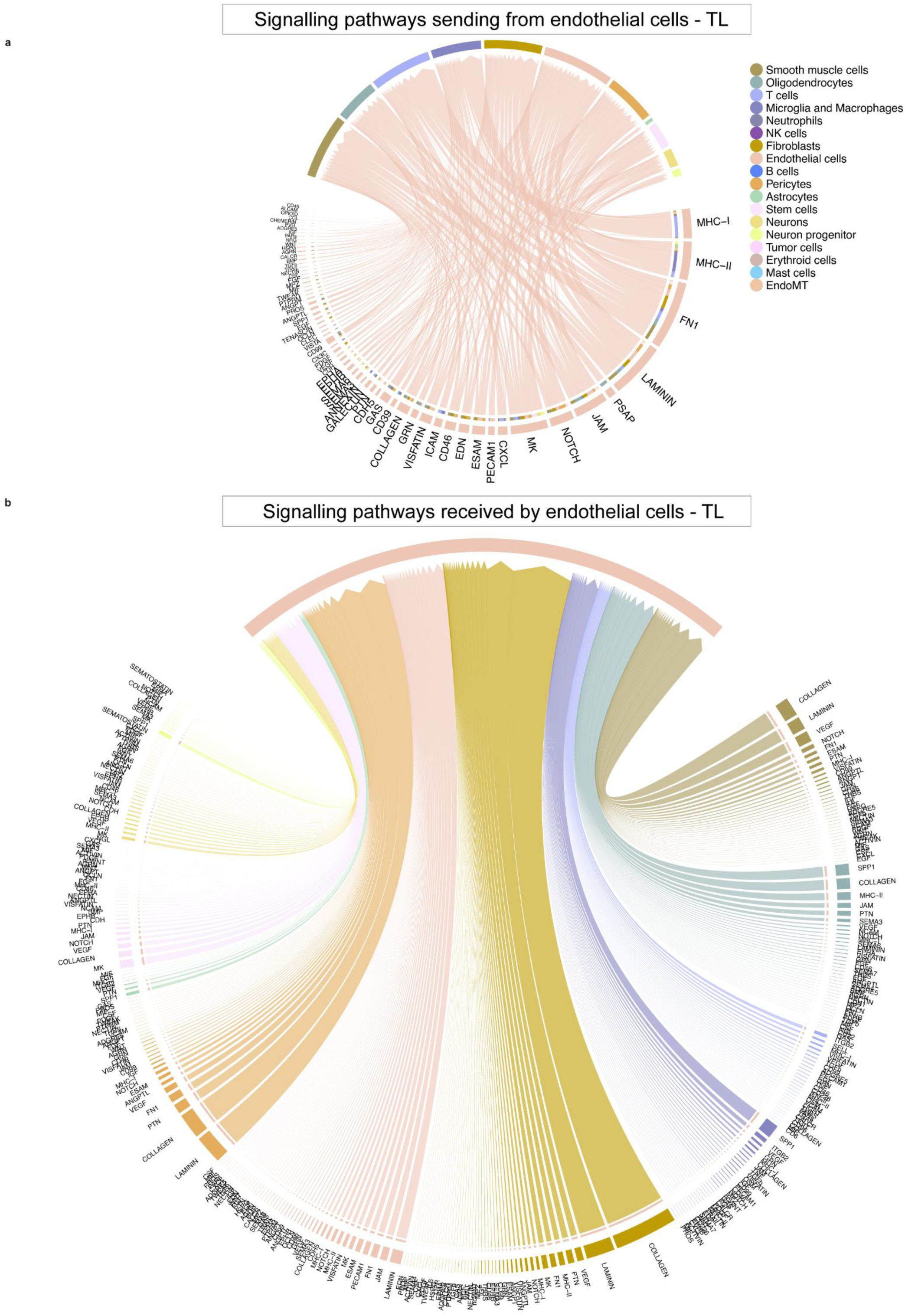
**a,b,** Chord plots showing the ligand-receptor signaling interactions between perivascular cells and ECs in adult/control brains; signaling pathways sending from (**a**) and receiving by endothelial cells (**b**). Edge thickness represents edge weights and edge color indicates the sender cell type.

**Extended Data Figure 44.**
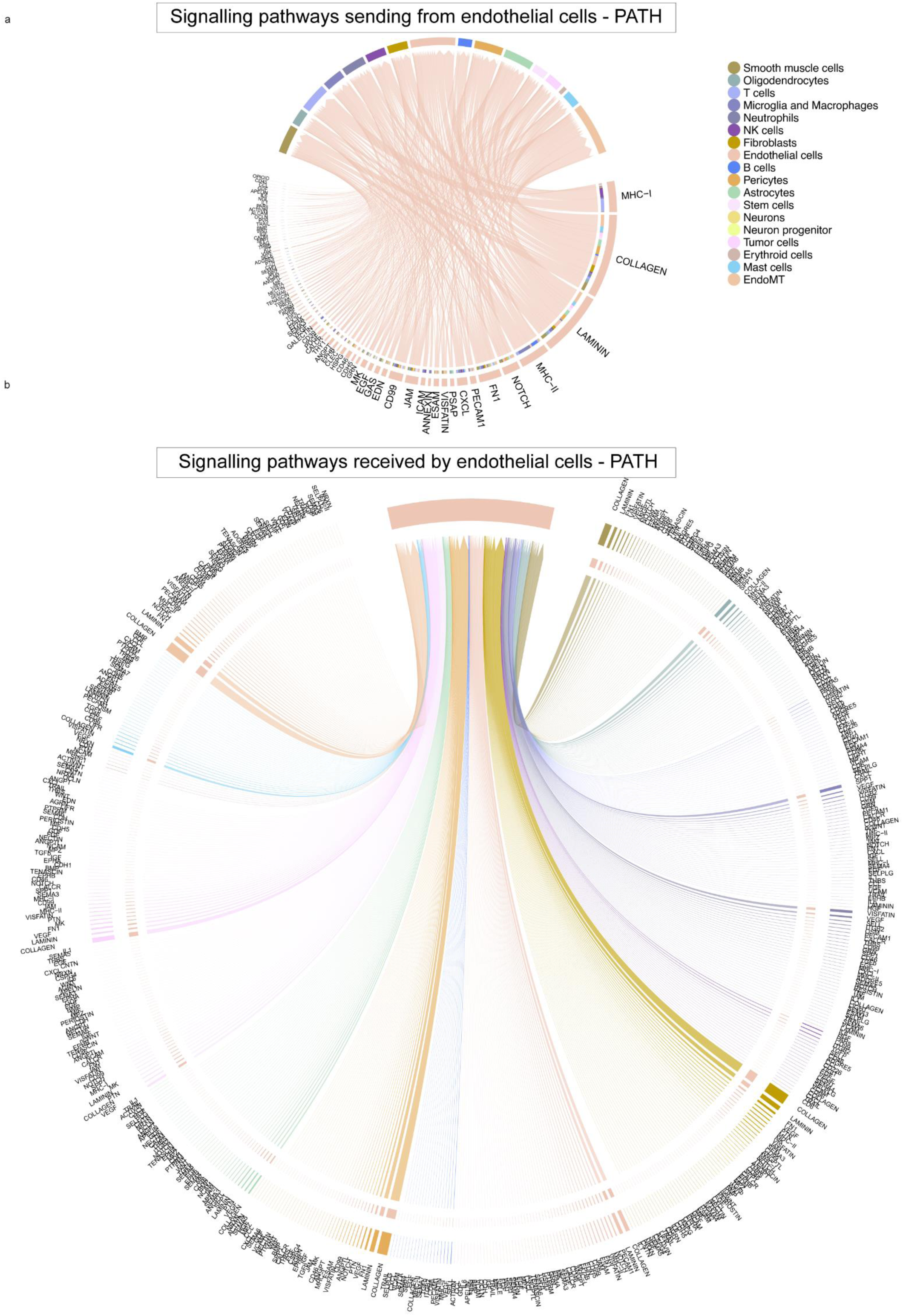
**a,b,** Chord plots showing the ligand-receptor signaling interactions between perivascular cells and ECs in pathological brains; signaling pathways sending from (**a**) and receiving by endothelial cells (**b**). Edge thickness represents edge weights and edge color indicates the sender cell type.

**Extended Data Figure 45.**
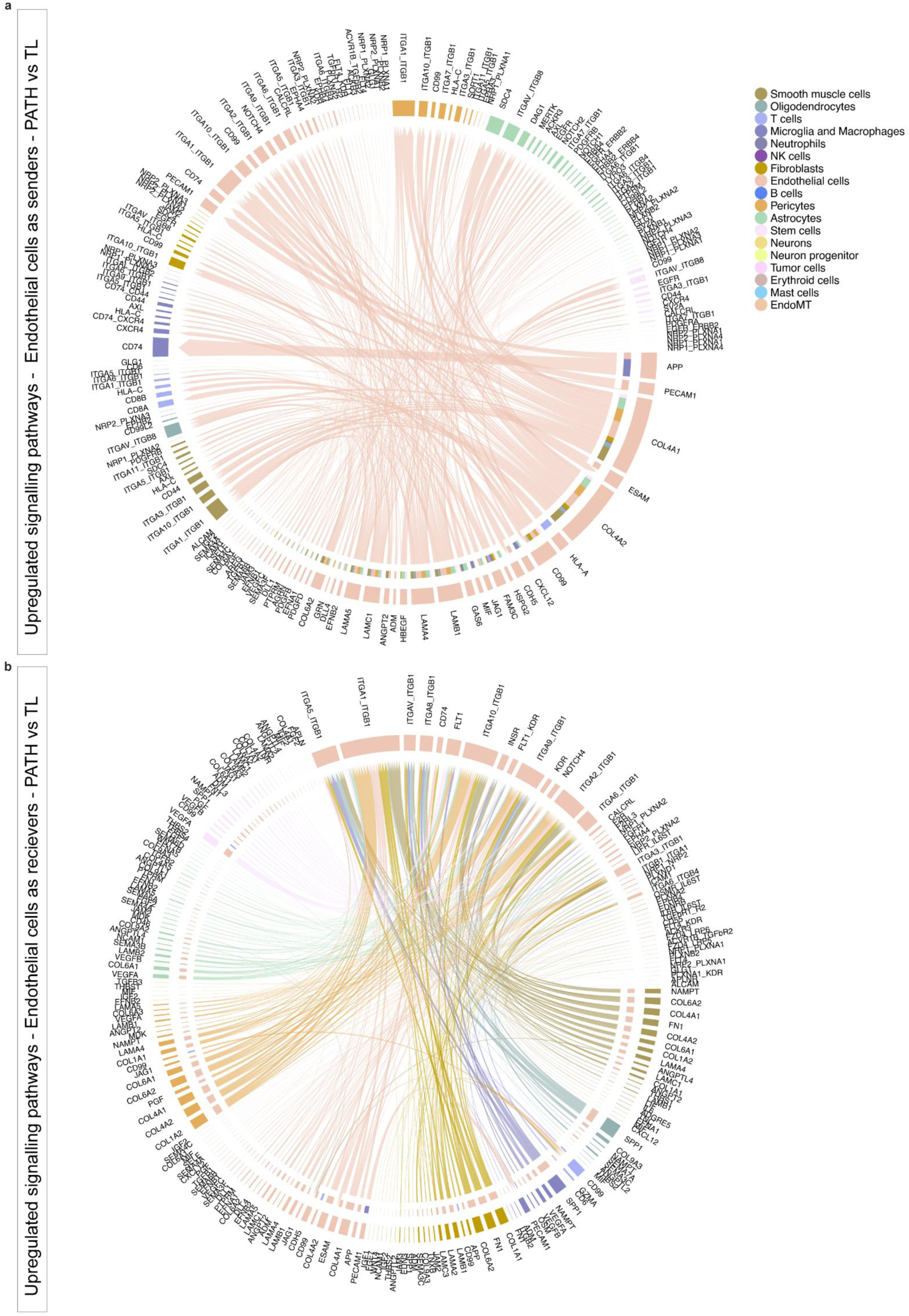
**a,b,** Chord plots showing the ligand-receptor signaling interactions between perivascular cells and ECs upregulated in pathological as compared to adult/control brains; signaling pathways sending from (**a**) and receiving by endothelial cells (**b**). Edge thickness represents edge weights and edge color indicates the sender cell type.

**Extended Data Figure 46.**
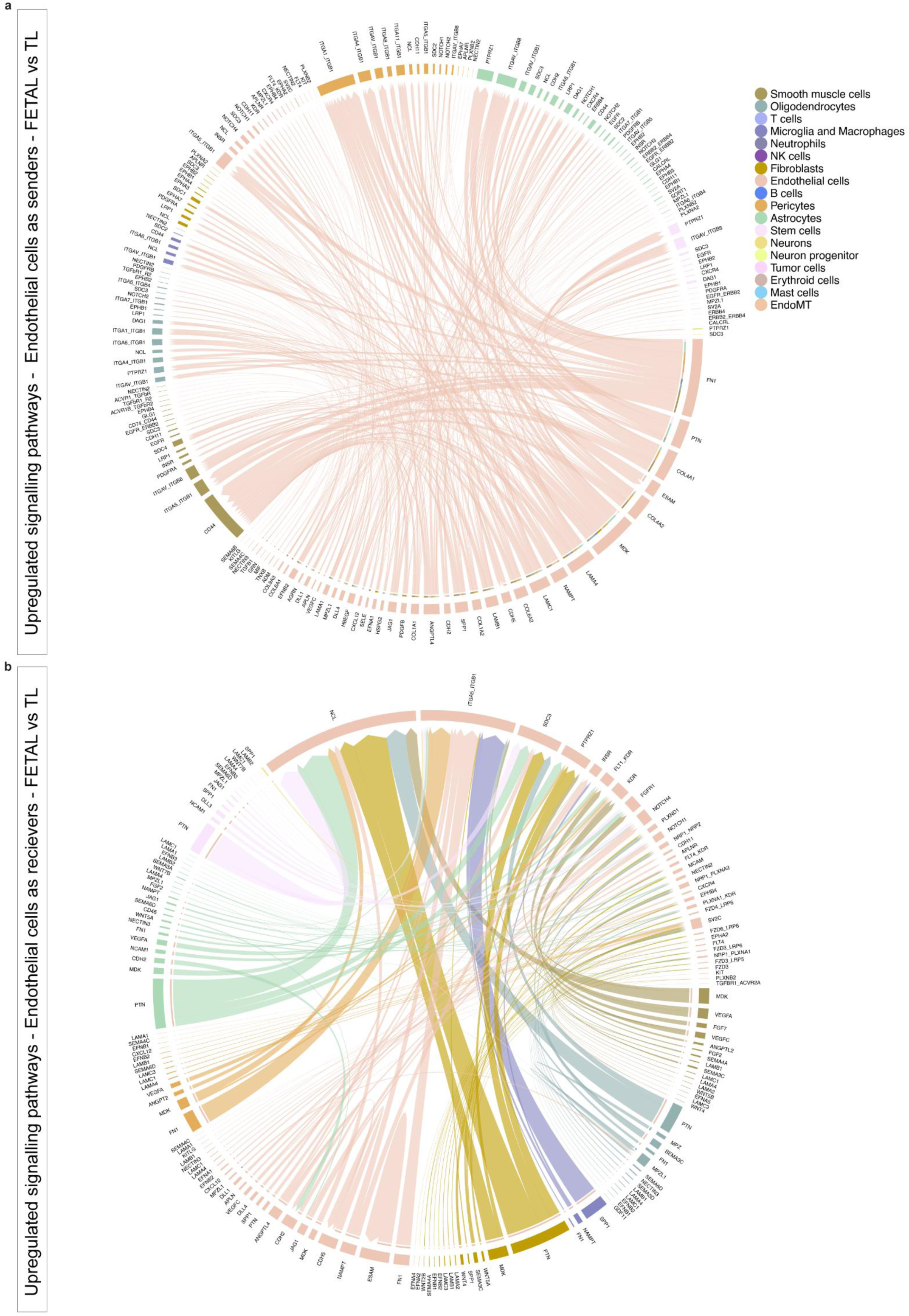
**a,b,** Chord plots showing the ligand-receptor signaling interactions between perivascular cells and ECs upregulated in fetal as compared to adult/control brains; signaling pathways sending from (**a**) and receiving by endothelial cells (**b**). Edge thickness represents edge weights and edge color indicates the sender cell type.

**Extended Data Figure 47.**
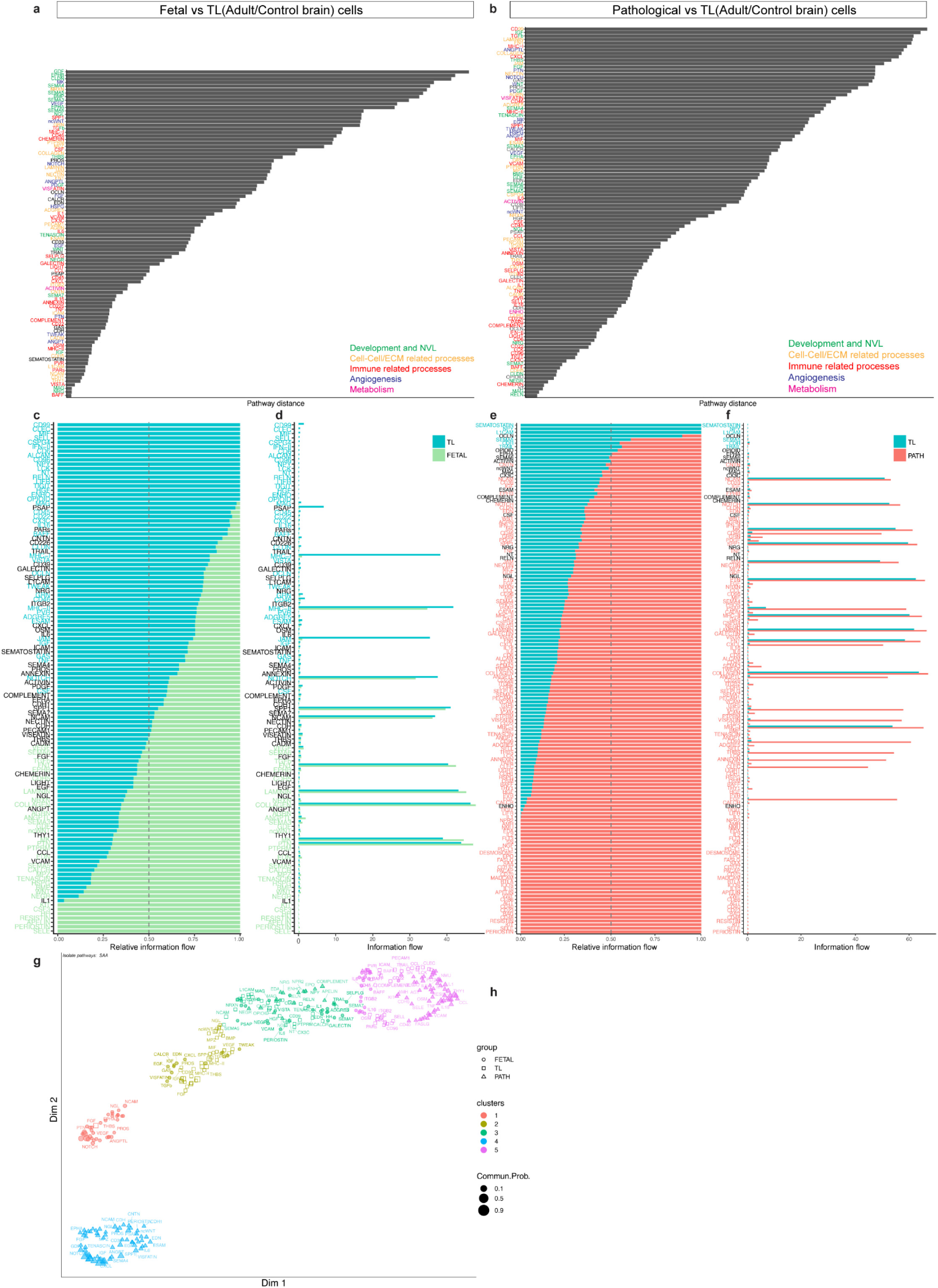
**a,b,** Barplot showing pathway distance; the overlapping signaling pathways between fetal and adult/control brains (**a**), pathological and adult/control brains (**b**) were ranked based on their pairwise Euclidean distance in the shared two-dimensional manifold. **c-f,** Barplot showing the relative (left panel) and absolute (right panel) information flow of all significant signaling pathways within the inferred networks between fetal and adult/control brain cells (**c,d**), and between pathological and adult/control brain cells (**e,f**). **g,** Jointly projecting and clustering signaling pathways from fetal, adult/control and pathological brains onto shared two-dimensional manifold according to their functional similarity of the inferred networks. Circles, squares and triangle symbols represent the signaling networks from fetal, adult/control and pathological brains. Each circle/square/triangle represents the communication network of one signaling pathway. Symbol size is proportional to the total communication probability of that signaling network. Different colors represent different groups of signaling pathways.

**Supplementary Figure 1**

**a,b,c,** Heatmap and hierarchical clustering of all significant genes comparing fetal brain and fetal periphery ECs (**a**), fetal and adult/control brain ECs (**b**), adult/control and pathological brain ECs (**c**).

**Supplementary Figure 2**

**a,** Volcano plot showing the differential expression analysis comparing endothelial cells adult/control brain (left) and brain vascular malformations (right), (Benjamini Hochberg correction; p-value < 0.05 and log2FC≥0.25 colored significant in red). **b,** Heatmap and hierarchical clustering of all significant genes comparing adult/control brain and brain vascular malformations endothelial cells. **c,** Enrichment map visualizing the significantly enriched pathways from the GSEA analysis performed on the ranked list of differentially expressed genes between brain vascular malformation and adult/control brain endothelial cells. Results include enriched genesets belonging to development and NVL, angiogenesis, cell-cell/extracellular matrix interaction, metabolism, and immune related processes.

**d,** Expression heatmap of the top 25 ranking marker genes in the indicated tissues. Color scale: red, high expression; white, intermediate expression; blue, low expression. **e,** Over-representation (enrichment) analysis shown as dotplots representing the top 50 pathways enriched in brain vascular malformations as compared to adult/control brain ECs.

**Supplementary Figure 3**

**a**, Volcano plot showing the differential expression analysis comparing endothelial cells from adult/control brains (left) and brain tumor (right), (Benjamini Hochberg correction; p-value < 0.05 and log2FC≥0.25 colored significant in red). **b**, Heatmap and hierarchical clustering of all significant genes comparing adult/control brain and brain tumor endothelial cells. **c**, Enrichment map visualizing the significantly enriched pathways from the GSEA analysis performed on the ranked list of differentially expressed genes between brain tumor and adult/control brain endothelial cells. Results include enriched genesets belonging to development and NVL, angiogenesis, cell-cell/extracellular matrix interaction, metabolism, and immune related processes. **d**, Expression heatmap of the top 25 ranking marker genes in the indicated tissues. Color scale: red, high expression; white, intermediate expression; blue, low expression. **e,** Over-representation (enrichment) analysis shown as dotplots representing the top 50 pathways enriched in brain tumor as compared to adult/control brain ECs.

**Supplementary Figure 4**

**a,** Volcano plot showing the differential expression analysis comparing endothelial cells from extra-axial tumors (left) and intra-axial tumor (right), (Benjamini Hochberg correction; p-value < 0.05 and log2FC≥0.25 colored significant in red). **b,** Heatmap and hierarchical clustering of all significant genes comparing extra-axial tumors (left) and intra-axial tumor (right) endothelial cells. **c,** Expression heatmap of the top 25 ranking marker genes in the indicated tissues. Color scale: red, high expression; white, intermediate expression; blue, low expression. **d,** Over-representation (enrichment) analysis shown as dotplots representing the top 50 pathways enriched in intra-axial tumors as compared to extra-axial tumors’ ECs.

**Supplementary Figure 5**

**a,** Volcano plot showing the differential expression analysis comparing endothelial cells from secondary brain tumors (left) and primary brain tumors(right), (Benjamini Hochberg correction; p-value < 0.05 and log2FC≥0.25 colored significant in red). **b,** Heatmap and hierarchical clustering of all significant genes comparing primary and secondary brain tumor endothelial cells. **c,** Enrichment map visualizing the significantly enriched pathways from the GSEA analysis performed on the ranked list of differentially expressed genes between primary and secondary brain tumor endothelial cells. Results include enriched genesets belonging to development and NVL, angiogenesis, cell-cell/extracellular matrix interaction, metabolism, and immune related processes. **d,** Expression heatmap of the top 25 ranking marker genes in the indicated tissues. Color scale: red, high expression; white, intermediate expression; blue, low expression. **e,** Over-representation (enrichment) analysis shown as dotplots representing the top 50 pathways enriched in primary as compared to secondary brain tumor ECs.

**Supplementary Figure 6**

**a,** Volcano plot showing the differential expression analysis comparing endothelial cells from low grade glioma (left) and glioblastoma (right), (Benjamini Hochberg correction; p-value < 0.05 and log2FC≥0.25 colored significant in red). **b,** Heatmap and hierarchical clustering of all significant genes comparing low grade glioma and glioblastoma endothelial cells. **c,** Enrichment map visualizing the significantly enriched pathways from the GSEA analysis performed on the ranked list of differentially expressed genes between glioblastoma and low grade glioma endothelial cells. Results include enriched genesets belonging to development and NVL, angiogenesis, cell-cell/extracellular matrix interaction, metabolism, and immune related processes. **d,** Expression heatmap of the top 25 ranking marker genes in the indicated tissues. Color scale: red, high expression; white, intermediate expression; blue, low expression. **e,** Over-representation (enrichment) analysis shown as dotplots representing the top 50 pathways enriched in glioblastoma as compared to low grade glioma ECs.

**Supplementary Figure 7**

**a,** Volcano plot showing the differential expression analysis comparing endothelial cells from adult/control brain (left) and cavernoma (right), (Benjamini Hochberg correction; p-value < 0.05 and log2FC≥0.25 colored significant in red). **b,** Enrichment map visualizing the significantly enriched pathways from the GSEA analysis performed on the ranked list of differentially expressed genes between cavernoma and adult/control brain endothelial cells. Results include enriched genesets belonging to development and NVL, angiogenesis, cell-cell/extracellular matrix interaction, metabolism, and immune related processes.

**Supplementary Figure 8**

**a,** Volcano plot showing the differential expression analysis comparing endothelial cells from adult/control brain (left) and arteriovenous malformation (right), (Benjamini Hochberg correction; p-value < 0.05 and log2FC≥0.25 colored significant in red). **b,** Heatmap and hierarchical clustering of all significant genes comparing adult/control brain and arteriovenous malformation endothelial cells. **c,** Enrichment map visualizing the significantly enriched pathways from the GSEA analysis performed on the ranked list of differentially expressed genes between arteriovenous malformation and adult/control brain endothelial cells. Results include enriched genesets belonging to development and NVL, angiogenesis, cell-cell/extracellular matrix interaction, metabolism, and immune related processes.

**Supplementary Figure 9**

**a,** Volcano plot showing the differential expression analysis comparing endothelial cells from adult/control brain (left) and hemangioblastoma (right), (Benjamini Hochberg correction; p-value < 0.05 and log2FC≥0.25 colored significant in red). **b,** Enrichment map visualizing the significantly enriched pathways from the GSEA analysis performed on the ranked list of differentially expressed genes between hemangioblastoma and adult/control brain endothelial cells. Results include enriched genesets belonging to development and NVL, angiogenesis, cell-cell/extracellular matrix interaction, metabolism, and immune related processes.

**Supplementary Figure 10**

**a,** Volcano plot showing the differential expression analysis comparing endothelial cells from adult/control brain (left) and low-grade glioma (right), (Benjamini Hochberg correction; p-value < 0.05 and log2FC≥0.25 colored significant in red). **b,** Heatmap and hierarchical clustering of all significant genes comparing adult/control brain and low-grade glioma endothelial cells.**c,** Enrichment map visualizing the significantly enriched pathways from the GSEA analysis performed on the ranked list of differentially expressed genes between low-grade glioma and adult/control brain endothelial cells. Results include enriched genesets belonging to development and NVL, angiogenesis, cell-cell/extracellular matrix interaction, metabolism, and immune related processes.

**Supplementary Figure 11**

**a,** Volcano plot showing the differential expression analysis comparing endothelial cells from adult/control brain (left) and glioblastoma (right), (Benjamini Hochberg correction; p-value < 0.05 and log2FC≥0.25 colored significant in red). **b,** Heatmap and hierarchical clustering of all significant genes comparing adult/control brain and glioblastoma endothelial cells.

**Supplementary Figure 12**

**a,** Volcano plot showing the differential expression analysis comparing endothelial cells from adult/control brain (left) and brain metastasis (right), (Benjamini Hochberg correction; p-value < 0.05 and log2FC≥0.25 colored significant in red). **b,** Heatmap and hierarchical clustering of all significant genes comparing adult/control brain and brain metastasis endothelial cells.

**Supplementary Figure 13**

**a,** Volcano plot showing the differential expression analysis comparing endothelial cells from adult/control brain (left) and meningioma (right), (Benjamini Hochberg correction; p-value < 0.05 and log2FC≥0.25 colored significant in red). **b,** Heatmap and hierarchical clustering of all significant genes comparing adult/control brain and meningioma endothelial cells. **c,** Enrichment map visualizing the significantly enriched pathways from the GSEA analysis performed on the ranked list of differentially expressed genes between meningioma and adult/control brain endothelial cells. Results include enriched genesets belonging to development and NVL, angiogenesis, cell-cell/extracellular matrix interaction, metabolism, and immune related processes.

**Supplementary Figure 14**

**a,b,** Heatmap showing outgoing signalling patterns of different EC subtypes in adult/control (**a**) and pathological (**b**) brains. **c,d,** Heatmap showing incoming signalling patterns of different EC subtypes in adult/control (**c**) and pathological (**d**) brains.

**Supplementary Figure 15**

**a,b,** Comparison of the significant ligand-receptor pairs between pathological and adult/control brains, which contribute to the signalling sending from angiogenic capillaries to all other EC subtypes (**a**) and to signalling received (**b**) by angiogenic capillaries from other EC subtypes. Left panel are pathways upregulated in pathological brain as compared to adult/control brain endothelial cells, while right panel are signalling pathways downregulated in pathological brain as compared to adult/control brain endothelial cells. Dot colour reflects communication probabilities and dot size represents computed p-values. Empty space means the communication probability is zero. p-values are computed from one-sided permutation test.

**Supplementary Figure 16**

**a,b,** Heatmap showing outgoing signalling patterns of different EC subtypes in adult/control (**a**) and fetal (**b**) brains. **c,d,** Heatmap showing incoming signalling patterns of different EC subtypes in adult/control (**c**) and fetal (**d**) brains.

**Supplementary Figure 17**

**a,b,** Dotplot heatmap showing the comparison of the significant ligand-receptor pairs between fetal and adult/control brains, which contribute to the signalling sending from angiogenic capillaries to all other EC subtypes (**a**) and to signalling received (**b**) by angiogenic capillaries from other EC subtypes. Left panel are pathways upregulated in fetal as compared to adult/control brain endothelial cells, while right panel are signalling pathways downregulated in pathological brain as compared to adult/control brain endothelial cells. Dot colour reflects communication probabilities and dot size represents computed p-values. Empty space means the communication probability is zero. p-values are computed from one-sided permutation test.

**Supplementary Figure 18**

**a-n,** Scatter plot showing the differential incoming and outgoing interaction strength of pathways in the different EC subtypes - identifying signalling changes in those cells in pathological as compared to adult/control brains.

**Supplementary Figure 19**

**a-n,** Scatter plot showing the differential incoming and outgoing interaction strength of pathways in the different EC subtypes - identifying signalling changes in those cells in fetal as compared to adult/control brains.

**Supplementary Figure 20**

**a-i,** Dotplot heatmap of MHC class I and II ligand–receptor interactions in the indicated entities analysed used cellphoneDB package; p-values are indicated by circle size, scale on right. The means of the average expression level of interacting ligands and receptors in the endothelial cells and the respective NVU cells are indicated by colour scale on the right (red, high; yellow, medium-high; blue, medium-low; black, low).

**Supplementary Figure 21**

**a-r,** Scatter plot showing the differential incoming and outgoing interaction strength of pathways in the cell types in adult/control and pathological brains - identifying signalling changes in those cells in pathological as compared to control conditions.

**Supplementary Figure 22**

**a-k,** Scatter plot showing the differential incoming and outgoing interaction strength of pathways in the cell types in fetal and adult/control brains - identifying signalling changes in those cells in developmental as compared to adult/control states.

**Supplementary Figure 23**

**a,b,** Heatmap showing outgoing signalling patterns of different cell types in adult/control (**a**) and pathological (**b**) brains. c,d, Heatmap showing incoming signalling patterns of different cell types in adult/control (**c**) and pathological (**d**) brains.

**Supplementary Figure 24**

**a,b,** Heatmap showing outgoing signalling patterns of different cell types in adult/control (**a**) and fetal (**b**) brains. **c,d,** Heatmap showing incoming signalling patterns of different cell types in adult/control (**c**) and fetal (**d**) brains.

**Supplementary Figure 25**

**a,b,** Dotplot heatmap showing the significantly upregulated ligand-receptor interactions in pathological as compared to adult/control brains, which contribute to the signalling sending from endothelial cells to all other cell types (**a**) and to signalling received (**b**) by endothelial cells from other cell types. Dot colour reflects communication probabilities and dot size represents computed p-values. Empty space means the communication probability is zero. p-values are computed from one-sided permutation test.

**Supplementary Figure 26**

**a,b,** Dotplot heatmap showing the comparison of a selection of significant ligand-receptor pairs between adult/control and pathological brains, which contribute to the signalling sending from endothelial cells to all other cell types (**a**) and to signalling received (**b**) by endothelial cells from other cell types. Dot colour reflects communication probabilities and dot size represents computed p-values. Empty space means the communication probability is zero. p-values are computed from one-sided permutation test.

**Supplementary Figure 27**

**a,b,** Dotplot heatmap showing the comparison of significant ligand-receptor pairs between fetal and adult/control brains, which contribute to the signalling sending from endothelial cells to all other cell types. Left panel are pathways upregulated in fetal brain as compared to adult/control brain cells (**a**), while right panel are signalling pathways downregulated in fetal brain as compared to adult/control brain cells (**b**). Dot colour reflects communication probabilities and dot size represents computed p-values. Empty space means the communication probability is zero. p-values are computed from one-sided permutation test.

**Supplementary Figure 28**

**a,b,** Dotplot heatmap showing the comparison of significant ligand-receptor pairs between fetal and adult/control brains, which contribute to the signalling received by endothelial cells from all other cell types. Left panel are pathways upregulated in fetal brain as compared to adult/control brain cells (**a**), while right panel are signalling pathways downregulated in fetal brain as compared to adult/control brain cells (**b**). Dot colour reflects communication probabilities and dot size represents computed p-values. Empty space means the communication probability is zero. p-values are computed from one-sided permutation test.

