## Supplementary figures for "Molecular atlas of the human brain vasculature at the single-cell level"

Supplementary Figure 1

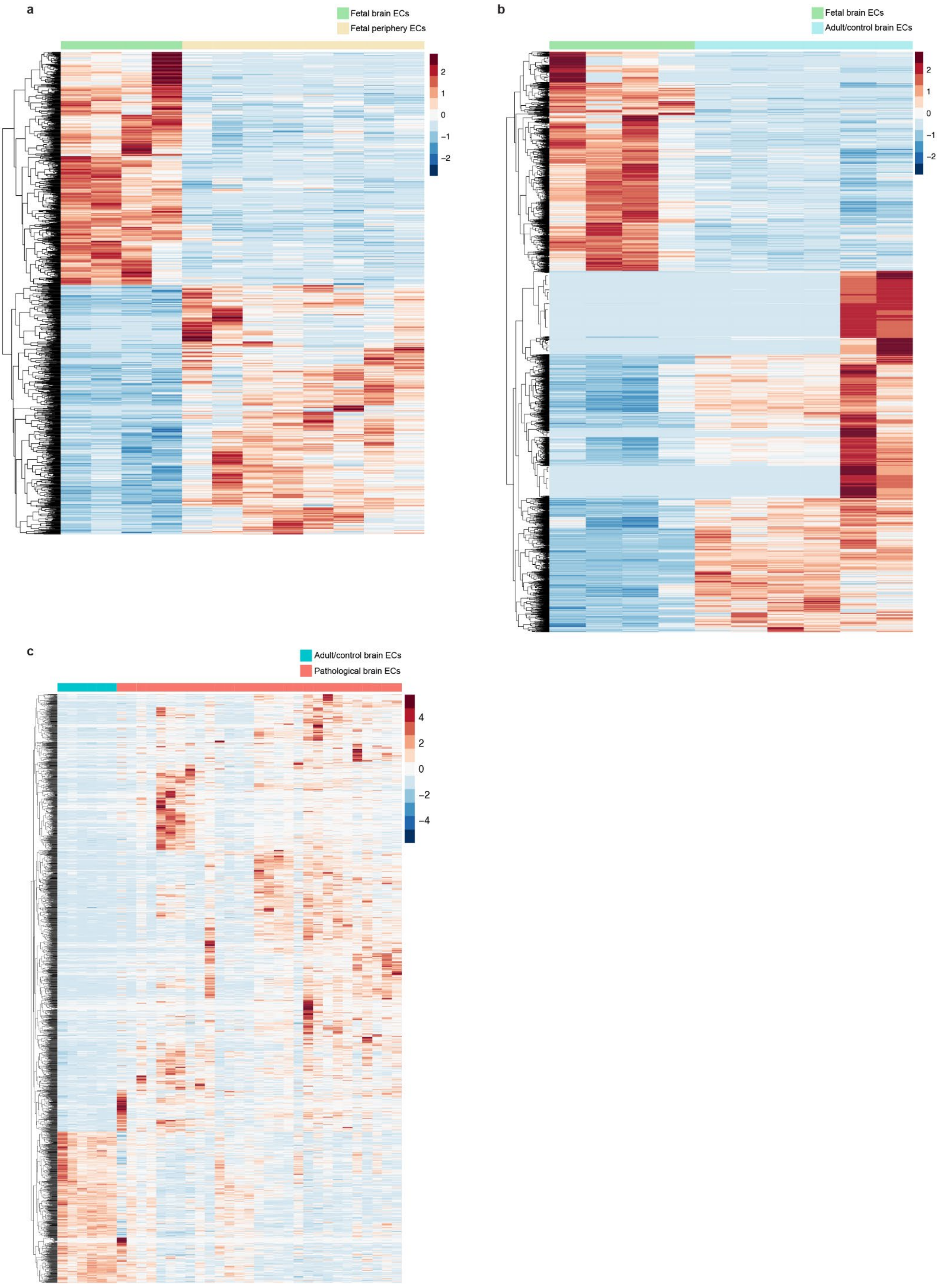

#### Supplementary Figure 2

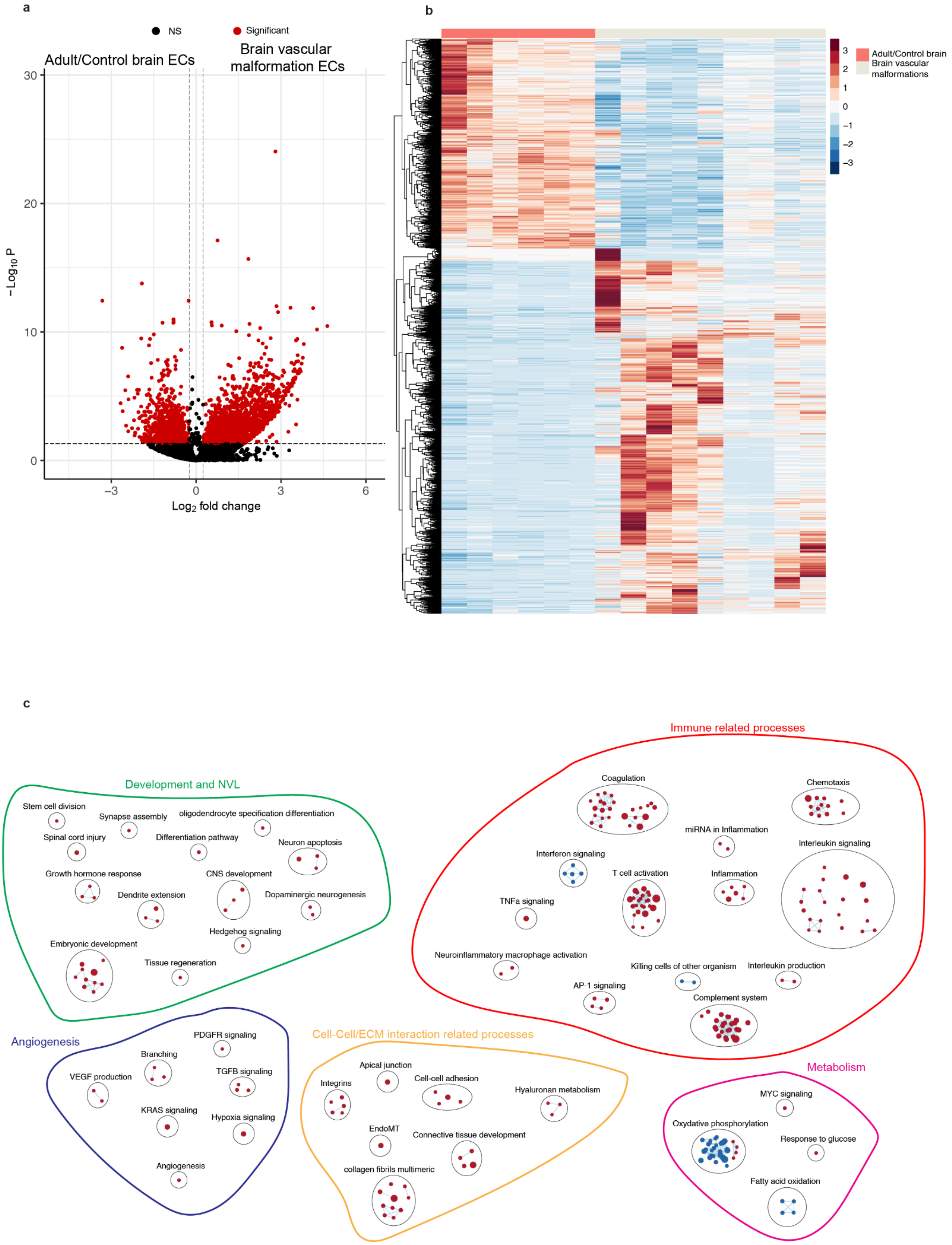

### Supplementary Figure 2

d e

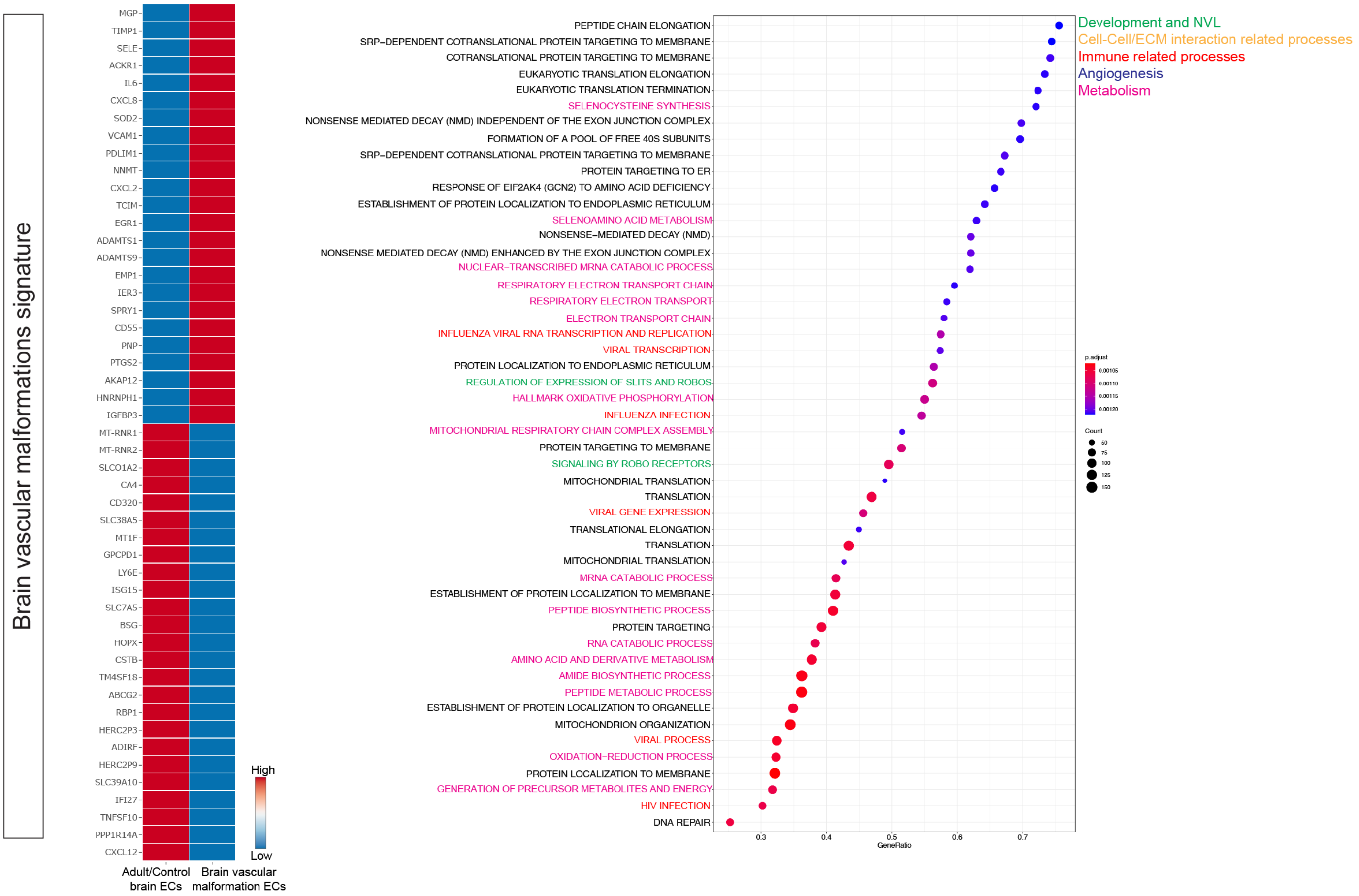

#### Supplementary Figure 3

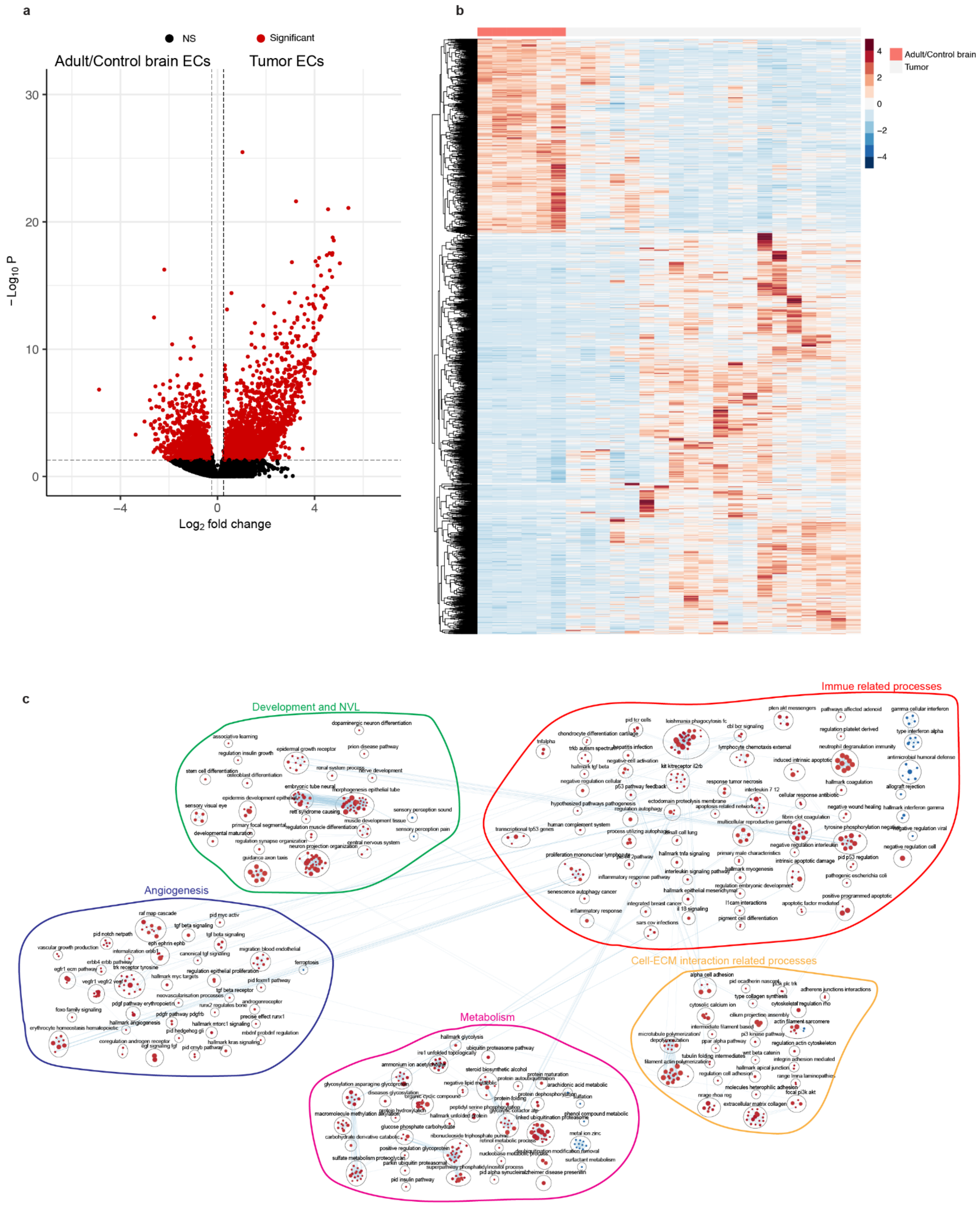

Supplementary Figure 3

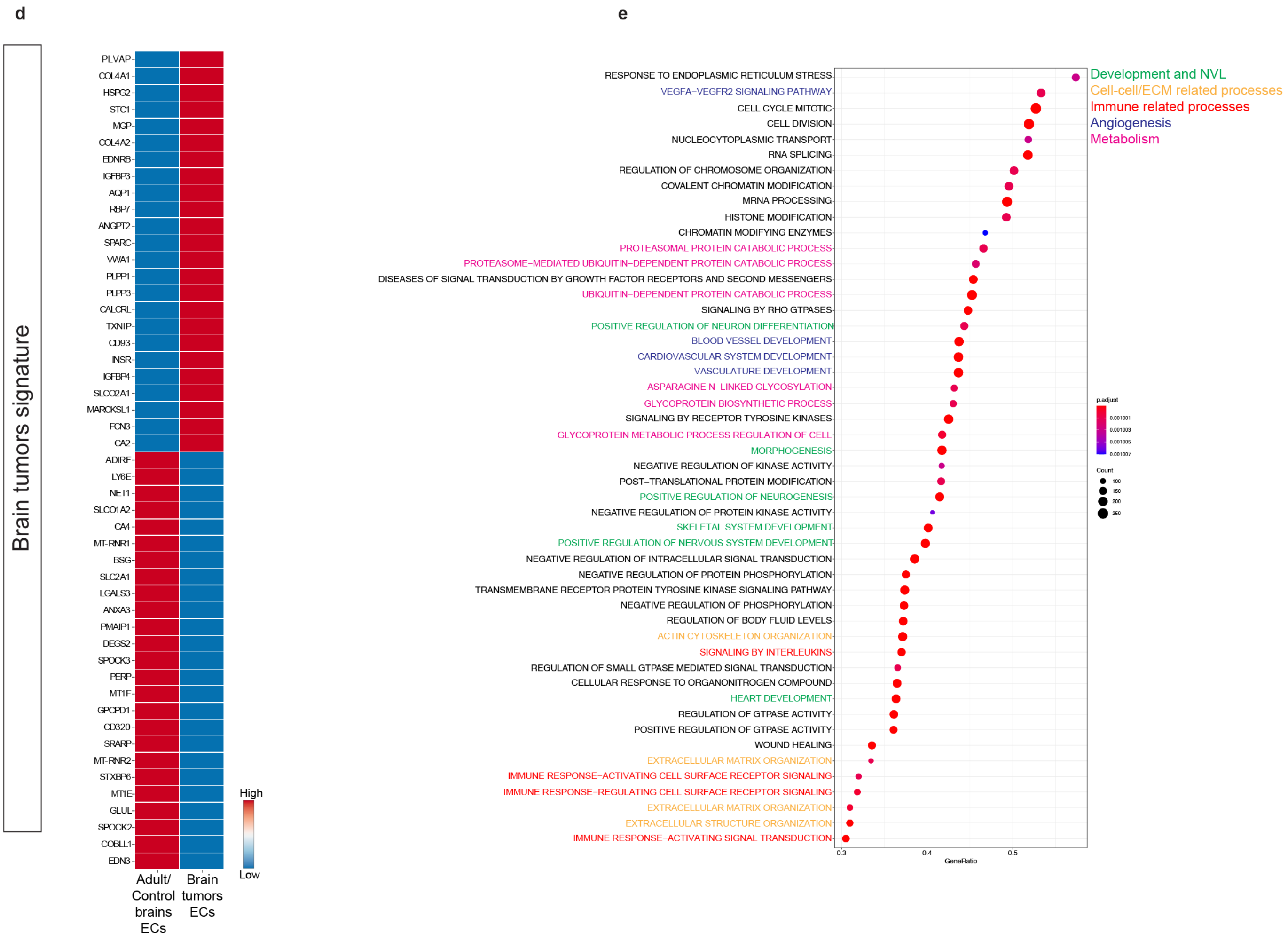

Supplementary Figure 4

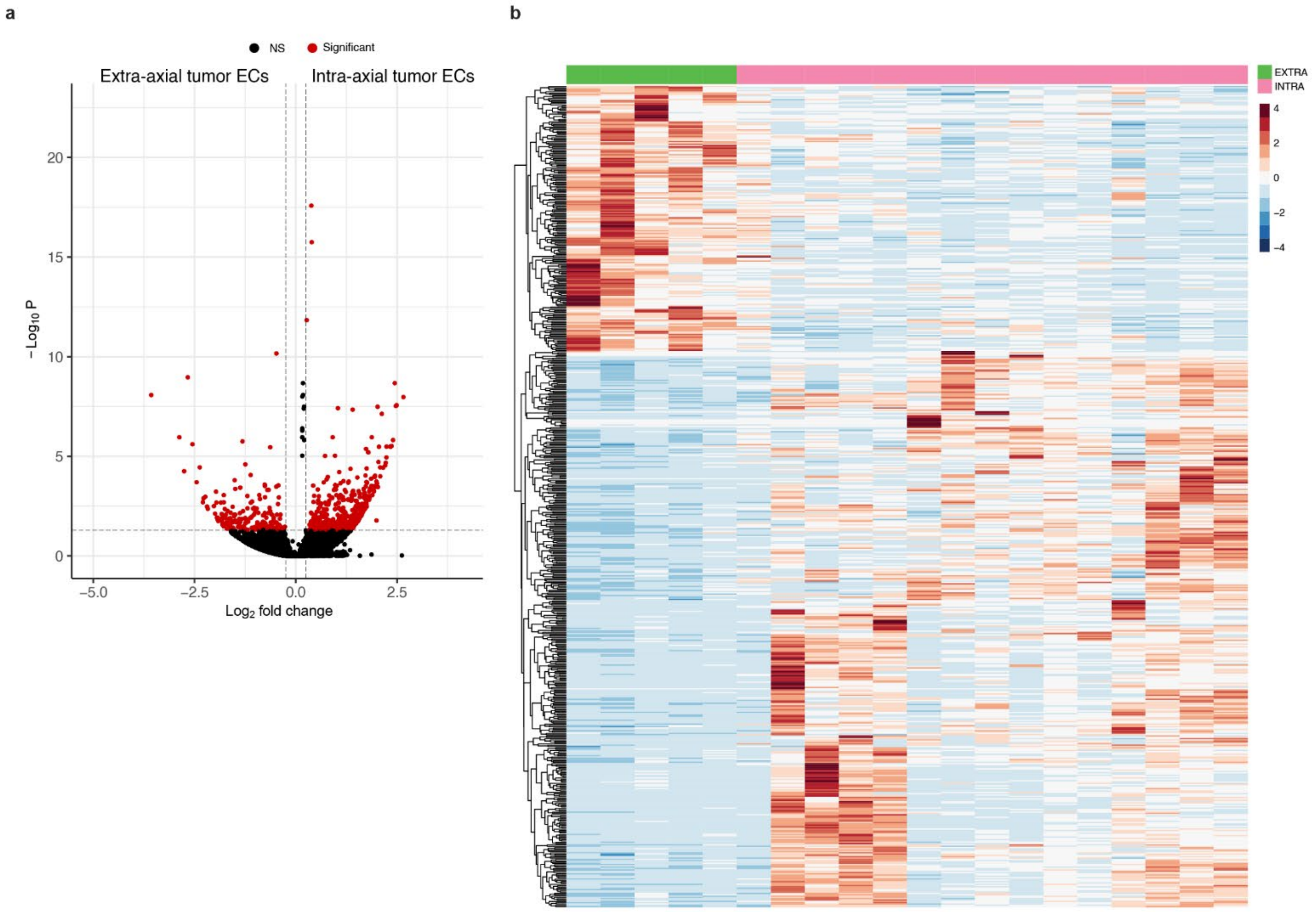

### Supplementary Figure 4

c

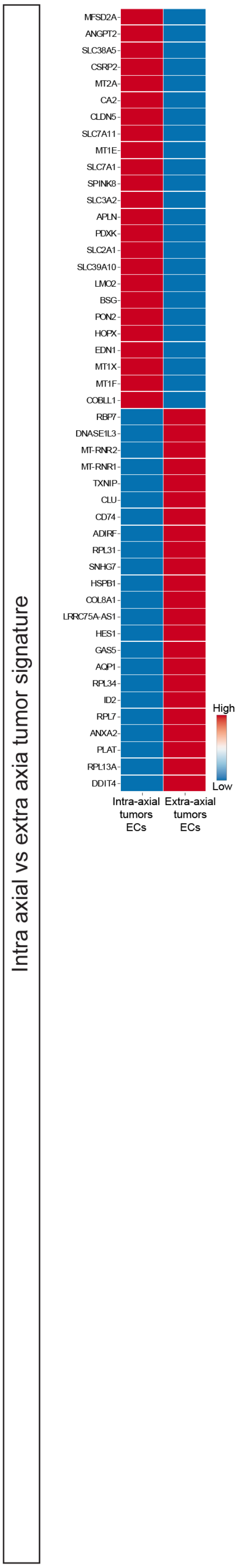


d

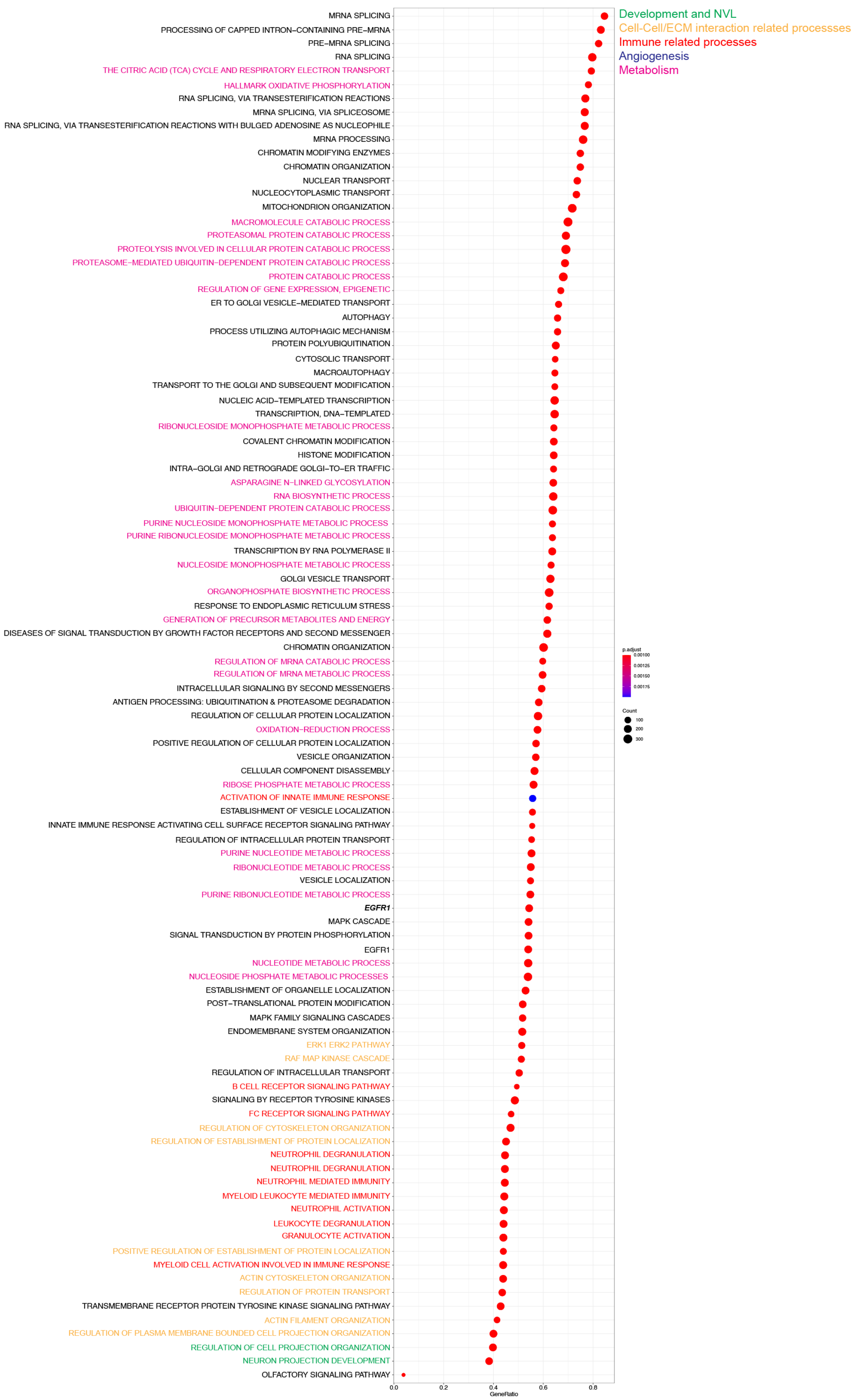

### Supplementary Figure 5

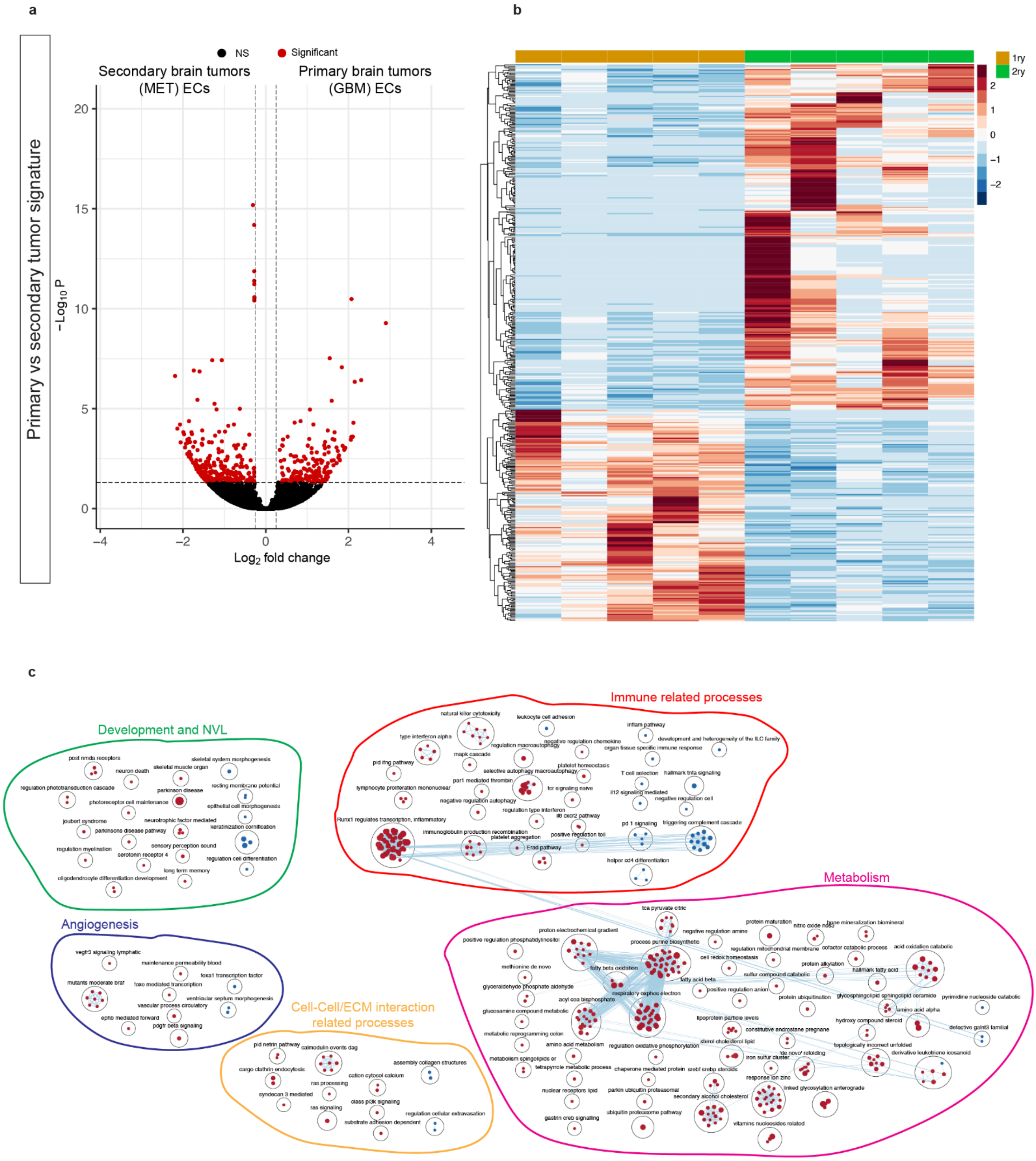

### Supplementary Figure 5

**d**

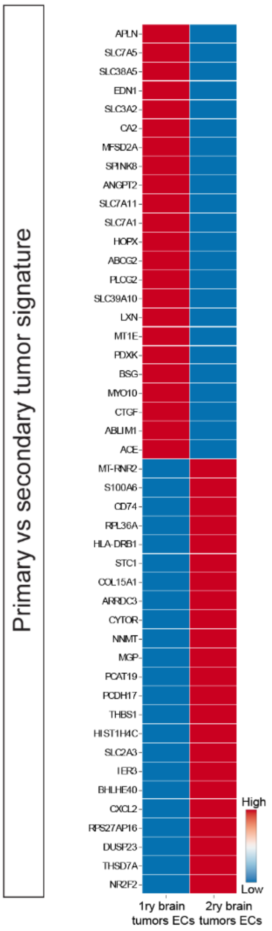


**e**

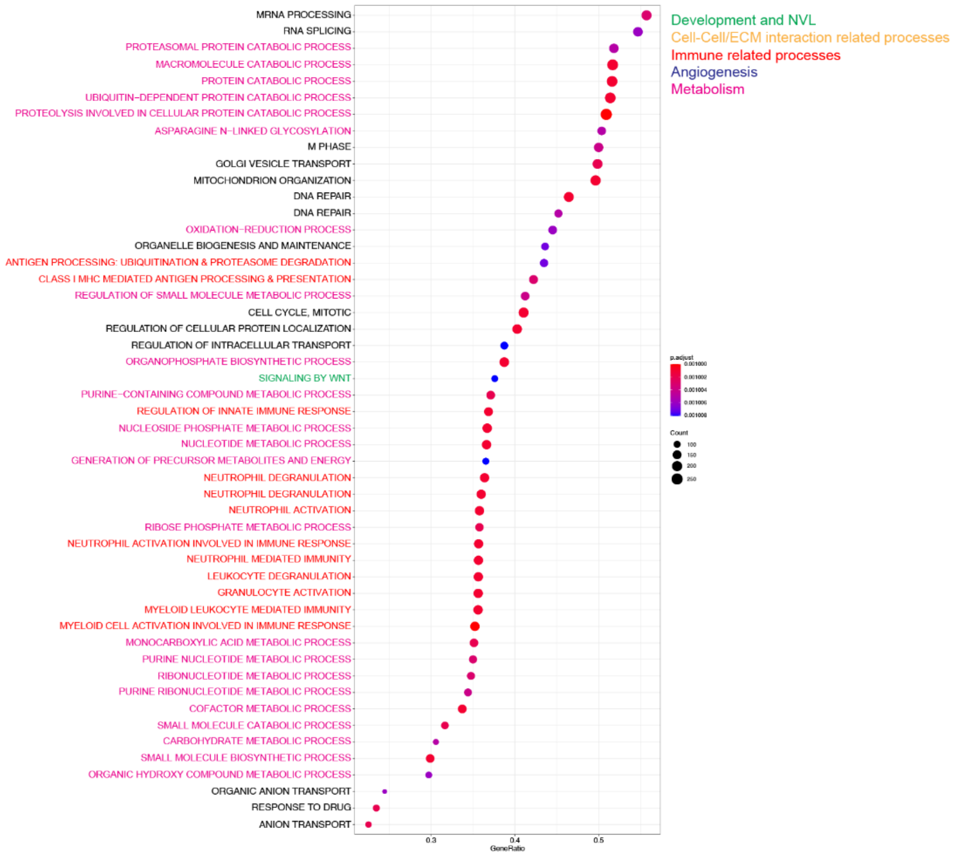

### Supplementary Figure 6

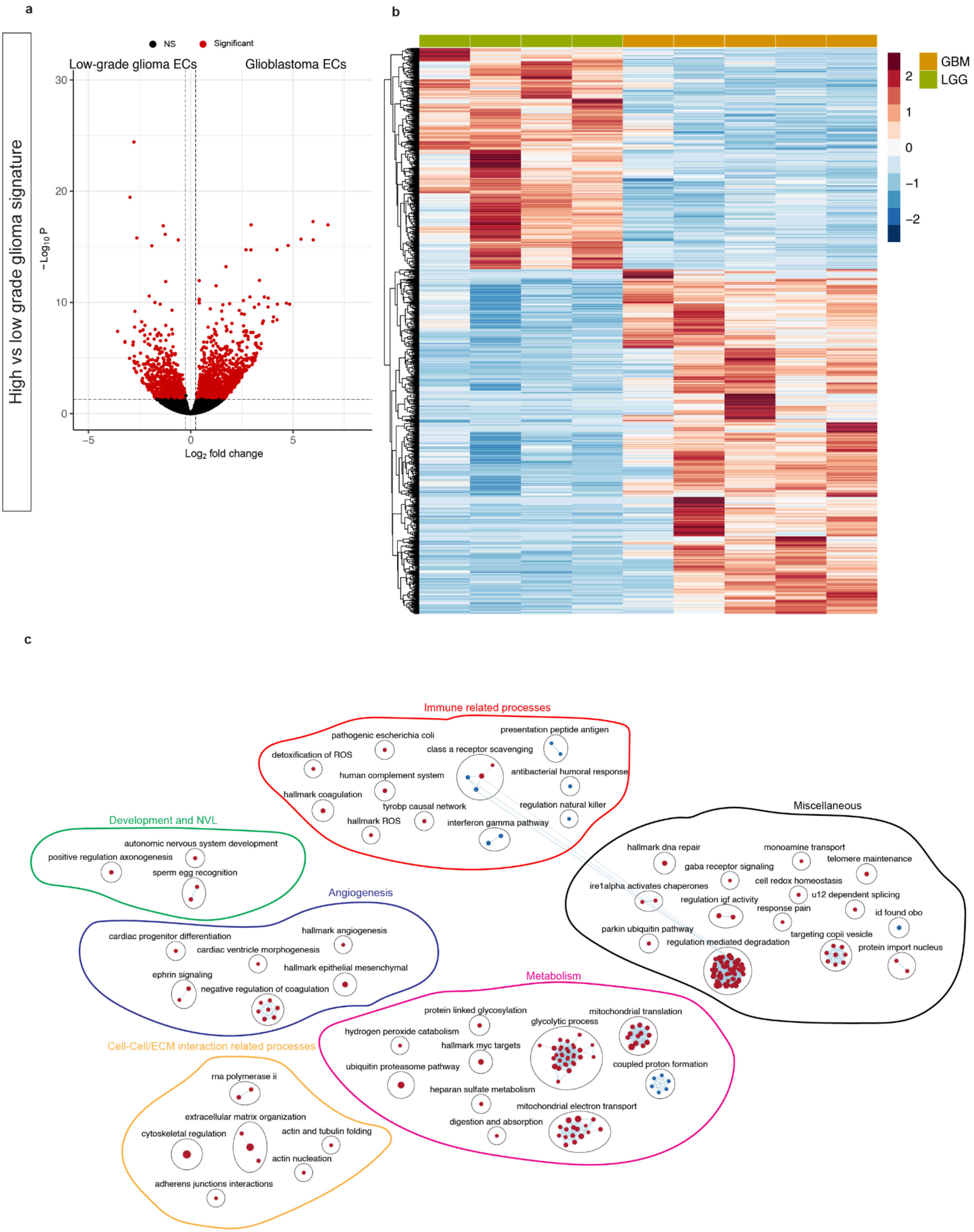

### Supplementary Figure 6

**d**

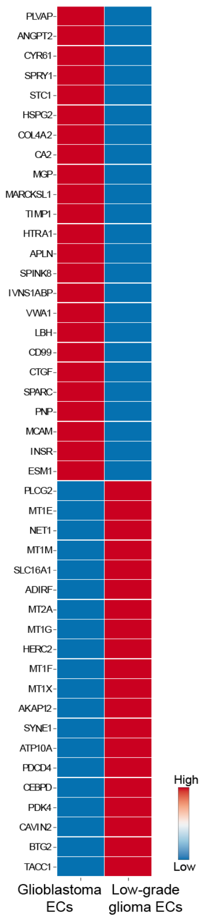


**e**

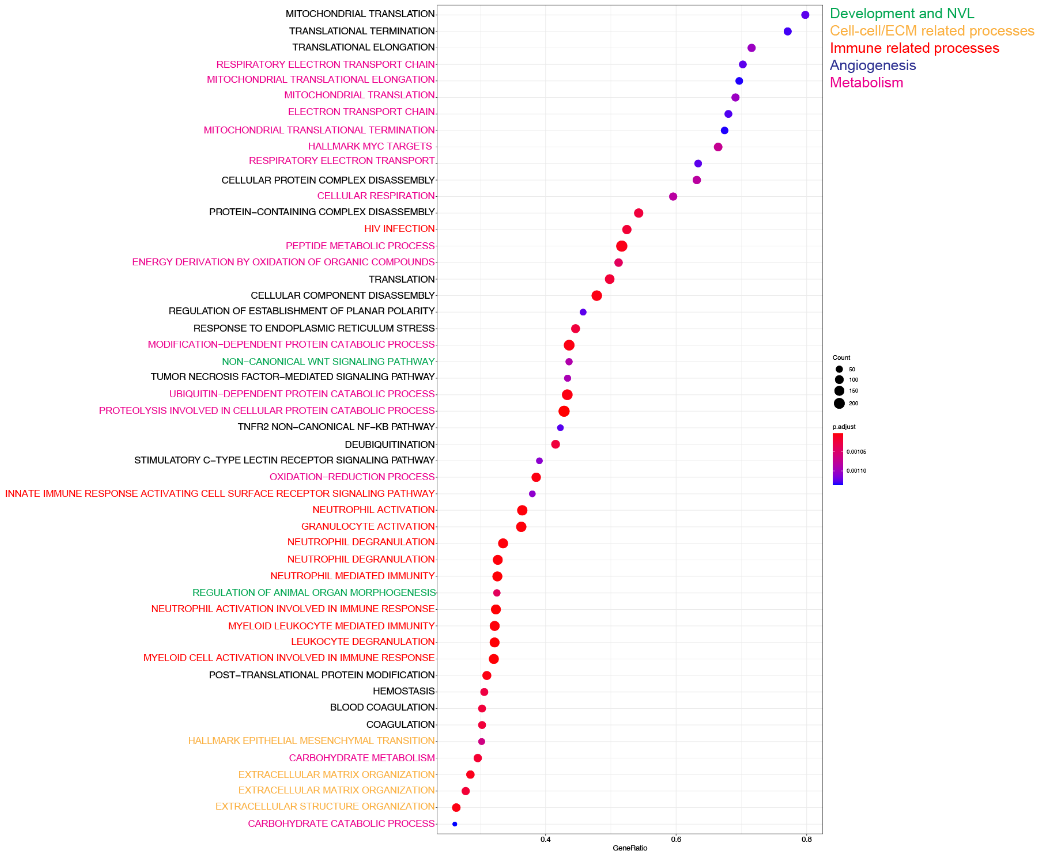

#### Supplementary Figure 7

**a**

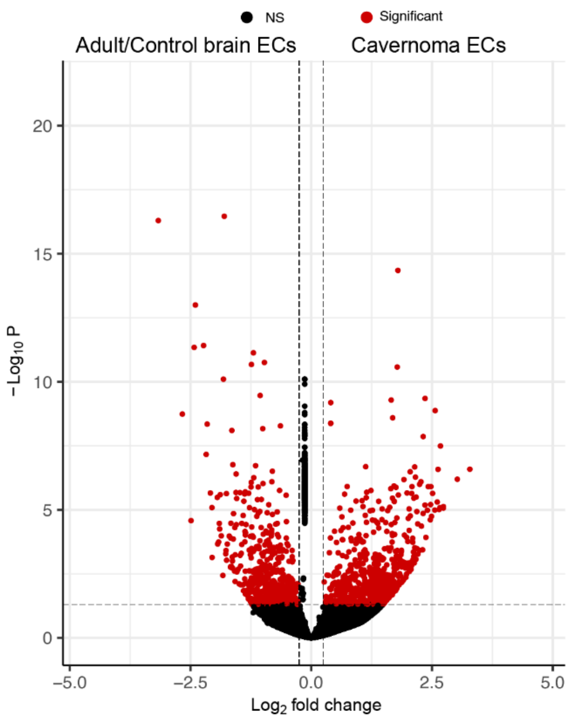


**b**

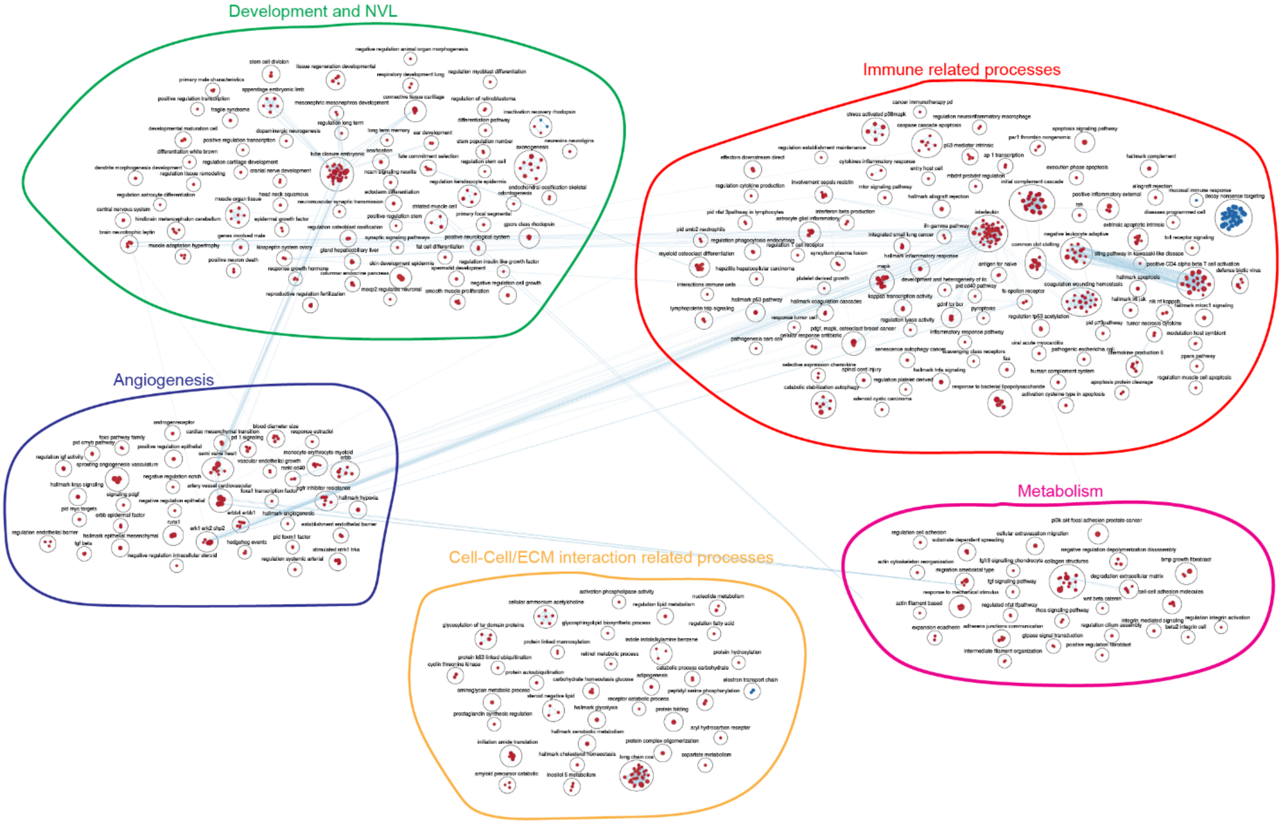

#### Supplementary Figure 8

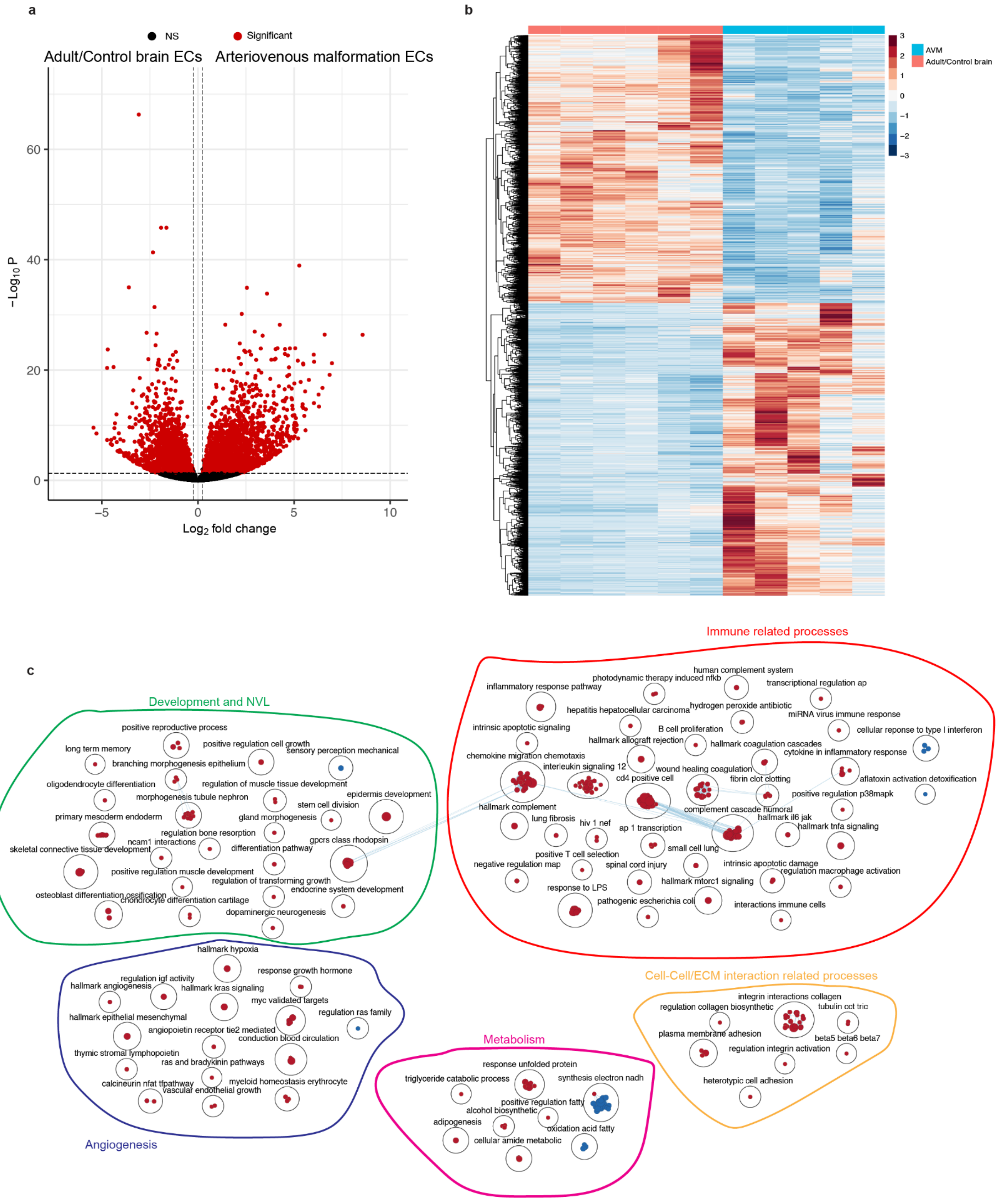

Supplementary Figure 9

a

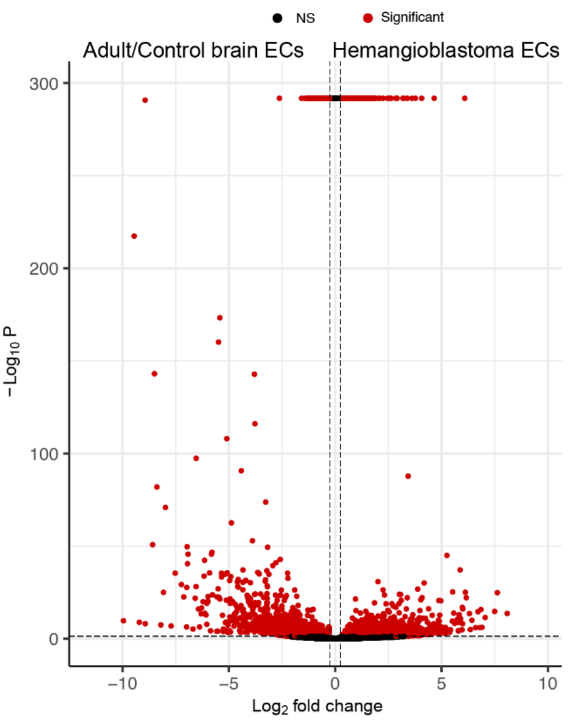


b

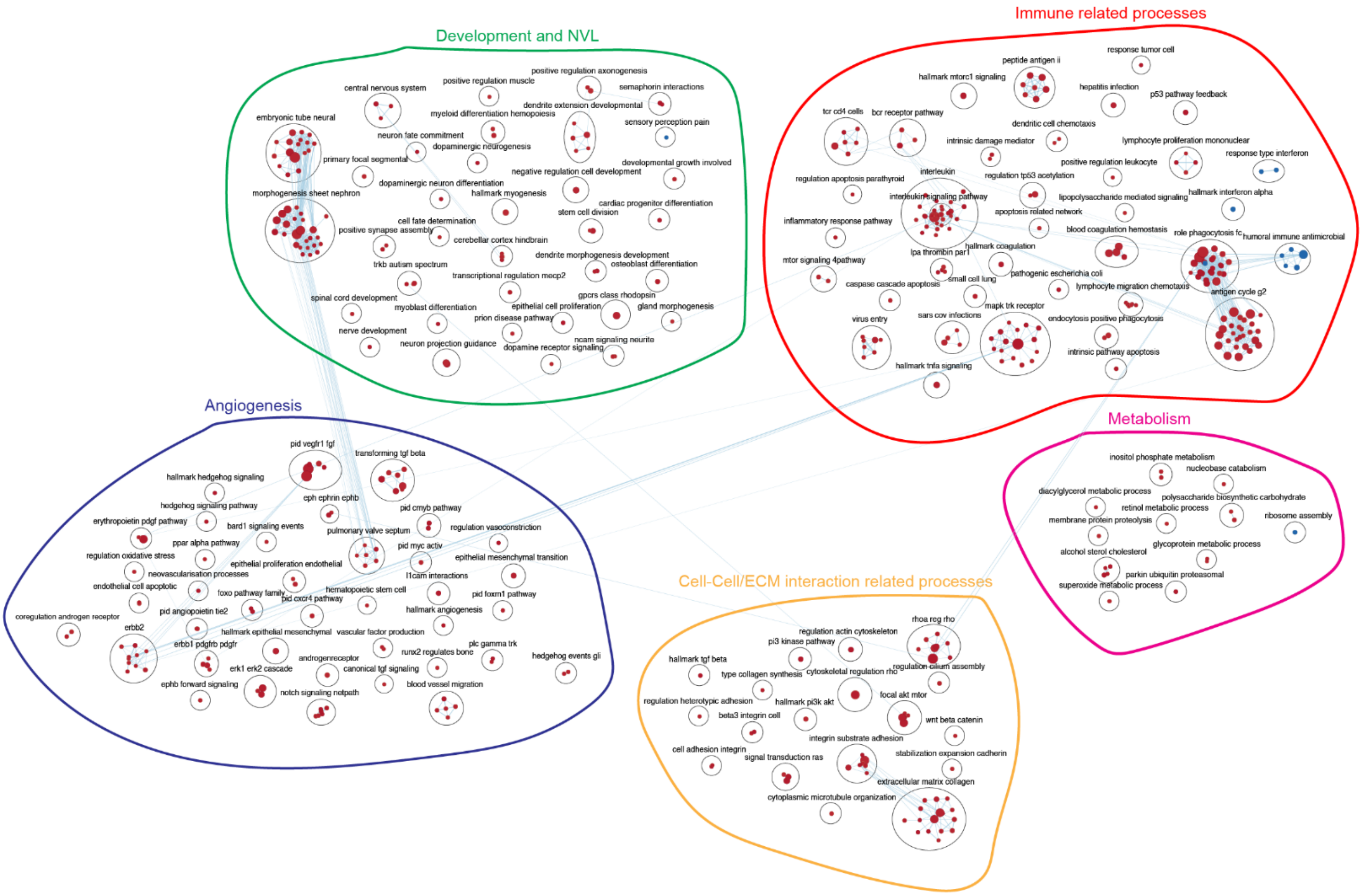

Supplementary Figure 10

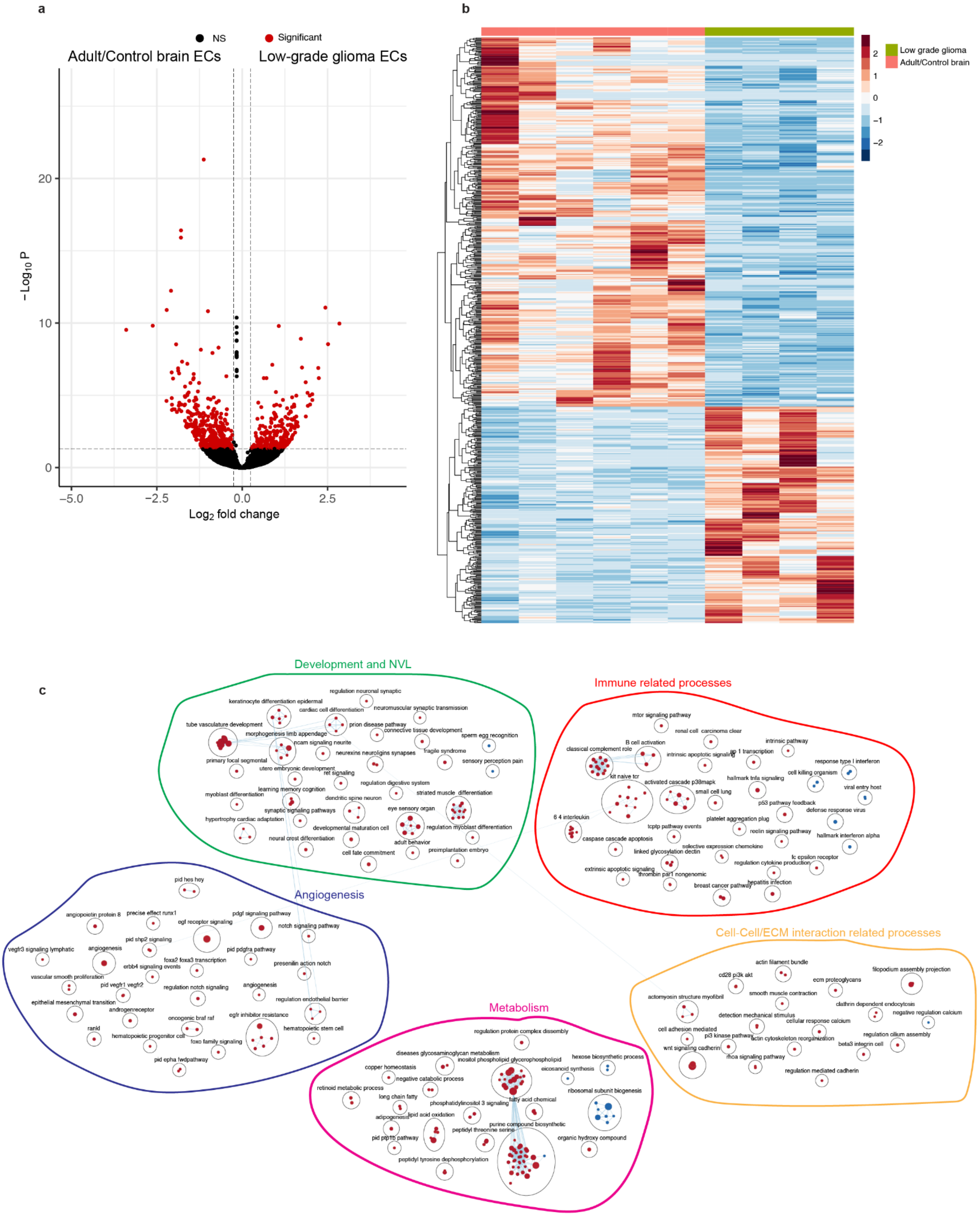

Supplementary Figure 11

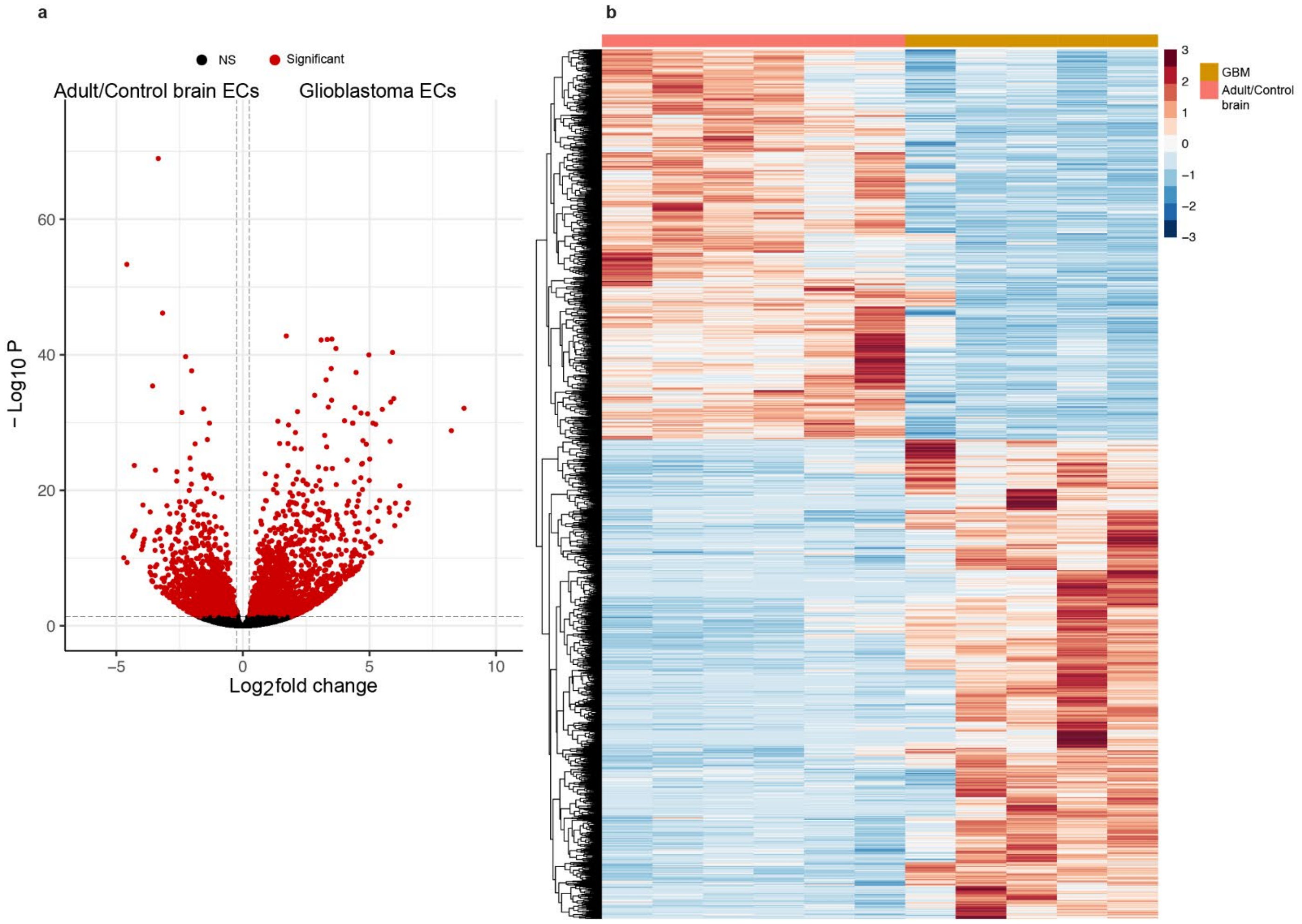

Supplementary Figure 12

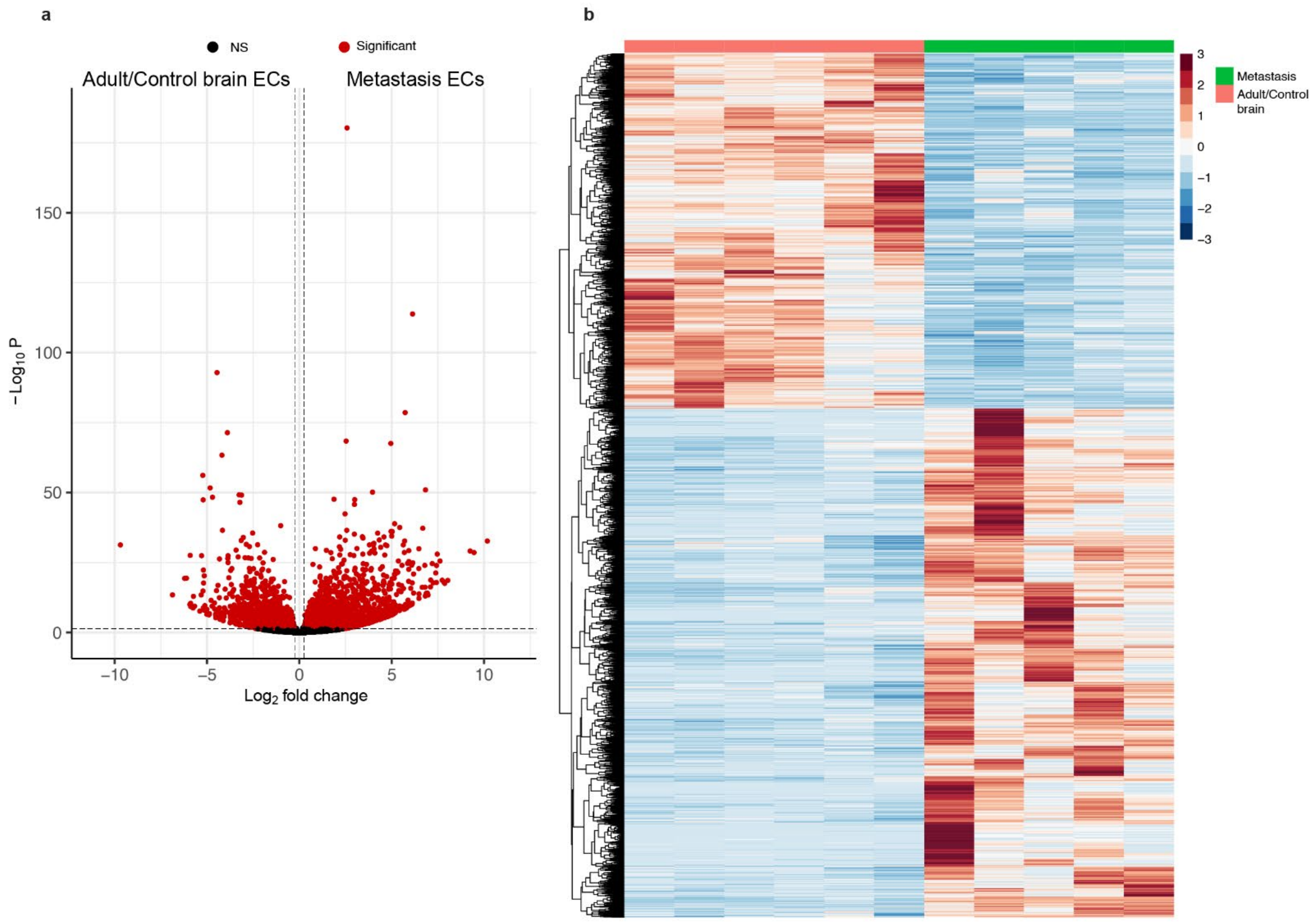

#### Supplementary Figure 13

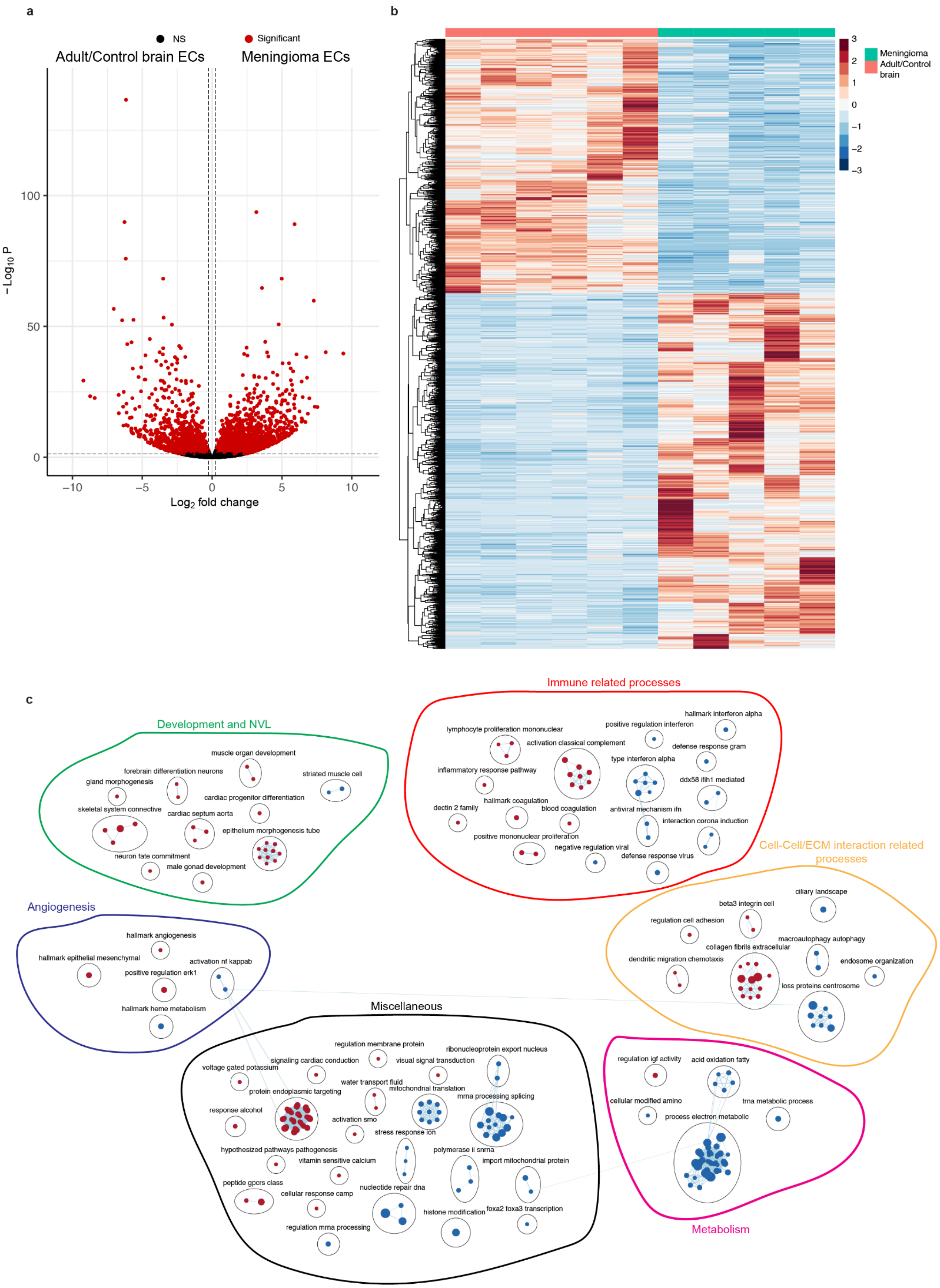

### Supplementary Figure 14

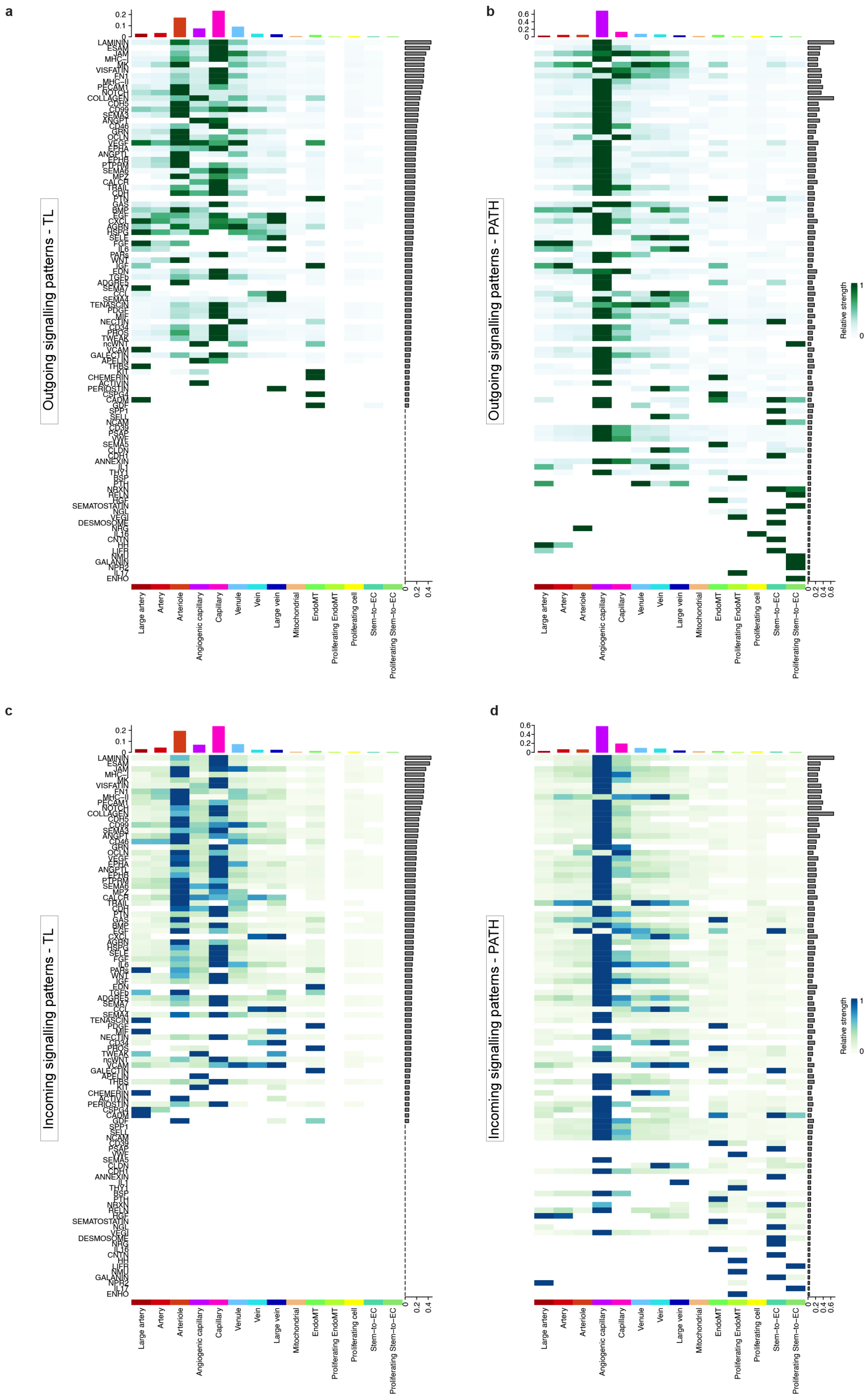

### Supplementary Figure 15

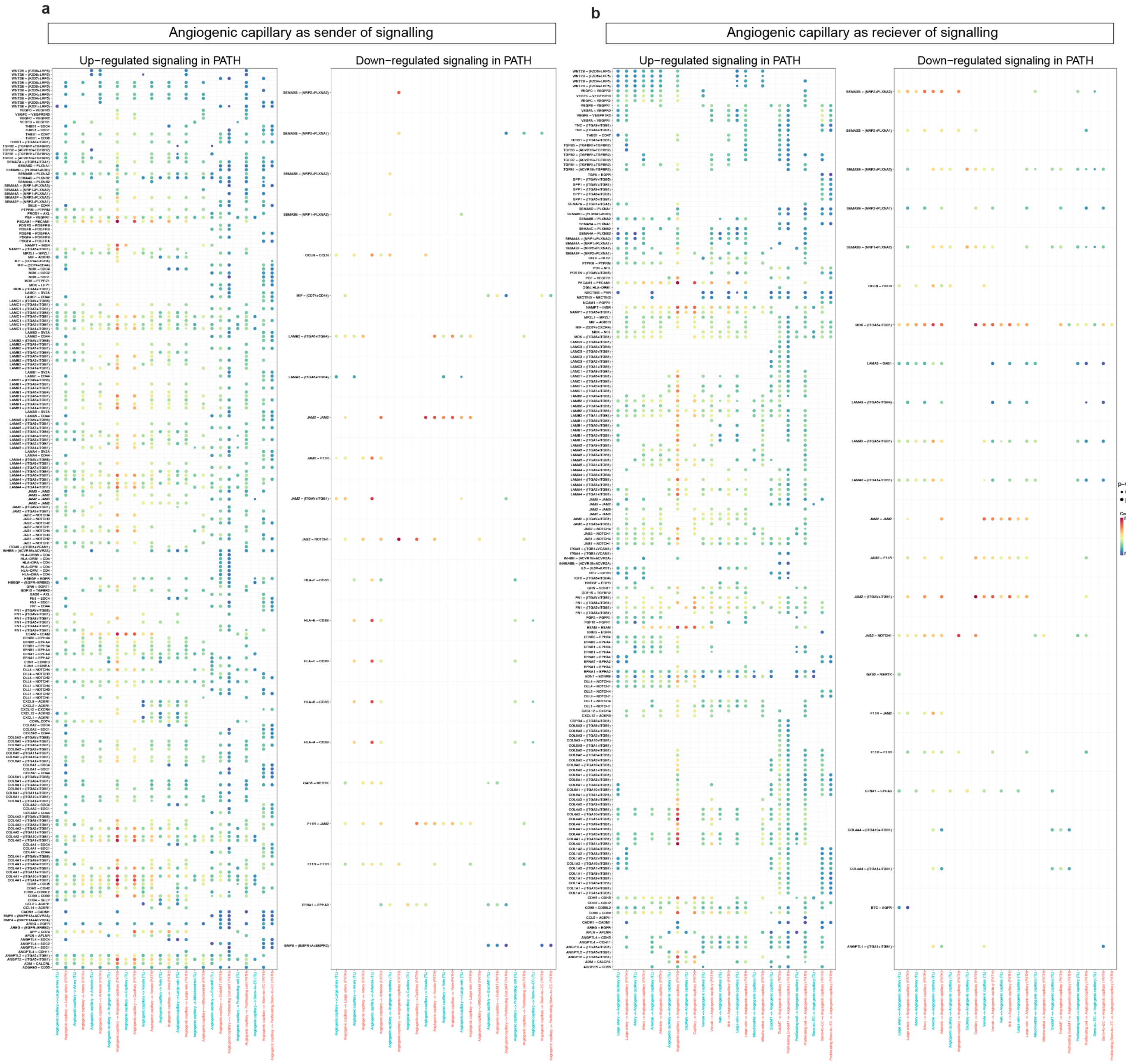

### Supplementary Figure 16

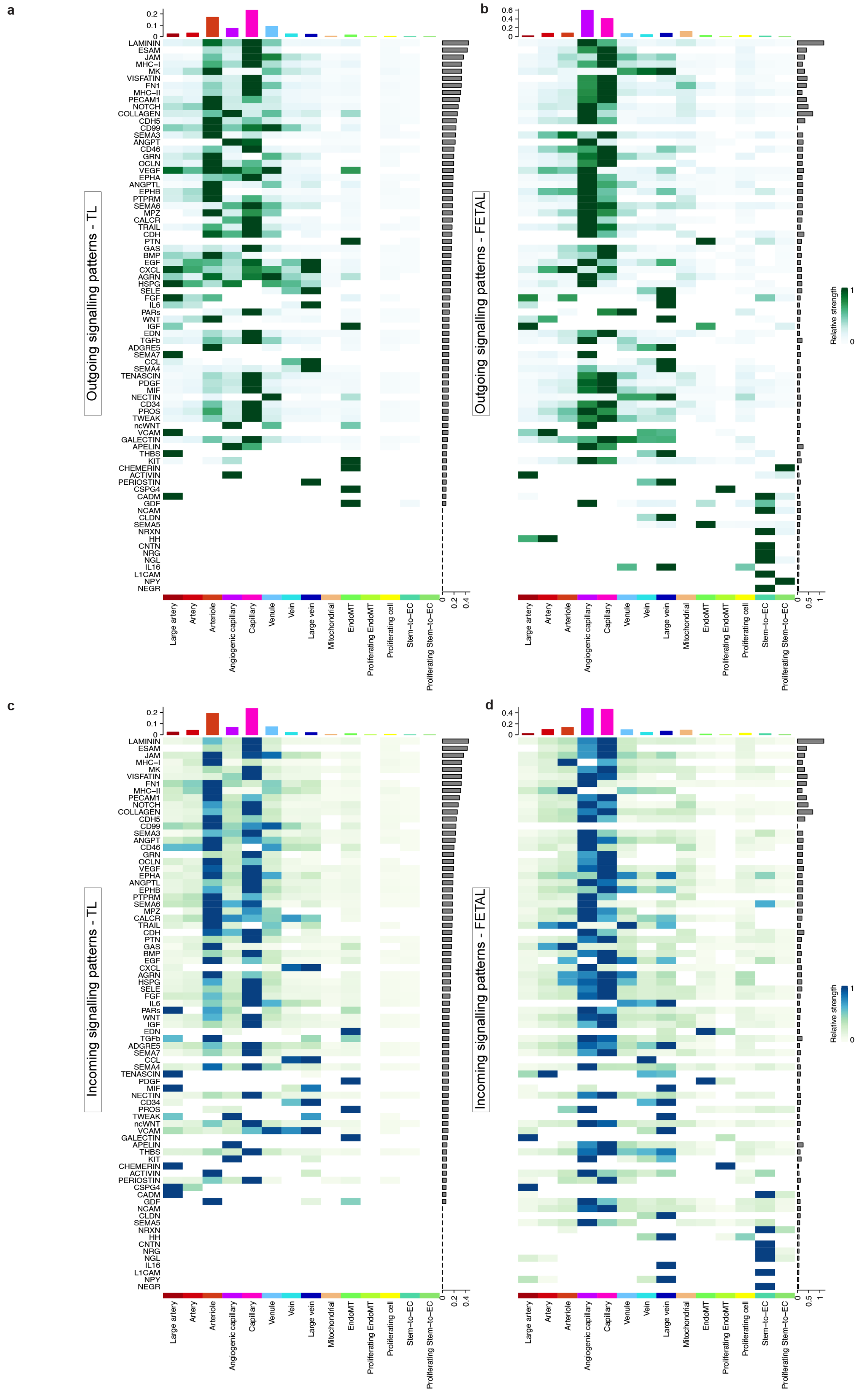

### Supplementary Figure 17

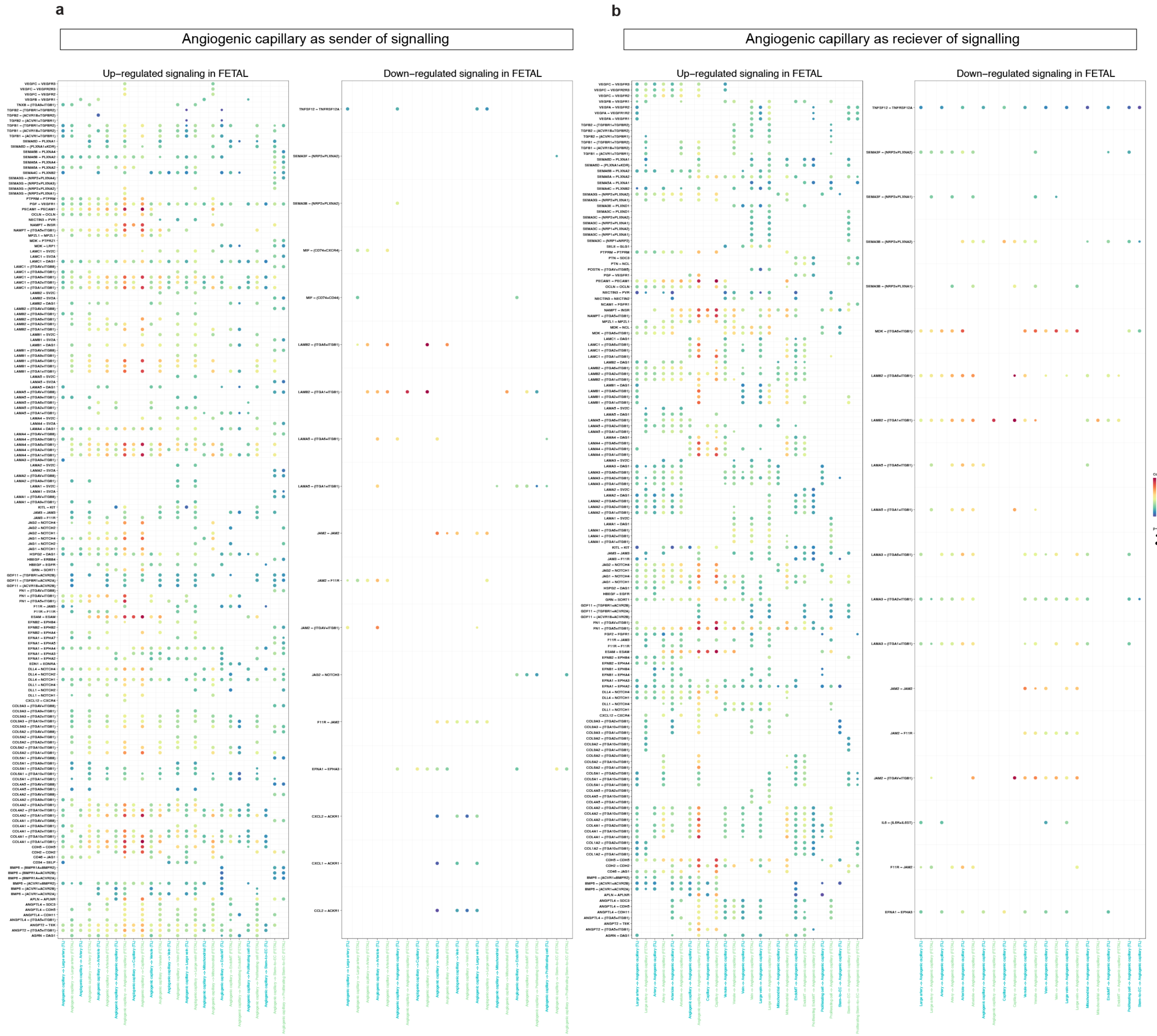

### Supplementary Figure 18

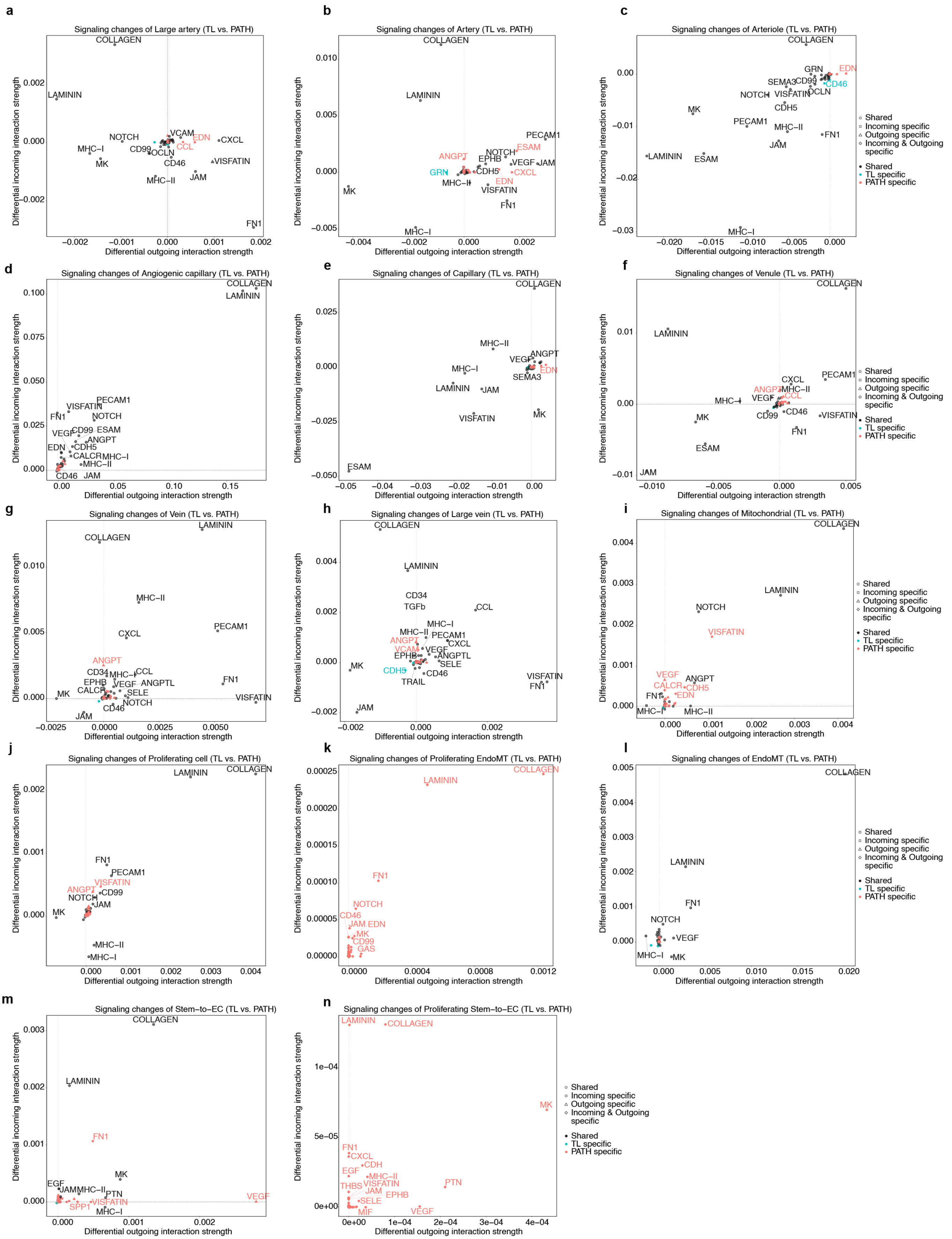

#### Supplementary Figure 19

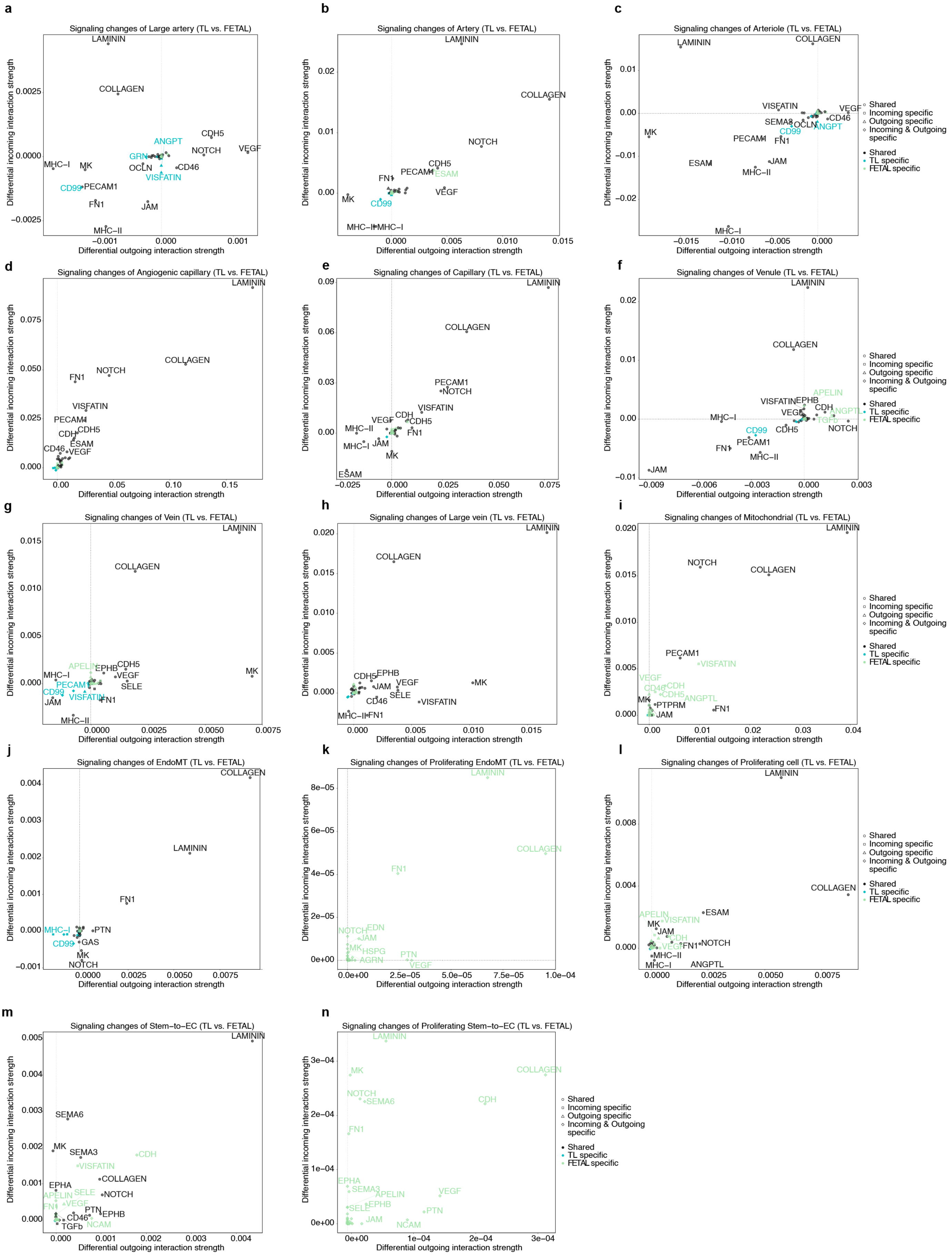

### Supplementary Figure 20

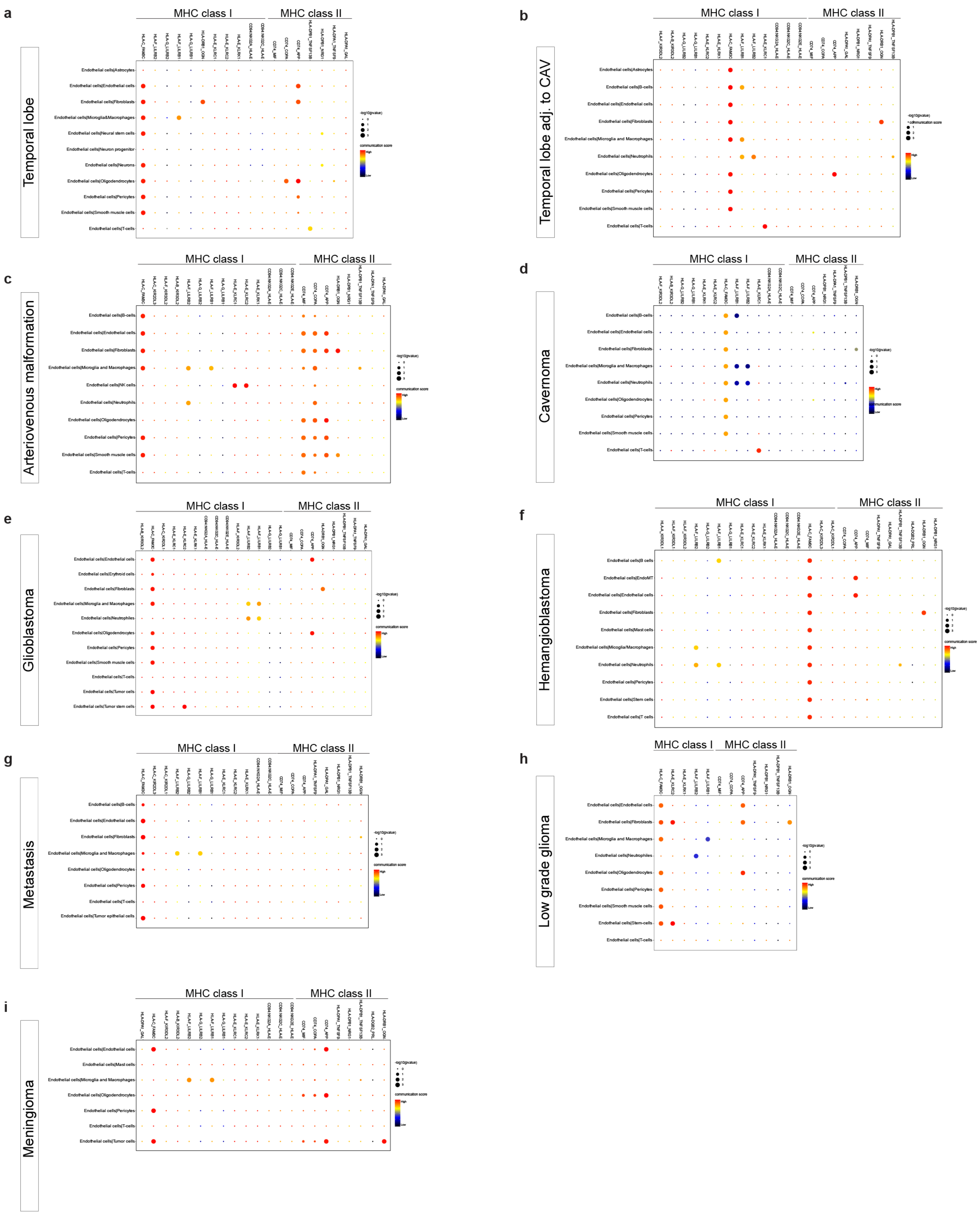

### Supplementary Figure 21

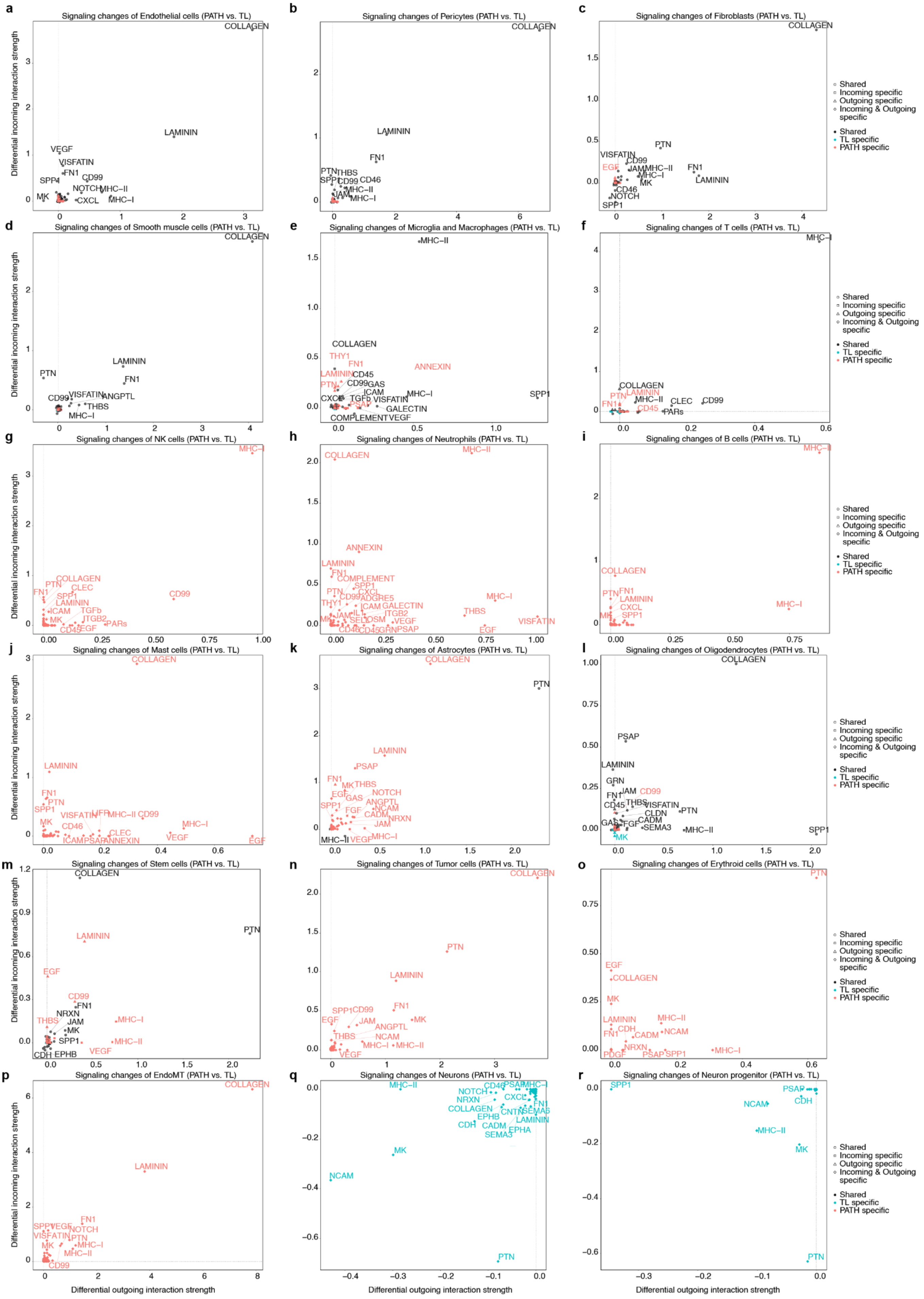

### Supplementary Figure 22

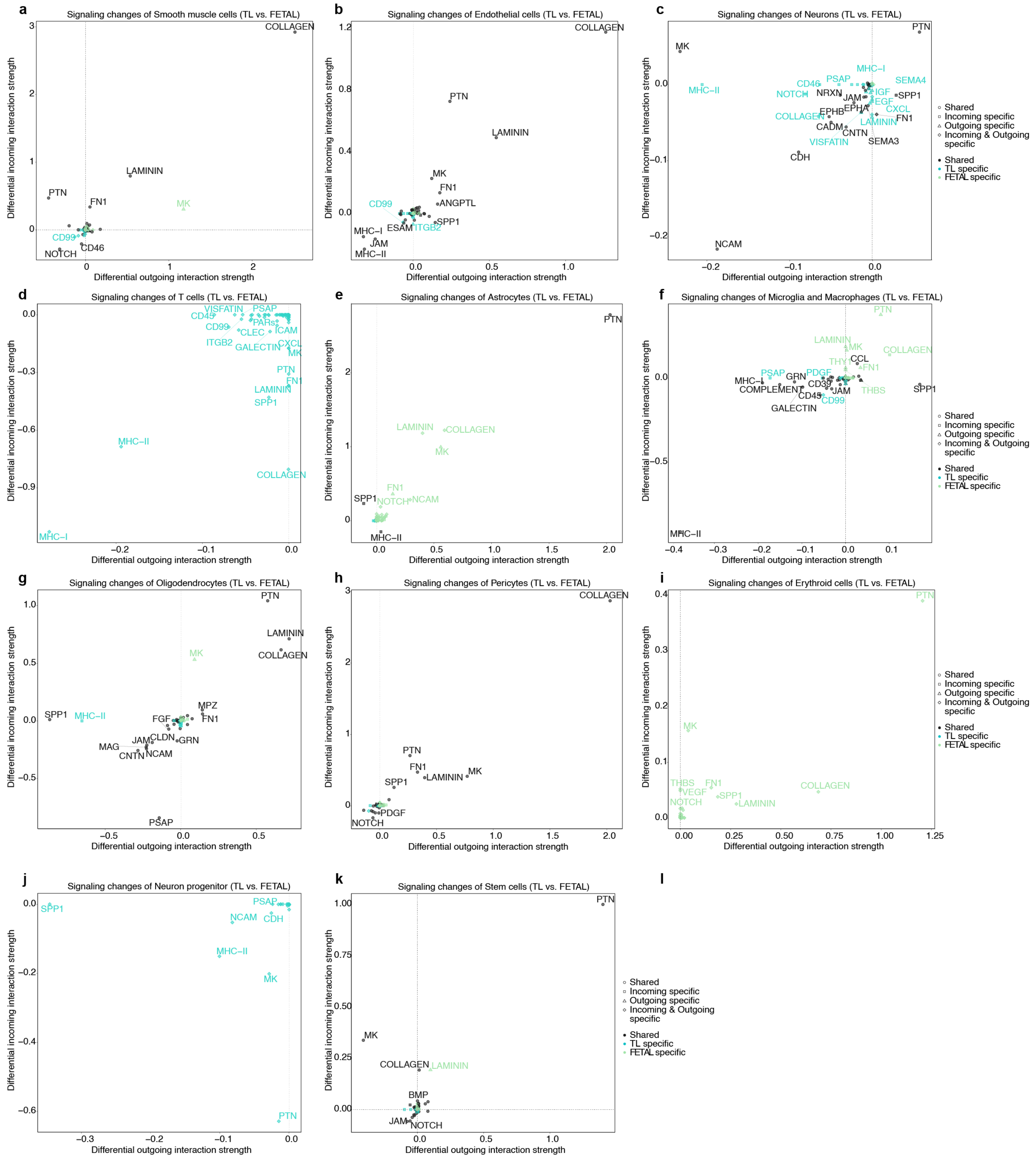

### Supplementary Figure 23

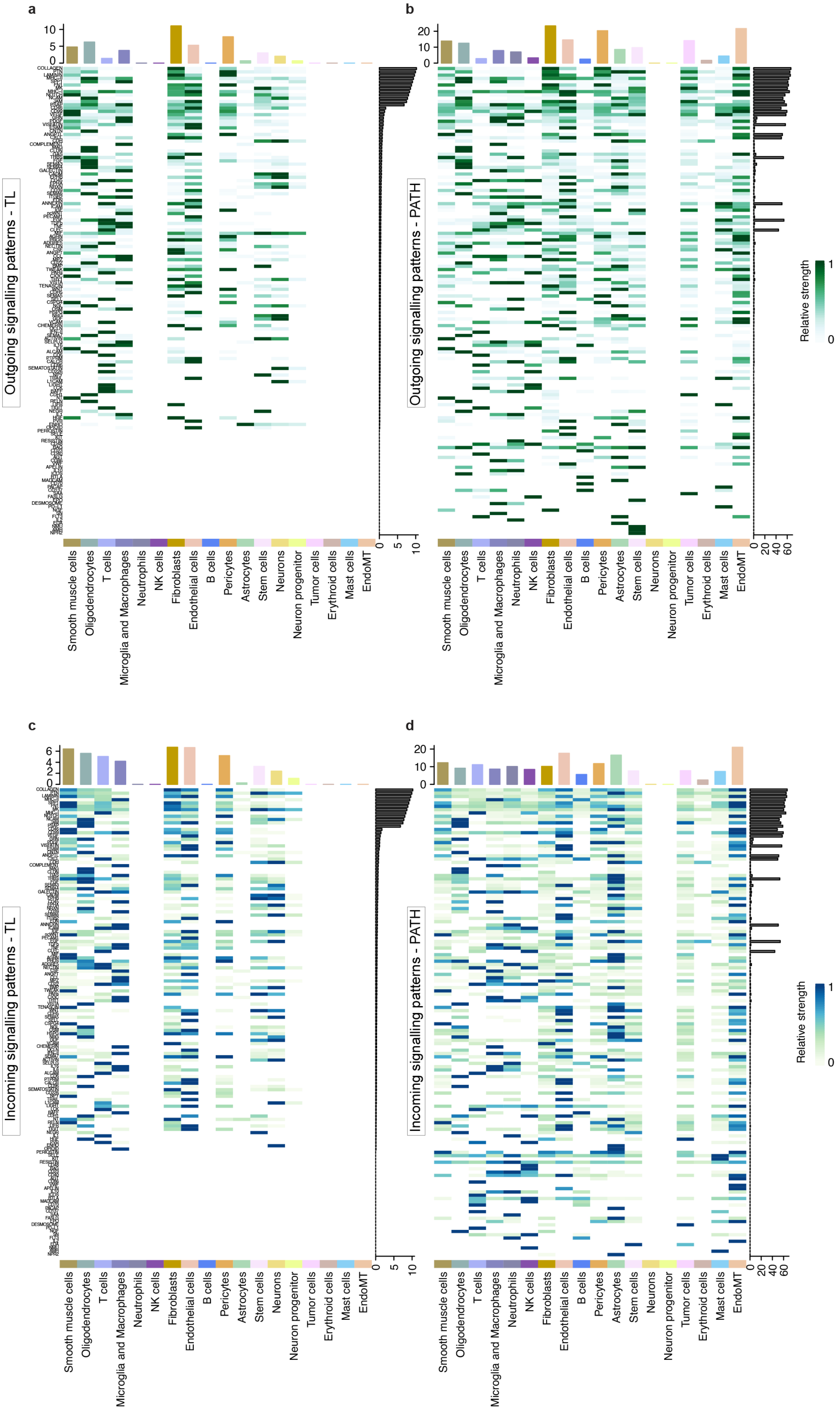

### Supplementary Figure 24

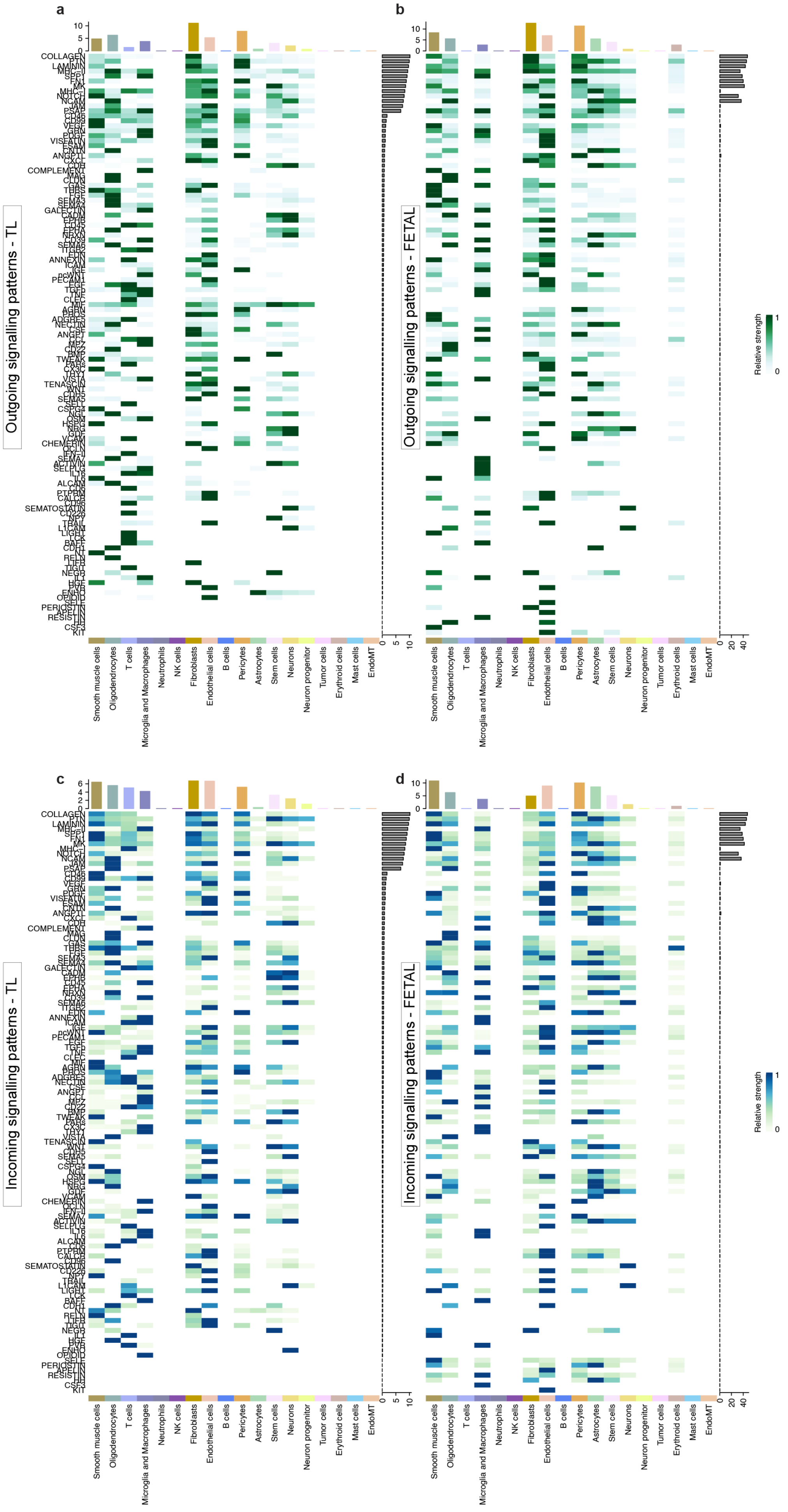

### Supplementary Figure 25

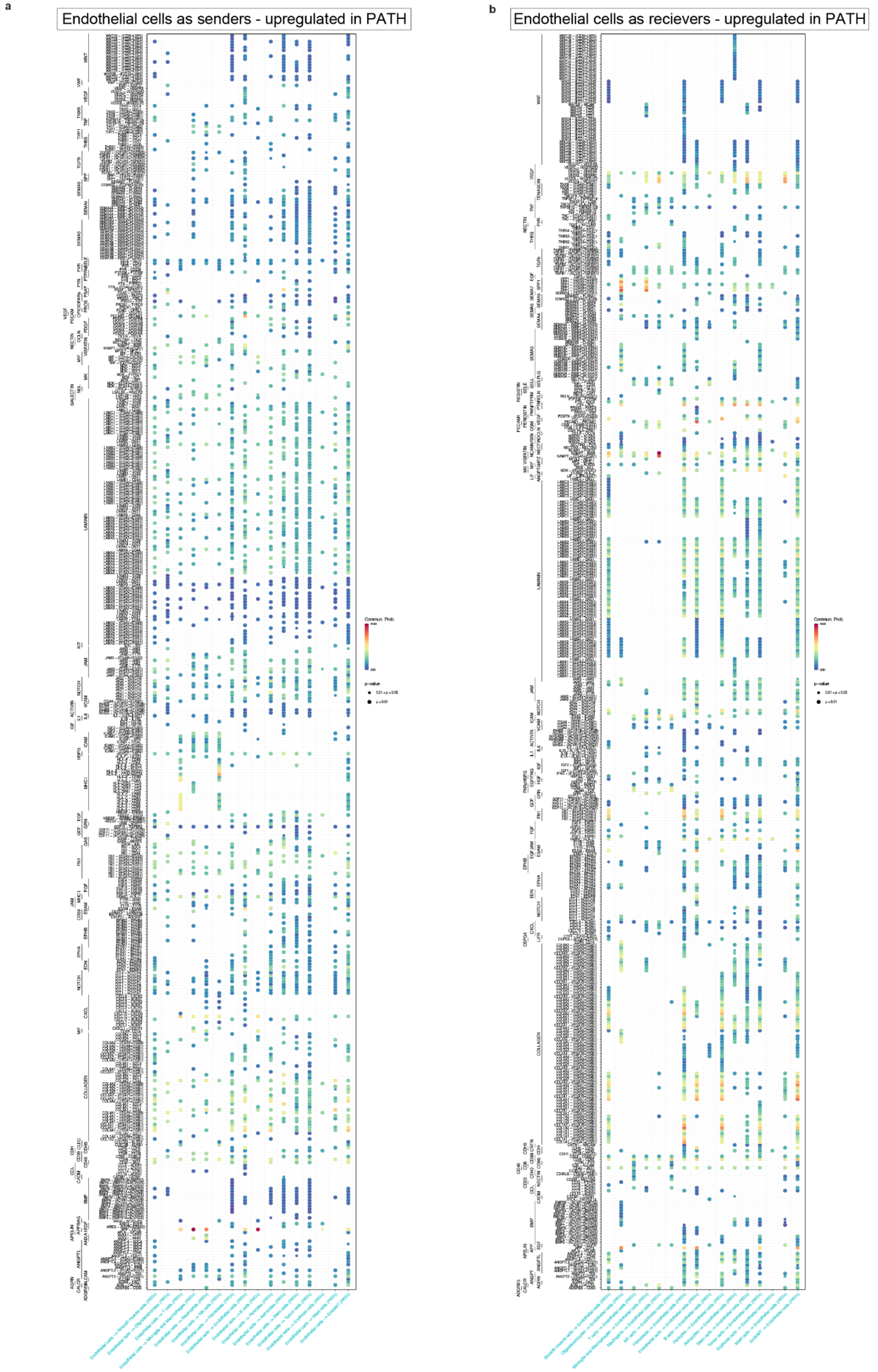

### Supplementary Figure 26

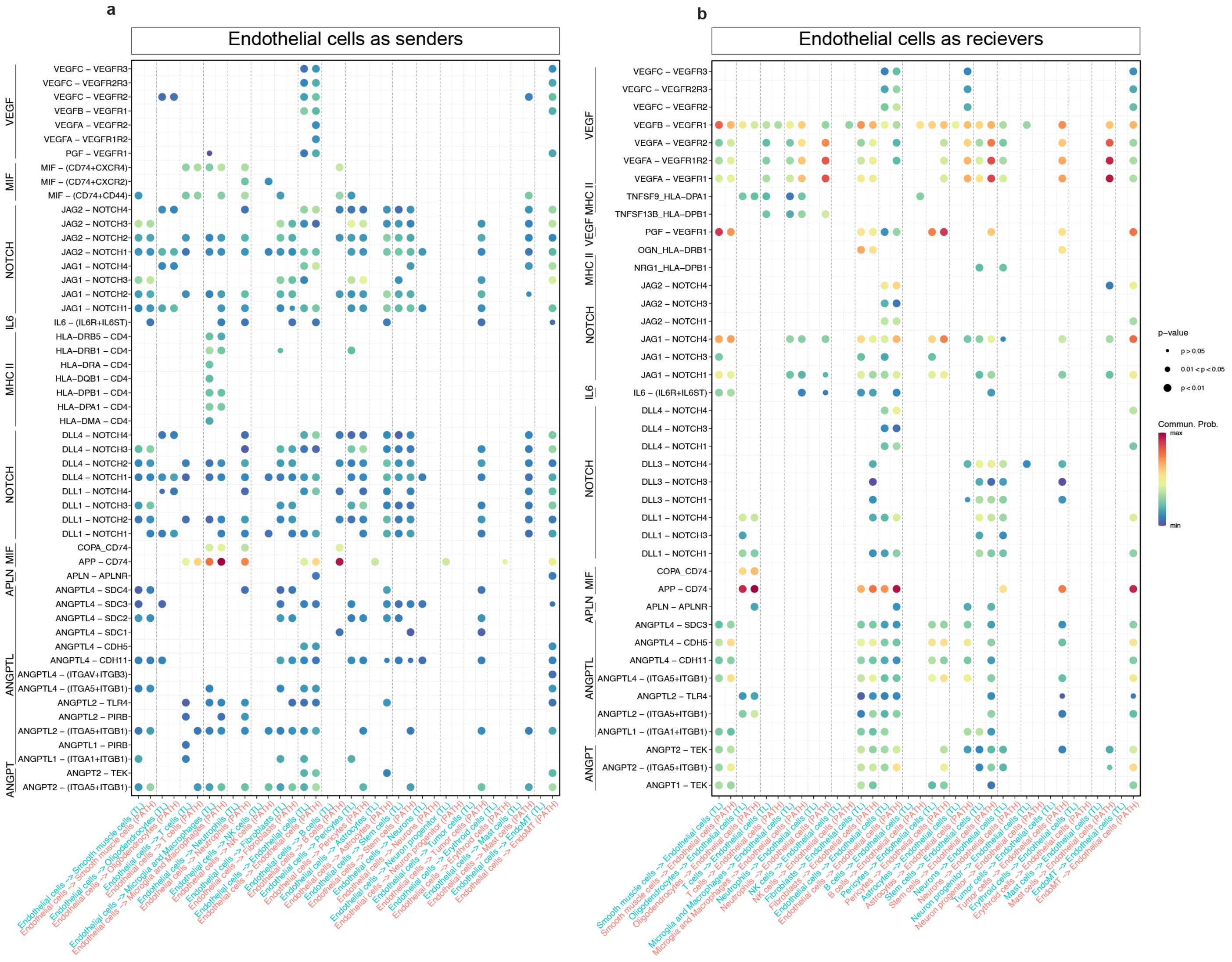

### Supplementary Figure 27

**a** Endothelial cells as sender - upregulated in FETAL

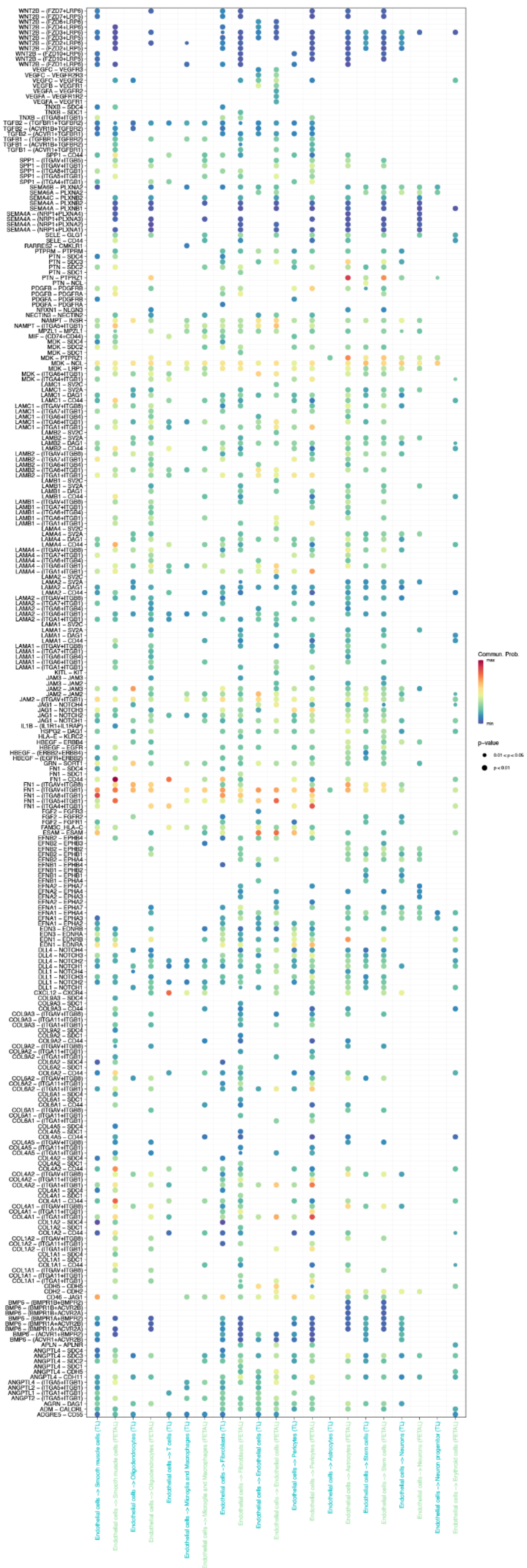

**b** Endothelial cells as sender - downregulated in FETAL

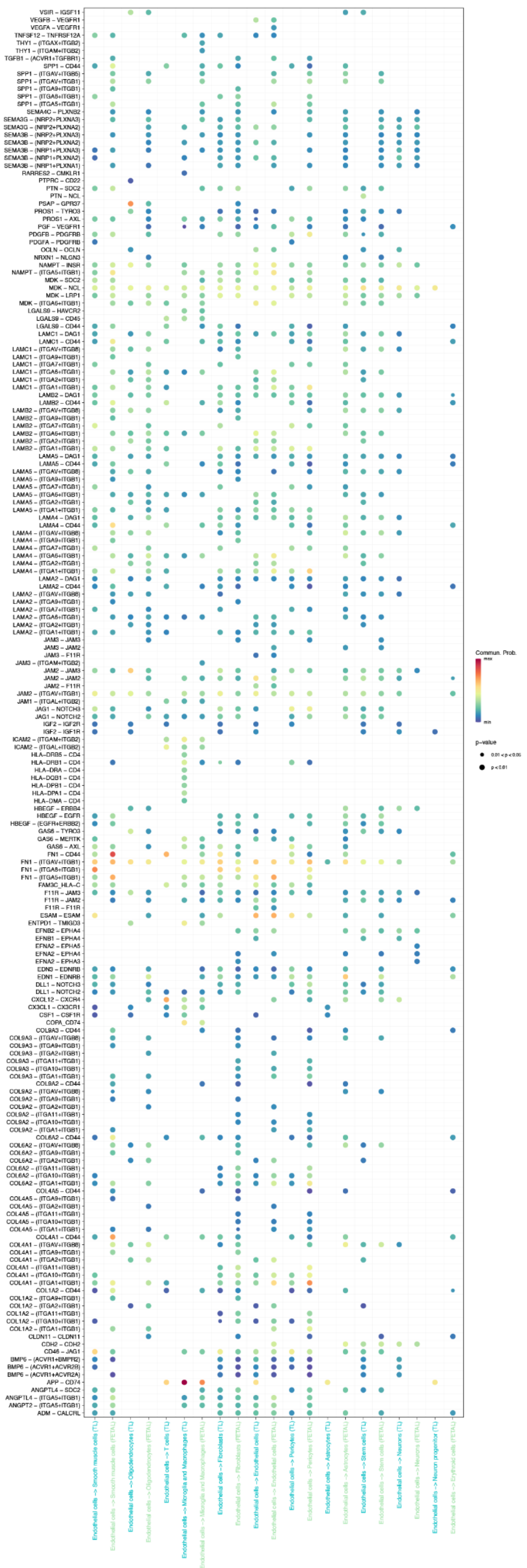

Supplementary Figure 28

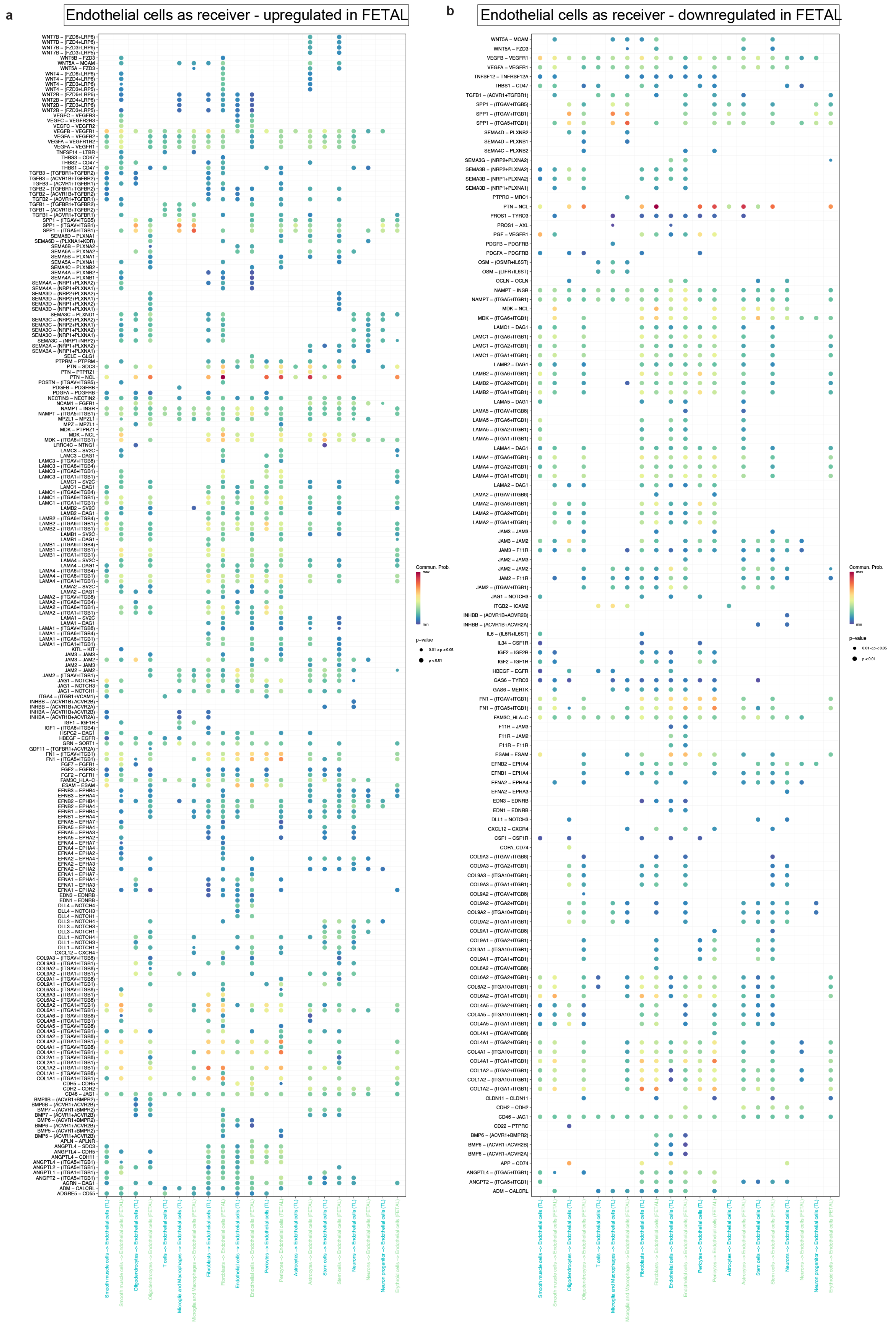
